# Estimating Divergence Times and Diversification Rates with the Unresolved Fossilized Birth-Death Process

**DOI:** 10.64898/2026.09.10.749957

**Authors:** Wonseop Lim, Levi Yoder Raskin, Jacky Kaiyuan Li, John P. Huelsenbeck, Rasmus Nielsen

**Affiliations:** Department of Integrative Biology, University of California, Berkeley, CA, USA; Biostatistics Division, University of California, Berkeley, CA, USA; Department of Statistics, University of California, Berkeley, CA, USA; Globe Institute, University of Copenhagen, Denmark

**Author notes:** Corresponding author: Wonseop Lim.

## Abstract

Tip-dating using Fossilized Birth-Death (FBD) models offers a favorable alternative to conventional node-dating analyses of the evolutionary timescale, bypassing various conceptual and technical difficulties. However, despite their increasing popularity in phylogenetic studies, their use has remained rather limited in taxonomic scope because of the difficulty in collecting morphological data, which is often required for tip-dating analyses, and computational intractability when a large number of fossils are present. Recent studies, however, suggest that FBD models can also be used in a principled way to provide prior information for molecular clock analyses, even in the absence of morphological data, relying only on temporal information and given clade assignments. Heath et al. (2014) introduced an “unresolved” FBD model in which uncertainty in fossil placements is analytically integrated out when morphological data are unavailable, providing a potential solution to this problem. Here, we show that the topological enumeration scheme used by Heath et al. (2014) to marginalize over unresolved fossil attachments overcounts configurations in which multiple fossil taxa can be resolved as a clade, and we provide a correction to this factor, yielding a valid marginalized unresolved likelihood. We apply our implementation to empirical datasets of Cetacea, Emydidae, and Palaeodictyopterida, and show that our method converges and yields reasonable estimates within a reasonable amount of time, even when hundreds of fossils are present.

## 1 Introduction

Reconstructing a dated phylogenetic tree is often the first step in understanding the evolutionary history of the clade of interest (dos Reis et al., 2016; Bromham et al., 2018). Given genetic or morphological data, one defines a model of evolution for sequence (e.g., DNA and amino acids) or morphological data. However, such models only resolve branch lengths in units of the expected number of substitutions per site, which equals the product of the substitution rate and the elapsed time. Thus, rate and time are always confounded: the same number of substitutions can arise from any arbitrary combination of rate and time. To make the model identifiable, one must therefore supply external prior information on divergence times. This external information can be derived from fossils (Benton & Donoghue, 2007; Parham et al., 2012; Wright et al., 2022) or, for measurably evolving organisms such as viruses, from samples with a known collection date (Drummond et al., 2003; Biek et al., 2015).

The earliest and most widely applied method for injecting external prior information on divergence times is to assign prior distributions to a selected set of internal nodes, an approach commonly referred to as “node dating” (Benton & Donoghue, 2007; Parham et al., 2012). In a Bayesian setting, one first identifies the node to be calibrated, gathers fossil specimens that can be assigned to the corresponding clade based on diagnostic morphological characters, together with their stratigraphic ages, and then specifies a prior distribution on the node age that reflects the uncertainty in this fossil evidence (Parham et al., 2012). In practice, however, defining a “reasonable” prior on a node age is extremely challenging, and poor choice of prior can yield biased divergence time estimates (Yang & Rannala, 2006; Hug & Roger, 2007; Ho & Phillips, 2009; Inoue et al., 2010; Lee & Skinner, 2011; Heads, 2012; Warnock et al., 2012; dos Reis & Yang, 2013; Warnock et al., 2015; Barba-Montoya et al., 2017; Brown & Smith, 2018), as there is no information in the sequence data to update prior calibration densities (Heath & Moore, 2014; Brown & Smith, 2018). Moreover, further complication of node dating can arise due to the unintended interaction between the specified calibration densities and the tree prior (Yang & Rannala, 2006; Inoue et al., 2010; Heled & Drummond, 2012; Rannala, 2016; Barba-Montoya et al., 2017). A line of research has sought to make calibration priors more objective, for example, by accounting for taphonomic processes (Marshall, 2008; Hedman, 2010; Claramunt & Cracraft, 2015; Warnock et al., 2017; Claramunt, 2022), diversification processes (Wilkinson et al., 2011; Nowak et al., 2013; Matschiner et al., 2017), or allowing calibrated nodes to change during analysis (Guindon, 2018). Even these approaches, however, can use only those fossils that can be confidently assigned to the crown node of the corresponding clade, thereby discarding a nontrivial portion of the paleontological record.

“Tip dating” (Pyron, 2011; Ronquist et al., 2012; Wright et al., 2022; Heckeberg et al., 2026) is an alternative to node dating that bypasses many of the problems described above: instead of specifying prior belief about divergence times to selected nodes, tip-dating approaches accommodate temporal information directly by incorporating heterochronously sampled tips into the tree. Its most widely used form relies on the fossilized birth-death (FBD) process (Stadler, 2010; Didier et al., 2012), which models the mechanistic joint process of speciation, extinction, and fossil sampling. The FBD process can be generalized by allowing the speciation, extinction, and sampling rates to vary through time (Stadler et al., 2013; Gavryushkina et al., 2014). Also, unlike models based on extant taxa only (Kubo & Iwasa, 1995; Louca & Pennell, 2020), it is statistically identifiable (Truman et al., 2025), provided that sampled ancestors (SA), sampled fossils that are direct ancestors of other sampled taxa (Foote, 1996; Stadler, 2010), are allowed, and that sampling does not affect edge survival.

In many cases, FBD models are applied to heterochronously sampled tips that carry associated sequence or morphological data. In a macroevolutionary context, however, non-extant tips seldom retain genetic data. Consequently, most studies have relied on morphological matrices scored from both extant and fossil taxa to apply the FBD process to date the tree. Yet morphological matrices are not always available, either because scoring characters is costly and time-consuming or because the morphological knowledge required to build a reliable matrix is simply lacking. Critiques of tip-dating have argued that the selective sampling of fossil taxa, often conducted due to the availability of morphological data or to include the oldest fossil representing the clade, can bias timescale estimates (Guindon, 2019; Matschiner, 2019), while simulation and empirical studies find that including complete fossil occurrences generally improves accuracy (O’Reilly & Donoghue, 2020; Luo et al., 2023; Nikolic et al., 2025). Thus, one would benefit from being able to include all available fossil record into the FBD process to date the phylogeny.

Recent studies have shown that the FBD process can supply a coherent, mechanistically motivated prior on node ages even when fossil tips carry no morphological data (Heath et al., 2014; Featherstone et al., 2021; Barido-Sottani et al., 2023; Nikolic et al., 2025; Capobianco et al., 2026). This coherence implies that, when the goal is simply to date divergences among extant species, including fossil taxa as data-free tips and letting the FBD process induce the joint node-age prior offers an effective alternative to morphology-based tip dating, which can sidestep the limitations described above. However, for the FBD process to induce a joint prior on the divergence times of extant taxa, it must marginalize over all possible attachments of the fossil tips. This can be highly inefficient when done using standard tree moves, and can become prohibitively slow when many fossil tips carry no morphological data, as the set of possible attachments to marginalize over then becomes large. Heath et al. (2014) sought to sidestep this by treating fossil attachments as “unresolved” and analytically integrating over their attachments to obtain the divergence-time prior.

Here, we show that the topological enumeration used by Heath et al. (2014) to construct the unresolved FBD likelihood overcounts the number of configurations by which multiple fossil taxa can join a clade, and that this overcounting induces bias that increases with the number of fossil tips. We derive the correct enumeration for the quantity and re-implement the model, coupling it with an approximate dating approach (Thorne et al., 1998; Thorne & Kishino, 2002; Seo et al., 2004; Guindon, 2010; dos Reis & Yang, 2011) for coherent and accelerated phylogenetic dating under the unresolved FBD prior. We also show that using this correct enumeration, one can estimate speciation, extinction, and sampling rate trajectories only from the fossil records and number of extant species under a bifurcating speciation model, similar to what has been implemented by Warnock et al. (2020) and Stolz (2024) using the budding speciation model (Stadler et al., 2018), and work of Didier et al. (2017), Didier and Laurin (2020), and Didier and Laurin (2024) but without or only partially requiring a prespecified phylogeny of taxa.

## 2 Theory

### 2.1 Fossilized Birth-Death Process

Here, we introduce the basic formulation of the FBD process (Stadler, 2010; Didier et al., 2012). We first write likelihoods in terms of oriented trees (Gernhard, 2008; Ford et al., 2009), which are objects obtained by distinguishing branches upon a branching event with left or right labels. Converting the likelihood of oriented trees to labeled trees can be easily done by multiplying by a conversion factor (see below).

#### Skyline FBD process

We assume a skyline FBD model (Stadler et al., 2013; Gavryushkina et al., 2014), where the process starts at the origin *t*_*or*_ = *x*_0_ with a single taxon. We partition time into bins, and for each bin [*t*_*i*_, *t*_*i*−1_) (*i* = 1, …, *l*, where *t*_0_ = ∞, and *t*_*l*_ = 0 is present), define speciation, extinction, and preservation rates as *λ*_*i*_, *µ*_*i*_, and *ψ*_*i*_. Generalizing the model to allow each rate to have different time bins is trivial. We also define the probability of extant taxa being sampled (via Bernoulli sampling) as *ρ*. Following Stadler (2010) and Stadler et al. (2013), we can write the master equation for *p*_*i*_(*t*): the probability that an edge at time *t* ∈ [*t*_*i*_, *t*_*i*−1_) has no sampled extinct or extant descendants, as follows:

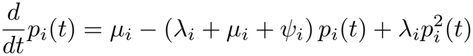

which gives a unique solution

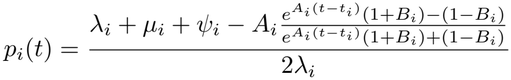

where

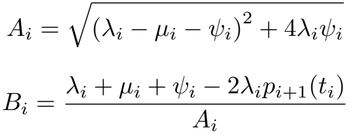

and *p*_*l*+1_(*t*_*l*_) = 1 − *ρ*. Now define *g*_*i,e*_(*t*) as the probability density that an edge in the tree at time *t* ∈ [*t*_*i*_, *t*_*i*−1_) corresponding to edge *e* evolved between *t* and the present (*t*_*l*_ = 0) as observed in the tree. Then the master equation for *g*_*i,e*_(*t*) is given as

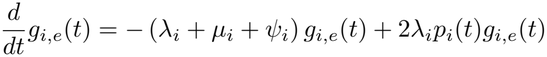

with, at the end of edge *e* at time *t*_*e*_:

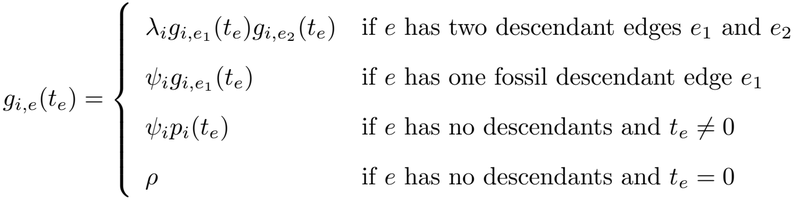

The solution to this equation, for *t* and *t*_*e*_ in the same bin *i*, is

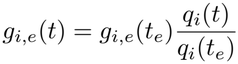

where

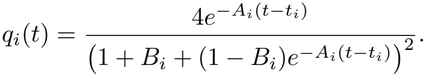

Since *q*_*i*_(*t*_*i*_) = 1, an edge spanning several bins accumulates these factors across bins. Writing 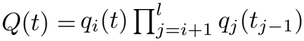 for *t* in bin *i*, we have 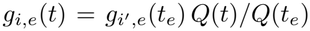 for *t* in bin *i* and *t*_*e*_ in bin *i*^′^ ≥ *i*.

The probability density of an observed oriented tree *T*_*or*_ is then enumerated by recursively using *g*_*i,e*_(*t*) from the time of origin *x*_0_ to the present, which gives

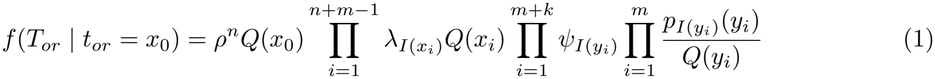

where *n* is the number of extant species, *m* is number of sampled fossil tips, *k* is number of sampled ancestors, *x*_*i*_ is the divergence times, *y*_*i*_ is the age of fossils, and *I* is a function where *I*(*x*) = *i* if *x* ∈ [*t*_*i*_, *t*_*i*−1_). The likelihood for the labeled tree, *T*_*lb*_, can be obtained by applying a conversion factor (Gernhard, 2008; Stadler, 2010): First, consider all possible orientations (factor 2^*n*+*m*−1^), and label the tips, assuming that labels are exchangeable within extant taxa and within fossil taxa (note that the likelihood is identical up to constant proportionality even when exchangeability is assumed between all taxa). Thus,

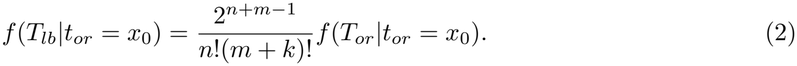

Note that *n* and *m* + *k* are invariant, and only *m* and *k* can change due to changes in the sampled ancestors (SA) configuration during the Markov chain Monte Carlo (MCMC) (Gavryushkina et al., 2014; Heath et al., 2014). So it can be effectively written *f* (*T*_*lb*_|*t*_*or*_ = *x*_0_) ∝ 2^*m*^*f* (*T*_*or*_|*t*_*or*_ = *x*_0_).

#### Conditioning

One often needs to condition the FBD process on specific events (Stadler, 2013). Stadler (2010) derived conditioning of the FBD process on survival event *S*, i.e., starting at the origin *x*_0_, the process leaves at least one sampled extant taxa. Given a tree *T* (oriented or labeled), define 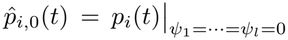, a probability that an edge at time *t* ∈ [*t*_*i*_, *t*_*i*−1_) has no sampled extant descendants; i.e., it is a solution to the master equation of *p*_*i*_(*t*) with *ψ*_*i*_ = 0, with the value 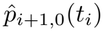 at *t*_*i*_ and 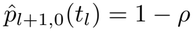 at the present. Then,

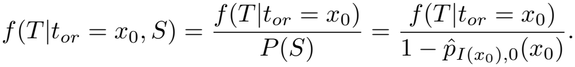

Stadler (2010) also derived an FBD process conditioned on the crown age, *x*_1_, and a corresponding density for conditioning on the event *S*. Apart from survival, the minimum conditioning one can do is the event of sampling at least one of any taxa: *A*. Since 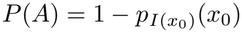,

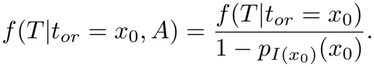

One may also condition on the event of having no extant samples, *E*, which is equivalent to extinction when *ρ* = 1. This equals 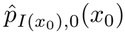, but for the process to be observable, we need at least some taxa sampled. Thus, we can write 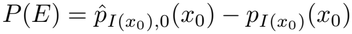, and therefore

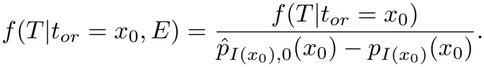

One can also condition on the number of extant taxa (Stadler, 2010, 2013), with conditioning on extinction viewed as a special case; however, this comes with some important caveats. Under the constant-rate birth-death process, it is known that swapping *λ* and *µ* changes the probability that the number of lineages goes from *N* to *M* only by the factor (*λ/µ*)^*N*−*M*^ (Waugh, 1958; Tavaré, 2018). Under *ρ* = 1, the density of a birth-death tree without fossil sampling conditioned on the number of extant tips is therefore the same when we swap *λ* and *µ* (Stadler & Steel, 2019). For the constant-rate FBD process with *ρ* = 1, it is not difficult to check that swapping *λ* and *µ* leaves *A*_1_ and *B*_1_ unchanged, and since *q*_1_, and thus *Q*, depend on *λ* and *µ* only through *A*_1_ and *B*_1_, they are unaffected by the swap. The numerator of *p*_1_ is likewise unchanged, while its denominator 2*λ* becomes 2*µ*, so the swap multiplies *p*_1_ by *λ/µ*. In Equation (1), the swap therefore turns each of the *n* + *m* − 1 factors of *λ* into *µ* and multiplies each of the *m* factors *p*_1_(*y*_*i*_) by *λ/µ*, so the swapped density equals the original density multiplied by (*µ/λ*)^*n*+*m*−1^(*λ/µ*)^*m*^ = (*λ/µ*)^1−*n*^.

When conditioning on *S*_*n*_, the event of having *n* extant species, we divide the density by *P* (*S*_*n*_) (Stadler, 2010, Theorem 3.3 & Corollary 3.6), which integrates the density over all trees with *n* extant tips, each of which is multiplied by the same factor (*λ/µ*)^1−*n*^ (see also Rannala (1997) and Stadler and Steel (2019)). Thus, conditioning on *S*_*n*_ leaves the constant-rate FBD density invariant under the swap. Extinction conditioning *E* is the subcase of *S*_0_ with at least one fossil sampled. With *ρ* = 1, having no extant sample means that no lineage survives to the present, so a tree under *E* has *n* = 0 and its density is multiplied by *λ/µ*. The probability 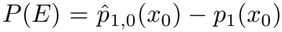 is multiplied by *λ/µ* as well, since 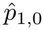 scales in the same way as *p*_1_, so the density conditioned on *E* is also invariant under the swap.

Under the general time-dependent FBD model with *ρ* = 1, we show in the Appendix (Theorem 3) that, for rates *λ*(*t*), *µ*(*t*), and *ψ*(*t*), the density conditioned on *S*_*n*_ or *E* is invariant under the swap if and only if *λ*(*t*) = *c µ*(*t*) for almost every *t*, for some constant *c >* 0. Under our skyline FBD implementation, swapping *λ*_*i*_ and *µ*_*i*_ in every interval therefore gives the same density under *S*_*n*_ or *E* exactly when *λ*_*i*_ = *c µ*_*i*_ in every interval for some constant *c >* 0. The enumeration factors of the unresolved likelihood do not contain the rates, so this applies to resolved and unresolved analyses alike. Thus, under our implementation, when *λ*_*i*_*/µ*_*i*_ varies little across intervals, the swapped state has nearly the same density, and the data will carry little information on the sign of *λ*_*i*_ − *µ*_*i*_. Conversely, when the posterior places *λ*_*i*_*/µ*_*i*_ far from constant across intervals, the swapped state has a different density and would be less of a concern.

### 2.2 Unresolved FBD Process

#### The original approach of Heath et al. (2014)

Unresolved fossils are associated with two values: for each fossil *f*, let *y*_*f*_ denote the stratigraphic (fossil) age, and *z*_*f*_ denote the “attachment age” of *f* . When *f* is a tip, *z*_*f*_ could roughly be understood as the timing of one of the speciation events that gave rise to *f* . When *f* is an SA, *z*_*f*_ equals *y*_*f*_ . Each fossil can flip between tip and SA states via reversible-jump MCMC (RJMCMC) (Green, 1995; Gavryushkina et al., 2014; Heath et al., 2014). Given node ages of a fixed backbone tree with *n* taxa and *z*_*f*_ values, one can calculate the likelihood of a corresponding oriented tree with the same fossil sampling and divergence times using Equation (1), by treating *z*_*f*_ values of tip fossils as divergence times. However, multiple trees can correspond to the same (**x, y, z**) configurations (where **x, y, z** are vectors of backbone node ages, fossil ages, and attachment ages, respectively). Given a fixed (**x, y, z**), one can enumerate the number of possible resolved oriented trees that induce it and sum their (equal) density to obtain the likelihood of the unresolved configuration, as follows (Heath et al., 2014): For each fossil *f*, let *γ*_*f*_ denote the number of edges compatible with the clade assignment of *f* that cross *f* ‘s attachment age *z*_*f*_ ; an edge crosses an age *t* when *t* lies strictly between the ages of its two endpoints. A tip fossil may bifurcate from any one of these *γ*_*f*_ edges and giving *γ*_*f*_ resolutions, and be either the left or the right descendant, giving 2*γ*_*f*_ distinct oriented resolutions; a SA fossil may lie on any one of the *γ*_*f*_ edges, with no orientation choice, giving *γ*_*f*_ resolutions. Within a resolution, we refer to the point on the selected edge at age *z*_*f*_ as the “attachment point” of *f* . All resolutions of a given (**x, y, z**) share the same density. Setting *T*_*ur*_ as an unresolved configuration and *T*_*or*_ as an arbitrarily resolved oriented tree (since arbitrary resolution shares the same density under fixed (**x, y, z**)), summing over the density, we get:

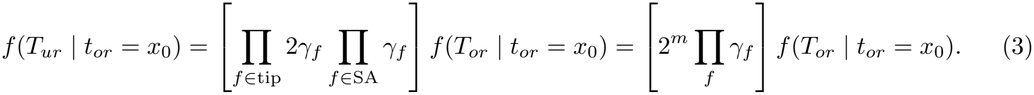

The 2^*m*^ term after the second equality is the equivalent factor as in the oriented–labeled conversion, resulting from our marginalization over orientations for *m* fossil tips. Accordingly, the oriented–labeled conversion for the unresolved configuration involves only the *n* − 1 backbone bifurcations, making the conversion factor 2^*n*−1^*/*(*n*!(*m* + *k*)!), and thus the labeled and oriented unresolved likelihoods are only proportional by a constant.

#### Correcting the enumeration

Equation (3) is correct under a fixed (**x, y, z**) when the clade assignments are not nested, but in practice we always marginalize over the latent attachment ages, **z**. Imagine a simple case where the backbone tree (*T*_*bb*_) consists of two extant taxa, and we have two fossils (*f*_1_, *f*_2_) with the same stratigraphic age (Fig. 1a). Now, imagine that we are sampling 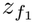 and 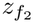 with the backbone node age *x*_1_ fixed, and consider two possible **z** configurations: in configuration 1, 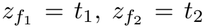, and in configuration 2, 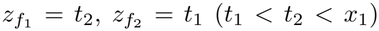 (Fig. 1a). Figure 1b shows the 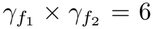 resolutions enumerated for each configuration under the approach of Heath et al. (2014). Among the 12 resolved labeled trees obtained across the two configurations, the two resolutions in which *f*_1_ and *f*_2_ form a cherry (shaded in Fig. 1b) each appear identically in both and are therefore counted twice. For a tip fossil, we refer to the lineage extending rootward from the fossil tip to its attachment point as its “stalk”. The two configurations differ only by exchanging the attachment ages *t*_1_ and *t*_2_ between the two fossils, and the cherry node sits at 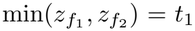 and the lineage rootward of the cherry node joins the backbone at 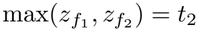 regardless of which fossil carries which value, so both configurations induce the identical resolved tree. The remaining resolutions, in which *f*_1_ and *f*_2_ attach to distinct edges, place the fossils at different ages under the two configurations and so stay distinct. We formalize this overcounting and derive a correction.

**Figure 1:**
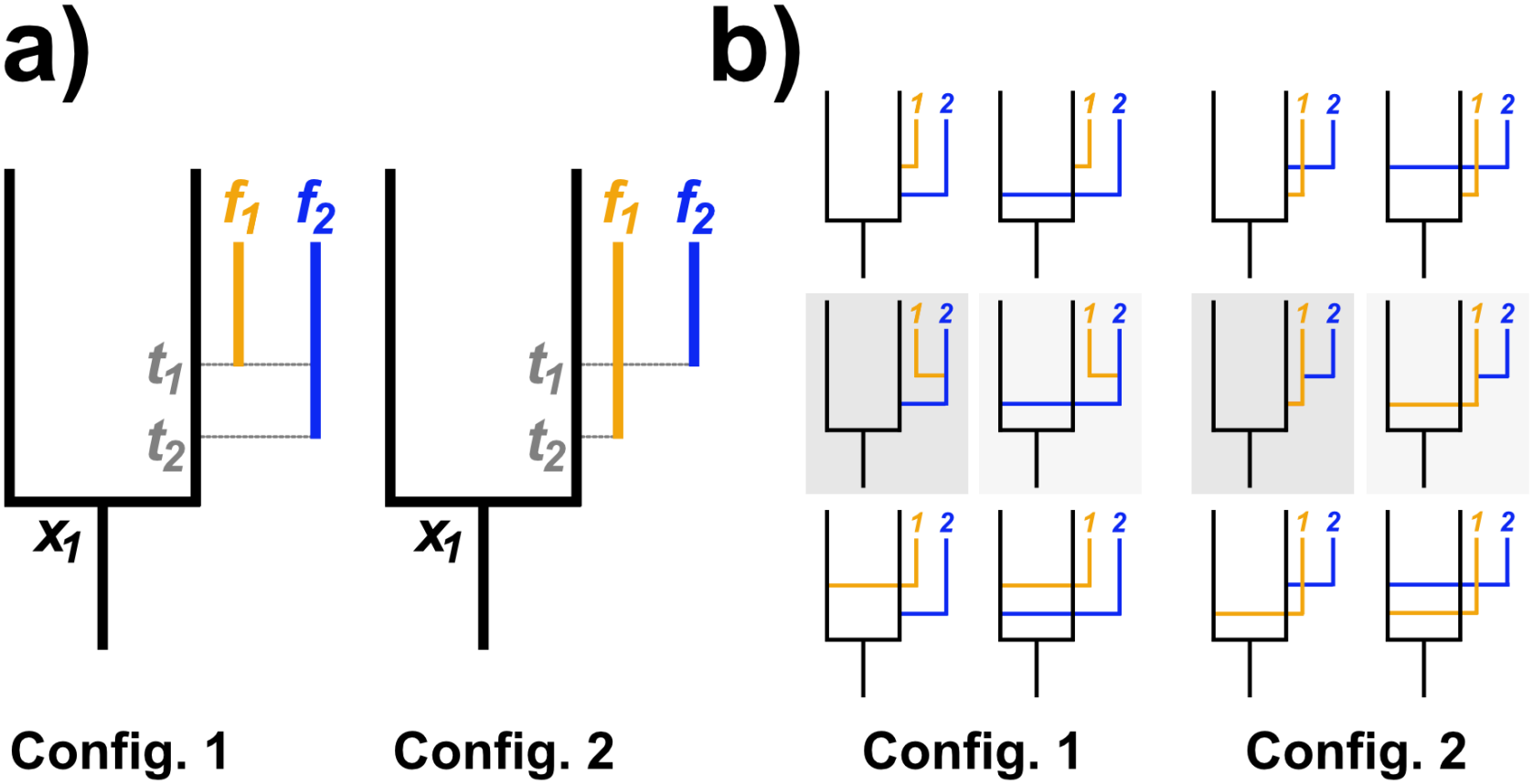
Example of overcounting of possible resolutions under the scheme of Heath et al. (2014). a) Visualization of configurations 1 and 2. Under configuration 1, 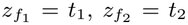, and under configuration 2, 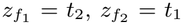, where *t*_1_ *< t*_2_ *< x*_1_ and *x*_1_ is crown node age of the backbone. b) 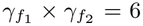 possible resolutions under each configuration. Resolutions with the same shaded background are the same resolved labeled tree, in which *f*_1_ and *f*_2_ form a cherry attached to the same backbone edge, so each of these two trees is counted twice across the two configurations.

We first define some notations. Each fossil *f* is assigned to a monophyletic clade of the backbone (Barido-Sottani et al., 2023), whose crown node we denote by *κ*_*f*_, either as a crown, a stem, or a total group member. The assignment fixes the “attachment zone” *A*_*f*_, the set of backbone edges *f* is allowed to attach to: the edges descending from *κ*_*f*_ for a crown group fossil; the backbone edge immediately rootward of *κ*_*f*_ for a stem group fossil; and both for a total group fossil. When *κ*_*f*_ is the crown node of the backbone itself, the backbone edge rootward of it is connected to the origin *x*_0_. Fix the backbone *T*_*bb*_ (with node ages **x**) and the fossil ages **y**. For given attachment ages **z**, (**x, y, z**) is compatible with a set R(**z**) of resolved labeled trees, all with equal density of Equation (2). Within any *T* ∈ *ℛ* (**z**), let *e*_*f*_ be the backbone edge that *f* reaches by following the chain of stalks it attaches to (Supplementary Figure S1). *T* is compatible with the clade assignments exactly when *e*_*f*_ ∈ *A*_*f*_ for every *f* . Within any *T* ∈ *ℛ* (**z**), define an “*f* -clade” as a maximal fossil-only subtree attached at a single point of a backbone edge (Supplementary Figure S1), so that fossil subtrees attached at different points of the same backbone edge are distinct *f* -clades, and denote *c*(*T* ) as the number of *f* -clades in *T* that contain a tip fossil.

##### Theorem 1

(Overcounting). *Marginalized over* **z**, *the enumeration of Heath et al. (2014), which gives equal weight to every resolution compatible with the clade assignments, assigns total weight* 2^*m*−*c*(T )^ *to each distinct resolved labeled tree* T ∈ *U*_**z**_ *ℛ* (**z**) *compatible with the clade assignments*.

*Proof*. See the Appendix.

For a backbone edge *e* lying in at least one attachment zone, let *Z*(*e*) be the attachment zone that contains *e* and is contained in every other attachment zone containing *e*, which exists and is unique (see Lemma 2 in the Appendix). Let *Ƶ* be the set of tuples **Z** = (*Z*_*f*_ )_*f*_ in which each *Z*_*f*_ is an attachment zone with *Z*_*f*_ ⊆ *A*_*f*_ . Given **Z**, let *b*_*f*_ be the number of backbone edges *e* with *Z*(*e*) = *Z*_*f*_ that cross *z*_*f*_, *s*_*f*_ the number of tip fossils *f* with 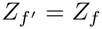 whose stalk crosses *z*_*f*_, and *γ*_*f*_ (**Z**) = *b*_*f*_ + *s*_*f*_ .

##### Theorem 2

(Corrected unresolved likelihood). *Marginalized over* **z**, *replacing γ*_*f*_ *in Equation* (3) *with* 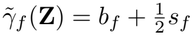 *for every tip fossil and with γ*_*f*_ (**Z**) *for every SA, and summing over* **Z** ∈ *Ƶ, weights each distinct resolved labeled tree compatible with the clade assignments exactly once, giving*

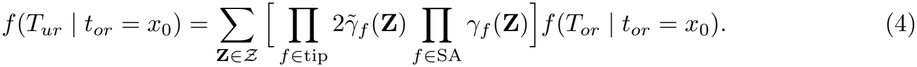

*Proof*. See the Appendix.

Algorithmically, Equation (4) is implemented by halving the weight of each tip fossil stalk crossing *z*_*f*_ when counting 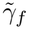, with backbone edges and SAs keeping full weight. Each fossil carries a second latent quantity beside its attachment age, the smallest attachment zone it has reached, *Z*(*e*_*f*_ ), and the counts are confined to *Z*(*e*_*f*_ ): *b*_*f*_ is counted over the backbone edges whose own smallest attachment zone is *Z*(*e*_*f*_ ), and *s*_*f*_ is counted over the stalks of the fossils that have reached *Z*(*e*_*f*_ ) via chain of attachment. The *Z*(*e*_*f*_ ) are sampled by MCMC, as the attachment ages are (see Supplementary Material 1, Section 1). Note that all results regarding corrected enumeration remain unchanged when fossil ages **y** are sampled to take stratigraphic uncertainty into account (Zhang et al., 2016; Barido-Sottani et al., 2019, 2020a), since 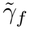 depends only on the current (**x, y, z**) and the *Z*(*e*_*f*_ ), and **y** is integrated over by MCMC.

#### Unsampled extants

It is straightforward to extend the logic for handling unresolved fossils to unsampled extant (UE) species by setting their tip ages to 0 and suppressing the SA-RJMCMC. Earlier approaches typically handled unsampled extant species either by assuming that extant species are sampled through Bernoulli random sampling (Nee et al., 1994; Stadler, 2009, 2010), or by assuming “diversified sampling”, which attempts to model researchers’ efforts to maximize the taxonomic coverage of a clade, and thus to sample as many deep divergences as possible, by assuming that all bifurcation events before some cutoff time are observed but none after it (Höhna et al., 2011; Höhna, 2014; Zhang et al., 2016, 2023). These assumptions can be difficult to meet in practice, and the UE formulation can bypass them at the cost of increased computation. Setting *ρ* = 1 is only possible if we know every extant species in the clade with certainty, and omitting the probability that these particular species were the ones sampled then results in an exact posterior when, within each clade to which UEs are assigned, the sampled species were chosen independently of the tree within that clade, for example by random sampling within the clade. This is the setting of clade-specific sampling fractions, which cannot be incorporated consistently when unsampled species are integrated out analytically (Stadler, 2009; Beaulieu, 2021). Diversified sampling cannot be represented exactly, but the analogous design, in which one sequences a single species per clade, or several chosen at random, and assigns the remaining species of each clade to that clade, can be treated exactly. We also note that data-augmentation approaches to clade-specific sampling fractions, as implemented in ClaDS (Maliet & Morlon, 2022) and the birth-death diffusion model (Quintero et al., 2024), differ from the UE formulation in that only the sampling fraction of each clade is used, while the number of unsampled species is random.

#### No-backbone cases

Given only fossil ages and the number of extant taxa without any backbone tree, we can also estimate speciation, extinction, and sampling rates by treating every given taxon as unresolved and integrating over possible tree topologies by marginalizing their attachment ages with proper topological enumeration (Equation (4)). To make sure that the FBD process starts properly at the origin *x*_0_, we select one taxon and connect it to the origin to form an artificial “spine” backbone, fixing its attachment age at *x*_0_ and excluding it from the attachment and SA moves. Its stalk is thereafter a backbone edge, and so contributes to *b*_*f*_ at full weight rather than to *s*_*f*_ for other fossils *f* . When the “absolute” youngest taxon exists, for example, when extant taxa are present, or the youngest taxon is a fossil whose upper stratigraphic bound is younger than the lower age bound of every other fossil, we can select such a taxon (breaking ties randomly if needed), and elect it as a spine (since the youngest taxa can never be SA). However, when there is no absolute youngest taxon, which can happen when there are no extant taxa and fossil ages are sampled from stratigraphic age bounds without the absolute youngest fossil, so that the youngest fossil taxon fluctuates across MCMC iterations, the spine taxon is re-elected dynamically at each iteration (see Supplementary Material 1, Section 1).

## 3 Implementation

We implemented the unresolved FBD model using the adaptive Metropolis-coupled MCMC (MC3) algorithm (Geyer, 1991; Gilks & Roberts, 1996; Miasojedow et al., 2013) (collapsing to regular MCMC when no heated chains are used) with deterministic even-odd (DEO) scheme (Okabe et al., 2001; Lingenheil et al., 2009; Syed et al., 2022), as we found that parallel tempering was often necessary to achieve convergence of certain parameters within a reasonable amount of time. Here, we describe the implemented algorithm briefly. Details can be found in Supplementary Material 1. Throughout, gamma and exponential priors are parameterized by rate, and normal and truncated normal priors by the mean and variance of the (untruncated) normal. Source code and Linux binary of the implemented method are available from: https://github.com/wnsplim/UFBD.

### FBD process

We sample node ages (**x**) and attachment ages (**z**) (plus fossil ages **y**, if sampled, and the *Z*(*e*_*f*_ ), if the assignments are nested) from the unresolved FBD density described above. SA-RJMCMC was implemented following Gavryushkina et al. (2014) and Heath et al. (2014). When skyline rates were set, we allowed for smoothing of the log-rates across bins with an Ornstein–Uhlenbeck (OU) process (Uhlenbeck & Ornstein, 1930), as we found it stabilized variance and improved MCMC convergence. Specifically, for each rates assigned an OU smoothing, for the log-rate *ξ*_*i*_ = log *r*_*i*_ for rate *r*_*i*_ (which can be either *λ*_*i*_, *µ*_*i*_, or *ψ*_*i*_) of bin *i* we write

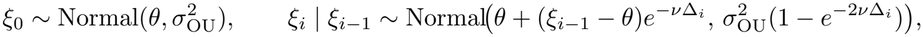

where *θ* is the long-term mean log-rate, 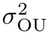 the stationary variance, *ν* the strength of mean reversion, and Δ_*i*_ the time between the midpoints of bins *i* − 1 and *i* (see Supplementary Material 1, Section 1 for further details including the prior specification for *θ*, 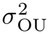, and *ν*); so that the log-rate of each bin reverts toward a long-term mean, and the log-rates of adjacent bins are correlated, with the correlation decaying as the time separation between the bins increases. This is similar to the OU rate smoothing implementation of du Plessis (2016) and Zhang et al. (2023), but parameterizing it by the stationary variance of the log-rates directly, rather than the per-unit-time variance of its fluctuations, from which the stationary variance follows only in combination with the reversion rate.

To efficiently calculate *γ*_*f*_ (**Z**) and 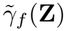 (**Z**) (Eq. 4) for every fossil, which changes whenever **x, y, z**, or a *Z*(*e*_*f*_ ) is updated, we compute these counts incrementally using sorted edge-age arrays, held one per *Z*(*e*) and assigned once by a subtree index, and recomputing only the fossils affected by each move, which reduces the per-step counting cost from O((*m* + *k*)*ñ*) to O((*m* + *k*) log *ñ*) for local moves, where *ñ* is the number of backbone edges and stalks of unresolved fossils and extants compatible with the fossil (see Supplementary Material 1, Section 2).

### Sequence likelihood

When the sequence is directly given, we calculate the likelihood of the sequence given the backbone tree topology using the standard pruning algorithm (Felsenstein, 1981) under the GTR model (Tavaré, 1986) with discrete gamma rate heterogeneity (Yang, 1994), with optional invariable sites (Churchill et al., 1992; Gu et al., 1995). We did not implement other substitution models, since it has been shown that phylogenetic analyses, including divergence time estimation, are relatively robust to substitution model choice, at least among GTR-class models (Huelsenbeck & Rannala, 2004; Barba-Montoya et al., 2020; Tao et al., 2020; Fabreti & Höhna, 2023). Because the backbone topology is fixed, we accelerate the pruning computation by detecting site repeats once at initialization and reusing them throughout the analysis, as described in Kobert et al. (2017), and also by omitting taxa absent from a partition (Stamatakis & Alachiotis, 2010).

We also implemented the approximate likelihood approach where the likelihood of the fixed topology is approximated as a multivariate normal density via maximum likelihood estimates of branch lengths and the Hessian of the log-likelihoods (Thorne et al., 1998; Thorne & Kishino, 2002; Seo et al., 2004; Guindon, 2010; dos Reis & Yang, 2011), which can be computed through external programs like PAML (Yang, 2007), IQ-TREE (Demotte et al., 2025), and phyloHessian (Wang & Meade, 2026). Following dos Reis and Yang (2011), we applied an arcsine transformation to the branch lengths before the normal approximation. We note that Stadler and Yang (2013) implemented an approximate likelihood approach under the FBD model, but under the fixed, fully resolved tree without sampled ancestors.

### Molecular clock

We implemented two relaxed clock models: an uncorrelated log-normal (UCLN) clock of Drummond et al. (2006), and an autocorrelated geometric Brownian motion (GBM) clock of Kishino et al. (2001). Across clock partitions, the vectors of clock-partition mean rates, (*r*_clock,*j*_)_*j*_, and clock-rate variances, 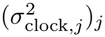, are each assigned a gamma-Dirichlet prior (dos Reis et al., 2014), which reduces to a gamma prior when there is a single clock partition.

The clock-rate variance 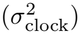 and the branch-specific rates (*r*_*b*_, where *b* is a branch index) can produce a funnel-shaped posterior geometry, which is difficult to explore efficiently (Neal, 2003). One can sample the log-rate of each branch directly, which works well when the sequence data strongly constrain the branch rates, because the branch rate is then determined largely by the data, so its posterior is approximately independent of 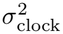. However, when the likelihood is diffuse, the branch rate is determined largely by the prior, and direct sampling retains a strong prior-induced dependence between the branch rates and 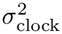. An alternative is to sample a standardized latent variable *u*_*b*_ ∼ Normal(0, 1^2^) and express the per-branch clock rate as log *r*_*b*_ = *η*_*b*_ + *s*_*b*_*u*_*b*_, where *η*_*b*_ and *s*_*b*_ are the prior mean and standard deviation of log *r*_*b*_. This non-centered parameterization can improve mixing of 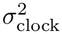 when branch-specific likelihoods are weak, because the standardized variables *u*_*b*_ are a priori independent of 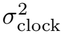 (Papaspiliopoulos et al., 2007). We observed that this non-centered form improved 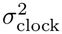 mixing under diffuse likelihoods, but not when the likelihood is concentrated; when the latter is the case, the sequence data effectively pin log *r*_*b*_, so any change in *s*_*b*_, and therefore in 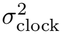, must be compensated by a corresponding change in *u*_*b*_ to keep log *r*_*b*_ near the likelihood-supported value. The non-centered parameterization can therefore introduce strong posterior dependence between *u*_*b*_ and 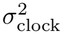, slowing the mixing. Which of the two forms mixes better therefore depends on how strongly the sequence data constrain the branch rates. Our implementation samples the log-rates directly (centered parameterization) by default.

## 4 Experiments and Examples

### 4.1 Simulation-based calibration checking

To demonstrate that our implemented algorithm and unresolved likelihood are well formulated (Mendes et al., 2025), we performed simulation-based calibration checking (SBC) (Cook et al., 2006; Talts et al., 2020; Modrák et al., 2025). We simulated the FBD process conditioned on survival from the origin under three rate models: (i) constant rates with *λ* ∼ Unif(0.03, 0.08), *µ* ∼ Unif(0.02, 0.06), *ψ* ∼ Unif(0.04, 0.2), and *x*_0_ ∼ LogNormal(4.0, 0.1^2^); (ii) skyline rates with interval boundaries at 20 and 40, with the rates of each interval drawn independently from the priors of (i); and (iii) the same intervals with OU-smoothed *λ*_*i*_ and *µ*_*i*_, with *θ* ∼ Normal(log 0.055, 0.15^2^) for *λ, θ* ∼ Normal(log 0.04, 0.15^2^) for *µ, σ*_OU_ ∼ Exp(30), and *ν* ∼ Exp(2), and with *ψ*_*i*_ drawn independently as in (ii). Under each rate model, each extant taxon was designated a UE with probability 0, 0.5, or 1, giving nine settings. Simulated trees that did not survive were discarded and resimulated with the same parameter values, which matches the survival conditioning used in inference. Fossils and UEs were then pruned from each simulated tree, leaving only labels and fossil ages with unresolved attachment. The three constant-rate settings were also analyzed with the enumeration of Heath et al. (2014). Four further settings, testing the sampling of fossil ages (**y**), the sampling of attachment zones (**Z**) under nested clade assignments, extinction conditioning, and dynamic spine election, are described in Supplementary Material 1, Section 3.

As test quantities, we used the rates of each interval, the OU parameters in rate model (iii), the number of sampled ancestors (*k*), the origin age *x*_0_, and, in replicates whose backbone has at least two tips, the crown age *x*_1_ and the age of one internal backbone node chosen uniformly at random. Because the node is chosen independently of the true values and using only the observed backbone, the random node age is a valid test quantity (Modrák et al., 2025). Following Modrák et al. (2025), we also used the joint log-likelihood and the joint log-prior density. Each setting comprised 5,000 simulated datasets, each analyzed with a single MCMC chain of 10 million iterations sampled every 2,000 iterations, with the first 10% of samples discarded as burn-in. No replicate failed or was dropped, and in every setting at least 99% of the replicates had bulk and tail ESS of at least 400 for all parameters.

We computed randomized normalized ranks (with ties broken at random) of each test quantity and assessed their uniformity with the empirical-CDF *γ* statistic of Säilynoja et al. (2022) (unrelated to the counts *γ*_*f*_ ), evaluated at the points *i/n* (*i* = 1, …, *n* − 1), where *n* is the number of replicates in which the test quantity is defined. Its *p*-value was obtained from the distribution of *γ* under uniformity, estimated from 40,000 simulated sets of *n* independent Unif(0, 1) ranks (note that thus *p*-value under our settings have a floor of 1*/*40,001). Because each setting has many test quantities, some *p*-values are expected to be small by chance even when the implementation is correct (Cook et al., 2006; Modrák et al., 2025). We therefore adjusted the *p*-values within each setting for multiple testing using the Bonferroni correction (Dunn, 1961) and the procedures of Benjamini and Hochberg (1995) (BH) and Benjamini and Yekutieli (2001) (BY), of which BH is the least conservative. We considered a test quantity to reject uniformity when its adjusted *p*-value was below 0.05.

### 4.2 Empirical datasets

We demonstrate analyses on three datasets representing three major use cases of the method: approximate likelihood path utilizing a Hessian file derived from the genome-scale alignment of Cetacea (whales, dolphins, and porpoises) from McGowen et al. (2020); full likelihood path using sequence alignment of turtle family Emydidae (pond turtles) extracted from Thomson et al. (2021) turtle phylogeny, and no-backbone case with extinct Paleozoic early winged insect group, Palaeodictyopterida (Prokop & Engel, 2019). We ran 4 parallel chains of MC3 with 4 coupled chains (one cold chain, three hot chains; Cetacea, Palaeodictyopterida), or plain MCMC without coupled chains (Emydidae), until the rank-normalized split 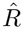 (Gelman & Rubin, 1992; Gelman et al., 2013; Vehtari et al., 2021) fell below 1.01, and the minimum bulk ESS (the effective sample size of the rank-normalized draws) and tail ESS (the smaller of the effective sample sizes of the 5% and 95% quantile indicators) exceeded 400 for all parameters, following the suggestions of Vehtari et al. (2021). Fossil data were primarily retrieved from the Paleobiology Database (https://paleobiodb.org; accessed July 13–26, 2026), with validity, clade assignment, and ages audited against the literature (see Supplementary Material 2). Each species entered the analyses once, as a single sampled fossil whose age range is that of its holo-type’s stratigraphic horizon or, where the holotype lacks a usable horizon, that of the best-dated validly referred specimen (see Supplementary Material 2), and fossils of extant species were excluded. This filtering was necessary for the standard FBD process implemented here, which assumes that every sampled fossil is a distinct labeled taxon and so cannot accommodate another occurrence of a species already in the tree, whether as a fossil or as an extant tip; incorporating multiple sampling instances of the same species will require extensions like the stratigraphic range FBD model (Stadler et al., 2018) (see Discussion). We chose clades in which most species are known from a single sampling event, so that this assumption is met as closely as possible. Of the species matched to Paleobiology Database records, 336 of 535 (Cetacea), 10 of 18 (Emydidae), and 408 of 446 (Palaeodictyopterida) were recorded from a single locality and horizon, counting collections that share coordinates and age range as one. Among the remaining species, 83 of 199 (Cetacea), 3 of 8 (Emydidae), and 30 of 38 (Palaeodictyopterida) were recorded from two sampling events.

For skyline analyses, the OU long-term mean log-rate, stationary log-standard deviation, and reversion rate were assigned *θ* ∼ Normal(log 0.2, 1.6^2^), *σ*_OU_ ∼ Exp(10*/*3), and *ν* ∼ Exp(2) priors (which equal the defaults of the implementation), respectively, for speciation and extinction rates. Under these priors, the long-term rate *e*^*θ*^ lies between about 0.009 and 4.6 events per lineage per Myr with 95% prior probability, the rate of a bin lies within a factor of 1.12 of *e*^*θ*^ with 50% prior probability and within a factor of 2.5 with 95% prior probability, and the log-rates of two bins whose midpoints are 1 Myr apart have a prior median correlation of about 0.7. The data rarely inform *ν*, so this prior largely sets the degree of smoothing. The cutoff of the open-ended interval was placed near the oldest fossil records, beyond which few fossil age ranges extend (1 of 535 beyond 50 Mya for Cetacea and 17 of 446 beyond 320 Mya for Palaeodictyopterida), and bin widths of 1 Myr (Cetacea) and 2.5 Myr (Palaeodictyopterida) were chosen based on fossil density. The binning scheme is a modeling choice that should, in practice, be made carefully with regard to the data in hand; importantly, the resolution at which the rate trajectory can be recovered will depend on both the binning scheme and the informativeness of the data. We note that our bins were intended to give a reasonably fine resolution and are not necessarily optimal. We did not assign OU smoothing to the preservation rate, as we believe that smoothing the preservation rate is less biologically defensible than speciation or extinction rates.

For Cetacea, we obtained the six-partition alignment and corresponding tree topology from Mc-Gowen et al. (2020) and pruned all non-cetacean outgroups. We used PAML v4.9h (Yang, 2007) to calculate maximum-likelihood branch lengths and the Hessian matrix under the optimization scheme of Yang (2000). We analyzed these data using a per-partition GBM clock with centered parameterization and 51 skyline intervals, with 1-Myr bins from 0 to 50 Mya and one open-ended interval beyond 50 Mya, with OU-smoothed *λ* and *µ*, independent *ψ*_*i*_ ∼ Exp(2), and *x*_0_ ∼ TruncNormal(55, 2^2^) bounded below at 50 Mya, based on the published estimates of the Cetacea–Hippopotamidae divergence (Meredith et al., 2011; Zurano et al., 2019) (see Supplementary Material 2). For Emydidae, we pruned all non-emydid species from the alignment and tree of Thomson et al. (2021) and analyzed the 15 loci directly under GTR+Γ_4_ and a per-partition UCLN clock under non-centered parameterization. Performance comparison with non-centered and centered parameterization for this dataset was additionally explored in Supplementary Material 1, Section 1. We used constant *λ, µ, ψ* ∼ Exp(10), as fossil records of Emydidae were relatively scarce (21 fossil species after filtering, when compared to 535 and 446 for Cetacea and Palaeodictyopterida respectively; see Supplementary Material 2) and *x*_0_ − 54.8 ∼ Exp(0.1), a relatively diffuse prior offset at the older bound of the age interval of the oldest fossil. For both Cetacea and Emydidae datasets, all extant species not present in the backbone tree were set as UE, and when multiple samples per species existed, only the sample with the highest sequence coverage was retained. The Palaeodictyopterida dataset was analyzed without a backbone under extinction conditioning with *ρ* = 1 at 250 Mya, so that no lineage survives to 250 Mya, as this group is widely accepted to have not survived the end-Permian extinction (Grimaldi & Engel, 2005; Prokop & Engel, 2019). To do so, all ages were shifted so that 250 Mya is the present of the process, and with *ρ* = 1, conditioning on *E* conditions on no lineage of the clade surviving to 250 Mya. 29 skyline intervals, with 2.5-Myr bins from 250 to 320 Mya and one open-ended interval beyond 320 Mya, with OU-smoothed *λ* and *µ*, independent *ψ*_*i*_ ∼ Exp(10), and *x*_0_ − 327 ∼ Exp(0.1), likewise offset just beyond the older bounds of the oldest fossil age intervals, were used. See Supplementary Material 2 for detailed MCMC settings and justifications for the origin prior.

To assess the downstream consequences of the overcounting for the estimated divergence times and rates, we reanalyzed all three datasets with the enumeration of Heath et al. (2014), with all other settings unchanged. To assess the information that the molecular data provide for the estimation of rate trajectories and node ages, we also analyzed the Cetacea dataset without the Hessian file and the clock model, so that the node ages are informed only by the FBD process given the backbone topology and the fossils. Finally, since the origin prior of the Cetacea analysis was centered on published estimates with relatively small variance, we tested the robustness of the Cetacea estimates to this prior by replacing it with a prior of the same form as for Emydidae and Palaeodictyopterida, *x*_0_ − 46 ∼ Exp(0.1), which was chosen based on the age of the oldest fossil record (see Supplementary Material 2). We summarize node ages by posterior means, matching the posterior mean trees, and rates and the number of sampled ancestors by posterior medians. All trees were visualized using FigTree v 1.4.4 (Rambaut, 2018), or the R package ape v 5.8.1 (Paradis & Schliep, 2019).

## 5 Results

### 5.1 Simulation-based calibration checking

The results of simulation-based calibration checking under constant rate FBD are shown in Figure 2. Under the corrected enumeration, none of the test quantities rejected uniformity in any of the three UE regimes under any of the three adjustments, and the smallest BH-adjusted *p*-value was 0.69. Under the enumeration of Heath et al. (2014), every test quantity except the log prior density rejected uniformity in every regime for all three *p*-value adjustments, except *x*_0_ at a UE probability of 0, and the discrepancy grew monotonically with the proportion of extant taxa designated as UE. The ranks of *λ* and *µ* concentrated at low values and those of *ψ* at high values, so under the constant-rate settings simulated here, the overcounting inflated the posteriors of the speciation and extinction rates while deflating that of the sampling rate, with the bias increasing as more taxa were left unresolved. Under skyline rates, again none of the test quantities rejected uniformity under any adjustment, either with independent rates or with OU-smoothed rates (Supplementary Figures. S2 and S3), and the smallest BH-adjusted *p*-value was 0.13.

**Figure 2:**
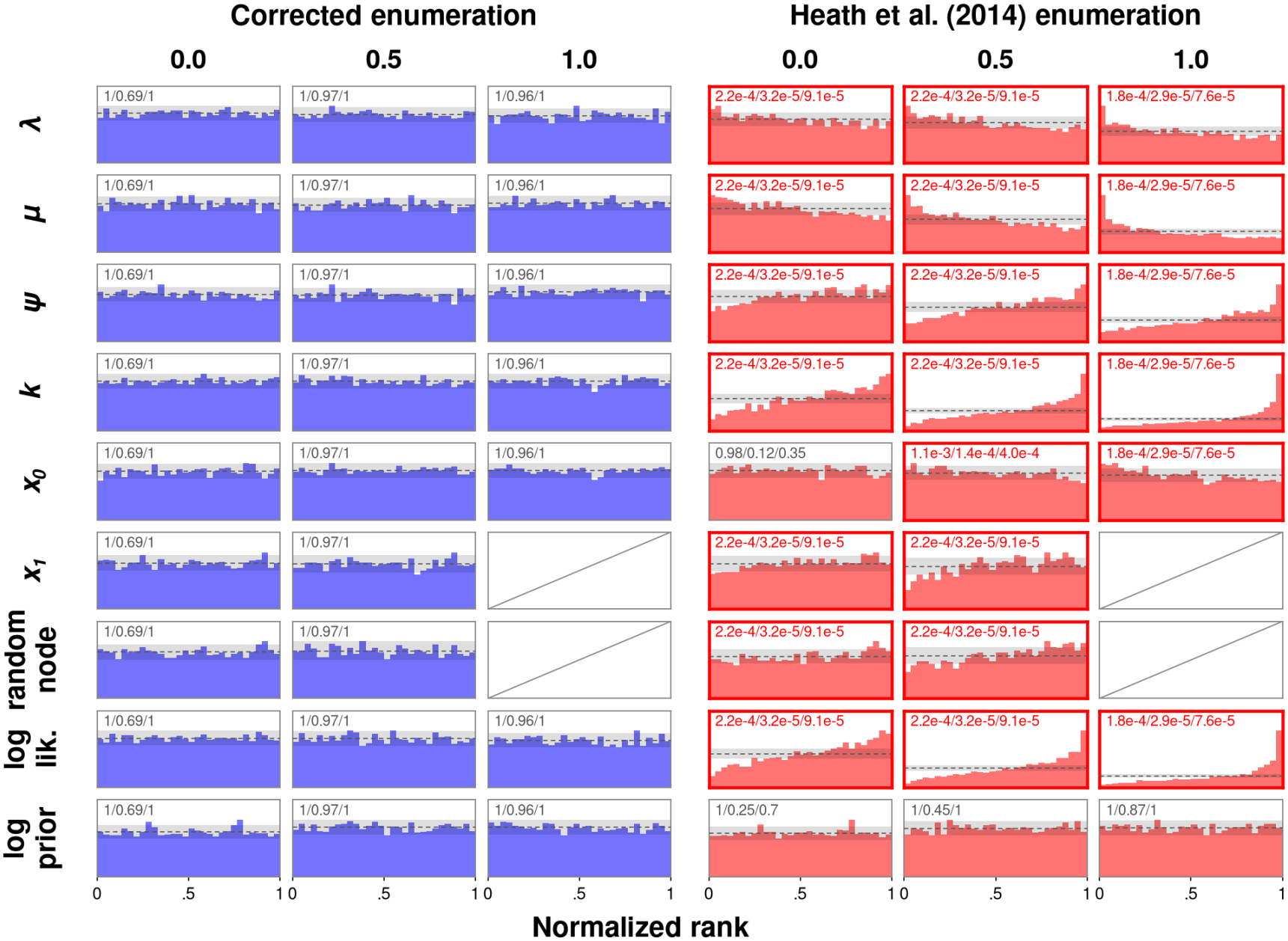
Results of the simulation-based calibration checking under constant rates FBD (model (i)). Each panel shows a histogram of the rank of the true value of a test quantity relative to the values sampled from the MCMC-inferred posterior, in repeated sampling, when sampling parameter values from the prior and simulating data conditionally on the sampled parameter values. Under a well-calibrated sampler, the ranks should be uniformly distributed (Cook et al., 2006; Talts et al., 2020). Rows correspond to the test quantities, with *k* the number of sampled ancestors, random node the age of one internal backbone node chosen uniformly at random, and log lik. and log prior the joint log-likelihood and log-prior density. Columns correspond to the probability with which each extant taxon was designated a UE; *x*_1_ and the random node are undefined when this probability is 1. The left block uses the corrected enumeration and the right block the enumeration of Heath et al. (2014). The dashed line is the expected bin count under uniformity and the grey band its pointwise 95% interval. The numbers in the top left of each panel are the Bonferroni-, BH-, and BY-adjusted *p*-values of the empirical-CDF *γ* statistic of Säilynoja et al. (2022), and thick frames mark panels with a BH-adjusted *p*-value below 0.05.

### 5.2 Empirical analyses

The Cetacea, Emydidae, and Palaeodictyopterida analyses converged after approximately 130, 23, and 725 million MCMC iterations, respectively, taking approximately 38, 45, and 25 hours of wall-clock time. Each used 16 cores on an Intel Xeon Platinum 8358 2.6 GHz processor. We note that the Cetacea and Emydidae runs used full clock partitioning, with a separate clock model assigned to each partition, which made convergence more challenging than when using a single shared clock model. Runtime can be reduced by using fewer clock partitions, at the cost of precision (Rannala & Yang, 2007; dos Reis & Yang, 2013; Angelis et al., 2018).

#### Cetacea

The resulting posterior mean tree for the Cetacea analysis is shown in Figure 3, and speciation, extinction, diversification, and sampling rate through time are plotted in Figure 4. Crown Cetacea was estimated to have appeared around ∼ 37.18 Mya, with a relatively tight 95% credible interval (CI) of 36.63–37.78 Mya (Fig. 3), overlapping the estimates of Zurano et al. (2019) (36.39 Mya; 95% HPD 36.05–36.88 Mya), McGowen et al. (2020) (36.72 Mya; 36.4–37.5 Mya), and the metatree analysis of Lloyd and Slater (2021) (36.9 Mya; 35.3–38.6 Mya), but older than Zhou et al. (2011) (34.43 Mya; 33.51–36.28 Mya). Crown age of Odontoceti was estimated to be ∼ 34.91 Mya (95% CI 33.12–36.61 Mya), overlapping McGowen et al. (2020) (34.13 Mya; 33.7–34.9 Mya), Zurano et al. (2019) (32.55 Mya; 29.70–35.10 Mya), and Zhou et al. (2011) (30.13 Mya; 25.75–33.84 Mya), but older than Lloyd and Slater (2021) (29.11 Mya; 27.7–30.5 Mya). Crown age of Mysticeti was estimated to be ∼ 22.73 Mya (95% CI 21.19–24.22 Mya), overlapping Zurano et al. (2019) (21.48 Mya; 20.58–22.62 Mya) and Lloyd and Slater (2021) (25.49 Mya; 23.4–27.4 Mya), but younger than McGowen et al. (2020) (25.75 Mya; 25.22–26.72 Mya).

**Figure 3:**
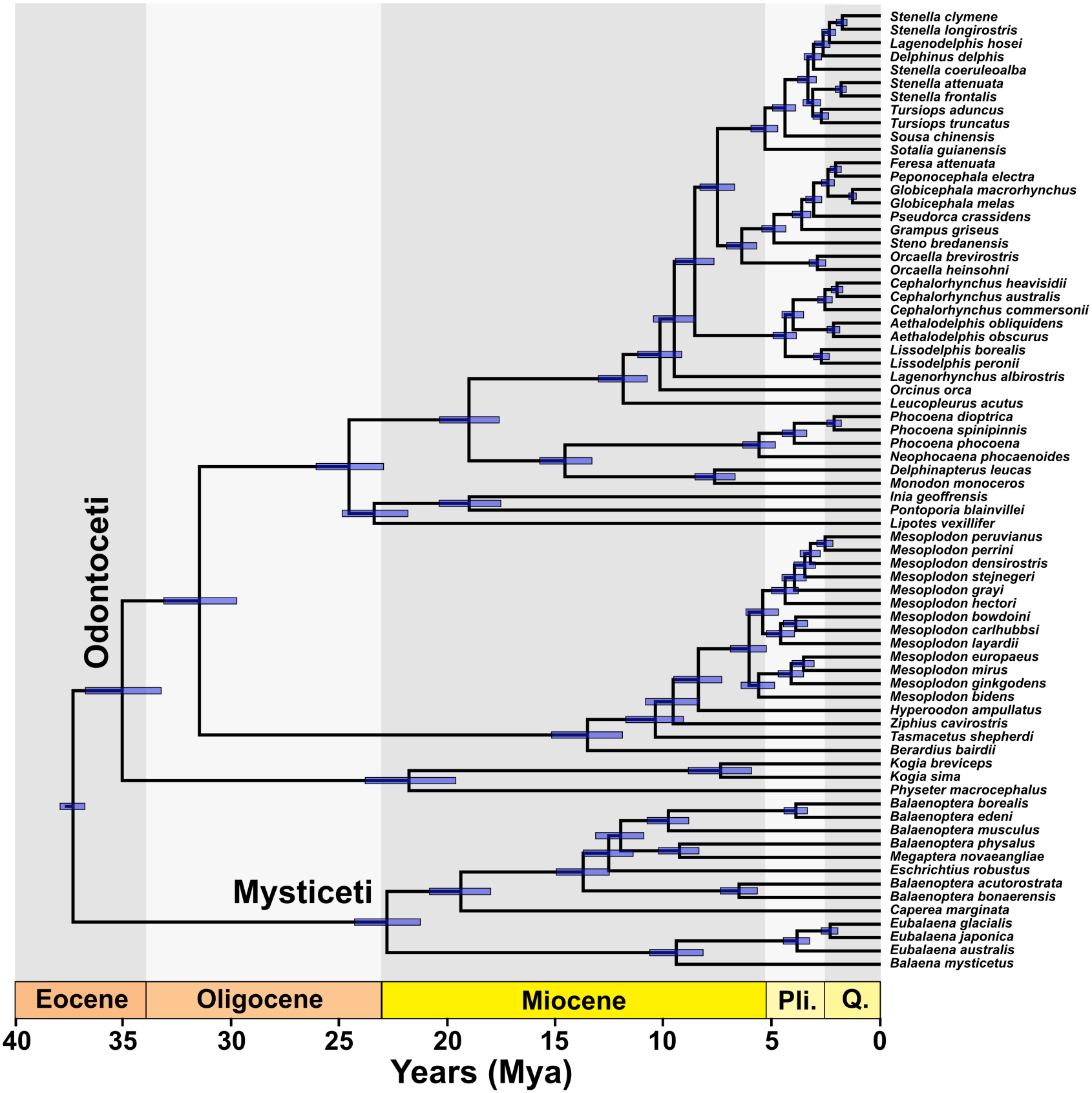
Posterior mean dated tree resulting from the Cetacea analysis. Node bars denote 95% credible intervals for that node age. Abbreviations: Pli: Pliocene, Q: Quaternary

**Figure 4:**
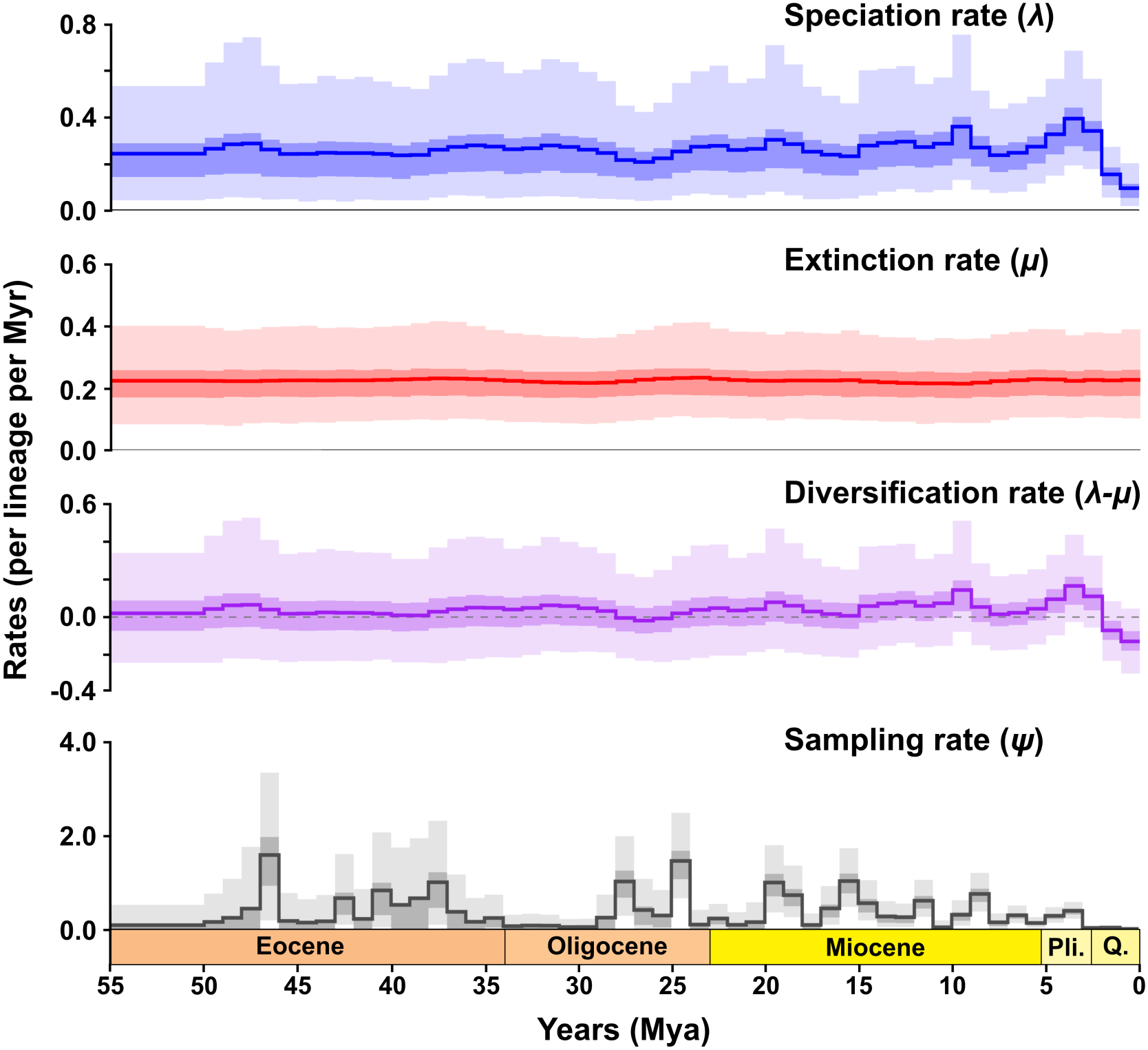
Speciation, extinction, diversification, and sampling rates resulting from the Cetacea analysis. Lines denote posterior medians, darker shades denote 50% credible intervals, and lighter shades denote 95% credible intervals. Abbreviations: Pli: Pliocene, Q: Quaternary

The posterior median extinction rate of Cetacea was nearly constant at about 0.22 events per lineage per Myr across all bins, and the posterior median speciation rate mostly stayed between 0.21 and 0.30 but dropped to 0.15 and 0.09 in the last 2 Myr (Fig. 4). The estimated sampling rate varied considerably over time (Fig. 4). Our speciation and extinction rate estimates were of a similar order to those of the metatree analysis of Lloyd and Slater (2021) (approximately 0.16–0.26 and 0.10– 0.25 events per lineage per Myr, respectively) and the extant species-only analysis of Mazet et al. (2023) (approximately 0.33, and increasing from near 0 to approximately 1.25 toward the present, respectively), although the estimated trajectories differed. We could not find convincing evidence of the elevated extinction around the Oligocene–Miocene transition noted by Bisconti et al. (2023) from the fossil record of mysticetes, nor the increased diversification rate during the late Miocene–early Pliocene suggested by Steeman et al. (2009), beyond short increases of about 0.1 in the posterior median speciation rate at 9–10 and 3–4 Mya.

Under the enumeration of Heath et al. (2014), the estimated rate trajectories of Cetacea differed markedly from those of the corrected enumeration, with posterior medians of *λ*_*i*_, *µ*_*i*_, and *ψ*_*i*_ about 1.0–12.1, 5.2–6.5, and 0.06–1.05 times those of the corrected enumeration and the median number of sampled ancestors falling from 390 to 72, yet the posterior mean node ages shifted by at most 0.77 Myr, all within the 95% CIs of the corrected enumeration (Supplementary Figure S4). This likely reflects the dense assignment of Cetacea fossils to fine clades, which constrains their attachments and, in turn, the node ages under either enumeration. Thus, to test the possibility of whether the enumeration of Heath et al. (2014) affects the node age estimates, we ran the analysis where we relaxed the assignments to the total groups of Odontoceti, Mysticeti, or Cetacea, with 5-Myr bins from 0 to 50 Mya and one open-ended interval beyond 50 Mya for faster convergence. Under the enumeration of Heath et al. (2014), this analysis did not converge within the maximum cutoff of 500 million iterations (maximum 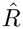 of 1.016, minimum bulk ESS of 194), with three of its four chains spending 17% of their samples with crown Mysticeti younger than 10 Mya, which the corrected enumeration never visited (Supplementary Figure S5).

Without the Hessian file and the clock model, the 95% CIs of node ages were a median 3.4 times wider, crown Odontoceti was estimated at 26.31 Mya instead of 34.91 Mya, and *λ*_*i*_ and *µ*_*i*_ were nearly constant across intervals (Supplementary Figure S6), so the molecular data informed both the node ages and the rate trajectory. Replacing the origin prior with *x*_0_ − 46 ∼ Exp(0.1) moved the posterior mean origin age from 54.08 to 51.14 Mya, but shifted the posterior mean node ages by at most 0.11 Myr (Supplementary Figure S7).

#### Emydidae

The resulting posterior mean tree is shown in Figure 5, with an inset showing the posterior distribution of speciation, extinction, and sampling rates. Crown Emydidae was estimated to have appeared around ∼ 40.20 Mya, with a 95% CI 34.54–46.65 Mya (Fig. 5), compared with 36.14 Mya (95% HPD 32.00–42.99 Mya) in the dated tree of Thomson et al. (2021) and 41.79 Mya (32–71.61 Mya) in Spinks et al. (2016). Two subfamilies, Emydinae and Deirochelyinae, were estimated to have crown ages of 18.51 Mya (95% CI 14.70–22.70 Mya) and 26.69 Mya (95% CI 19.73–34.31 Mya), respectively, compared with 16.48 (11.57–21.59) and 23.24 (17.96–28.92) Mya in Thomson et al. (2021) and 28.84 (17.98–51.13) and 31.08 (20.93–54.33) Mya in Spinks et al. (2016). All six intervals overlapped our result. The posterior medians of the speciation, extinction, and diversification rates were 0.109 (95% CI 0.074–0.149), 0.012 (95% CI 0.000–0.049), and 0.093 (95% CI 0.058–0.130) events per lineage per Myr, respectively (see also the inset in Figure 5), with speciation and diversification rate being larger than the typical values for turtles reported in previous studies only based on the timetree of extant species only (Colston et al., 2020; Thomson et al., 2021). Under the enumeration of Heath et al. (2014), the Emydidae estimates were essentially unchanged, since both enumerations favored configurations in which nearly all fossils are sampled ancestors (Supplementary Figure S8).

**Figure 5:**
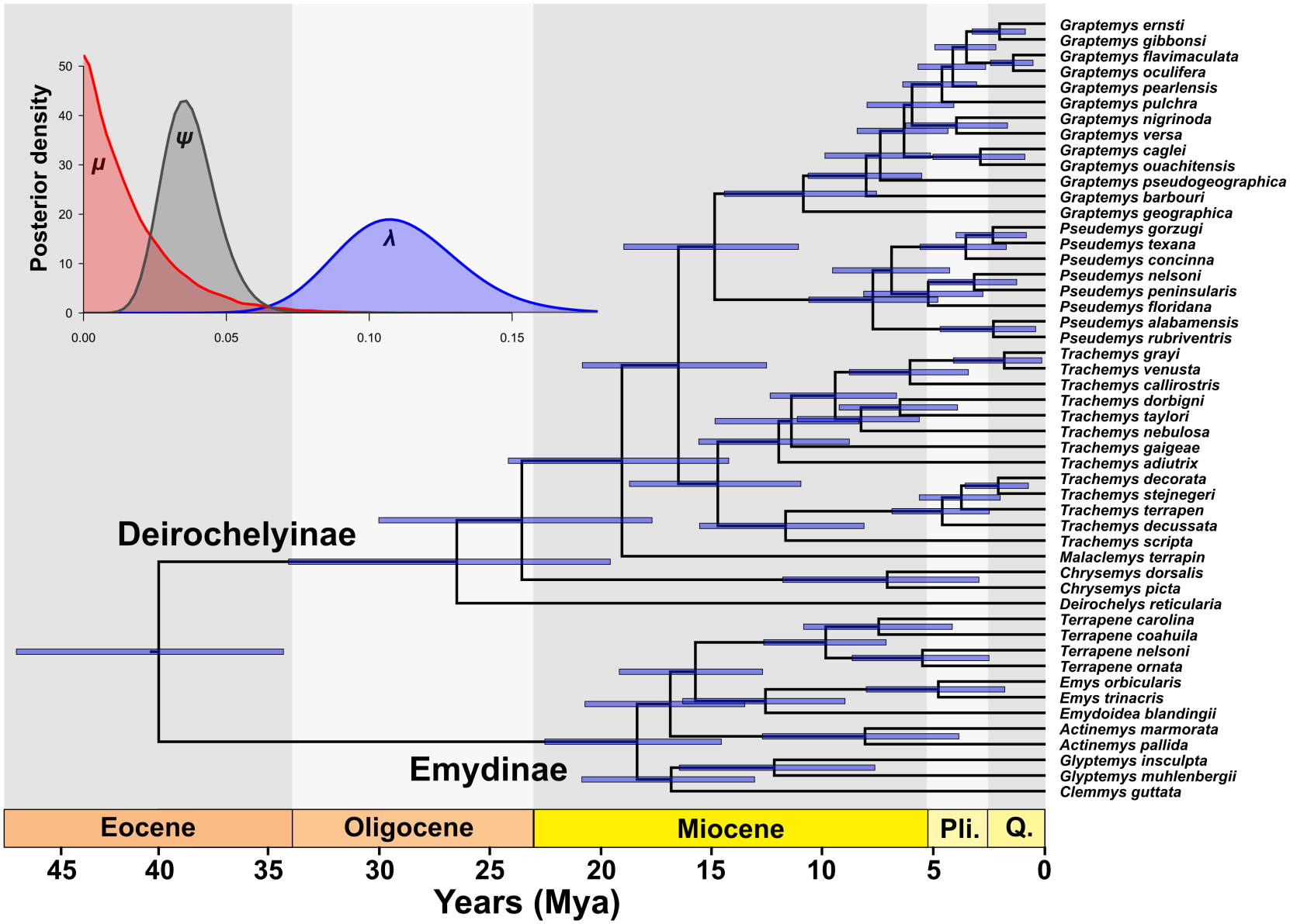
Posterior mean dated tree resulting from the Emydidae analysis. Node bars denote 95% credible intervals for that node age. Top left inset shows the posterior distribution of speciation (*λ*), extinction (*µ*), and sampling (*ψ*) rates. Abbreviations: Pli: Pliocene, Q: Quaternary

#### Palaeodictyopterida

The resulting speciation, extinction, diversification, and sampling rate through time for the Palaeodictyopterida analysis are plotted in Figure 6. Because the Palaeodictyopterida dataset was analyzed under extinction conditioning, the interpretation of the sign of its diversification rate requires care (see “Conditioning” paragraph above). The posterior median of *λ*_*i*_*/µ*_*i*_ ranged from 0.49 to 2.10 across the 29 intervals, being far from constant across intervals. To test whether the data support the swapped configuration, we took the final state of the four cold chains and exchanged *λ*_*i*_ and *µ*_*i*_ in every interval, together with their OU parameters. The exchange did not change the log prior density of each state but lowered its log (FBD)-likelihood by 44 to 516. We then resumed all four runs for 5 million iterations from these swapped states. Across the 29 intervals, the posterior probability of a negative diversification rate (*λ*_*i*_ *< µ*_*i*_) computed from all samples post-resume, including the ones immediately after the swap, differed from that in the original chains by at most 0.10, and in the open interval older than 320 Mya it was 0.003 in the original chains and *<* 0.001 in the resumed chains, suggesting that the data strongly disfavor the swapped configuration.

**Figure 6:**
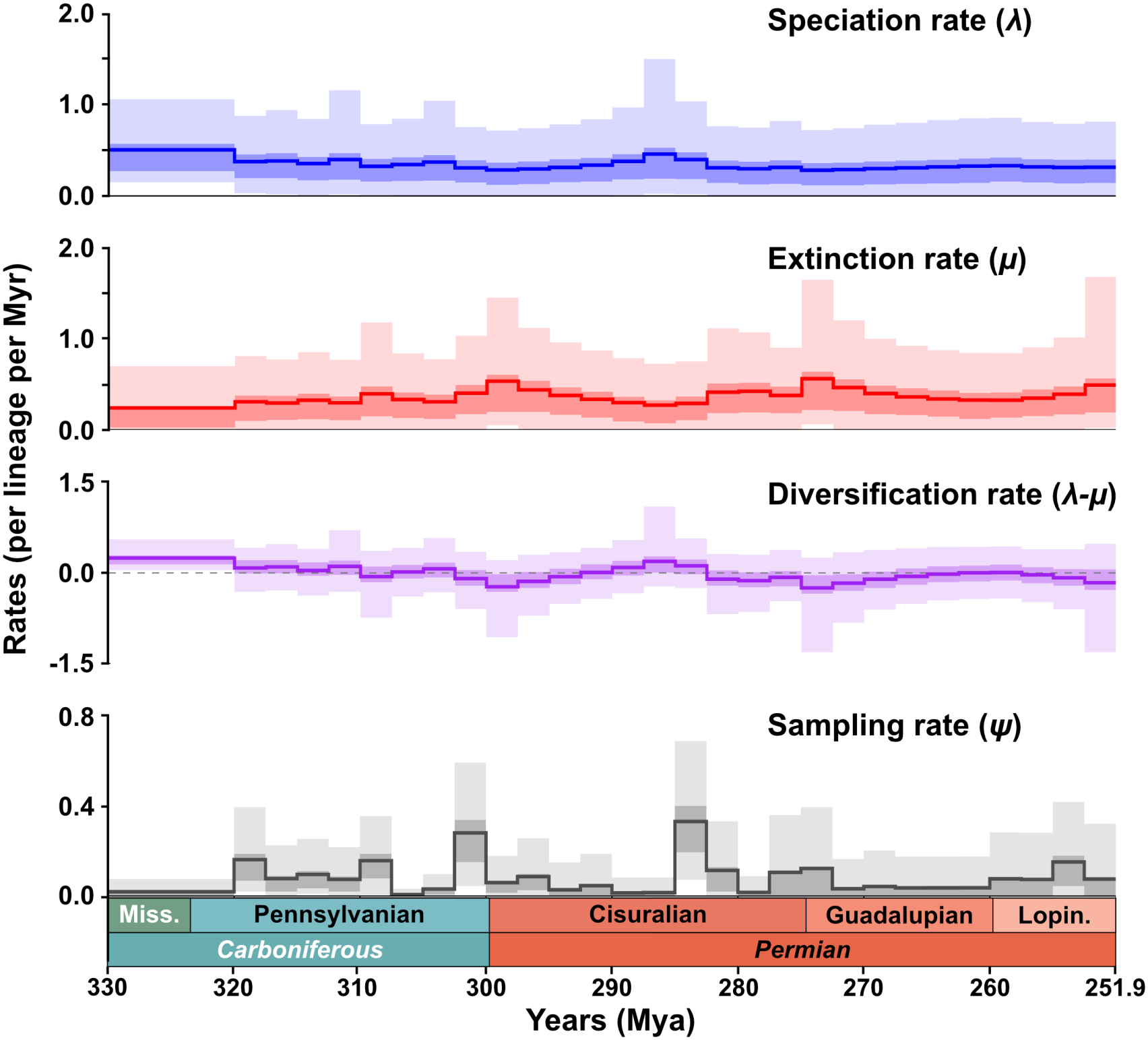
Speciation, extinction, diversification, and sampling rates resulting from the Palaeodictyopterida analysis. Lines denote posterior medians, darker shades denote 50% credible intervals, and lighter shades denote 95% credible intervals. Abbreviations: Miss: Mississippian, Lopin: Lopingian

The posterior median speciation rate was between 0.28 and 0.40 events per lineage per Myr in 27 of the 29 intervals, and higher in the open interval older than 320 Mya (0.51) and at 285–287.5 Mya (0.46), whereas the posterior median extinction rate varied between 0.24 and 0.56, with 95% CIs overlapping across intervals. The 95% CI of the diversification rate excluded zero only in the open interval older than 320 Mya (median 0.25, 95% CI 0.05–0.55). The estimated sampling rate fluctuated considerably throughout time, partly reflecting the typical lagerstätten nature of insect fossil records (Labandeira & Sepkoski, 1993).

Under the enumeration of Heath et al. (2014), the Palaeodictyopterida analysis did not converge within the maximum cutoff of 1 billion iterations (maximum 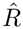 of 1.018, minimum bulk ESS of 231), but its posterior again differed markedly from that of our implementation of the corrected enumeration (Supplementary Figure S9). The posterior medians of *λ*_*i*_ and *µ*_*i*_ were about 2.0–4.2 and 2.1–3.7 times those of the corrected enumeration, with the lower of *ψ*_*i*_ being 0.07–0.95 times, while the number of sampled ancestors fell from a median of 174 (95% CI 106–242) to 21 (10–33).

## 6 Discussion

Here, we presented the corrected formulation of the unresolved FBD process first explored by Heath et al. (2014) and implemented it with several methodological enhancements (see Supplementary Material 1). Using three datasets (Cetacea, Emydidae, Palaeodictyopterida) varying in size and scope, we showed that our implementation of the method achieves reasonable runtimes even when hundreds of unresolved taxa are present, and in cases where the speciation, extinction, and sampling rates are estimated through the FBD process with only fossil occurrence data present. When morphological data are abundant and well described by the morphological model, total-evidence tip dating can place fossils and date the tree jointly. In practice, however, an adequate fit of morphological models is often difficult to achieve, morphological matrices are not available for many clades, and even where characters can be scored, how to code them often lacks consensus and requires expert knowledge (Lee & Palci, 2015; Khakurel et al., 2024). Ideally, every available fossil would be scored for morphological characters under an adequate model, but this is rarely the case, and depending on the objective, placing fossils in the tree may not be one’s primary goal; instead, one may use them only to estimate divergence times among extant species, infer diversification rates, or address other related questions. In such cases, the unresolved FBD process offers a practical and computationally efficient solution, as it uses every fossil that can be assigned to a clade and integrates over its attachments without requiring morphological data.

While the unresolved FBD process (or the FBD process in general) provides a model-based frame-work for divergence time estimation without relying on subjective node priors, it is important to note that it does not solve the fundamental identifiability problem of the molecular clock and thus its statistical consequences (Rannala & Yang, 2007; dos Reis & Yang, 2013). Moreover, it shares the same caveats with all other model-based approaches: models can only be an approximation to the truth. For example, the true trajectory of speciation, extinction, and sampling rates need not be piecewise constant, nor shared across all lineages at the same time. While the FBD process is identifiable without conditioning (Truman et al., 2025) (though not under conditioning on *S*_*n*_ or *E* when the swap condition of Theorem 3 holds), this nonparametric identifiability does not guarantee stable estimation. With finite data, short intervals hold few or no events, which can give each per-bin rate a wide posterior. Fine binning coupled with OU smoothing was one way we came up with to overcome this, with a bias–variance tradeoff. More generally, narrower interval width might be more realistic, but lower variance (at the cost of bias) and shorter runtime favor wider ones, so intervals should be chosen appropriately with regard to the dataset in hand, and any patterns in rate changes finer than the data can resolve cannot be expected to be recovered at any interval width.

As with all joint analyses of paleontological and neontological data, the FBD process is also exposed to conflicts in species concepts and assumed speciation modes between extant and fossil species (Simpson, 1951; Ezard et al., 2012; Silvestro et al., 2018; Stadler et al., 2018), which require caution when interpreting the results, particularly when the taxonomic practices applied to fossil and extant species are expected to differ greatly. Moreover, the original formulation of the FBD process by Stadler (2010) and its subsequent implementations (Gavryushkina et al., 2014; Heath et al., 2014) treat each sampled fossil as a distinct labeled taxon, and so cannot accommodate another occurrence of a species already in the tree, whether as a fossil or as an extant tip. All three of our analyses therefore represented each species by a single occurrence and excluded fossils of extant species, which discards sampling events and makes *ψ* the rate at which species are recorded, and this can be problematic for taxa with high fossil preservation potential.

For estimating diversification rates from fossil occurrences alone, as in our Palaeodictyopterida analysis, the widely used program PyRate (Silvestro et al., 2014, 2019) can use multiple occurrences of each species, but its birth-death model counts the speciation and extinction events of the observed species only, as if every species had been sampled, and its performance degrades at low sampling rates and high turnover (Warnock et al., 2020), and when singletons dominate the fossil record (Černý et al., 2022). The stratigraphic range FBD (Stadler et al., 2018) instead accounts for unsampled species while using repeated occurrences under budding, bifurcating, and anagenetic speciation, but its existing implementations only assume budding speciation, are restricted to constant rates (Warnock et al., 2020), remain experimental (Flannery-Sutherland et al., 2022), or require sampling the topology (Stolz, 2024; Stolz et al., 2025). Under the budding speciation model, each species attaches asymmetrically to its ancestor, so the symmetric configurations behind the overcounting corrected here do not arise, whereas bifurcating or mixed speciation will require symmetric splits to be enumerated without redundant counting. Extending the unresolved formulation presented here to the stratigraphic range FBD under these speciation modes would remove the restriction to a single occurrence per species.

Under the scenario where a rich morphological matrix is available for the clade of interest (at least for some taxa), it might be desirable to integrate it into the analysis depending on the study objective. Moreover, when one does not have a robust backbone phylogeny, one might wish to integrate over uncertainties in the backbone topology. With well-designed definitions of and proposals for the attachment zones of unresolved fossils, it would be possible to accommodate tree proposals for the resolved portion of the phylogeny while integrating out only the attachments of fossils that lack morphological information using the unresolved FBD formulation. Another source of information that we did not account for is fossils that have not been assigned to the species level. In some cases, the oldest fossil of a clade, which plays a crucial role in determining the timeline of that clade, may not be identified to the species level. Such a fossil can enter the FBD process with a clade assignment at the level to which it is identified (Barido-Sottani et al., 2023), but only if it is confidently known to represent a species distinct from every other sampled taxon. A specimen identified only to genus may instead be another occurrence of a species already in the tree, which the standard FBD process cannot accommodate, so this information cannot be readily integrated without special treatment. The “occurrence birth-death process” (OBDP), introduced by Vaughan et al. (2019) and Gupta et al. (2020) in a phylodynamic context and later extended to a macroevolutionary setting (Andréoletti et al., 2022), aims to incorporate such unlabeled occurrences into the FBD framework and could, in principle, be integrated into the unresolved FBD framework.

It is widely acknowledged that speciation and extinction rates are heterogeneous throughout the tree of life (Morlon et al., 2011; Kopperud et al., 2026). Barido-Sottani and Morlon (2026) recently extended the FBD process to incorporate a multi-type branching process in which speciation, extinction, and sampling rates may vary across the tree (Barido-Sottani et al., 2020b). Extending the unresolved FBD formulation to this setting would allow uncertainty in fossil placement to be integrated while accommodating heterogeneity in diversification and sampling rates, although it would also raise challenges in how rate shifts should be handled at unresolved lineages.

## 7. Appendix

### 7.1 Proofs of Theorems 1 and 2

#### Lemma 1.

*Each resolved labeled tree T is realized by exactly* 2^*m*−*c*(T )^ *distinct pairs* (**z, h**), *where the resolution* **h** = (*h*_*f*_ )_*f*_ *assigns each fossil f the edge or stalk h*_*f*_ *it attaches to, all of identical density*.

*Proof*. Distinct *f* -clades are independent, so it suffices to count pairs within one *f* -clade and multiply. Note that an *f* -clade may contain SAs, but they do not participate in any bifurcation event and thus cannot contribute to overcounting due to multiple attachment realizations leading to the same resolved labeled tree. We fix an *f* -clade of *c* tip fossils and use induction on *c*. For *c* = 1, there is only one way for its single fossil to attach to its backbone branch. For *c* ≥ 2, the first bifurcation event joins two subclades of sizes *a* and *c* − *a*. Either subclade can attach to a stalk of the other, and both possible attachments result in the same resolved labeled tree. The subclades are realized independently in 2^*a*−1^ and 2^*c*−*a*−1^ ways by the induction, giving 2 · 2^*a*−1^ · 2^*c*−*a*−1^ = 2^*c*−1^. Defining *c*_*j*_ as count of the tip fossils of the *f* -clade *j*, and multiplying over *f* -clades and using Σ_*j*_ *c*_*j*_ = *m* gives 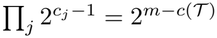. All such pairs share **x, y**, and differ only by permuting the attachment ages **z** among each *f* -clade’s tip fossils. Since the density does not depend on which fossil owns which attachment age, the density is identical across them.

#### Proof of Theorem 1

First, fix **z**. Every resolution compatible with **z** and the clade assignments has the same density, and the enumeration of Heath et al. (2014) gives each of them weight one, so the marginalization is the integration of *f* (*T*_*lb*_ | *t*_*or*_ = *x*_0_) over all pairs (**z, h**) where **h** is such a resolution. Group these pairs by the resolved labeled tree *T* ∈ *U*_**z**_ R(**z**) each one realizes. A resolution is compatible with the clade assignments exactly when the *T* it realizes is, so every pair realizing a compatible *T* is counted. By Lemma 1, every compatible *T* is realized by exactly 2^*m*−*c*(T )^ such pairs, of equal density. The factor 2^*m*−*c*(T )^ can thus be pulled out of the integral, so the enumeration places total weight 2^*m*−*c*(T )^ on each distinct compatible *T* .

#### Lemma 2.

*For any backbone edge e lying in at least one attachment zone, the attachment zones containing e form a chain under inclusion. Thus, Z*(*e*) *exists and is unique*.

*Proof*. Two monophyletic clades of *T*_*bb*_ are nested or disjoint (Semple and Steel (2003), Theorem 3.5.2). Within one clade, its crown and stem attachment zones are disjoint, and both lie in its total attachment zone. If a clade with crown node *κ*^′^ is nested strictly within one with crown node *κ*, then *κ*^′^ and the backbone edge rootward of it descend from *κ*, and since *κ*^′^ is strictly nested, that edge terminates at a node still within the crown of *κ*, so every attachment zone of *κ*^′^ lies in the crown attachment zone of *κ*, and thus in its total attachment zone, and is disjoint from its stem attachment zone. The attachment zones of disjoint clades are disjoint. Distinct attachment zones are therefore nested or disjoint, so any two attachment zones that both contain *e* are nested, and the attachment zones containing *e* form a chain. This chain is finite, so it has a unique smallest element, which is *Z*(*e*).

#### Lemma 3.

*Given* **Z** ∈ *Ƶ, the resolutions with attachment zones* **Z** *are in bijection with the tuples* (*h*_*f*_ )_*f*_ *in which h*_*f*_ *is one of the b*_*f*_ *edges or one of the s*_*f*_ *stalks, so that each fossil f has γ*_*f*_ (**Z**) *possible attachments. Every such resolution is compatible with the clade assignments*.

*Proof*. Say that a resolution has attachment zones **Z** ∈ *Ƶ* when *Z*(*e*_*f*_ ) = *Z*_*f*_ for every fossil *f* . A resolution with attachment zones **Z** assigns each fossil *f* an attachment *h*_*f*_ . Attaching *f* to a backbone edge *e* gives *e*_*f*_ = *e*, so *Z*(*e*) = *Z*_*f*_, and attaching it to the stalk of *f*^*’*^ gives 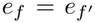, so 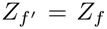. Thus, *h*_*f*_ is one of the *b*_*f*_ edges or one of the *s*_*f*_ stalks. Conversely, let (*h*_*f*_ )_*f*_ be such a tuple. A fossil attached to another fossil’s stalk has the strictly smaller attachment age, so following the attachments from *f* ends at some fossil attached to a backbone edge *e*; every fossil on that chain has *e*_*f*_ = *e*, and 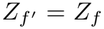 whenever *f* attaches to the stalk of *f*, by the definition of *s*_*f*_, so *Z*(*e*_*f*_ ) = *Z*(*e*) = *Z*_*f*_ . The tuple therefore gives a resolution with attachment zones **Z**. Finally, *e*_*f*_ ∈ *Z*(*e*_*f*_ ) = *Z*_*f*_ ⊆ *A*_*f*_, so every such resolution is compatible with the clade assignments.

#### Proof of Theorem 2

A compatible resolution has *e*_*f*_ ∈ *A*_*f*_, so *e*_*f*_ lies in at least one attachment zone, and by Lemma 2, *Z*(*e*_*f*_ ) exists and *Z*(*e*_*f*_ ) ⊆ *A*_*f*_ ; the resolution therefore has attachment zones **Z** = (*Z*(*e*_*f*_ ))_*f*_ for a single **Z** ∈ *Ƶ*. By Lemma 3, the sum over *Ƶ* enumerates every compatible resolution exactly once, and the resolutions with attachment zones **Z** are in bijection with the tuples of attachments, so their total weight is the product over the fossils of the total weight of the attachments compatible to each other, which is the bracket of Equation (4) without its factor 2^*m*^. Under 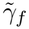, an attachment of a tip fossil to a stalk weighs 1*/*2, and every other attachment weighs 1. A pair (**z, h**) realizing *T* attaches one tip fossil of each *f* -clade to a backbone edge and the remaining *c*_*j*_ − 1 to tip fossil stalks, and Σ_*j*_ (*c*_*j*_ − 1) = *m* − *c*(T ), so it weighs 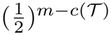. By Lemma 1 there are 2^*m*−*c*(*T* )^ of them, so *T* receives total weight 1.

#### 7.2 Swap invariance

For rates *λ*(*t*), *µ*(*t*), and *ψ*(*t*) varying through time, *p*(*t*) and *Q*(*t*) are the solutions of the master equations of the main text with the per-bin rates replaced by *λ*(*t*), *µ*(*t*), and *ψ*(*t*), with *p*(0) = 1 − *ρ* and *Q*(0) = 1, and 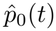 is *p*(*t*) with *ψ* ≡ 0; the density of an oriented tree is Equation (1) with 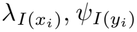, and 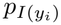 replaced by *λ*(*x*_*i*_), *ψ*(*y*_*i*_), and *p*(*y*_*i*_) (MacPherson et al., 2022; Truman et al., 2025). The skyline FBD model explored in the main text is a simple, tractable subcase of this general model.

##### Theorem 3

(Swap invariance). *Let ρ* = 1, *and let λ*(*t*), *µ*(*t*), *and ψ*(*t*) *be non-negative and Lebesgue-integrable on* (0, *x*_0_), *and positive at almost every t* ∈ (0, *x*_0_). *Fix n* ≥ 0. *Swapping λ*(*t*) *and µ*(*t*) *for all t* ∈ (0, *x*_0_) *leaves the FBD density of every tree with n extant tips conditioned on S*_*n*_ *unchanged if and only if λ*(*t*) = *c µ*(*t*) *for almost every t* ∈ (0, *x*_0_), *for some constant c >* 0. *The same holds for the density of every tree without extant tips conditioned on E*.

*Proof*. First assume *λ*(*t*) = *c µ*(*t*) for almost every *t* ∈ (0, *x*_0_). We use the subscript *σ* to denote the quantities after swapping *λ*(*t*) and *µ*(*t*). For example, *p*_*σ*_(*t*) is *p*(*t*) after the swapping. We show that under this assumption, swapping *λ*(*t*) and *µ*(*t*) gives *p*_*σ*_ = *cp*, 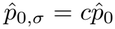, and *Q*_*σ*_ = *Q*, which implies swap invariance under *S*_*n*_ or *E* conditioning. Trivially, *cp* and *p*_*σ*_ solve 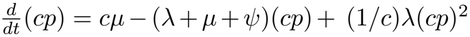 and 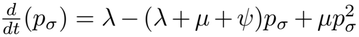 . It is easy to note that these two agree since *λ* = *cµ* for almost every *t*, and both equal 0 at *t* = 0 since *ρ* = 1. Since the rates are only Lebesgue-integrable, *cp* and *p*_*σ*_ are extended solutions of this equation, i.e., absolutely continuous on [0, *x*_0_] and satisfying it at almost every *t*, and as *ρ* = 1 *>* 0, this equation has a unique extended solution on [0, *x*_0_] (Truman et al., 2025, Proposition 1), so *p*_*σ*_ = *cp*. The same argument holds for 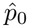. For *Q*, the master equation of *g* gives 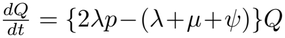 with 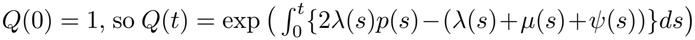, and likewise 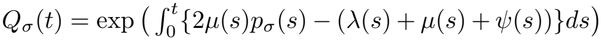. Since *µp*_*σ*_ = *cµp* = *λp* at almost every *t*, the two integrands agree almost everywhere, and *Q*_*σ*_ = *Q*.

Conversely, assume that the swap leaves the density conditioned on *C*, which is either *S*_*n*_ for some *n* or *E*, unchanged for every tree compatible with such a conditioning scheme. Write *f* (*T* ) = *f* (*T* | *t*_*or*_ = *x*_0_) for the FBD density of a tree *T* and *f*_*σ*_(*T* ) for the density after the swap, so that the density conditioned on *C* is *f* (*T* )*/P* (*C*). Let *T* be such a tree with a lineage alive at age *x*, and let *T* ^′^ be a tree obtained by adding to that lineage, at age *x*, an edge ending in a fossil tip of age *y < x*, so that *T* ^′^ is compatible with the same conditioning scheme. The added edge contributes to the FBD density the factor *λ*(*x*)*Q*(*x*) for the branching event and *ψ*(*y*)*p*(*y*)*/Q*(*y*) for the fossil sampling event, so *f* (*T* ^′^)*/f* (*T* ) = *λ*(*x*)*Q*(*x*)*ψ*(*y*)*p*(*y*)*/Q*(*y*). As the conditioned densities of *T* and *T* ^′^ are unchanged by the assumption, *f*_*σ*_(*T* )*/P*_*σ*_(*C*) = *f* (*T* )*/P* (*C*) and *f*_*σ*_(*T* ^′^)*/P*_*σ*_(*C*) = *f* (*T* ^′^)*/P* (*C*), so *f*_*σ*_(*T* )*/f* (*T* ) = *f*_*σ*_(*T* ^′^)*/f* (*T* ^′^). Since *λ*(*x*) *>* 0 and *ψ*(*y*) *>* 0 at almost every *x* and *y*, dividing the two sides gives, for almost every *x > y*,

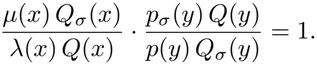

The first factor depends only on *x* and the second factor only depends on *y*, so both must be positive constants almost everywhere, i.e., *cµQ*_*σ*_ = *λQ* and *c*^′^*p*_*σ*_*Q* = *pQ*_*σ*_ with *cc*^′^ = 1 and *c, c*^′^ *>* 0, which gives *µp*_*σ*_ = *λp*. Then 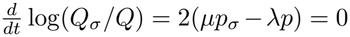 with *Q*_*σ*_(0) = *Q*(0) = 1, so *Q*_*σ*_ = *Q* and *λ* = *cµ* at almost every *t*.

## Author contributions

Conceptualization: WL; Data curation: WL; Formal analysis: WL; Funding acquisition: RN; Investigation: WL; Methodology: WL, LYR, JKL; Software: WL, LYR; Supervision: JPH, RN; Visualization: WL; Writing – original draft: WL; Writing – review & editing: WL, LYR, JKL, JPH, RN.

## Conflict of interest

None declared.

## Funding

This work was supported by the National Institutes of Health [NIH R35 GM153400 to R.N.].

## Acknowledgments

Claude Code, running primarily Opus 5.5 and, to a lesser extent, Fable 5.1, was used to write and refine the program code, design simulation experiments and validation procedures, prepare code for generating figures, and conduct a mechanical audit of fossil occurrence records against the PBDB and published literature. ChatGPT (Astra 6.2) and Claude (Fable 5.1) were used to identify potential logical and notational gaps in the proofs and to improve their rigor. The authors directed all AI-assisted work and reviewed, tested, and validated all code, analyses, figures, and audit results produced with these tools. The authors take full responsibility for the content of this article.

## Data availability statement

The data, input files, analysis configurations, and outputs for the empirical analyses, and the scripts for the SBC, are available from the Dryad repository (Lim et al., 2026): https://doi.org/10.5061/dryad.w6m905r4v. The original Cetacea and Emydidae sequence data are available from the sources cited in the text.

Fossil occurrence records were obtained from the Paleobiology Database as described in the “Empirical datasets” section and Supplementary Material 2. The source code and Linux binary of the implemented method are also available from: https://github.com/wnsplim/UFBD.

## Notes

### Competing Interest Statement

The authors have declared no competing interest.

### Summary of Updates

More treatments on swap, and improved empirical analyses

## References

Andréoletti J, Zwaans A, Warnock RCM, Aguirre-Fernández G, Barido-Sottani J, Gupta A, Stadler T, Manceau M. 2022. The occurrence birth–death process for combined-evidence analysis in macroevolution and epidemiology. Systematic Biology. 71 (6):1440–1452. 10.1093/sysbio/syac037

Angelis K, Álvarez-Carretero S, dos Reis M, Yang Z. 2018. An evaluation of different partitioning strategies for Bayesian estimation of species divergence times. Systematic Biology. 67 (1):61–77. 10.1093/sysbio/syx061

Barba-Montoya J, dos Reis M, Yang Z. 2017. Comparison of different strategies for using fossil calibrations to generate the time prior in Bayesian molecular clock dating. Molecular Phylogenetics and Evolution. 114 :386–400. 10.1016/j.ympev.2017.07.005

Barba-Montoya J, Tao Q, Kumar S. 2020. Using a GTR+ff substitution model for dating sequence divergence when stationarity and time-reversibility assumptions are violated. Bioinformatics. 36 (Supplement 2):i884–i894. 10.1093/bioinformatics/btaa820

Barido-Sottani J, Aguirre-Fernández G, Hopkins MJ, Stadler T, Warnock R. 2019. Ignoring stratigraphic age uncertainty leads to erroneous estimates of species divergence times under the fossilized birth-death process. Proceedings of the Royal Society B: Biological Sciences. 286 (1902):20190685. 10.1098/rspb.2019.0685

Barido-Sottani J, Morlon H. 2026. Integrating fossils samples with heterogeneous diversification rates: A combined multi-type fossilized birth-death model. Systematic Biology. 75 (6):1175–1195. 10.1093/sysbio/syag011

Barido-Sottani J, Pohle A, De Baets K, Murdock D, Warnock RCM. 2023. Putting F into FBD analysis: Tree constraints or morphological data? Palaeontology. 66 (6):e12679. 10.1111/pala.12679

Barido-Sottani J, van Tiel NMA, Hopkins MJ, Wright DF, Stadler T, Warnock RC. 2020a. Ignoring fossil age uncertainty leads to inaccurate topology and divergence time estimates in time calibrated tree inference. Frontiers in Ecology and Evolution. 8 :183. 10.3389/fevo.2020.00183

Barido-Sottani J, Vaughan TG, Stadler T. 2020b. A multitype birth–death model for Bayesian inference of lineage-specific birth and death rates. Systematic Biology. 69 (5):973–986. 10.1093/sysbio/syaa016

Beaulieu JM. 2021. The problem with clade-specific sampling fractions.HiSSE Vignette. https://speciationextinction.info/articles/Clade-specific-sampling.html

Benjamini Y, Hochberg Y. 1995. Controlling the false discovery rate: A practical and powerful approachto multiple testing. Journal of the Royal Statistical Society: Series B (Methodological). 57 (1):289–300. 10.1111/j.2517-6161.1995.tb02031.x

Benjamini Y, Yekutieli D. 2001. The control of the false discovery rate in multiple testing under dependency. The Annals of Statistics. 29 (4):1165–1188. 10.1214/aos/1013699998

Benton MJ, Donoghue PCJ. 2007. Paleontological evidence to date the tree of life. Molecular Biologyand Evolution. 24 (1):26–53. 10.1093/molbev/msl150

Biek R, Pybus OG, Lloyd-Smith JO, Didelot X. 2015. Measurably evolving pathogens in the genomic era. Trends in Ecology & Evolution. 30 (6):306–313. 10.1016/j.tree.2015.03.009

Bisconti M, Pellegrino L, Carnevale G. 2023. The chronology of mysticete diversification (Mammalia,Cetacea, Mysticeti): Body size, morphological evolution and global change. Earth-Science Reviews. 239 :104373. 10.1016/j.earscirev.2023.104373

Bromham L, Duchêne S, Hua X, Ritchie AM, Duchêne DA, Ho SYW. 2018. Bayesian molecular dating: Opening up the black box. Biological Reviews. 93 (2):1165–1191. 10.1111/brv.12390

Brown JW, Smith SA. 2018. The past sure is tense: On interpreting phylogenetic divergence timeestimates. Systematic Biology. 67 (2):340–353. 10.1093/sysbio/syx074

Capobianco A, Darlim G, Höhna S. 2026. How to date a molecular phylogeny: Comparison of effectivepriors between node calibration and fossilized birth-death. Proceedings of the Royal Society B: Biological Sciences. 293 (2070):20253255. 10.1098/rspb.2025.3255

Černý D, Madzia D, Slater GJ. 2022. Empirical and methodological challenges to the model-basedinference of diversification rates in extinct clades. Systematic Biology. 71 (1):153–171. 10.1093/sysbio/syab045

Churchill GA, von Haeseler A, Navidi WC. 1992. Sample size for a phylogenetic inference. Molecular Biology and Evolution. 9 (4):753–769. 10.1093/oxfordjournals.molbev.a040757

Claramunt S. 2022. CladeDate: Calibration information generator for divergence time estimation. Meth-ods in Ecology and Evolution. 13 (11):2331–2338. 10.1111/2041-210X.13977

Claramunt S, Cracraft J. 2015. A new time tree reveals Earth history’s imprint on the evolution ofmodern birds. Science Advances. 1 (11):e1501005. 10.1126/sciadv.1501005

Colston TJ, Kulkarni P, Jetz W, Pyron RA. 2020. Phylogenetic and spatial distribution of evolutionary diversification, isolation, and threat in turtles and crocodilians (non-avian archosauromorphs). BMCEvolutionary Biology. 20 :81. 10.1186/s12862-020-01642-3

Cook SR, Gelman A, Rubin DB. 2006. Validation of software for Bayesian models using posterior quantiles. Journal of Computational and Graphical Statistics. 15 (3):675–692. 10.1198/106186006X136976

Demotte P, Panchaksaram M, Kumarasinghe H, Ly-Trong N, dos Reis M, Minh BQ. 2025. IQ2MC:A new framework to infer phylogenetic time trees using IQ-TREE 3 and MCMCTree with mixture models. EcoEvoRxiv. 10.32942/X2CD2X

Didier G, Fau M, Laurin M. 2017. Likelihood of tree topologies with fossils and diversification rateestimation. Systematic Biology. 66 (6):964–987. 10.1093/sysbio/syx045

Didier G, Laurin M. 2020. Exact distribution of divergence times from fossil ages and tree topologies.Systematic Biology. 69 (6):1068–1087. 10.1093/sysbio/syaa021

Didier G, Laurin M. 2024. Testing extinction events and temporal shifts in diversification and fossilization rates through the skyline fossilized birth-death (FBD) model: The example of some mid-Permian synapsid extinctions. Cladistics. 40 (3):282–306. 10.1111/cla.12577

Didier G, Royer-Carenzi M, Laurin M. 2012. The reconstructed evolutionary process with the fossil record. Journal of Theoretical Biology. 315 :26–37. 10.1016/j.jtbi.2012.08.046

dos Reis M, Donoghue PCJ, Yang Z. 2016. Bayesian molecular clock dating of species divergences inthe genomics era. Nature Reviews Genetics. 17 (2):71–80. 10.1038/nrg.2015.8

dos Reis M, Yang Z. 2011. Approximate likelihood calculation on a phylogeny for Bayesian estimationof divergence times. Molecular Biology and Evolution. 28 (7):2161–2172. 10.1093/molbev/msr045

dos Reis M, Yang Z. 2013. The unbearable uncertainty of Bayesian divergence time estimation. Journal of Systematics and Evolution. 51 (1):30–43. 10.1111/j.1759-6831.2012.00236.x

dos Reis M, Zhu T, Yang Z. 2014. The impact of the rate prior on Bayesian estimation of divergencetimes with multiple loci. Systematic Biology. 63 (4):555–565. 10.1093/sysbio/syu020

Drummond AJ, Ho SYW, Phillips MJ, Rambaut A. 2006. Relaxed phylogenetics and dating withconfidence. PLoS Biology. 4 (5):e88. 10.1371/journal.pbio.0040088

Drummond AJ, Pybus OG, Rambaut A, Forsberg R, Rodrigo AG. 2003. Measurably evolving populations. Trends in Ecology & Evolution. 18 (9):481–488. 10.1016/S0169-5347(03)00216-7

du Plessis L. 2016. Understanding the spread and adaptation of infectious diseases using genomic sequencing data. Doctoral Thesis. Zürich: ETH Zürich. 10.3929/ethz-a-010866338

Dunn OJ. 1961. Multiple comparisons among means. Journal of the American Statistical Association.56 (293):52–64. 10.1080/01621459.1961.10482090

Ezard THG, Pearson PN, Aze T, Purvis A. 2012. The meaning of birth and death (in macroevolutionary birth–death models). Biology Letters. 8 (1):139–142. 10.1098/rsbl.2011.0699

Fabreti LG, Höhna S. 2023. Nucleotide substitution model selection is not necessary for Bayesianinference of phylogeny with well-behaved priors. Systematic Biology. 72 (6):1418–1432. 10.1093/sysbio/syad041

Featherstone LA, Di Giallonardo F, Holmes EC, Vaughan TG, Duchêne S. 2021. Infectious disease phylodynamics with occurrence data. Methods in Ecology and Evolution. 12 (8):1498–1507. 10.1111/2041-210X.13620

Felsenstein J. 1981. Evolutionary trees from DNA sequences: A maximum likelihood approach. Journal of Molecular Evolution. 17 (6):368–376. 10.1007/BF01734359

Flannery-Sutherland JT, Silvestro D, Benton MJ. 2022. Global diversity dynamics in the fossil recordare regionally heterogeneous. Nature Communications. 13 :2751. 10.1038/s41467-022-30507-0

Foote M. 1996. On the probability of ancestors in the fossil record. Paleobiology. 22 (2):141–151. 10.1017/S0094837300016146

Ford D, Matsen FA, Stadler T. 2009. A method for investigating relative timing information on phylogenetic trees. Systematic Biology. 58 (2):167–183. 10.1093/sysbio/syp018

Gavryushkina A, Welch D, Stadler T, Drummond AJ. 2014. Bayesian inference of sampled ancestortrees for epidemiology and fossil calibration. PLoS Computational Biology. 10 (12):e1003919. 10.1371/journal.pcbi.1003919

Gelman A, Carlin JB, Stern HS, Dunson DB, Vehtari A, Rubin DB. 2013. Bayesian Data Analysis.3rd ed. Boca Raton: CRC Press.

Gelman A, Rubin DB. 1992. Inference from iterative simulation using multiple sequences. Statistical Science. 7 (4):457–472. 10.1214/ss/1177011136

Gernhard T. 2008. The conditioned reconstructed process. Journal of Theoretical Biology. 253 (4):769– 778. 10.1016/j.jtbi.2008.04.005

Geyer CJ. 1991. Markov chain Monte Carlo maximum likelihood. In: Computing Science and Statistics: Proceedings of the 23rd Symposium on the Interface. Interface Foundation of North America. p. 156–163.

Gilks WR, Roberts GO. 1996. Strategies for improving MCMC. In: Gilks WR, Richardson S, Spiegel-halter DJ, editors. Markov Chain Monte Carlo in Practice. London: Chapman and Hall. p. 89–114.

Green PJ. 1995. Reversible jump Markov chain Monte Carlo computation and Bayesian model determination. Biometrika. 82 (4):711–732. 10.1093/biomet/82.4.711

Grimaldi D, Engel MS. 2005. Evolution of the Insects. Cambridge: Cambridge University Press.

Gu X, Fu YX, Li WH. 1995. Maximum likelihood estimation of the heterogeneity of substitution rateamong nucleotide sites. Molecular Biology and Evolution. 12 (4):546–557. 10.1093/oxfordjournals.molbev.a040235

Guindon S. 2010. Bayesian estimation of divergence times from large sequence alignments. Molecular Biology and Evolution. 27 (8):1768–1781. 10.1093/molbev/msq060

Guindon S. 2018. Accounting for calibration uncertainty: Bayesian molecular dating as a “doublyintractable” problem. Systematic Biology. 67 (4):651–661. 10.1093/sysbio/syy003

Guindon S. 2019. Rates and rocks: Strengths and weaknesses of molecular dating methods. Frontiersin Genetics. 11 :526. 10.3389/fgene.2020.00526

Gupta A, Manceau M, Vaughan T, Khammash M, Stadler T. 2020. The probability distribution of the reconstructed phylogenetic tree with occurrence data. Journal of Theoretical Biology. 488 :110115.10.1016/j.jtbi.2019.110115

Heads M. 2012. Bayesian transmogrification of clade divergence dates: A critique. Journal of Biogeography. 39 (10):1749–1756. 10.1111/j.1365-2699.2012.02784.x

Heath TA, Huelsenbeck JP, Stadler T. 2014. The fossilized birth-death process for coherent calibrationof divergence-time estimates. Proceedings of the National Academy of Sciences of the United States of America. 111 (29):E2957–E2966. 10.1073/pnas.1319091111

Heath TA, Moore BR. 2014. Bayesian inference of species divergence times. In: Chen MH, Kuo L,Lewis PO, editors. Bayesian Phylogenetics: Methods, Algorithms, and Applications. Boca Raton: CRC Press. p. 277–318.

Heckeberg NS, Capobianco A, Khakurel B, Darlim G, Höhna S. 2026. Practical guide and review of fossil tip-dating in phylogenetics. Systematic Biology. 75 (1):156–192. 10.1093/sysbio/syaf050

Hedman MM. 2010. Constraints on clade ages from fossil outgroups. Paleobiology. 36 (1):16–31. 10.1666/0094-8373-36.1.16

Heled J, Drummond AJ. 2012. Calibrated tree priors for relaxed phylogenetics and divergence time estimation. Systematic Biology. 61 (1):138–149. 10.1093/sysbio/syr087

Ho SYW, Phillips MJ. 2009. Accounting for calibration uncertainty in phylogenetic estimation ofevolutionary divergence times. Systematic Biology. 58 (3):367–380. 10.1093/sysbio/syp035

Höhna S. 2014. Likelihood inference of non-constant diversification rates with incomplete taxon sampling. PLoS One. 9 (1):e84184. 10.1371/journal.pone.0084184

Höhna S, Stadler T, Ronquist F, Britton T. 2011. Inferring speciation and extinction rates under different sampling schemes. Molecular Biology and Evolution. 28 (9):2577–2589. 10.1093/molbev/msr095

Huelsenbeck JP, Rannala B. 2004. Frequentist properties of Bayesian posterior probabilities of phylogenetic trees under simple and complex substitution models. Systematic Biology. 53 (6):904–913.10.1080/10635150490522629

Hug LA, Roger AJ. 2007. The impact of fossils and taxon sampling on ancient molecular dating analyses. Molecular Biology and Evolution. 24 (8):1889–1897. 10.1093/molbev/msm115

Inoue J, Donoghue PCJ, Yang Z. 2010. The impact of the representation of fossil calibrations on Bayesian estimation of species divergence times. Systematic Biology. 59 (1):74–89. 10.1093/sysbio/syp078

Khakurel B, Grigsby C, Tran TD, Zariwala J, Höhna S, Wright AM. 2024. The fundamental role of character coding in Bayesian morphological phylogenetics. Systematic Biology. 73 (5):861–871. 10.1093/sysbio/syae033

Kishino H, Thorne JL, Bruno WJ. 2001. Performance of a divergence time estimation method under a probabilistic model of rate evolution. Molecular Biology and Evolution. 18 (3):352–361. 10.1093/oxfordjournals.molbev.a003811

Kobert K, Stamatakis A, Flouri T. 2017. Efficient detection of repeating sites to accelerate phylogenetic likelihood calculations. Systematic Biology. 66 (2):205–217. 10.1093/sysbio/syw075

Kopperud BT, Capobianco A, Clarke JT, Palazzesi L, Höhna S. 2026. The nature and prevalence of diversification rate shifts across the tree of life. Evolution Letters. 10 (2):217–227. 10.1093/evlett/qrag005

Kubo T, Iwasa Y. 1995. Inferring the rates of branching and extinction from molecular phylogenies.Evolution. 49 (4):694–704. 10.1111/j.1558-5646.1995.tb02306.x

Labandeira CC, Sepkoski Jr. JJ. 1993. Insect diversity in the fossil record. Science. 261 (5119):310–315. 10.1126/science.11536548

Lee MSY, Palci A. 2015. Morphological phylogenetics in the genomic age. Current Biology. 25 (19):R922– R929. 10.1016/j.cub.2015.07.009

Lee MSY, Skinner A. 2011. Testing fossil calibrations for vertebrate molecular trees. Zoologica Scripta.40 (5):538–543. 10.1111/j.1463-6409.2011.00488.x

Lim W, Raskin L, Li J, Huelsenbeck J, Nielsen R. 2026. Data and code from: Estimating divergence times and diversification rates with the unresolved fossilized birth-death process. Dryad. 10.5061/dryad.w6m905r4v

Lingenheil M, Denschlag R, Mathias G, Tavan P. 2009. Efficiency of exchange schemes in replicaexchange. Chemical Physics Letters. 478 (1–3):80–84. 10.1016/j.cplett.2009.07.039

Lloyd GT, Slater GJ. 2021. A total-group phylogenetic metatree for Cetacea and the importance of fossil data in diversification analyses. Systematic Biology. 70 (5):922–939. 10.1093/sysbio/syab002

Louca S, Pennell MW. 2020. Extant timetrees are consistent with a myriad of diversification histories.Nature. 580 (7804):502–505. 10.1038/s41586-020-2176-1

Luo A, Zhang C, Zhou QS, Ho SYW, Zhu CD. 2023. Impacts of taxon-sampling schemes on Bayesian tip dating under the fossilized birth-death process. Systematic Biology. 72 (4):781–801. 10.1093/sysbio/syad011

MacPherson A, Louca S, McLaughlin A, Joy JB, Pennell MW. 2022. Unifying phylogenetic birth–death models in epidemiology and macroevolution. Systematic Biology. 71 (1):172–189. 10.1093/sysbio/syab049

Maliet O, Morlon H. 2022. Fast and accurate estimation of species-specific diversification rates using data augmentation. Systematic Biology. 71 (2):353–366. 10.1093/sysbio/syab055

Marshall CR. 2008. A simple method for bracketing absolute divergence times on molecular phylogeniesusing multiple fossil calibration points. The American Naturalist. 171 (6):726–742. 10.1086/587523

Matschiner M. 2019. Selective sampling of species and fossils influences age estimates under the fossilized birth-death model. Frontiers in Genetics. 10 :1064. 10.3389/fgene.2019.01064

Matschiner M, Musilová Z, Barth JMI, Starostová Z, Salzburger W, Steel M, Bouckaert R. 2017.Bayesian phylogenetic estimation of clade ages supports trans-Atlantic dispersal of cichlid fishes.Systematic Biology. 66 (1):3–22. 10.1093/sysbio/syw076

Mazet N, Morlon H, Fabre PH, Condamine FL. 2023. Estimating clade-specific diversification rates and palaeodiversity dynamics from reconstructed phylogenies. Methods in Ecology and Evolution.14 (10):2575–2591. 10.1111/2041-210X.14195

McGowen MR, Tsagkogeorga G, Álvarez-Carretero S, dos Reis M, Struebig M, Deaville R, Jepson PD, Jarman S, Polanowski A, Morin PA, et al. 2020. Phylogenomic resolution of the cetacean tree of life using target sequence capture. Systematic Biology. 69 (3):479–501. 10.1093/sysbio/syz068

Mendes FK, Bouckaert R, Carvalho LM, Drummond AJ. 2025. How to validate a Bayesian evolutionary model. Systematic Biology. 74 (1):158–175. 10.1093/sysbio/syae064

Meredith RW, Janečka JE, Gatesy J, Ryder OA, Fisher CA, Teeling EC, Goodbla A, Eizirik E, Simão TLL, Stadler T, et al. 2011. Impacts of the Cretaceous terrestrial revolution and KPg extinction on mammal diversification. Science. 334 (6055):521–524. 10.1126/science.1211028

Miasojedow B, Moulines E, Vihola M. 2013. An adaptive parallel tempering algorithm. Journal of Computational and Graphical Statistics. 22 (3):649–664. 10.1080/10618600.2013.778779

Modrák M, Moon AH, Kim S, Bürkner P, Huurre N, Faltejsková K, Gelman A, Vehtari A. 2025. Simulation-based calibration checking for Bayesian computation: The choice of test quantitiesshapes sensitivity. Bayesian Analysis. 20 (2):461–488. 10.1214/23-BA1404

Morlon H, Parsons TL, Plotkin JB. 2011. Reconciling molecular phylogenies with the fossil record.Proceedings of the National Academy of Sciences of the United States of America. 108 (39):16327– 16332. 10.1073/pnas.1102543108

Neal RM. 2003. Slice sampling. The Annals of Statistics. 31 (3):705–767. 10.1214/aos/1056562461

Nee S, May RM, Harvey PH. 1994. The reconstructed evolutionary process. Philosophical Transactions of the Royal Society B: Biological Sciences. 344 (1309):305–311. 10.1098/rstb.1994.0068

Nikolic MC, Warnock RCM, Hopkins MJ. 2025. Combining fossil taxa with and without morphological data improves dated phylogenetic analyses. Biology Letters. 21 (8):20250205. 10.1098/rsbl.2025.0205

Nowak MD, Smith AB, Simpson C, Zwickl DJ. 2013. A simple method for estimating informative node age priors for the fossil calibration of molecular divergence time analyses. PLoS One. 8 (6):e66245.10.1371/journal.pone.0066245

O’Reilly JE, Donoghue PCJ. 2020. The effect of fossil sampling on the estimation of divergence times with the fossilized birth-death process. Systematic Biology. 69 (1):124–138. 10.1093/sysbio/syz037

Okabe T, Kawata M, Okamoto Y, Mikami M. 2001. Replica-exchange Monte Carlo method for the isobaric–isothermal ensemble. Chemical Physics Letters. 335 (5–6):435–439. 10.1016/S0009-2614(01)00055-0

Papaspiliopoulos O, Roberts GO, Sköld M. 2007. A general framework for the parametrization of hierarchical models. Statistical Science. 22 (1):59–73. 10.1214/088342307000000014

Paradis E, Schliep K. 2019. ape 5.0: An environment for modern phylogenetics and evolutionary anal-yses in R. Bioinformatics. 35 (3):526–528. 10.1093/bioinformatics/bty633

Parham JF, Donoghue PCJ, Bell CJ, Calway TD, Head JJ, Holroyd PA, Inoue JG, Irmis RB, Joyce WG, Ksepka DT, et al. 2012. Best practices for justifying fossil calibrations. Systematic Biology. 61 (2):346–359. 10.1093/sysbio/syr107

Prokop J, Engel MS. 2019. Palaeodictyopterida. Current Biology. 29 (9):R306–R309. 10.1016/j.cub.2019.02.056

Pyron PA. 2011. Divergence time estimation using fossils as terminal taxa and the origins of Lissamphibia. Systematic Biology. 60 (4):466–481. 10.1093/sysbio/syr047

Quintero I, Lartillot N, Morlon H. 2024. Imbalanced speciation pulses sustain the radiation of mam-mals. Science. 384 (6699):1007–1012. 10.1126/science.adj2793

Rambaut A. 2018. FigTree v 1.4.4. https://tree.bio.ed.ac.uk/software/figtree/

Rannala B. 1997. Gene genealogy in a population of variable size. Heredity. 78 (4):417–423. 10.1038/hdy.1997.65

Rannala B. 2016. Conceptual issues in Bayesian divergence time estimation. Philosophical Transactions of the Royal Society B: Biological Sciences. 371 (1699):20150134. 10.1098/rstb.2015.0134

Rannala B, Yang Z. 2007. Inferring speciation times under an episodic molecular clock. Systematic Biology. 56 (3):453–466. 10.1080/10635150701420643

Ronquist F, Klopfstein S, Vilhelmsen L, Schulmeister S, Murray DL, Rasnitsyn AP. 2012. A total-evidence approach to dating with fossils, applied to the early radiation of the Hymenoptera. Systematic Biology. 61 (6):973–999. 10.1093/sysbio/sys058

Säilynoja T, Bürkner PC, Vehtari A. 2022. Graphical test for discrete uniformity and its applications in goodness-of-fit evaluation and multiple sample comparison. Statistics and Computing. 32 (2):32.10.1007/s11222-022-10090-6

Semple C, Steel M. 2003. Phylogenetics. Oxford: Oxford University Press.

Seo TK, Kishino H, Thorne JL. 2004. Estimating absolute rates of synonymous and nonsynonymous nucleotide substitution in order to characterize natural selection and date species divergences.Molecular Biology and Evolution. 21 (7):1201–1213. 10.1093/molbev/msh088

Silvestro D, Salamin N, Antonelli A, Meyer X. 2019. Improved estimation of macroevolutionary rates from fossil data using a Bayesian framework. Paleobiology. 45 (4):546–570. 10.1017/pab.2019.23

Silvestro D, Schnitzler J, Liow LH, Antonelli A, Salamin N. 2014. Bayesian estimation of speciation and extinction from incomplete fossil occurrence data. Systematic Biology. 63 (3):349–367. 10.1093/sysbio/syu006

Silvestro D, Warnock RCM, Gavryushkina A, Stadler T. 2018. Closing the gap between palaeontological and neontological speciation and extinction rate estimates. Nature Communications. 9 :5237. 10.1038/s41467-018-07622-y

Simpson GG. 1951. The species concept. Evolution. 5 (4):285–298. 10.1111/j.1558-5646.1951.tb02788.x

Spinks PQ, Thomson RC, McCartney-Melstad E, Shaffer HB. 2016. Phylogeny and temporal diversifi-cation of the New World pond turtles (Emydidae). Molecular Phylogenetics and Evolution. 103 :85–97. 10.1016/j.ympev.2016.07.007

Stadler T. 2009. On incomplete sampling under birth-death models and connections to the sampling-based coalescent. Journal of Theoretical Biology. 261 (1):58–66. 10.1016/j.jtbi.2009.07.018

Stadler T. 2010. Sampling-through-time in birth-death trees. Journal of Theoretical Biology. 267 (3):396–404. 10.1016/j.jtbi.2010.09.010

Stadler T. 2013. How can we improve accuracy of macroevolutionary rate estimates? Systematic Biology. 62 (2):321–329. 10.1093/sysbio/sys073

Stadler T, Gavryushkina A, Warnock RCM, Drummond AJ, Heath TA. 2018. The fossilized birth-death model for the analysis of stratigraphic range data under different speciation modes. Journalof Theoretical Biology. 447 :41–55. 10.1016/j.jtbi.2018.03.005

Stadler T, Kühnert D, Bonhoeffer S, Drummond AJ. 2013. Birth-death skyline plot reveals temporal changes of epidemic spread in HIV and hepatitis C virus (HCV). Proceedings of the NationalAcademy of Sciences of the United States of America. 110 (1):228–233. 10.1073/pnas.1207965110

Stadler T, Steel M. 2019. Swapping birth and death: Symmetries and transformations in phylodynamic models. Systematic Biology. 68 (5):852–858. 10.1093/sysbio/syz039

Stadler T, Yang Z. 2013. Dating phylogenies with sequentially sampled tips. Systematic Biology.62 (5):674–688. 10.1093/sysbio/syt030

Stamatakis A, Alachiotis N. 2010. Time and memory efficient likelihood-based tree searches on phylogenomic alignments with missing data. Bioinformatics. 26 (12):i132–i139. 10.1093/bioinformatics/btq205

Steeman ME, Hebsgaard MB, Fordyce RE, Ho SYW, Rabosky DL, Nielsen R, Rahbek C, Glenner H, Sørensen MV, Willerslev E. 2009. Radiation of extant cetaceans driven by restructuring of the oceans. Systematic Biology. 58 (6):573–585. 10.1093/sysbio/syp060

Stolz U. 2024. Beyond the binary: Generalised trees for phylodynamics. Doctoral Thesis. Zürich: ETH Zürich. 10.3929/ethz-b-000706825

Stolz U, Gavryushkina A, Vaughan TG, Stadler T, Allen BJ. 2025. Enhancing evolutionary timelines:The impact of stratigraphic range information on phylogenetic inference. bioRxiv. 10.1101/2025.04.17.649084

Syed S, Bouchard-Côté A, Deligiannidis G, Doucet A. 2022. Non-reversible parallel tempering: A scalable highly parallel MCMC scheme. Journal of the Royal Statistical Society Series B: Statistical Methodology. 84 (2):321–350. 10.1111/rssb.12464

Talts S, Betancourt M, Simpson D, Vehtari A, Gelman A. 2020. Validating Bayesian inference algorithms with simulation-based calibration. arXiv. 10.48550/arXiv.1804.06788

Tao Q, Barba-Montoya J, Huuki LA, Durnan MK, Kumar S. 2020. Relative efficiencies of simple and complex substitution models in estimating divergence times in phylogenomics. Molecular Biology and Evolution. 37 (6):1819–1831. 10.1093/molbev/msaa049

Tavaré S. 1986. Some probabilistic and statistical problems in the analysis of DNA sequences. In: Miura RM, editor. Some Mathematical Questions in Biology: DNA Sequence Analysis. Vol. 17. American Mathematical Society. p. 57–86.

Tavaré S. 2018. The linear birth–death process: An inferential retrospective. Advances in Applied Probability. 50 :253–269. 10.1017/apr.2018.84

Thomson RC, Spinks PQ, Shaffer HB. 2021. A global phylogeny of turtles reveals a burst of climate-associated diversification on continental margins. Proceedings of the National Academy of Sciences of the United States of America. 118 (7):e2012215118. 10.1073/pnas.2012215118

Thorne JL, Kishino H. 2002. Divergence time and evolutionary rate estimation with multilocus data. Systematic Biology. 51 (5):689–702. 10.1080/10635150290102456

Thorne JL, Kishino H, Painter IS. 1998. Estimating the rate of evolution of the rate of molecular evolution. Molecular Biology and Evolution. 15 (12):1647–1657. 10.1093/oxfordjournals.molbev.a025892

Truman K, Vaughan T, Gavryushkin A, Gavryushkina AS. 2025. The fossilized birth-death model is identifiable. Systematic Biology. 74 (1):112–123. 10.1093/sysbio/syae058

Uhlenbeck GE, Ornstein LS. 1930. On the theory of Brownian motion. Physical Review. 36 (5):823–841.10.1103/PhysRev.36.823

Vaughan TG, Leventhal GE, Rasmussen DA, Drummond AJ, Welch D, Stadler T. 2019. Estimating epidemic incidence and prevalence from genomic data. Molecular Biology and Evolution. 36 (8):1804–1816. 10.1093/molbev/msz106

Vehtari A, Gelman A, Simpson D, Carpenter B, Bürkner PC. 2021. Rank-normalization, folding, and localization: An improved R for assessing convergence of MCMC. Bayesian Analysis. 16 (2):667– 718. 10.1214/20-BA1221

Wang S, Meade A. 2026. Molecular clock dating using complex mixture models: Applied to ancientsymbionts. Molecular Biology and Evolution. 43 (3):msag039. 10.1093/molbev/msag039

Warnock RCM, Heath TA, Stadler T. 2020. Assessing the impact of incomplete species sampling on estimates of speciation and extinction rates. Paleobiology. 46 (2):137–157. 10.1017/pab.2020.12

Warnock RCM, Parham JF, Joyce WG, Lyson TR, Donoghue PCJ. 2015. Calibration uncertainty in molecular dating analyses: There is no substitute for the prior evaluation of time priors. Proceedings of the Royal Society B: Biological Sciences. 282 (1798):20141013. 10.1098/rspb.2014.1013

Warnock RCM, Yang Z, Donoghue PCJ. 2012. Exploring uncertainty in the calibration of the molecular clock. Biology Letters. 8 (1):156–159. 10.1098/rsbl.2011.0710

Warnock RCM, Yang Z, Donoghue PCJ. 2017. Testing the molecular clock using mechanistic models offossil preservation and molecular evolution. Proceedings of the Royal Society B: Biological Sciences. 284 (1857):20170227. 10.1098/rspb.2017.0227

Waugh WAO. 1958. Conditioned Markov processes. Biometrika. 45 (1–2):241–249. 10.1093/biomet/45.1-2.241

Wilkinson RD, Steiper ME, Soligo C, Martin RD, Yang Z, Tavaré S. 2011. Dating primate divergences through an integrated analysis of palaeontological and molecular data. Systematic Biology. 61 (1):16–31. 10.1093/sysbio/syq054

Wright AM, Bapst DW, Barido-Sottani J, Warnock RCM. 2022. Integrating fossil observations into phylogenetics using the fossilized birth-death model. Annual Review of Ecology, Evolution, and Systematics. 53 :251–273. 10.1146/annurev-ecolsys-102220-030855

Yang Z. 1994. Maximum likelihood phylogenetic estimation from DNA sequences with variable rates over sites: Approximate methods. Journal of Molecular Evolution. 39 (3):306–314. 10.1007/BF00160154

Yang Z. 2000. Maximum likelihood estimation on large phylogenies and analysis of adaptive evolution in Human Influenza Virus A. Journal of Molecular Evolution. 51 (5):423–432. 10.1007/s002390010105

Yang Z. 2007. PAML 4: Phylogenetic Analysis by Maximum Likelihood. Molecular Biology and Evolution. 24 (8):1586–1591. 10.1093/molbev/msm088

Yang Z, Rannala B. 2006. Bayesian estimation of species divergence times under a molecular clock using multiple fossil calibrations with soft bounds. Molecular Biology and Evolution. 23 (1):212–226.10.1093/molbev/msj024

Zhang C, Ronquist F, Stadler T. 2023. Skyline fossilized birth-death model is robust to violations ofsampling assumptions in total-evidence dating. Systematic Biology. 72 (6):1316–1336. 10.1093/sysbio/syad054

Zhang C, Stadler T, Klopfstein S, Heath TA, Ronquist F. 2016. Total-evidence dating under the fossilized birth-death process. Systematic Biology. 65 (2):228–249. 10.1093/sysbio/syv080

Zhou X, Xu S, Yang Y, Zhou K, Yang G. 2011. Phylogenomic analyses and improved resolution of Cetartiodactyla. Molecular Phylogenetics and Evolution. 61 (2):255–264. 10.1016/j.ympev.2011.02.009

Zurano JP, Magalhães FM, Asato AE, Silva G, Bidau CJ, Mesquita DO, Costa GC. 2019. Cetartiodactyla: Updating a time-calibrated molecular phylogeny. Molecular Phylogenetics and Evolution. 133 :256–262. 10.1016/j.ympev.2018.12.015

