## Supplementary Figure for "Estimating Divergence Times and Diversification Rates with the Unresolved Fossilized Birth-Death Process"

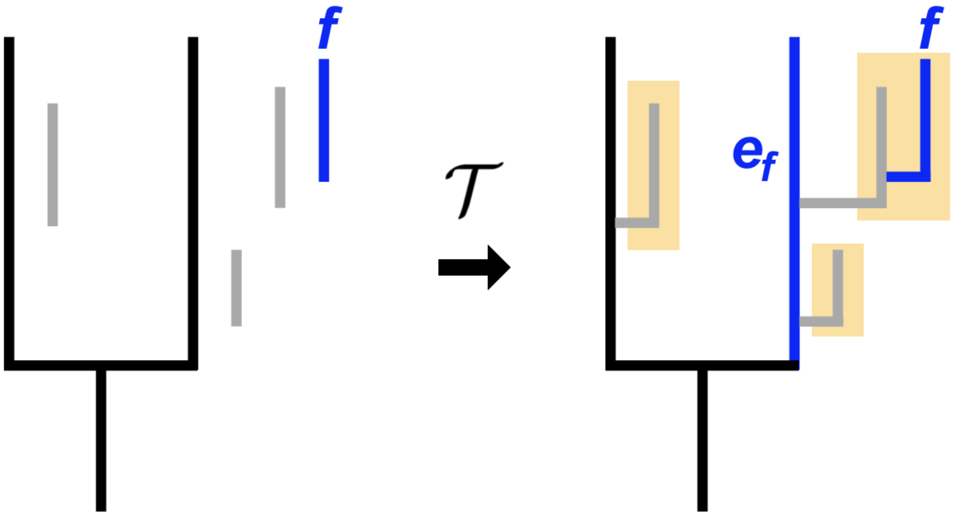


Supplementary Figure S1: A resolution, its $f$-clades, and $e_f$. Left, a backbone tree of two extant taxa and four unresolved fossils, with one of the fossils labeled $f$. Right, one resolved labeled tree $\mathcal{T} \in \mathcal{R}(\mathbf{z})$, in which each fossil attaches either to a backbone edge or to the stalk of another fossil. Shaded boxes mark the three $f$-clades. The fossil $f$ attaches to the stalk of another fossil, which in turn attaches to a backbone edge. Following this chain of stalks from $f$ reaches $e_f$.

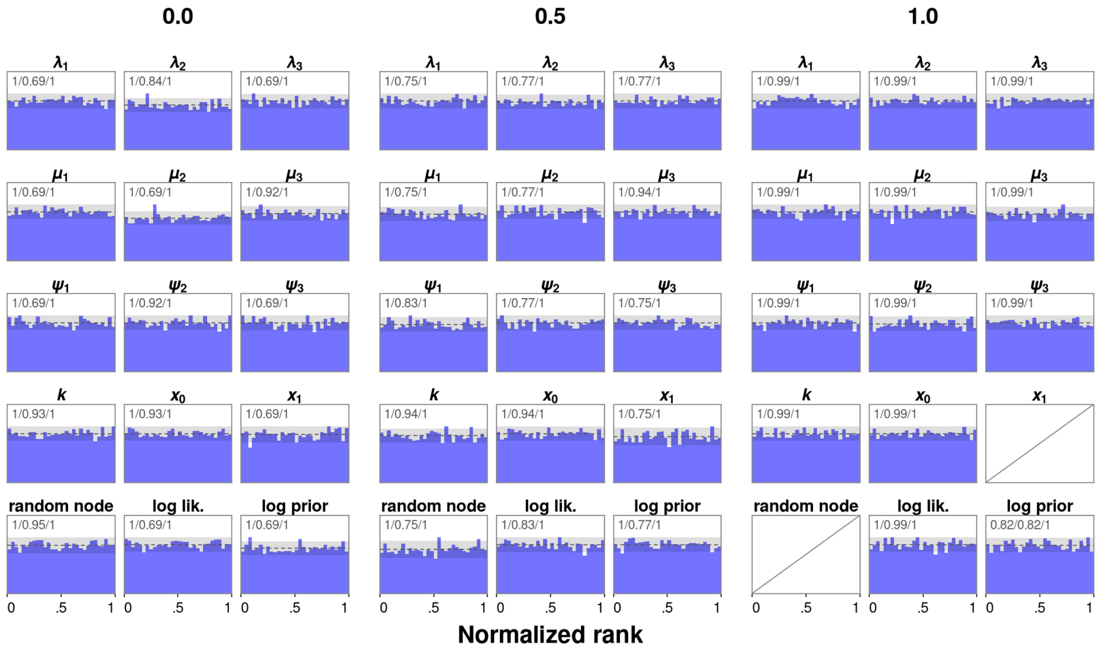


Supplementary Figure S2: Results of the simulation-based calibration checking under skyline rates drawn independently for each interval (rate model (ii)). Each panel shows a histogram of the rank of the true value of the test quantity named above it. Blocks correspond to the probability with which each extant taxon was designated a UE. $\lambda_i$, $\mu_i$, and $\psi_i$ are the rates of interval $i$, numbered from the oldest, and the other test quantities are as in Figure 2 of the main text; $x_1$ and the random node are undefined when the probability is 1. The dashed line, grey band, numbers in the top left of each panel, and frames are as in Figure 2 of the main text.

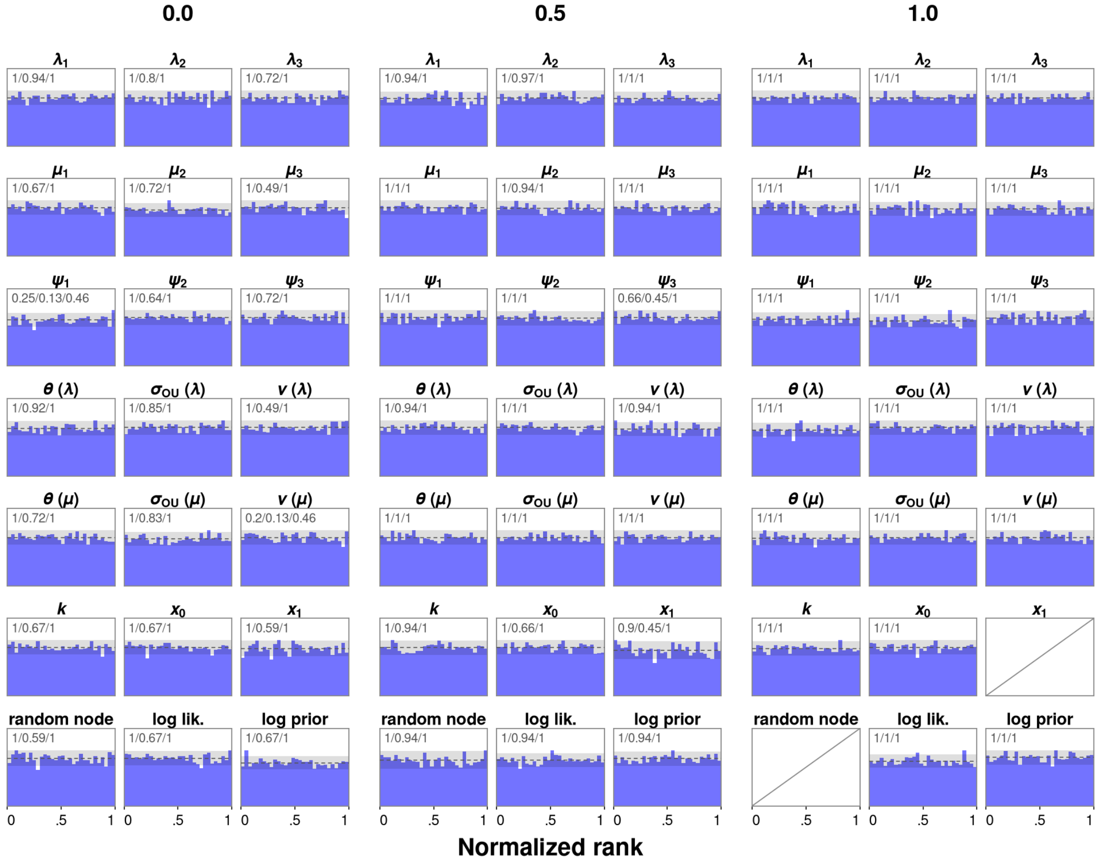


Supplementary Figure S3: Results of the simulation-based calibration checking under skyline rates with OU-smoothed $\lambda_i$ and $\mu_i$ (rate model (iii)), shown as in Supplementary Figure S2. $\theta$, $\sigma_{\mathrm{OU}}$, and $\nu$ are the OU parameters of the rate given in parentheses.

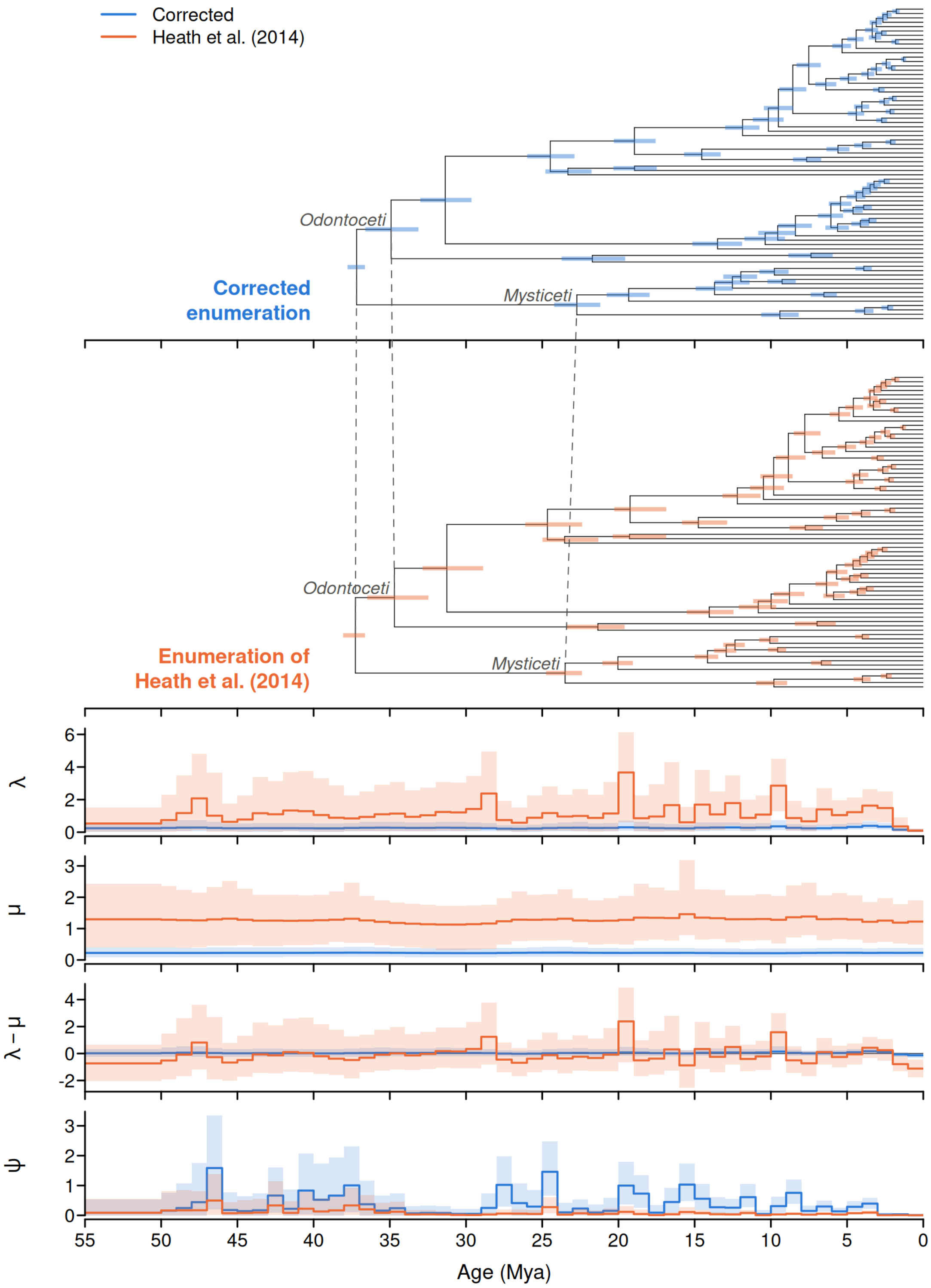


Supplementary Figure S4: Cetacea dataset analyzed with the corrected enumeration (top tree, blue) and with the enumeration of Heath et al. (2014) (bottom tree, orange). Trees are posterior mean trees, and node bars denote the 95% credible intervals of node ages, and dashed lines connect the crown nodes of Cetacea, Odontoceti, and Mysticeti between the two trees. The lower panels show the speciation ($\lambda$), extinction ($\mu$), diversification ($\lambda - \mu$), and sampling ($\psi$) rates through time, with lines denoting posterior medians and shades denoting 95% credible intervals.

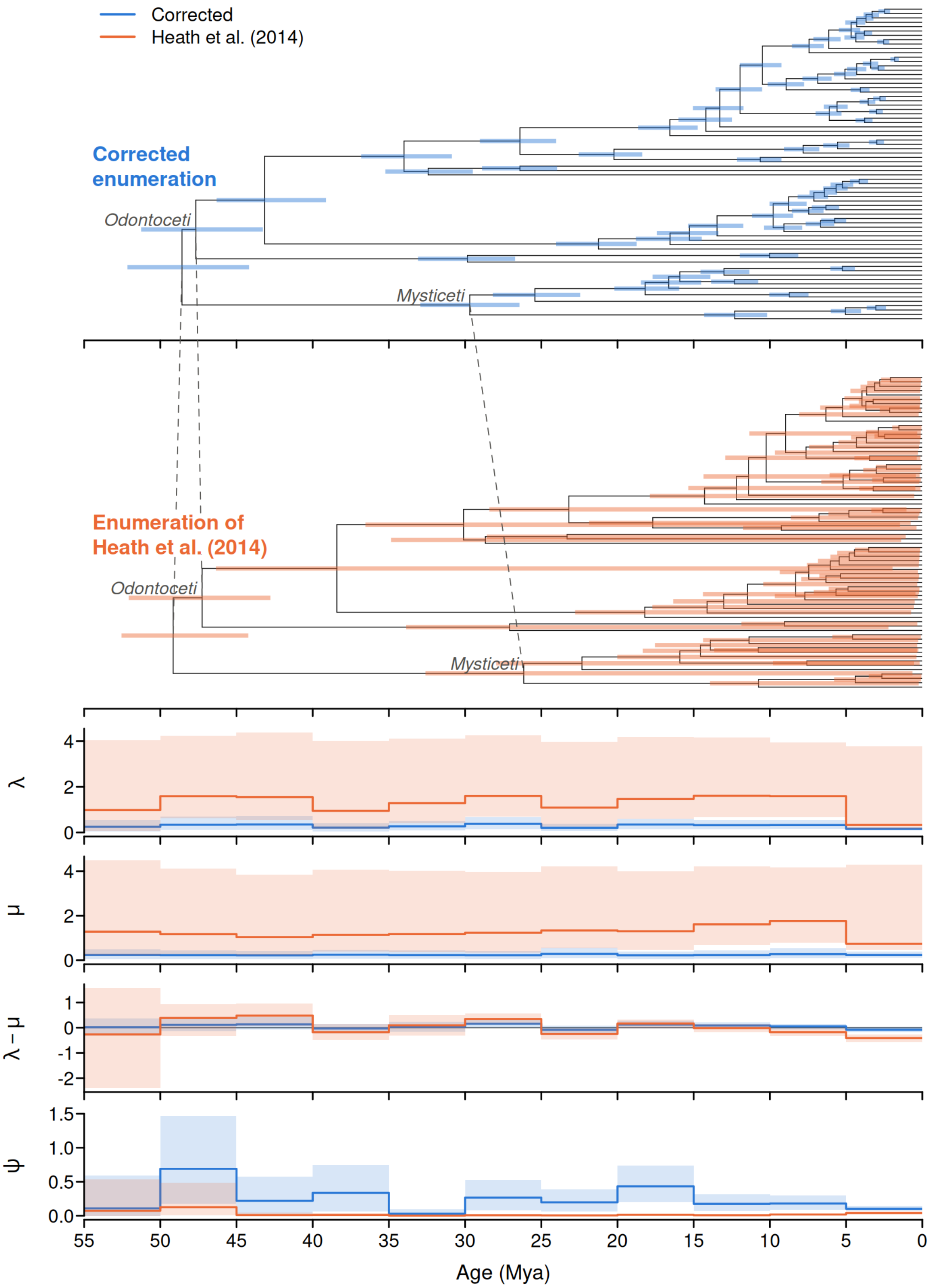


Supplementary Figure S5: Cetacea dataset with relaxed fossil assignments and 5-Myr rate intervals, analyzed with the corrected enumeration (top tree, blue) and with the enumeration of Heath et al. (2014) (bottom tree, orange), shown as in Supplementary Figure S4. Each fossil was assigned to the total group of Odontoceti or Mysticeti if its original clade was that suborder or nested within it, and to the total group of Cetacea otherwise.

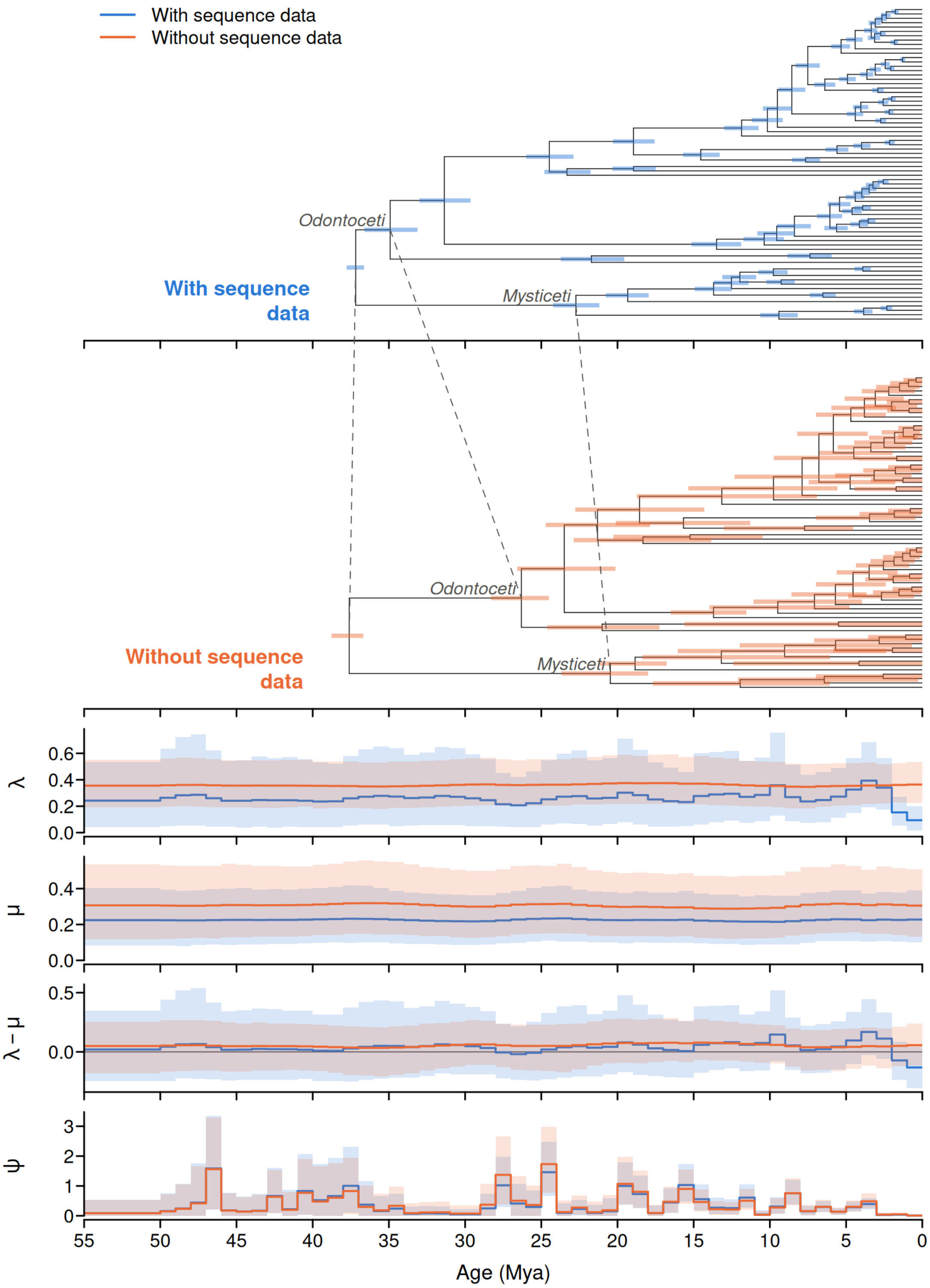


Supplementary Figure S6: Cetacea dataset analyzed with the sequence data (top tree, blue) and without the Hessian file and the clock model (bottom tree, orange), shown as in Supplementary Figure S4.

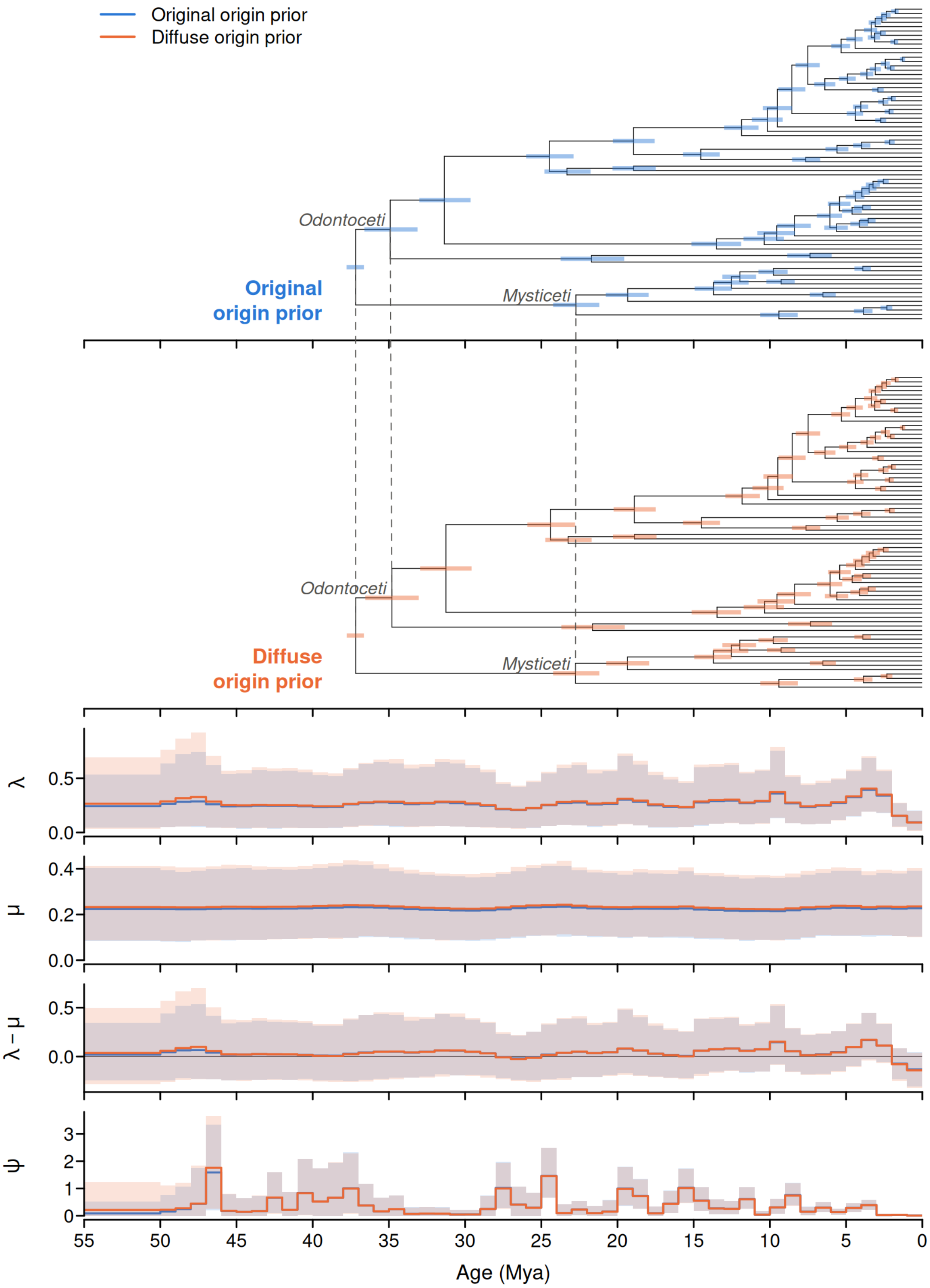


Supplementary Figure S7: Cetacea dataset analyzed with the origin prior $x_0 \sim \mathrm{TruncNormal}(55, 2^2)$ bounded below at 50 Mya (top tree, blue; "Original origin prior") and with $x_0 - 46 \sim \mathrm{Exp}(0.1)$ (bottom tree, orange; "Diffuse origin prior"), shown as in Supplementary Figure S4.

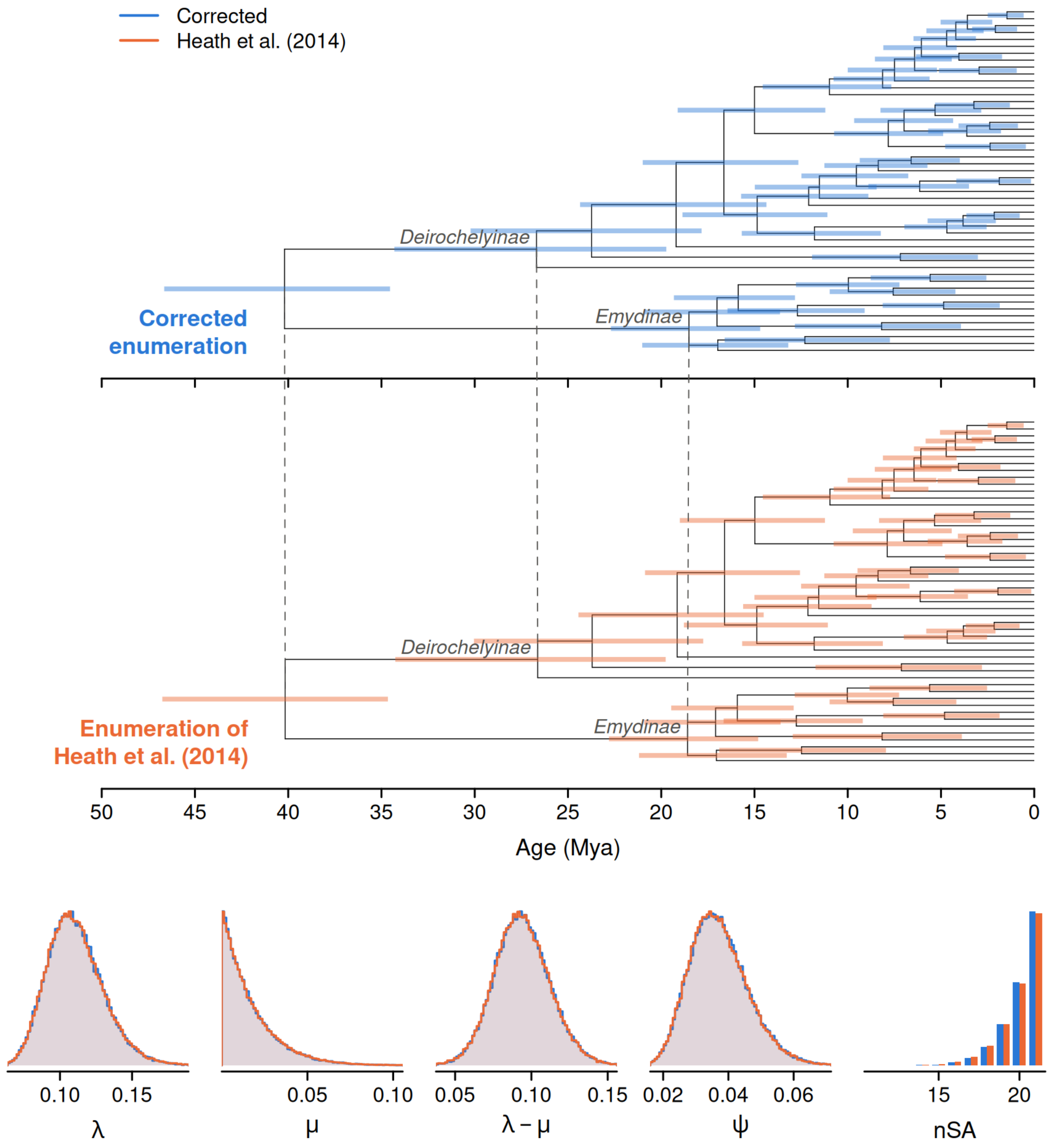


Supplementary Figure S8: Emydidae dataset analyzed with the corrected enumeration (top tree, blue) and with the enumeration of Heath et al. (2014) (bottom tree, orange), shown as in Supplementary Figure S4; dashed lines connect the crown nodes of Emydidae, Emydinae, and Deirochelyinae. The bottom panels show the posterior distributions of $\lambda$, $\mu$, $\lambda - \mu$, $\psi$, and the number of sampled ancestors (nSA). The total number of Emydidae fossils included in the analysis is 21.

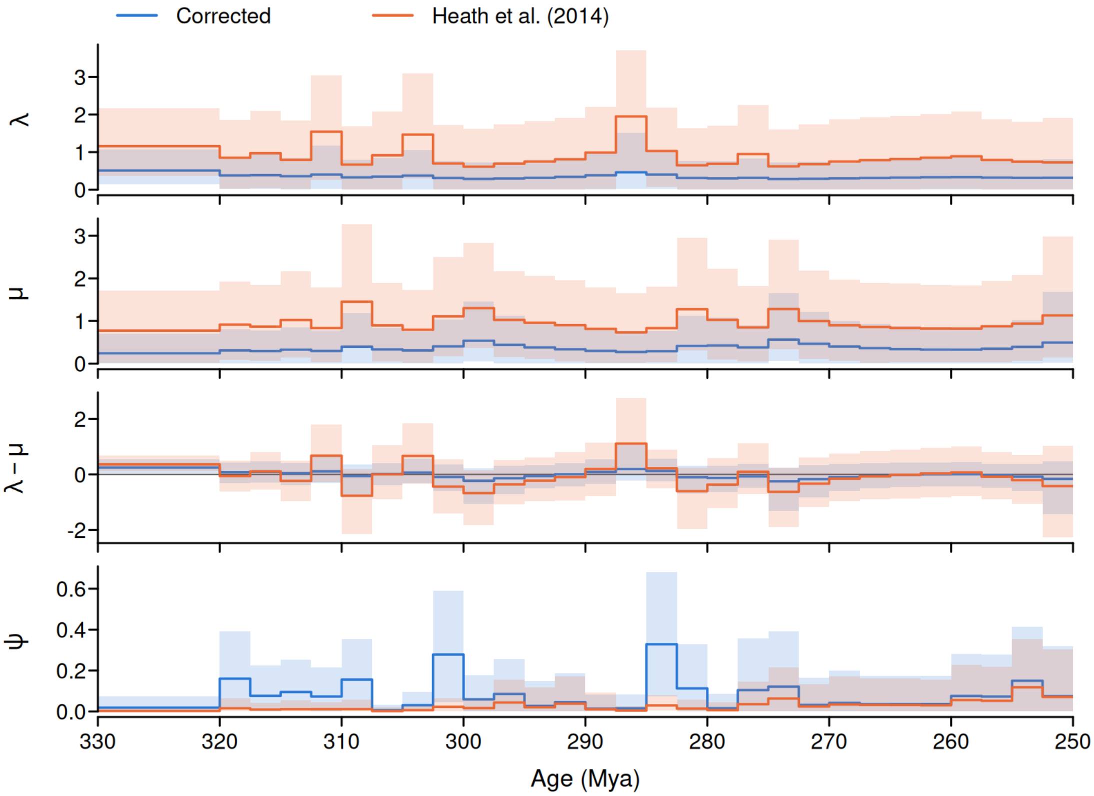


Supplementary Figure S9: Speciation ($\lambda$), extinction ($\mu$), diversification ($\lambda - \mu$), and sampling ($\psi$) rates of the Palaeodictyopterida dataset estimated with the corrected enumeration (blue) and with the enumeration of Heath et al. (2014) (orange), shown as in Supplementary Figure S4. The analysis under the enumeration of Heath et al. (2014) did not converge within $10^9$ iterations.
