## Supplementary Material 1 for "Estimating Divergence Times and Diversification Rates with the Unresolved Fossilized Birth-Death Process"

#### 1 Details of the MCMC algorithm

**Adaptive Metropolis-coupled Markov chain Monte Carlo** Metropolis-coupled MCMC (MC3) runs several chains in parallel, the $k$th targeting the posterior raised to a power $\beta_k$ (the cold chain at $\beta_0 = 1$), and periodically proposes swaps of their states so that the heated chains help the cold chain traverse the posterior more effectively (Geyer, 1991; Gilks & Roberts, 1996; Altekar et al., 2004). Adaptive variants tune the power coefficients during the run to control the swap acceptance (Miasojedow et al., 2013; Müller & Bouckaert, 2020). Swaps are proposed between chains at adjacent power coefficients under a deterministic even-odd (DEO) scheme (Okabe et al., 2001; Lingenheil et al., 2009), alternating the parity of the swapped pairs on successive sweeps, so the swap chain is non-reversible: the range of power coefficients is updated by a Robbins–Monro update (Robbins & Monro, 1951) toward a mean adjacent-swap acceptance of 0.40 (the optimum for the DEO kernel; Lingenheil et al. (2009)), and their interior values so that this acceptance is spread evenly across the adjacent chains (Syed et al., 2022).

**Proposal kernels and adaptive proposals** Unless noted otherwise, multiplicative moves use a Bactrian scale kernel (Yang & Rodríguez, 2013; Thawornwattana et al., 2018): a positive parameter $x$ is proposed as $x' = xe^{s\delta}$, with $\delta$ drawn from the symmetric two-component normal mixture $\frac{1}{2}\text{Normal}(-m, 1 - m^2) + \frac{1}{2}\text{Normal}(m, 1 - m^2)$, where $m = 0.95$. Each step size $s$ is adapted by a Robbins–Monro update on $\log s$ (Roberts & Rosenthal, 2007; Andrieu & Thoms, 2008), toward a target acceptance probability of 0.3. The constant-distance node-age move (see "Node age proposals") maintains a separate, independently adapted step size for each node, and the clock moves maintain step size for each sequence partition.

**FBD rate proposals** Each skyline rate $\lambda_i, \mu_i, \psi_i$ is updated by the Bactrian scale move. Two joint moves act on the whole vector of one rate in $\{\lambda, \mu, \psi\}$, independent across bins or OU-smoothed, chosen uniformly at random, proposed together with probability 0.20: a "vector scale" multiplies every bin by $e^{s\delta}$ (giving log-Jacobian $ls\delta$ for an $l$-bin vector; under OU smoothing it also adds $s\delta$ to $\theta$, so each bin's deviation from $\theta$ is unchanged), and a "shrink–expand" maps each bin's log-rate as $\log r_i \mapsto \overline{\log r} + e^{s\delta}(\log r_i - \overline{\log r})$ about the vector's mean log-rate $\overline{\log r}$ (log-Jacobian $(l-1)\,s\delta$). When $\lambda$ or $\mu$ is being smoothed by the OU process (Uhlenbeck & Ornstein, 1930) (details below), a joint "level shift" is proposed with probability 0.10: the additive amount $d = s\delta$ is applied to the level of both $\lambda$ and $\mu$ at once, added to each bin for an independent vector, or applied to the OU smoothing's mean rate while its increments are held fixed. Level shift was motivated by the observation that $\lambda_i - \mu_i$ is usually tightly constrained, and perturbing $\lambda$ or $\mu$ alone moves off this ridge and is rarely accepted, whereas the level shift keeps the difference fixed by shifting both rates together. Because the number of SAs covaries with rates, the level shift is paired with a batch of SA state flips (toward SAs when $d < 0$, toward tips when $d > 0$), carrying the reversible-jump Jacobian and the corresponding combinatorial factor (Green, 1995; Gavryushkina et al., 2014; Heath et al., 2014) (see below).

We allow for smoothing of $\lambda$, $\mu$, or $\psi$ via OU (Uhlenbeck & Ornstein, 1930) process on its log-rates. Writing $\xi_i = \log r_i$ for the log-rate of bin $i$,

$$\xi_0 \sim \text{Normal}(\theta, \sigma^2_{\text{OU}}), \qquad \xi_i \mid \xi_{i-1} \sim \text{Normal}\big(\theta + (\xi_{i-1} - \theta)e^{-\nu\Delta_i},\, \sigma^2_{\text{OU}}(1 - e^{-2\nu\Delta_i})\big),$$

where $\theta$ is the long-term mean log-rate, $\sigma^2_{\text{OU}}$ the stationary variance, $\nu$ the strength of mean reversion, and $\Delta_i$ the time between the midpoints of bins $i-1$ and $i$; each term carries a Jacobian $1/r_i$ from the change of variables to the rate. By default, $\theta \sim \text{Normal}(\log 0.2, 1.6^2)$, $\sigma_{\text{OU}} \sim \text{Exp}(10/3)$, and $\nu \sim \text{Exp}(2)$, with $\nu$ per Myr. Under the default prior on $\nu$, the log-rates of two bins whose midpoints are 1, 5, and 10 Myr apart have prior median correlations of 0.71, 0.18, and 0.03. This parametrizes the process by the stationary variance $\sigma^2_{\text{OU}}$. Parameters are updated by per-bin and hyperparameter moves: with probability $5/(l+5)$ for $l$ bins, one of five hyperparameter moves is chosen uniformly, and otherwise a single bin chosen uniformly is updated by the Bactrian scale move. $\theta$ is updated by adding a Bactrian step. $\sigma_{\text{OU}}$ and $\nu$ each have two moves: first updates $\sigma_{\text{OU}}$ or $\nu$ via Bactrian scale move and leaves the log-rates unchanged. When the data is uninformative about the log-rates this can hamper mixing as they follow their prior closely, and changing $\sigma_{\text{OU}}$ or $\nu$ alone makes them improbable under the new prior and thus making the mixing difficult. The second move shifts every log-rate together with $\sigma_{\text{OU}}$ or $\nu$, so that each log-rate stays the same number of prior standard deviations from its prior mean using the non-centered parameterization (Papaspiliopoulos et al., 2003, 2007): for $\sigma_{\text{OU}}$ it first proposes $\sigma'_{\mathrm{OU}} = e^{s\delta}\sigma_{\mathrm{OU}}$ and updates every log-rate as $\xi'_i = \theta + e^{s\delta}(\xi_i - \theta)$ for $i = 0, \ldots, l-1$, leaving $\theta$ and $\nu$ unchanged, with log-Jacobian $(l+1)s\delta + \sum_{i=0}^{l-1}(\xi'_i - \xi_i)$. For $\nu$ it first proposes $\nu' = e^{s\delta}\nu$, leaves $\theta$, $\sigma_{\mathrm{OU}}$, and $\xi_0$ unchanged, and updates $\xi_1, \ldots, \xi_{l-1}$ in turn as

$$\xi'_i = \theta + (\xi'_{i-1} - \theta)e^{-\nu'\Delta_i} + \left(\xi_i - \theta - (\xi_{i-1} - \theta)e^{-\nu\Delta_i}\right)\sqrt{\frac{1 - e^{-2\nu'\Delta_i}}{1 - e^{-2\nu\Delta_i}}},$$

with log-Jacobian $s\delta + \sum_{i=1}^{l-1}\left[\frac{1}{2}\log\{(1 - e^{-2\nu'\Delta_i})/(1 - e^{-2\nu\Delta_i})\} + \xi'_i - \xi_i\right]$.

$p_i(t)$ is evaluated from its solution, which is divided by $2\lambda_i$. When $\lambda_i \to 0$, this can result in numerical instability, so we replace it with asymptotic limits when $\lambda_i < \epsilon$ for some small $\epsilon$: taking the limit gives

$$p_i(t) \;=\; \frac{\mu_i}{\mu_i + \psi_i} + \left(p_{i+1}(t_i) - \frac{\mu_i}{\mu_i + \psi_i}\right) e^{-(\mu_i + \psi_i)(t - t_i)}.$$

We set $\epsilon$ based on the relative magnitude $\lambda_i/A_i$, since the numerator divided by $2\lambda_i$ is of order $\lambda_i$ but is formed by canceling terms of order $A_i$: we set $\epsilon$ to $10^{-7}A_i$ for $p_i$ and $1 - p_i$, which usually become unstable sooner, and to $10^{-9}A_i$ for $1 - p_{I(x_0)}(x_0)$ when conditioning on $A$.

**Fossil and attachment age proposals** Each unresolved fossil $f$ carries a stratigraphic age $y_f$ and a latent attachment age $z_f$ (for UE, $z_f$ only) (Heath et al., 2014). At each step, a fossil is chosen uniformly, and one of three moves is applied with equal probability: (1) the fossil age is redrawn uniformly within its stratigraphic bracket, accommodating stratigraphic uncertainty; (2) the attachment age is redrawn uniformly over the fossil's compatible interval; (3) the sampled-ancestor move (SA-RJMCMC) is applied, which is a reversible jump move (Green, 1995) that flips the fossil state between a sampled ancestor and a terminal tip, following Gavryushkina et al. (2014) and Heath et al. (2014), to which we refer for the Jacobian and acceptance ratio. Blocks of attachment ages are additionally rescaled jointly with the tree by the age-scaling moves below.

**Attachment zone proposals** When the clade assignments are nested, each fossil carries the smallest attachment zone it has reached, $Z(e_f)$, updated by three moves. The first draws $Z(e_f)$ from its full conditional (i.e., Gibbs sampling): the density $f(T_{or} \mid t_{or} = x_0)$ does not involve the attachment zones, so the full conditional of $Z(e_f)$ is proportional to $\prod_{g \in \mathrm{tip}} 2\tilde{\gamma}_g(\mathbf{Z}) \prod_{g \in \mathrm{SA}} \gamma_g(\mathbf{Z})$ evaluated at each attachment zone available to $f$ in $A_f$, the attachment zone fixed by $f$'s clade assignment. Only $f$ and the fossils whose attachment age is crossed by $f$'s stalk need to be recounted, and the candidate drawn from it is accepted with probability one. Since that conditional is zero at an attachment zone none of whose edges or stalks cross the current $z_f$, a second move proposes $Z(e_f)$ and $z_f$ jointly: a candidate zone is drawn uniformly among the other attachment zones available to $f$, and $z_f$ uniformly from $I(Z')$, where $Z'$ is the candidate zone. For an attachment zone $Z$, let $I(Z)$ be the union of the age intervals of the backbone edges whose own smallest attachment zone is $Z$ and of the stalks of the fossils that are not sampled ancestors and whose attachment zone is $Z$, each intersected with $f$'s permitted attachment age interval, and $M(Z)$ is the total length of $I(Z)$; then the Hastings ratio is $M(Z')/M(Z(e_f))$. The third resamples attachment zones within the slide of a single internal age, the crown node age scale, the constant-distance move, and the sequential scan of internal node ages (see "Node age proposals"). A boundary of an attachment zone sits at a backbone node age, so a node-age move that holds $Z(e_f)$ fixed can leave no edge of $Z(e_f)$ crossing $z_f$, which gives $\gamma_f(\mathbf{Z}) = 0$, making the proposal inefficient. When a node age moves from $x$ to $x'$, let $F$ be the set of fossils whose clade assignment spans more than one attachment zone, whose attachment age lies between $x$ and $x'$, and that can attach to an edge connected to the moved node. For each $f \in F$, a new attachment zone $Z'(e_f)$ is sampled from the distribution

$$\varphi_f(Z \mid \mathbf{x}') = \frac{\tilde{\gamma}_f(Z, \mathbf{x}')}{\sum_{W \subseteq A_f} \tilde{\gamma}_f(W, \mathbf{x}')},$$

with $\gamma_f(Z, \mathbf{x}')$ in place of $\tilde{\gamma}_f(Z, \mathbf{x}')$ when $f$ is an SA, holding every fossil other than $f$ at its current attachment zone, with the log Hastings ratio of the node-age move

$$\sum_{f \in F} \log \frac{\varphi_f(Z(e_f) \mid \mathbf{x})}{\varphi_f(Z'(e_f) \mid \mathbf{x}')}.$$

The numerator is the probability of drawing $f$'s current attachment zone back from the proposed state, so the other fossils in $F$ are held at their proposed attachment zones.

**Sequence likelihood computation** Because the backbone topology is fixed, the likelihood computation via the Felsenstein pruning algorithm (Felsenstein, 1981) can be evaluated incrementally. A partial-likelihood vector is stored at every backbone node, and a move that changes only some branch lengths recomputes the partial likelihoods only for the nodes on the path from the change to the root, together with the transition matrices of the branches below them, and reuses the cached vectors elsewhere. Following Kobert et al. (2017), on the fixed backbone, the site-repeat structure is computed once at initialization: each node stores its partial likelihoods over its distinct repeat classes, a parent indexes each child through a precomputed per-class map, and the per-site sum at the root reads each pattern's class. Tips absent from a partition are further skipped from computation (Stamatakis & Ott, 2008; Stamatakis & Alachiotis, 2010).

For the approximate-likelihood path, full sequence likelihood evaluation is replaced with a second-order approximation of the log-likelihood about the pre-inferred maximum-likelihood branch lengths

(Thorne et al., 1998; Thorne & Kishino, 2002; Seo et al., 2004; Guindon, 2010; dos Reis & Yang, 2011), expressed in the arcsine-transformed branch lengths using the gradient and Hessian for the fixed topology (dos Reis & Yang, 2011). Each likelihood evaluation is a quadratic function of the difference between the current transformed branch lengths and their estimates. Because most moves change only a few branch lengths, we cache this difference and its product with the Hessian, and at each evaluation, update that product only through the Hessian columns of the branches whose lengths changed, using compensated summation (Kahan, 1965; Neumaier, 1974; Ogita et al., 2005) to bound the accumulated round-off.

**Substitution model proposals** We set a prior for exchangeabilities and the estimated base frequencies as uniform over the simplex, the gamma shape as an exponential with mean one, and, when invariable sites are included, the proportion of invariable sites as uniform on $(0, 1)$. The exchangeabilities and frequencies are updated by a "delta-exchange": two components of the simplex are chosen uniformly, and a Bactrian-distributed amount is transferred from one to the other, leaving their sum unchanged and the proposal symmetric, and a proposal that takes either component outside $(0, 1)$ is rejected. The gamma shape and the proportion of invariable sites are updated by the Bactrian scale move.

**Clock model proposals** Across clock partitions, the vectors $(r_{\text{clock},j})_j$ and $(\sigma^2_{\text{clock},j})_j$ each follow a gamma-Dirichlet prior described in dos Reis et al. (2014) (see also Jin and Brown (2018)), under which the mean over the $L$ clock partitions ($\bar{r}_{\text{clock}}$ or $\overline{\sigma^2}_{\text{clock}}$) follows a gamma prior and the proportions of the $L$ values follow a Dirichlet prior; it reduces to a gamma prior when there is a single clock partition. Given clock partition $j$'s mean $r_{\text{clock},j}$ and variance $\sigma^2_{\text{clock},j}$, its per-branch rates are independent under the UCLN clock (Drummond et al., 2006), with $\log r_{b,j} \sim \text{Normal}(\log r_{\text{clock},j} - \sigma^2_{\text{clock},j}/2,\ \sigma^2_{\text{clock},j})$; under the GBM clock (Kishino et al., 2001) the log-rate instead diffuses as autocorrelated Brownian motion along the tree, so each node's log-rate is normal about its ancestor's with variance $\sigma^2_{\text{clock},j}t$ and mean offset $-\sigma^2_{\text{clock},j}t/2$ (keeping the expected rate equal to the ancestor's), where $t$ is the time between the parent and daughter branch midpoints, and the two daughter lineages of a node are correlated through their shared ancestor. Each per-branch rate $r_{b,j}$ and the mean $r_{\text{clock},j}$ are updated by the Bactrian scale move, and a global move that multiplies $r_{\text{clock},j}$ and every branch rate of clock partition $j$ by a common factor (log-Jacobian $(1 + B)s\delta$ for $B$ branches). We additionally allow for more flexible partitioning of the clock model as well (Duchêne & Ho, 2014, 2015; Foster & Ho, 2017).

The variance $\sigma^2_{\text{clock},j}$ is updated in one of two parameterizations (Papaspiliopoulos et al., 2003, 2007). In the centered parameterization, used by default, it is scaled directly by the Bactrian move with the branch rates held fixed. In the non-centered parameterization, the standardized residuals $u_{b,j} = (\log r_{b,j} - \eta_{b,j})/s_{b,j}$ are held fixed while $\sigma^2_{\text{clock},j}$ is scaled by the Bactrian move and every branch rate is recomputed as $\log r'_{b,j} = \eta'_{b,j} + s'_{b,j} u_{b,j}$, where $\eta_{b,j}$ and $s_{b,j}$ are the prior mean and standard deviation of $\log r_{b,j}$, both functions of $\sigma^2_{\text{clock},j}$; the move carries the log-Jacobian $\sum_b \log(r'_{b,j}/r_{b,j})$ in addition to that of the scale proposal.

We tested the consequences of choosing between centered and non-centered parameterizations for the clock-rate variance on the Emydidae dataset, as its sequence partitions were relatively short (median 855 bp; range 593–1,528 bp) compared with those of the Cetacea dataset, for which the Hessian file was derived from an alignment containing hundreds of kilobases per partition, where centered parameterization would be favored. After more than two weeks of running, the Emydidae dataset under centered parameterization did not converge within the maximum cutoff of 200 million MCMC iterations, with $\hat{R}$ of 1.021, minimum bulk ESS of 187, and minimum tail ESS of 286, with the slowest-converging parameters all being clock-rate, in contrast to non-centered parameterization that converged at around 23 million iterations.

**Node age proposals** Backbone node ages are updated by a uniform slide of a single internal age within the interval bounded by its oldest child and its parent (Hastings ratio equal to 1), and the crown node age by a multiplicative scale (Hastings ratio equal to the scale factor); the origin, when present, is scaled by a separate move. To move node ages without disturbing the sequence likelihood, we use the rate-compensated operators of Zhang and Drummond (2020) (see also Thorne et al. (1998)), summarized briefly here (full Jacobians and acceptance ratios are given in Zhang and Drummond (2020)): the internal-node "constant-distance" move perturbs one node age and rescales its three incident branch rates so that all incident genetic distances are preserved, and its crown node analogues "simple distance" (both clocks), and "small pulley" (used only for the UCLN clock, as it is most efficient only when the rates of branches directly above the crown node are not constrained by rate autocorrelation) act on the crown node age and the rates of branches directly above the crown node. A clade-block variant rescales the ages of an entire randomly chosen subtree together with its branch rates and interior attachment ages, as implemented in RevBayes (Höhna et al., 2016).

To further improve the mixing of deep node ages, all internal node ages are updated one at a time in a sequential scan, each by a uniform proposal within its (oldest-child, parent) interval, with branch rates held fixed. Finally, an up-down move jointly scales all node ages (including the fossil attachment ages and origin) in one direction and the rates in the other (Thorne et al., 1998). The constant-distance move and the sequential scan are two ways to update the internal node ages. We adaptively choose between them, weighting each by its mean squared jump in the log node-ages per node updated (Pasarica & Gelman, 2010), with a 0.05 floor on each. The other node-age moves target different quantities and are proposed at fixed probabilities.

**Dynamic spine election** When no backbone tree is supplied, one fossil must anchor the origin. Because the youngest taxon can never be a sampled ancestor, we designate the fossil with the absolute youngest age as a "spine" whose attachment age is fixed to the origin, $z_{\text{spine}} = x_0$, and which is excluded from the attachment and sampled-ancestor moves. When an absolute youngest fossil exists, it is chosen once at initialization (ties broken uniformly at random). However, when fossil ages are sampled from stratigraphic bounds and no fossil is unambiguously youngest, the identity of the youngest fossil fluctuates, so the spine is re-elected within the fossil-age move: after proposing a new age for fossil $f$, the current youngest fossil is recomputed; if it differs from the original spine, the new spine's attachment age is set to $x_0$ and the former spine is assigned a finite attachment age inherited from the new spine's previous configuration, leaving the proposal reversible (Hastings ratio 1).

### 2 Gamma-counting algorithm

Evaluating $f(T_{ur} \mid t_{or} = x_0)$ requires, at every likelihood evaluation, the count $\tilde{\gamma}_f(\mathbf{Z})$ for each tip fossil and UE and $\gamma_f(\mathbf{Z})$ for each SA. A naive implementation obtains each by a linear scan over the $\tilde{n}$ candidate attachment edges or stalks, so a full evaluation costs $\mathcal{O}((m+k)\tilde{n})$. We instead maintain indices that evaluate a count in $\mathcal{O}((m+k)\log\tilde{n})$ time and that limit recomputation after a local move to the fossils whose count can change.

**Indices** Three objects are maintained over the current $(\mathbf{x}, \mathbf{y}, \mathbf{z})$, and $\mathbf{Z}$.

(i) A depth-first preorder index of $T_{bb}$. An iterative traversal assigns each backbone node $v$ a preorder rank $\text{pre}(v)$, its position in the visiting order, and a subtree size $\text{size}(v)$, the number of nodes in the subtree rooted at $v$, including $v$. Because the traversal completes an entire subtree before leaving it, the nodes of that subtree occupy the contiguous block $[\text{pre}(v), \text{pre}(v) + \text{size}(v))$, so the test "$u$ lies in the subtree rooted at $v$" is the constant-time comparison $\text{pre}(v) \le \text{pre}(u) < \text{pre}(v) + \text{size}(v)$. This decides whether an edge lies in an attachment zone, and is used once, at setup, to enumerate the distinct attachment zones, to assign each backbone edge its $Z(e)$, and to record for each fossil the attachment zones contained in $A_f$, which are the values $Z(e_f)$ may take.

(ii) For each attachment zone, two sorted arrays holding the tipward and the rootward endpoint ages of every backbone edge $e$ whose $Z(e)$ is that zone. Reading the arrays of $Z(e_f)$, $b_f$ is the difference of two binary searches, $\#\{\text{tipward} < z_f\} - \#\{\text{rootward} \le z_f\}$. The edge running from the crown to the origin $x_0$ is carried in the arrays of its own $Z(e)$.

(iii) For each attachment zone, the analogous sorted arrays of the stalk endpoints $(y_g, z_g)$ of the tip fossils and UEs $g$ whose $Z(e_g)$ is that zone. Reading the arrays of $Z(e_f)$, $s_f$ is $\#\{y_g < z_f\} - \#\{z_g \leq z_f\}$.

**Assembling a count** A count reads only the arrays of the fossil's own $Z(e_f)$: $b_f$ from (ii), $s_f$ from (iii), then $\tilde{\gamma}_f(\mathbf{Z}) = b_f + \frac{1}{2}s_f$ for a tip fossil or UE and $\gamma_f(\mathbf{Z}) = b_f + s_f$ for an SA. In the no-backbone case one taxon is elected to anchor the origin, and its stalk is thereafter a backbone edge, so it is held in the arrays of (ii) and enters $b_f$ at full weight.

**Incremental maintenance** After a move that perturbs a backbone node age, an attachment age or the origin age, flips a fossil's SA state, or changes a $Z(e_f)$, the indices and the cached log-counts are incrementally updated rather than rebuilt from scratch. The arrays of (ii) are updated by erasing and reinserting only the endpoints of the perturbed edges in the arrays of their own $Z(e)$, unless the move perturbs more than $\lfloor \log_2 \tilde{n} \rfloor$ edges, in which case they are rebuilt and re-sorted; a local move therefore touches only its own edges and a global move triggers a single rebuild. The arrays of (iii) are updated from the current $(\mathbf{y}, \mathbf{z})$, and $\mathbf{Z}$ at a cost of $\mathcal{O}(m \log m)$. A fossil's cached count is flagged stale when its own $y_f$, $z_f$, SA state or $Z(e_f)$ changes, and also when a perturbed edge begins or ceases to cross its attachment age: an attachment-age move flags the fossils whose attachment age falls in the interval swept by the perturbed stalk, an origin move flags the whole-tree total group fossils whose attachment age lies between the old and the new origin age, and a node-age move flags the fossils whose attachment age falls in the interval of a perturbed edge. Only the flagged fossils are recounted. Since each count depends only on the current $(\mathbf{x}, \mathbf{y}, \mathbf{z})$ and $\mathbf{Z}$, this equals the direct evaluation.

### 3 Additional SBC

Each setting below was analyzed and tested as in the main text.

**Fossil ages** To check the sampling of fossil ages within their age ranges, each simulated fossil $f$ with stratigraphic age $y_f$ was given an age range of fixed width $w$ placed uniformly at random around $y_f$. With $u_f \sim \text{Unif}(0, 1)$ drawn independently, the upper bound was set to $h_f = y_f + u_f w$ giving the age range $R_f = [\max(0, h_f - w), h_f]$. Given $y_f$, $h_f$ is uniform on $[y_f, y_f + w]$, so the density of reporting $R_f$ is

$$P(R_f \mid y_f) = w^{-1} \mathbf{1}\{y_f \in R_f\},$$

which does not depend on $\lambda$, $\mu$, $\psi$, $\mathbf{x}$, and $\mathbf{z}$ and takes the same value at every age within $R_f$, including a range being truncated at the present. The probability of the reported age ranges is therefore

$$\int f(T_{ur} \mid t_{or} = x_0) \prod_f P(R_f \mid y_f)\, d\mathbf{y} = w^{-(m+k)} \int_{R_1 \times \cdots \times R_{m+k}} f(T_{ur} \mid t_{or} = x_0)\, d\mathbf{y},$$

which equals, up to a constant, the likelihood the sampler uses when it samples each $y_f$ uniformly within its age range. Thus, a correct MCMC implementation will give uniform randomized normalized ranks of the test quantities.

We simulated the constant-rate FBD model (i) of the main text with UE probability 0.5 and gave each fossil an age range of width $w = 5$. In addition to the test quantities of the main text, we used the age of one fossil chosen uniformly at random in each replicate. As with the random backbone node in the main text, the fossil is chosen independently of the true values, so its age is a valid test quantity. A fossil fixed by its label would not be, because the simulated fossils are labeled in simulation order, which the model treats as uninformative. None of the test quantities, including the fossil age, rejected uniformity under any of the three adjustments, and the smallest BH-adjusted $p$-value was 0.39 (Figure A1-1).

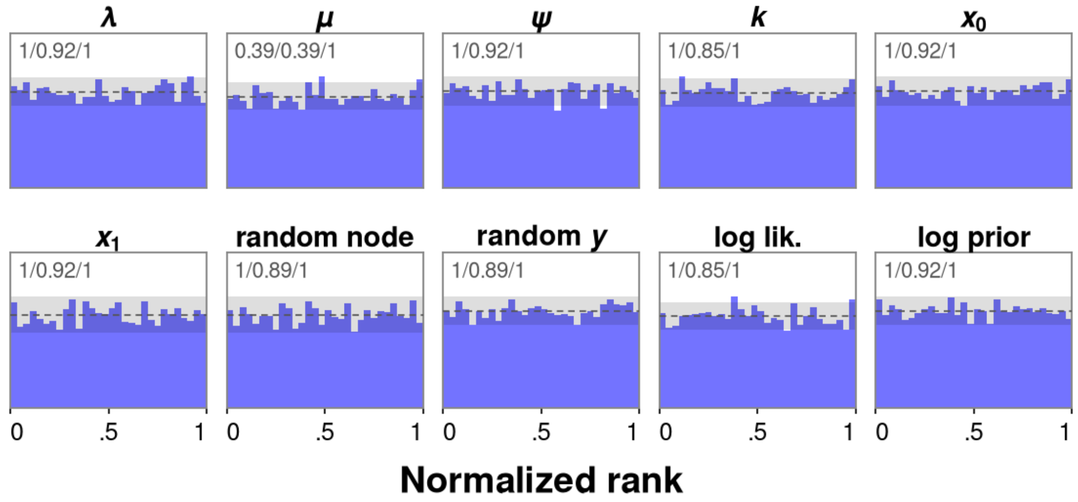


Figure A1-1: Results of the simulation-based calibration checking with sampled fossil ages. The test quantities, the dashed line, grey band, numbers in the top left of each panel, and frames are as in Figure 2 of the main text.

**Attachment zones** To check the sampling of $Z(e_f)$ under nested clade assignments, each simulated fossil $f$ was assigned to a clade by a walk along the backbone toward $e_f$ from the root, the backbone edge that $f$ reaches by following its chain of stalks. Let $u$ be the backbone node immediately tipward of $e_f$, and let $x_1 = v_0, v_1, \ldots, v_K = u$ be the backbone nodes on the path from the crown node of the backbone tree to $u$. Starting at $v_0$, the walk stops at $v_j$ with probability $r$ and assigns $f$ to the clade with crown node $v_j$, so that $A_f$ is the set of edges descending from $v_j$ together with the edge rootward of $v_j$; otherwise it moves on to $v_{j+1}$. If it reaches $u$ without stopping, $A_f = \{e_f\}$. When $u$ is a tip, the attachment zone of its clade is $\{e_f\}$, and the walk stops there. Every edge in the attachment zone of the clade with crown node $v_j$ is reached through the same nodes $v_0, \ldots, v_j$, so the probability of reporting $A_f$ is

$$P(A_f \mid e_f) = \begin{cases} r^{1-\mathrm{tip}(v_j)}(1-r)^j \, \mathbf{1}\{e_f \in A_f\}, & A_f \text{ is the zone of the clade with crown node } v_j, \\ (1-r)^{K+1} \, \mathbf{1}\{e_f \in A_f\}, & A_f = \{e_f\}, \end{cases}$$

where $\mathrm{tip}(v_j) = 1$ when $v_j$ is a tip and 0 otherwise. This depends on $A_f$ alone and not on which edge of $A_f$ carries $f$, nor on $\lambda$, $\mu$, $\psi$, $\mathbf{x}$, $\mathbf{y}$, and $\mathbf{z}$, so the clade assignments are coarsened at random (Heitjan & Rubin, 1991). Since $e_f$ is defined only within a resolved labeled tree $\mathcal{T}$, we write the probability of the reported clade assignments as an integral over these trees. Because $P(A_f \mid e_f)$ is zero unless $e_f \in A_f$ and otherwise takes a value set by $A_f$ alone, the product $\prod_f P(A_f \mid e_f)$ is the same for every tree compatible with the clade assignments and factors out. Let $\mathfrak{T}$ be the set of resolved labeled trees $\mathcal{T}$ compatible with the clade assignments, that is, with $e_f \in A_f$ for every $f$. Then

$$\int f(T_{lb} \mid t_{or} = x_0) \prod_f P(A_f \mid e_f) \, d\mathcal{T} \;\propto\; \int_{\mathfrak{T}} f(T_{lb} \mid t_{or} = x_0) \, d\mathcal{T}.$$

The right-hand side is, up to a constant proportionality, the marginal distribution of $f(T_{ur} \mid t_{or} = x_0)$ over $\mathbf{z}$. Since $r$ is fixed and thus distinct from the model parameters, by Theorem 1 (ii) of Heitjan and Rubin (1991), the posterior based on this likelihood is the correct posterior. Thus, a correct MCMC implementation gives uniform randomized normalized ranks of the test quantities. Starting the walk at $e_f$ and moving rootward instead would make the probability of reporting a clade depend on the number of nodes between $e_f$ and the crown node of that clade, so the reported clade would carry information on the position of $e_f$ within $A_f$ that the likelihood does not use.

We simulated the constant-rate FBD model (i) of the main text with UE probability 0 and assigned each fossil to a clade with $r = 0.3$. In addition to the test quantities of the main text, we used the attachment age of one fossil chosen uniformly at random in each replicate, a valid test quantity for the same reason as the fossil age above. None of the test quantities, including the attachment age, rejected uniformity under any of the three adjustments, and the smallest BH-adjusted $p$-value was 0.89 (Figure A1-2).

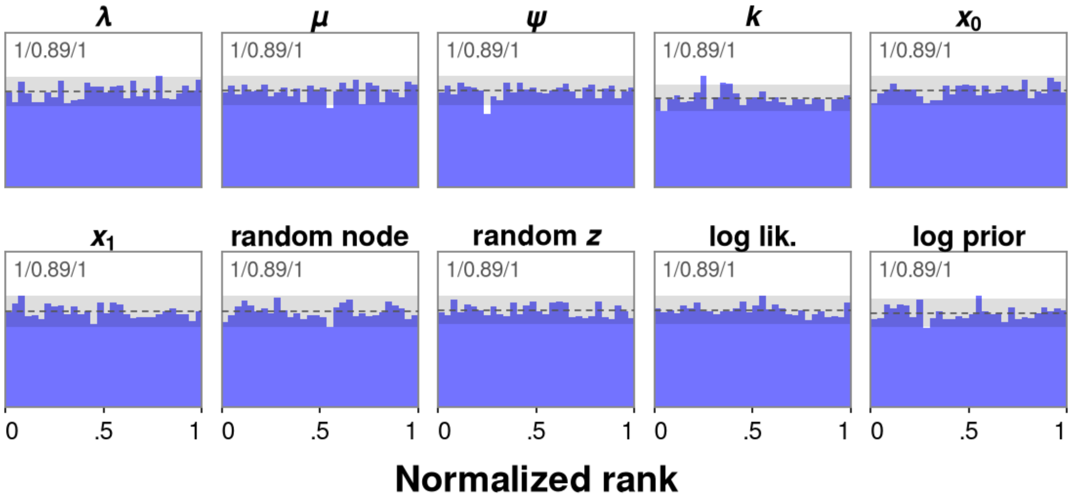


Figure A1-2: Results of the simulation-based calibration checking with nested clade assignments, shown as in Figure A1-1.

**Extinction conditioning and dynamic spine election** We simulated the FBD process conditioned on extinction with constant rates, $\lambda \sim \text{Unif}(0.03, 0.08)$, $\mu \sim \text{Unif}(0.02, 0.06)$, $\psi \sim \text{Unif}(0.7, 2.0)$, and $x_0 \sim \text{LogNormal}(4.0, 0.1^2)$. Simulated trees that did not satisfy the extinction condition were discarded and resimulated with the same parameter values, which matches the extinction conditioning. We ran two settings with extinct conditioning: first with fixed fossil age, and second with age ranges of width $w = 5$ as described above, in which the youngest fossil, and hence the spine, changes during the MCMC (dynamic spine election). In the latter, we additionally used the age of one fossil chosen uniformly at random in each replicate as a test quantity. In both settings, none of the test quantities rejected uniformity under any of the three adjustments, and the smallest BH-adjusted $p$-values were 0.10 and 0.90, respectively (Figure A1-3).

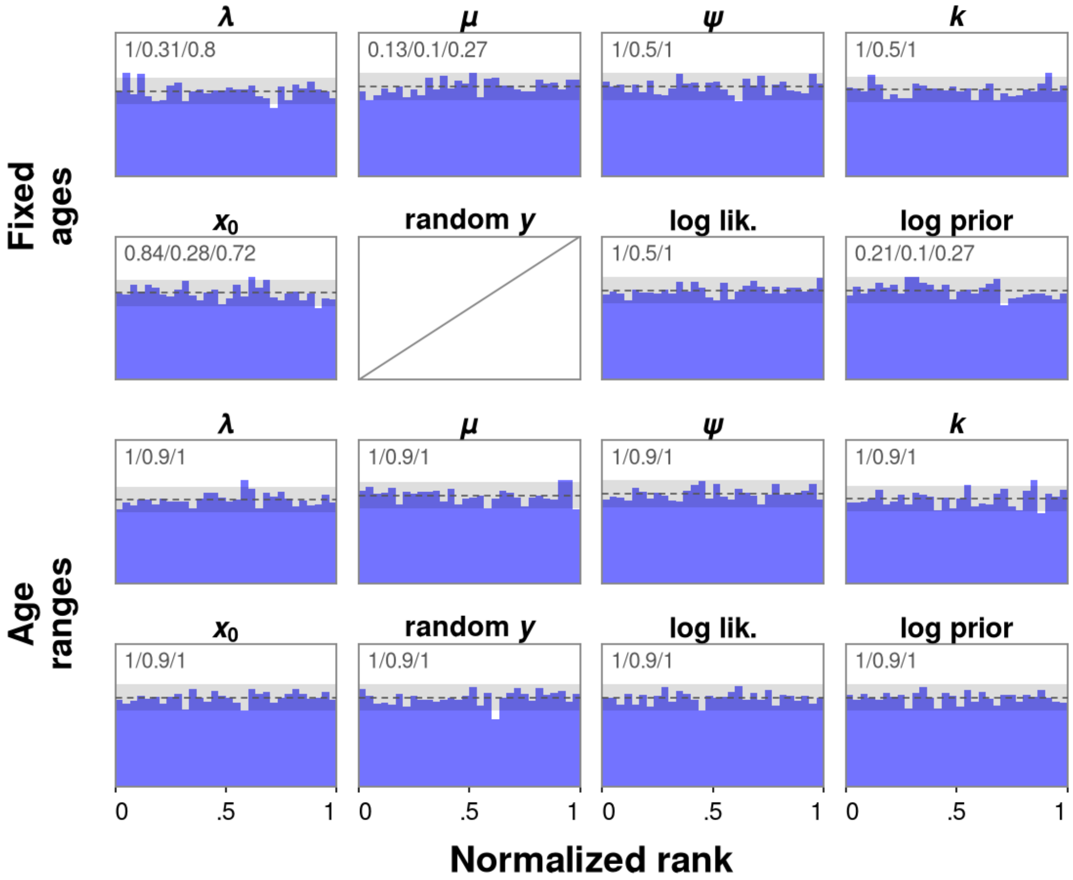


Figure A1-3: Results of the simulation-based calibration checking under extinction conditioning, with fixed fossil ages (top) and with age ranges (bottom), shown as in Figure A1-1.

### References


Altekar G, Dwarkadas S, Huelsenbeck JP, Ronquist F. 2004. Parallel Metropolis coupled Markov chain Monte Carlo for Bayesian phylogenetic inference. *Bioinformatics*. *20*(3):407–415. https://doi.org/10.1093/bioinformatics/btg427

Andrieu C, Thoms J. 2008. A tutorial on adaptive MCMC. *Statistics and Computing*. *18*(4):343–373. https://doi.org/10.1007/s11222-008-9110-y

dos Reis M, Yang Z. 2011. Approximate likelihood calculation on a phylogeny for Bayesian estimation of divergence times. *Molecular Biology and Evolution*. *28*(7):2161–2172. https://doi.org/10.1093/molbev/msr045

dos Reis M, Zhu T, Yang Z. 2014. The impact of the rate prior on Bayesian estimation of divergence times with multiple loci. *Systematic Biology*. *63*(4):555–565. https://doi.org/10.1093/sysbio/syu020

Drummond AJ, Ho SYW, Phillips MJ, Rambaut A. 2006. Relaxed phylogenetics and dating with confidence. *PLoS Biology*. *4*(5):e88. https://doi.org/10.1371/journal.pbio.0040088

Duchêne S, Ho SYW. 2014. Using multiple relaxed-clock models to estimate evolutionary timescales from DNA sequence data. *Molecular Phylogenetics and Evolution*. *77*:65–70. https://doi.org/10.1016/j.ympev.2014.04.010

Duchêne S, Ho SYW. 2015. Mammalian genome evolution is governed by multiple pacemakers. *Bioinformatics*. *31*(13):2061–2065. https://doi.org/10.1093/bioinformatics/btv121

Felsenstein J. 1981. Evolutionary trees from DNA sequences: A maximum likelihood approach. *Journal of Molecular Evolution*. *17*(6):368–376. https://doi.org/10.1007/BF01734359

Foster CSP, Ho SYW. 2017. Strategies for partitioning clock models in phylogenomic dating: Application to the angiosperm evolutionary timescale. *Genome Biology and Evolution*. *9*(10):2752–2763. https://doi.org/10.1093/gbe/evx198

Gavryushkina A, Welch D, Stadler T, Drummond AJ. 2014. Bayesian inference of sampled ancestor trees for epidemiology and fossil calibration. *PLoS Computational Biology*. *10*(12):e1003919. https://doi.org/10.1371/journal.pcbi.1003919

Geyer CJ. 1991. Markov chain Monte Carlo maximum likelihood. In: *Computing Science and Statistics: Proceedings of the 23rd Symposium on the Interface*. Interface Foundation of North America. p. 156–163.

Gilks WR, Roberts GO. 1996. Strategies for improving MCMC. In: Gilks WR, Richardson S, Spiegelhalter DJ, editors. *Markov Chain Monte Carlo in Practice*. London: Chapman and Hall. p. 89–114.

Green PJ. 1995. Reversible jump Markov chain Monte Carlo computation and Bayesian model determination. *Biometrika*. *82*(4):711–732. https://doi.org/10.1093/biomet/82.4.711

Guindon S. 2010. Bayesian estimation of divergence times from large sequence alignments. *Molecular Biology and Evolution*. *27*(8):1768–1781. https://doi.org/10.1093/molbev/msq060

Heath TA, Huelsenbeck JP, Stadler T. 2014. The fossilized birth-death process for coherent calibration of divergence-time estimates. *Proceedings of the National Academy of Sciences of the United States of America*. *111*(29):E2957–E2966. https://doi.org/10.1073/pnas.1319091111

Heitjan DF, Rubin DB. 1991. Ignorability and coarse data. *The Annals of Statistics*. *19*(4):2244–2253. https://doi.org/10.1214/aos/1176348396

Höhna S, Landis MJ, Heath TA, Boussau B, Lartillot N, Moore BR, Huelsenbeck JP, Ronquist F. 2016. RevBayes: Bayesian phylogenetic inference using graphical models and an interactive model-specification language. *Systematic Biology*. *65*(4):726–736. https://doi.org/10.1093/sysbio/syw021

Jin Y, Brown RP. 2018. Partition number, rate priors and unreliable divergence times in Bayesian phylogenetic dating. *Cladistics*. *34*(5):568–573. https://doi.org/10.1111/cla.12223

Kahan W. 1965. Further remarks on reducing truncation errors. *Communications of the ACM*. *8*(1):40. https://doi.org/10.1145/363707.363723

Kishino H, Thorne JL, Bruno WJ. 2001. Performance of a divergence time estimation method under a probabilistic model of rate evolution. *Molecular Biology and Evolution*. *18*(3):352–361. https://doi.org/10.1093/oxfordjournals.molbev.a003811

Kobert K, Stamatakis A, Flouri T. 2017. Efficient detection of repeating sites to accelerate phylogenetic likelihood calculations. *Systematic Biology*. *66*(2):205–217. https://doi.org/10.1093/sysbio/syw075

Lingenheil M, Denschlag R, Mathias G, Tavan P. 2009. Efficiency of exchange schemes in replica exchange. *Chemical Physics Letters*. *478*(1–3):80–84. https://doi.org/10.1016/j.cplett.2009.07.039

Miasojedow B, Moulines E, Vihola M. 2013. An adaptive parallel tempering algorithm. *Journal of Computational and Graphical Statistics*. *22*(3):649–664. https://doi.org/10.1080/10618600.2013.778779

Müller NF, Bouckaert RR. 2020. Adaptive Metropolis-coupled MCMC for BEAST 2. *PeerJ*. *8*:e9473. https://doi.org/10.7717/peerj.9473

Neumaier A. 1974. Rundungsfehleranalyse einiger Verfahren zur Summation endlicher Summen. *Zeitschrift für Angewandte Mathematik und Mechanik*. *54*:39–51.

Ogita T, Rump SM, Oishi S. 2005. Accurate sum and dot product. *SIAM Journal on Scientific Computing*. *26*(6):1955–1988. https://doi.org/10.1137/030601818

Okabe T, Kawata M, Okamoto Y, Mikami M. 2001. Replica-exchange Monte Carlo method for the isobaric–isothermal ensemble. *Chemical Physics Letters*. *335*(5–6):435–439. https://doi.org/10.1016/S0009-2614(01)00055-0

Papaspiliopoulos O, Roberts GO, Sköld M. 2003. Non-centered parameterizations for hierarchical models and data augmentation. In: Bernardo JM, Bayarri MJ, Berger JO, Dawid AP, Heckerman D, Smith AFM, West M, editors. *Bayesian Statistics 7: Proceedings of the Seventh Valencia International Meeting*. Oxford University Press. p. 307–326. https://doi.org/10.1093/oso/9780198526155.003.0017

Papaspiliopoulos O, Roberts GO, Sköld M. 2007. A general framework for the parametrization of hierarchical models. *Statistical Science*. *22*(1):59–73. https://doi.org/10.1214/088342307000000014

Pasarica C, Gelman A. 2010. Adaptively scaling the Metropolis algorithm using expected squared jumped distance. *Statistica Sinica*. *20*(1):343–364.

Robbins H, Monro S. 1951. A stochastic approximation method. *The Annals of Mathematical Statistics*. *22*(3):400–407. https://doi.org/10.1214/aoms/1177729586

Roberts GO, Rosenthal JS. 2007. Coupling and ergodicity of adaptive MCMC. *Journal of Applied Probability*. *44*(2):458–475. https://doi.org/10.1239/jap/1183667414

Seo TK, Kishino H, Thorne JL. 2004. Estimating absolute rates of synonymous and nonsynonymous nucleotide substitution in order to characterize natural selection and date species divergences. *Molecular Biology and Evolution*. *21*(7):1201–1213. https://doi.org/10.1093/molbev/msh088

Stamatakis A, Alachiotis N. 2010. Time and memory efficient likelihood-based tree searches on phylogenomic alignments with missing data. *Bioinformatics*. *26*(12):i132–i139. https://doi.org/10.1093/bioinformatics/btq205

Stamatakis A, Ott M. 2008. Efficient computation of the phylogenetic likelihood function on multi-gene alignments and multi-core architectures. *Philosophical Transactions of the Royal Society B: Biological Sciences*. *363*(1512):3977–3984. https://doi.org/10.1098/rstb.2008.0163

Syed S, Bouchard-Côté A, Deligiannidis G, Doucet A. 2022. Non-reversible parallel tempering: A scalable highly parallel MCMC scheme. *Journal of the Royal Statistical Society Series B: Statistical Methodology*. *84*(2):321–350. https://doi.org/10.1111/rssb.12464

Thawornwattana Y, Dalquen DA, Yang Z. 2018. Designing simple and efficient Markov chain Monte Carlo proposal kernels. *Bayesian Analysis*. *13*(4):1037–1063. https://doi.org/10.1214/17-BA1084

Thorne JL, Kishino H. 2002. Divergence time and evolutionary rate estimation with multilocus data. *Systematic Biology*. *51*(5):689–702. https://doi.org/10.1080/10635150290102456

Thorne JL, Kishino H, Painter IS. 1998. Estimating the rate of evolution of the rate of molecular evolution. *Molecular Biology and Evolution*. *15*(12):1647–1657. https://doi.org/10.1093/oxfordjournals.molbev.a025892

Uhlenbeck GE, Ornstein LS. 1930. On the theory of Brownian motion. *Physical Review*. *36*(5):823–841. https://doi.org/10.1103/PhysRev.36.823

Yang Z, Rodríguez CE. 2013. Searching for efficient Markov chain Monte Carlo proposal kernels. *Proceedings of the National Academy of Sciences of the United States of America*. *110*(48):19307–19312. https://doi.org/10.1073/pnas.1311790110

Zhang R, Drummond AJ. 2020. Improving the performance of Bayesian phylogenetic inference under relaxed clock models. *BMC Evolutionary Biology*. *20*:54. https://doi.org/10.1186/s12862-020-01609-4
