## Supplementary Material 2 for "Estimating Divergence Times and Diversification Rates with the Unresolved Fossilized Birth-Death Process"

### Supplementary Material 2: Paleontological Records for Empirical Datasets

#### 1 Fossil Records

Here, we describe the fossil records and, for the two analyses with a backbone tree (Cetacea, Emydidae), the unsampled extant taxa, of the three empirical datasets, together with the justification for each record's age, and when the analysis used a fixed backbone, its clade assignment. Candidate occurrences were compiled from the Paleobiology Database (https://paleobiodb.org; accessed July 13–26, 2026), and then audited individually against the primary literature, including records uncharted on PBDB when needed. We assigned valid records that were identified to species level for which a reasonable age estimate could be established, and excluded records such as *nomina dubia* and *nomina nuda*, occurrences assigned to a clade above species level, and taxa without stratigraphic age control (e.g., excluding whale fossils from trawl surveys; Bianucci et al. (2007)). We also excluded all records entirely bracketed within the Holocene, or records with stratigraphic range hitting 0, and fossils of extant species. When there were multiple fossil specimens per species, we equated each fossil's age range with the age range of its holotype's. When the holotype lacked an age estimate, we used the age estimate of the best-dated, validly described specimen. When no reliable specimen existed, we retain the fossil with a conservative wide age bracket. Ages are given as a minimum–maximum interval spanning the stratigraphic uncertainty. For the datasets with backbone phylogeny, each fossil additionally carries a phylogenetic placement: i.e., stem, crown, or total group.

##### 1.1 Cetacea

***Acrophyseter deinodon* Lambert et al. 2008** Holotype (MNHN SAS 1626) from the Montemar horizon of the Pisco Formation (Lambert et al., 2008), placed in the late Miocene by the Pisco Formation chronology of Ochoa et al. (2021), giving 6.6–7.9 Mya (superseding the older estimate of de Muizon & DeVries, 1985). Stem Physeteroidea: a macroraptorial *Zygophyseter*-clade sperm whale recovered outside the crown in all recent analyses (Peri et al., 2022; Lambert et al., 2023).

***Acrophyseter robustus* Lambert et al. 2017** Holotype (MUSM 1399) from the "La Bruja vertebrate level" (basal allomember P2) at Cerro la Bruja, Pisco Formation (Lambert et al., 2017a), dated to the late Miocene by $^{40}$Ar/$^{39}$Ar tephrochronology within the allomember (P1 8.9–9.5 Mya; P2 <8.5 Mya) (Di Celma et al., 2017), giving 7.45–9.0 Mya. (The describers instead give a >9.2 Mya late Serravallian–Tortonian age for the type horizon; we follow the sequence-stratigraphic allomember placement.) Stem Physeteroidea: see *Acrophyseter deinodon*.

***Adicetus latus* (Kellogg 1941)** Holotype from Adiça beach (Costa de Caparica, Almada, Portugal), in Cotter's division VIIb of the Lisbon Miocene (Kellogg, 1941; Figueiredo et al., 2024), placed in the lower Tortonian at ca. 10 Mya (Figueiredo et al., 2024), giving 7.25–11.63 Mya. Crown Mysticeti: recovered within Cetotheriidae *sensu stricto* by Bisconti et al. (2025a).

***Adicetus vandelli* (Van Beneden & Gervais 1868)** Holotype from Adiça beach (Costa de Caparica, Almada, Portugal), in Cotter's division VIIb of the Lisbon Miocene (Van Beneden & Gervais, 1880; Figueiredo et al., 2024), placed in the lower Tortonian at ca. 10 Mya (Figueiredo et al., 2024), giving 7.25–11.63 Mya. Crown Mysticeti: see *Adicetus latus*.

***Aegicetus gehennae* Gingerich et al. 2019** Holotype (CGM 60584) from the lower whale-bearing interval of the Gehannam Formation at Wadi Al Hitan (Gingerich et al., 2019), in planktonic-foraminiferal Zone E14, calibrated to 38.0–37.54 Mya (Wade et al., 2021), giving 37.54–38.00 Mya. Stem Cetacea.

***Aegyptocetus tarfa* Bianucci & Gingerich 2011** Holotype (MSNTUP I-15459), a cranium with associated postcranium recovered from commercial facing stone traced to the Gebel Hof Formation at Wadi Tarfa, Eastern Desert of Egypt (Bianucci & Gingerich, 2011), of early Bartonian (Mokattamian MK4–MK6) age, giving 40.00–41.03 Mya. Total group Cetacea.

***Aetiocetus cotylalveus* Emlong 1966** Holotype from the Yaquina Formation of Oregon (Emlong, 1966), dated to the late Rupelian–early Chattian (magnetochrons C12n–C9r) of the formation (Poust & Boessenecker, 2018), giving 27.4–30.6 Mya. Stem Mysticeti: an aetiocetid, member of one of the basal clades of archaic toothed mysticetes (Marx & Fordyce, 2015).

***Aetiocetus palmadentis* (Solís-Añorve et al. 2019)** Holotype (MUEcSj 5/24/123) from the San Juan Member of the El Cien Formation (Solís-Añorve et al., 2019), dated to the late Oligocene by a U–Pb zircon age on a tuff in the San Juan Member (Rochín-Bañaga et al., 2024), giving 27.79–28.11

Mya. Stem Mysticeti: see *Aetiocetus cotylalveus.*

***Aetiocetus polydentatus* Barnes et al. 1995** Holotype from the uppermost Upper Tuffaceous Siltstone Member of the Morawan Formation, Hokkaido, assigned to the late Oligocene (Barnes et al., 1995), giving 23.04–27.3 Mya. Stem Mysticeti: see *Aetiocetus cotylalveus.*

***Aetiocetus weltoni* Barnes et al. 1994** Holotype from near the top of the Yaquina Formation, close to the contact with the overlying Nye Mudstone (Barnes et al., 1995), of late Oligocene age, giving 27.4–30.6 Mya. Stem Mysticeti: see *Aetiocetus cotylalveus.*

***Aglaocetus burtinii* (Van Beneden 1859)** Lectotype (IRScNB CtM 677) probably from the Antwerpen Sand Member of the Berchem Formation, Antwerp (Van Beneden, 1859; Steeman, 2010), dinocyst-dated to the Langhian–Serravallian (Louwye et al., 2000, 2020), giving 12.8–15.97 Mya. Total group Mysticeti: a "cetothere"-grade baleen whale Steeman (2010).

***Aglaocetus latifrons* (Van Beneden 1880)** Lectotype (IRScNB CtM 567/Reg 198, a left periotic) from the Miocene of the Antwerp area, Belgium (Van Beneden, 1880; Steeman, 2010), placed in the middle Miocene (Langhian–Serravallian) of the southern North Sea Basin (Steeman, 2010), giving 11.63–15.98 Mya. Total group Mysticeti: see *Aglaocetus burtinii.*

***Aglaocetus longifrons* (Van Beneden 1880)** Lectotype (IRScNB CtM 719a-c) probably from the Antwerpen Sand Member of the Berchem Formation, Antwerp (Van Beneden, 1880; Steeman, 2010), dinocyst-dated to the Langhian–Serravallian (Louwye et al., 2000, 2020), giving 12.8–15.97 Mya. Total group Mysticeti: see *Aglaocetus burtinii.*

***Aglaocetus moreni* (Lydekker 1894)** Holotype from the Gaiman Formation of Chubut, Argentina (Lydekker, 1894; Buono et al., 2017); the Gaiman overlies the Colhuehuapian Trelew Member of the Sarmiento Formation and so does not predate late Aquitanian (Buono et al., 2017), giving 15.98–20.45 Mya. Total group Mysticeti: see *Aglaocetus burtinii.*

***Aglaocetus rotundus* (Van Beneden 1880)** Lectotype (IRScNB CtM 594) probably from the Deurne Sand Member of the Diest Formation, Antwerp (Van Beneden, 1880; Steeman, 2010), mid–late Tortonian by the Deurne Sands dinocyst chronology (Louwye, 2002; Goolaerts et al., 2020), giving 7.6–9.5 Mya. Total group Mysticeti: transferred from *Amphicetus* to *Aglaocetus* by Steeman (2010); see *Aglaocetus burtinii.*

***Agorophius pygmaeus* (Müller 1849)** The type skull, now lost, and the referred cranium SC 2015.51.1 from the Ashley Formation near Charleston, South Carolina (Müller, 1849; Godfrey et al., 2016), dated by $^{87}$Sr/$^{86}$Sr correlation of the Ashley Formation (Weems et al., 2016; Boessenecker & Geisler, 2018), giving 26.57–29.0 Mya. Total group Odontoceti.

***Albacetus salvifactus* Bisconti et al. 2025** Holotype (MCEA G1621), a partial skull with associated cervical and thoracic vertebrae, from the Sant'Agata Fossili Marls at Piana Biglini near Alba (Piedmont, Italy) (Bisconti et al., 2025a); the horizon is reported as upper Tortonian, but no horizon-specific age is published, so it is bracketed conservatively by the Tortonian stage at 7.246–11.63 Mya. Crown Mysticeti: a cetotheriid *sensu stricto* (Bisconti et al., 2025a).

***Albertocetus meffordorum* Uhen 2008** Holotype (USNM 525001), an ex-situ concretion assigned to the upper Oligocene Belgrade Formation at Onslow Beach, North Carolina (Uhen, 2008), its age constrained by $^{87}$Sr/$^{86}$Sr dating of the Belgrade Formation and of referred Ashley Formation material (Boessenecker et al., 2017a), giving 26.2–29.0 Mya. Stem Odontoceti: a xenorophid, the earliest-diverging clade within Odontoceti (Uhen, 2008; Boessenecker et al., 2017a).

***Albicetus oxymycterus* (Kellogg 1925)** Holotype (USNM 10923), the only known specimen, from the lower calcareous unit of the Monterey Formation on the sea cliffs at Santa Barbara, California (Kellogg, 1925a; Boersma & Pyenson, 2015), dated to the early middle Miocene (Langhian) by the Relizian–Luisian benthic foraminiferal stages and calcareous nannofossil zone CN4 (Boersma & Pyenson, 2015), giving 13.82–15.98 Mya. Stem Physeteroidea: recovered as a stem physeteroid by Boersma and Pyenson (2015).

***Albireo savagei* Barnes 2008** Holotype (UCMP 112048) from the Squire Member of the Pismo Formation, San Luis Obispo County, California (Barnes, 2008), of late Pliocene (Blancan) age (Boessenecker, 2013), giving 2–3 Mya. Total group Delphinoidea: see *Albireo whistleri*.

***Albireo whistleri* Barnes 1984** Holotype (UCR 14589, a skull and associated skeleton) from Kilmer's Discovery Locality in the lower member of the Almejas Formation, Isla Cedros, Baja California (Barnes, 1984), dated to the late Miocene by correlation of the lower Almejas Formation to 6–8 Mya (Boessenecker, 2013). Total group Delphinoidea: the type of the aberrant family Albireonidae.

***Allodelphis pratti* Wilson 1935** Holotype from the Pyramid Hill Sand Member of the Jewett Sand, Kern County, California (Wilson, 1935), dated by the San Joaquin Basin chronostratigraphy of

Scheirer and Magoon (2008), giving 24–25 Mya. Crown Odontoceti: an allodelphinid, recovered within crown Odontoceti (Boersma & Pyenson, 2016).

***Allodelphis woodburnei* Barnes & Reynolds 2009** Holotype (SBCM L3210-1) from the lower Vaqueros Formation at Cajon Pass, San Bernardino County, California, of early Miocene (Aquitanian) age (Barnes & Reynolds, 2009), giving 20.45–23.04 Mya. Crown Odontoceti: see *Allodelphis pratti*.

***Ambulocetus natans* Thewissen et al. 1994** Holotype (H-GSP 18507), a partial skeleton, from the Kuldana Formation of the Kala Chitta Hills, northern Pakistan (Thewissen et al., 1994), of middle Eocene age, giving 46–49 Mya. Stem Cetacea.

***Amphicetus later* Van Beneden 1880** Lectotype (IRScNB CtM 575) probably from the Deurne Sand Member of the Diest Formation, Antwerp (Van Beneden, 1880; Steeman, 2010), mid–late Tortonian (Louwye, 2002; Goolaerts et al., 2020), giving 7.6–9.5 Mya. Total group Mysticeti: retained as a "cetothere"-grade mysticete by Steeman (2010).

***Anacharsis orbus* Bogachev 1956** Holotype, a nearly complete skeleton from the Sarmatian near Stavropol in the northern Caucasus (Bogachev, 1956), giving 9.6–12.1 Mya (Palcu et al., 2021). Total group Odontoceti.

***Ancalecetus simonsi* Gingerich & Uhen 1996** Holotype (CGM 42290) from locality ZV-81 in the Birket Qarun Formation at Wadi Al Hitan (Gingerich & Uhen, 1996), Priabonian (nannofossil Zone NP19/20), giving 33.90–37.71 Mya. Stem Cetacea.

***Andrewsiphius sloani* (Sahni & Mishra 1972)** Holotype (LUVP 11002), a partial skull, from the Harudi Formation near Baranda, Kutch, western India (Sahni & Mishra, 1972, 1975), of middle Eocene age, giving 37.71–46.5 Mya. Stem Cetacea.

***Angelocetus cursiensis* Peri et al. 2022** Holotype (MSNUP I-16954) from a limestone quarry near the Cursi graveyard in the Pietra Leccese, Lecce Province, dated by its foraminiferal content to between the first and last common occurrences of *Paragloborotalia acrostoma* (Burdigalian) (Peri et al., 2022), giving 16.24–19.52 Mya. Crown Physeteroidea: recovered within the crown sperm whale clade, outside both Physeteridae and Kogiidae, by Peri et al. (2022).

***Ankylorhiza tiedemani* (Allen 1887)** Holotype (AMNH 10445) from the upper Oligocene Ashley Formation of South Carolina (Allen, 1887), with a referred skeleton from the younger Chandler Bridge

Formation (Boessenecker et al., 2020), both units $^{87}Sr/^{86}Sr$-dated (Boessenecker & Geisler, 2018), giving 23.5–29.0 Mya. Stem Odontoceti: recovered outside the crown as a large-bodied basal odontocete by Boessenecker et al. (2020).

***Anoplonassa forcipata* Cope 1869** Holotype (MCZ 8766), a mandibular symphysis from Chatham County, Georgia, near Savannah, with no recorded formation and coarse stratigraphic control. Thus we assigned the conservative age bracket of 2.58–23.03 Mya. Total group Ziphiidae.

***Antaecetus aithai* (Gingerich & Zouhri 2015)** Holotype (FSAC Bouj-6, a series of associated thoracic vertebrae) from the Aridal Formation at Gueran, south of Boujdour, southwestern Morocco (Gingerich & Zouhri, 2015), of middle Eocene (Bartonian) age, giving 37.71–41.03 Mya. Stem Cetacea.

***Antwerpibalaena liberatlas* Duboys de Lavigerie et al. 2020** Holotype (IRSNB M.2325), a partial skeleton from the base of the Oorderen Sand Member of the Lillo Formation at Kieldrecht Lock (Duboys de Lavigerie et al., 2020), its dinocyst assemblage giving a Piacenzian 2.76–3.21 Mya (Louwye et al., 2004; Duboys de Lavigerie et al., 2020). Total group Balaenidae: described as a stem balaenid (Duboys de Lavigerie et al., 2020) but recovered in a clade with the crown taxon *Eubalaena ianitrix* by Tanaka et al. (2025) and near *Balaenula*, and placed inside the crown by Bisconti et al. (2017).

***Aondelphis talen* Viglino et al. 2018** Holotype (MPEF-PV 517), a partial skull with tympanoperiotic, from the lower part of the Gaiman Formation at Bryn Gwyn, Chubut, Argentina (Viglino et al., 2018b); the lower Gaiman lies on the Colhuehuapian Trelew Member and is early Miocene (Buono et al., 2017; Viglino et al., 2018b), giving 15.98–20.45 Mya. Total group Odontoceti.

***Aporotus dicyrtus* (du Bus 1868)** Holotype (IRSNB 3808-M.541) from Antwerp with no recorded type horizon (Du Bus, 1872; Lambert, 2005c); the bracket spans the Antwerpen Sand Member base to the top of the Kattendijk Formation, giving 4.4–15.98 Mya. Total group Ziphiidae: a poorly known beaked whale recovered in the stem "*Messapicetus* clade" by Bianucci et al. (2016), assigned to the total group.

***Aporotus recurvirostris* (du Bus 1868)** Holotype (IRSNB 3812-M.1887) from the Miocene of the Antwerp area, Belgium (Du Bus, 1868), placed in the Serravallian–Tortonian of the southern North Sea Basin (Lambert, 2005c), giving 7.6–15.97 Mya. Total group Ziphiidae: see *Aporotus dicyrtus*.

***Aprixokogia kelloggi* Whitmore & Kaltenbach 2008** Holotype (USNM 187015) from the Yorktown Formation at the Lee Creek Mine, North Carolina (Whitmore & Kaltenbach, 2008), the Lee Creek Yorktown given a corrected early Pliocene age (Boessenecker et al., 2019), giving 3.92–4.9 Mya. Total group Kogiidae.

***Araeodelphis natator* Kellogg 1957** Holotype (USNM 10478) from Shattuck Zone 4–9, Plum Point Member of the Calvert Formation (Kellogg, 1957); along Calvert Cliffs, the species spans the Popes Creek Sand to Shattuck Zone 7, which the dinocyst chronology of Godfrey and Lambert (2023) brackets between the base of the Popes Creek Sand and the top of dinocyst zone DN3, giving 16.0–19.5 Mya. Crown Odontoceti: an early platanistid (Platanistoidea), nested within the crown odontocete (Godfrey & Lambert, 2023).

***Archaebalaenoptera castriarquati* Bisconti 2007** Holotype (SBAER 240536) from the first sandy level of the Rio Carbonari section at Castell'Arquato, Piacenza (Bisconti, 2007a), placed in the lower Piacenzian by ecobiostratigraphy (Bisconti, 2007a; Bisconti et al., 2020), giving 3.10–3.55 Mya. Total group Balaenopteroidea.

***Archaebalaenoptera eusebioi* Bisconti et al. 2021** Holotype, a nearly complete skull, from the Aguada de Lomas level of the Pisco Formation, southern Peru (Bisconti et al., 2021c), whose lowermost succession is bracketed to 8.85–7.93 Mya by K–Ar dating of two tuffs (Ochoa et al., 2021), giving 7.93–8.85 Mya. Total group Balaenopteroidea.

***Archaebalaenoptera liesselensis* Bisconti et al. 2020** Holotype (MAB002286) from Liessel; dated from the matrix of glauconitic sand cemented onto the skull to the late Tortonian Dinozone SNS M14 of the Diessen Formation (Bisconti et al., 2020), giving 7.5–8.1 Mya. Total group Balaenopteroidea.

***Archaeobalaena dosanko* Tanaka et al. 2020** Holotype (FCCP 1049) from the lower Chippubetsu Formation, below the T1 tuff (Tanaka et al., 2020), dated to the early Pliocene, giving 3.5–5.2 Mya. Total group Balaenidae.

***Archaeodelphis patrius* Allen 1921** Holotype from the "Tiger Leap" locality near Charleston, South Carolina, without a mapped formation (Allen, 1921a); the only direct age evidence is from the dinoflagellate-cyst assemblage in the adhering matrix, and the bracket spans the Charleston Oligocene cetacean-bearing units from the Ashley to the Chandler Bridge Formation (Weems et al., 2016), giving

23.5–29.0 Mya. Total group Odontoceti.

***Archaeophocaena teshioensis* Murakami et al. 2012** Holotype from the upper Miocene Koetoi Formation, Hokkaido, dated to diatom Zone NPD 7A (Murakami et al., 2012b), giving 5.5–6.4 Mya. Total group Phocoenidae: recovered outside crown Phocoenidae by Colpaert et al. (2015).

***Archaeoziphius microglenoideus* Lambert & Louwye 2006** Holotype (IRSNB 3856-M.1853), a cranium from Kessel in the Antwerpen Sand Member of the Berchem Formation (Lambert & Louwye, 2006), the cetacean-bearing sediment dated to the late Langhian–early Serravallian (Louwye et al., 2000; Lambert & Louwye, 2006), giving 12.8–15.97 Mya. Total group Ziphiidae: recovered within the subfamily Berardiinae (with *Berardius* and *Microberardius*), inside crown Ziphiidae, by Bianucci et al. (2016) and Kawatani and Kohno (2021), but its position seems unstable.

***Archaeschrichtius ruggieroi* Bisconti & Varola 2006** Holotype from the Cisterna quarry in the upper Pietra Leccese, Apulia (Bisconti & Varola, 2006), dated to the early Tortonian on the cetacean-bearing levels of that quarry (Bianucci & Landini, 2006), giving 8.14–10.5 Mya. Total group Balaenopteroidea.

***Argyrocetus bakersfieldensis* (Wilson 1935)** Holotype from the Pyramid Hill Sand Member of the Jewett Sand, Kern County, California (Wilson, 1935), dated by the San Joaquin Basin chronostratigraphy of Scheirer and Magoon (2008), giving 24–25 Mya. Total group Odontoceti.

***Argyrocetus joaquinensis* Kellogg 1932** Holotype from the upper Vedder Sand near its contact with the Jewett Sand, Kern County, California (Kellogg, 1932), dated by the San Joaquin Basin chronostratigraphy of Scheirer and Magoon (2008), giving 24–25 Mya. Total group Odontoceti.

***Argyrocetus patagonicus* Lydekker 1894** Holotype from the lower part of the Gaiman Formation at Bryn Gwyn, Chubut, Argentina (Lydekker, 1894; Lambert et al., 2025); the Gaiman overlies the Colhuehuapian Trelew Member and does not predate late Aquitanian (Buono et al., 2017), giving 15.98–20.45 Mya. Total group Odontoceti: tentatively grouped with the eastern Pacific homodont clade (*Chilcacetus*, *Eoplatanista*, eurhinodelphinids) (Lambert et al., 2025).

***Arimidelphis sorbinii* Bianucci 2005** Holotype from the upper Pliocene of the Marecchia River valley (Poggio Berni), central-eastern Italy (Bianucci, 2005), giving 1.80–3.60 Mya. Crown Delphinidae: recovered within Orcininae, in a clade with the extant *Orcinus* and *Pseudorca*, by Bianucci (2005).

***Arktocara yakataga* Boersma & Pyenson 2016** Holotype (USNM 214830), a partial skull from the Poul Creek Formation of the Yakataga district, southeastern Alaska (Boersma & Pyenson, 2016), dated to the Oligocene by detrital-zircon fission-track ages on the formation that do not exclude a Rupelian (Boersma & Pyenson, 2016), giving 24.0–29.0 Mya. Crown Odontoceti: recovered within Platanistoidea as an allodelphinid by Boersma and Pyenson (2016).

***Artiocetus clavis* Gingerich et al. 2001** Holotype (GSP-UM 3458), a skull with much of the skeleton, from the upper Habib Rahi Formation at Kunvit, Kohlu District, eastern Balochistan, Pakistan (Gingerich et al., 2001b), of early middle Eocene (Lutetian) age, giving 45.5–48 Mya. Stem Cetacea.

***Ashleycetus planicapitis* Sanders & Geisler 2015** Holotype (ChM PV6003), a partial skull from the Ashley Formation near Charleston, South Carolina (Sanders & Geisler, 2015), dated by $^{87}Sr/^{86}Sr$ correlation of the Ashley Formation (Weems et al., 2016; Boessenecker & Geisler, 2018), giving 26.57–29.0 Mya. Total group Odontoceti.

***Astadelphis gastaldii* (Brandt 1874)** Holotype (MGPT-PU 13883) from the Sabbie d'Asti (Asti Sands) at Ca' Lunga, Asti (Brandt, 1874; Bianucci, 1996), a late Zanclean–early Piacenzian unit, giving 3.00–4.00 Mya. Crown Delphinidae: recovered within Delphininae by Bianucci (2013).

***Atlanticetus lavei* Bisconti et al. 2021** Holotype, partial skeleton from the Burdigalian Pietra da Cantoni Group of Piedmont (Bisconti et al., 2021a), giving 16.0–19.0 Mya. Total group Mysticeti.

***Atlanticetus patulus* (Kellogg 1968)** Holotype (USNM 23690), a skull with tympanic bullae, left periotic and vertebrae, from Shattuck Zone 14 of the Calvert Formation at Stratford Bluffs, Westmoreland County, Virginia (Kellogg, 1968), dated to the middle Miocene by the Potomac River cliffs stratigraphy (Weems et al., 2017), giving 13.5–14.6 Mya. Total group Mysticeti.

***Atocetus iquensis* Muizon 1988** Holotype from the late Miocene Pisco Formation of the Sacaco Basin, southern Peru (de Muizon, 1988), dated by the Sacaco-Basin vertebrate-level chronology of Ochoa et al. (2021), giving 7.45–9.0 Mya. Total group Delphinida.

***Atocetus nasalis* (Barnes 1985)** Holotype (LACM 30093) from the upper Monterey Formation at Laguna Niguel, Orange County, California (Barnes, 1985b), dated to the late Miocene by the diatom biostratigraphy of the upper Orange County Monterey assemblage (Parham et al., 2022), giving 8.0–10.0 Mya. Total group Delphinida.

***Atropatenocetus posteocenicus* Aslanova 1977** Holotype from the Oligocene Maikop Group of Azerbaijan (Aslanova, 1977), giving 23.04–27.3 Mya. Total group Odontoceti.

***Attockicetus praecursor* Thewissen & Hussain 2000** Holotype (H-GSP 96232), a fragmentary skull, from a horizon near the Kuldana–Kohat formational contact in the Kala Chitta Hills, northern Pakistan (Thewissen & Hussain, 2000), of middle Eocene age, giving 46–49 Mya. Stem Cetacea.

***Aulophyseter mediatlanticus* (Cope 1895)** Holotype (USNM V 9463), a partial skull with proximal rostrum, from the St. Marys Formation at Drum Point, near the mouth of the Patuxent River, Calvert County, Maryland (Cope, 1895; Case, 1904), from the Little Cove Point Member dated to the early Tortonian by the Calvert Cliffs dinocyst stratigraphy (Kidwell et al., 2015; Weems et al., 2017; Godfrey & Lambert, 2023), giving 8.7–11.9 Mya. Total group Physeteroidea.

***Aulophyseter morricei* Kellogg 1927** Holotype from the Sharktooth Hill bonebed of the Round Mountain Silt (Kellogg, 1927b), correlated with magnetochron C5Br by the magnetostratigraphy of Pyenson et al. (2009), giving 15.2–16.0 Mya. Crown Physeteroidea: placed within Physeteridae (Peri et al., 2022; Lambert et al., 2023).

***Aureia rerehua* Meekin et al. 2024** Holotype (OU 22553) from the Otekaike Limestone at Haughs Quarry, Hakataramea Valley (Meekin et al., 2024), directly $^{87}$Sr/$^{86}$Sr-dated on the holotype to 23.10 ± 0.45 Mya (Marx et al., 2024) and independently constrained by foraminifera composition in the matrix to 22–23 Mya (Meekin et al., 2024), giving 22.0–23.55 Mya. Total group Odontoceti: a waipatiid-grade platanistoid whose crown-versus-stem position is unresolved, recovered outside crown Odontoceti by Meekin et al. (2024).

***Auroracetus bakerae* Gibson & Geisler 2009** Holotype from the Sunken Meadow Member of the Yorktown Formation at the Aurora (Lee Creek) Mine, North Carolina (Gibson & Geisler, 2009) giving 3.92–4.9 Mya (Boessenecker et al., 2019). Total group Inioidea.

***Australithax intermedia* Muizon 1988** Holotype from the El Jahuay (ELJ) horizon of the Pisco Formation (de Muizon, 1988), dated to the Tortonian by U–Pb zircon geochronology (Ochoa et al.,

2022), giving 9.0–9.6 Mya. Stem Phocoenidae: recovered as a stem member by Colpaert et al. (2015), falling outside crown Phocoenidae.

***Australodelphis mirus* Fordyce et al. 2002** Holotype (CPC 25730) from the Sørsdal Formation, Vestfold Hills, East Antarctica (Fordyce et al., 2002), of early Pliocene age, giving 4.1–4.5 Mya. Total group Delphinidae.

***Awadelphis hirayamai* Murakami 2016** Holotype from the uppermost Miocene Senhata Formation on the Boso Peninsula, Chiba (Murakami, 2016), dated by fission-track ages on the tephras bracketing its horizon (Murakami, 2016), giving 5.7–6.3 Mya. Total group Inioidea.

***Awamokoa tokarahi* Tanaka & Fordyce 2016** Holotype (OU 22125) from the Kokoamu Greensand at Tokarahi, Waitaki Valley (Tanaka & Fordyce, 2016), of Duntroonian age (Boessenecker & Richards, 2024; Robinson et al., 2024), giving 25.2–27.3 Mya. Total group Odontoceti.

***Babiacetus indicus* Trivedi & Satsangi 1984** Holotype (GSI 19647), a mandible, from the Harudi Formation of the Babia Hills, Kutch, western India (Trivedi & Satsangi, 1984), of late middle Eocene (Bartonian) age, giving 37.71–42 Mya. Stem Cetacea.

***Babiacetus mishrai* Bajpai & Thewissen 1998** Holotype (RUSB 2512), a partial skull, from the Harudi Formation of Kutch, western India (Bajpai & Thewissen, 1998), of late middle Eocene (Bartonian) age, giving 37.71–42 Mya. Stem Cetacea.

***Balaena meneghinii* (Capellini 1876)** Holotype from the Pliocene of Tuscany (Capellini, 1876), giving 2.58–5.33 Mya. Total group Balaenidae.

***Balaena montalionis* (Capellini 1904)** Holotype (MSNUP I-12357) from the Pliocene marine deposits near Montaione, Tuscany (Capellini, 1904), giving 2.58–5.33 Mya. Total group Balaenidae: recovered as the sister taxon of *Balaenella brachyrhynus* by Bisconti et al. (2017).

***Balaena ricei* Westgate & Whitmore 2002** Holotype from the Yorktown Formation at Rice's Pit, Hampton, Virginia (Westgate & Whitmore, 2002), spanning the Rushmere and Morgarts Beach Members, dated to the Piacenzian by the Yorktown dinoflagellate chronology (Dowsett et al., 2021), giving 3.15–3.3 Mya. Total group Balaenidae.

***Balaenella brachyrhynus* Bisconti 2005** Holotype (NMB 42001), an incomplete cranium from Kallo (Eerste Kanaaldok, 18 m b.s.l.) in the Kattendijk Formation, benthic-foraminiferal zone BFN 4 (Bisconti, 2005), early Zanclean by the Kattendijk dinocyst chronology (Louwye et al., 2004), giving 4.4–5.0 Mya. Total group Balaenidae: recovered closer to the crown genus *Balaena* than to *Eubalaena* by Bisconti (2005), and as sister to "*Balaena*" *montalionis* by Bisconti et al. (2017).

***Balaenoptera cephalus* (Cope 1867)** Holotype (ANSP 12691), paired mandibles from the Calvert Formation on the James T. Thomas farm near Hughesville, Charles County, Maryland (Cope, 1867), dated by a dinoflagellate flora recovered from the holotype's matrix to the upper part of dinoflagellate zone DN5 (Shattuck Zones 14–16A) (Weems & Edwards, 2007; Godfrey & Lambert, 2023), giving 13.0–14.6 Mya. Total group Balaenopteroidea.

***Balaenoptera davidsonii* (Cope 1872)** Holotype from the San Diego Formation of California (Cope, 1872), dated to the Pliocene–early Pleistocene range of the formation (Boessenecker, 2013), giving 1.8–4.2 Mya. Total group Balaenopteroidea.

***Balaenoptera siberi* Pilleri 1989** Holotype (skull and skeleton, A. Pezzia Asserto private collection) from the Aguada de Lomas (AGL) horizon of the Pisco Formation (Pilleri & Siber, 1989), dated to the Tortonian by K–Ar ages on tuffs within the AGL level (7.93 and 8.85 Mya) (Ochoa et al., 2021), giving 7.9–8.8 Mya. Total group Balaenopteroidea: a valid species according to Marx and Kohno (2016), but genus assignment likely needs revision (Bosselaers & Post, 2010).

***Balaenoptera sursiplana* Cope 1895** Holotype, a single tympanic bulla stated to be from the "Yorktown formation of Maryland" (Cope, 1895); the named unit is Pliocene but is not mapped in Maryland, whose Chesapeake Group there is no younger than the Tortonian, and considering both records together give 3.0–11.63 Mya. Total group Balaenopteroidea.

***Balaenotus insignis* (Van Beneden 1872)** Holotype from the lower Pliocene of the Antwerp Basin (Stuyvenberg), Belgium (Van Beneden, 1872), giving 2.58–5.33 Mya. Total group Balaenidae.

***Balaenotus lawleyi* (Capellini 1876)** Holotype from the Pliocene of the Siena area, Tuscany (Capellini, 1876), giving 2.58–5.33 Mya. Total group Balaenidae.

***Balaenotus orcianensis* Pilleri 1987** Holotype from the Pliocene of Orciano Pisano, Tuscany (Pilleri, 1987), giving 2.58–3.60 Mya. Total group Balaenidae.

***Balaenula astensis* Trevisan 1942** Holotype from the Sabbie d'Asti at Portacomaro, Asti (Trevisan, 1942), a late Zanclean–early Piacenzian unit, giving 3.00–4.00 Mya. Total group Balaenidae: recovered outside the balaenid crown by Buono et al. (2017), who note the paraphyly of *Balaenula*.

***Balaenula balaenopsis* (Van Beneden 1872)** Holotype from the lower Pliocene of the Antwerp area, Belgium (Van Beneden, 1872), giving 2.6–5.3 Mya. Total group Balaenidae.

***Balaenula forsythmajori* (Pilleri 1987)** Holotype from the Pliocene of Tuscany (Pilleri, 1987), giving 2.58–3.60 Mya. Total group Balaenidae.

***Basilosaurus cetoides* (Owen 1839)** Holotype from the upper Eocene (Priabonian) Yazoo Formation of Louisiana (Owen, 1839; Uhen, 2013), giving 33.9–37.7 Mya. Stem Cetacea.

***Basilosaurus isis* (Andrews 1904)** Holotype from the Birket Qarun Formation at the west end of Birket Qarun (Gingerich, 1992); the species is represented at Wadi Al Hitan by abundant articulated skeletons in the late transgressive systems tract of the first Priabonian sequence (Peters et al., 2009), giving 33.90–37.71 Mya. Stem Cetacea.

***Basiloterus hussaini* Gingerich et al. 1997** Holotype (GSP-UM 3190), two lumbar vertebrae, from the Drazinda Formation at Bari Nadi, Sulaiman Range, Punjab, Pakistan (Gingerich et al., 1997), of late middle Eocene (Bartonian) age, giving 37.71–41.03 Mya. Stem Cetacea.

***Belemnoziphius compressus* (Huxley 1864)** Holotype, a phosphatised rostrum from the basal nodule bed of the Red Crag Formation, Suffolk (Huxley, 1864); the nodule bed is a condensed reworked lag whose cetacean bone is likely derived from older (Miocene) deposits, so its Gelasian deposition (2.58–3.2 Mya) can only servea as a minimum age. Total group Ziphiidae.

***Belonodelphis peruanus* Muizon 1988** Holotype from the upper Miocene Pisco Formation of the Sacaco Basin, southern Peru (de Muizon, 1988), dated by the Sacaco-Basin chronology of Ochoa et al. (2021), giving 6.71–9.5 Mya. Total group Odontoceti.

***Beneziphius brevirostris* Lambert 2005** Holotype (IRSNB ED002-M.1885), a cranium from the Antwerp area; palynology of sediment from its mesorostral groove gives the *Achomosphaera andalousiensis* Zone, early to mid-Serravallian (12.8–13.2 Mya) (Miján et al., 2017). Total group Ziphiidae.

***Berardius kobayashii* Kawatani & Kohno 2021** Holotype (SCM 5530-6) from the Tsurushi Formation at Sobama Beach, Ogi, Sado Island, Niigata (Kawatani & Kohno, 2021), dated to planktonic foraminiferal Zone PF4, above the Ogi Basalt Member (11.7–14.1 Mya) (Kawatani & Kohno, 2021), giving 11.5–12.3 Mya. Crown Ziphiidae: recovered nested within the extant subfamily Berardiinae, as sister to *Berardius minimus* in most of the most-parsimonious trees, by Kawatani and Kohno (2021).

***Bohaskaia monodontoides* Velez-Juarbe & Pyenson 2012** Holotype from the Morgarts Beach Member of the Yorktown Formation at Rice's Pit, Hampton, Virginia (Vélez-Juarbe & Pyenson, 2012), dated to the Piacenzian by the Yorktown dinoflagellate chronology (Dowsett et al., 2021), giving 3.0–3.6 Mya. Crown Monodontidae: placed within Monodontidae by Vélez-Juarbe and Pyenson (2012).

***Borealodon osedax* Shipps et al. 2019** Holotype (USNM 539939) from the Pysht Formation, Clallam County, Washington (Shipps et al., 2019), dated to the Oligocene by the Twin River Group magnetostratigraphy of Prothero et al. (2001), giving 26.5–30.5 Mya. Stem Mysticeti: recovered as a stem mysticete by Shipps et al. (2019).

***Brabocetus gigaseorum* Colpaert et al. 2015** Holotype (IRSNB M.2171), a partial skull from the Beverentunnel, found in the Kattendijk Formation 2 m below the Oorderen Sand Member in gray glauconitic sand (above the reworked basal gravel) (Colpaert et al., 2015), early Zanclean (Louwye et al., 2004), giving 4.4–5.0 Mya. Total group Phocoenidae: Colpaert et al. (2015) place it on the phocoenid stem and state that "no fossil species falls within crown Phocoenidae", but as this is the only analysis to include it, the total group is assigned.

***Brachydelphis jahuayensis* Lambert & Muizon 2013** Holotype (MNHN PPI 268a–c) from the El Jahuay vertebrate level of the Pisco Formation, Sacaco Basin, southern Peru (Lambert & de Muizon, 2013), dated by the El Jahuay chronology of Ochoa et al. (2022), giving 9.0–9.6 Mya. Total group Inioidea: a monophyletic Pontoporiidae including *Brachydelphis* (crown) is recovered by Lambert et al. (2018a), but Pyenson et al. (2015) leave it in an unresolved basal polytomy (Pyenson et al., 2015; Lambert et al., 2018a).

***Brachydelphis mazeasi* Muizon 1988** Holotype (MNHN.F.PPI266) from the "La Bruja vertebrate level" (basal allomember P2) at Cerro la Bruja, Pisco Formation (de Muizon, 1988), dated to the late Miocene by the basal-P2 allomember age (Di Celma et al., 2017), giving 7.45–9.0 Mya. Total group Inioidea: see *Brachydelphis jahuayensis*.

***Brandtocetus chongulek* Gol'din & Startsev 2014** Holotype from the Sarmatian of the Kerch Peninsula, Crimea (Gol'din & Startsev, 2014), giving 7.65–9.6 Mya (Palcu et al., 2021). Crown Mysticeti: recovered within Cetotheriidae by Gol'din and Startsev (2014, 2017).

***Brevirostrodelphis dividum* (True 1912)** Holotype (USNM 7278), a skeleton from the Calvert Cliffs one mile south of Chesapeake Beach, Maryland, with no horizon recorded (True, 1912); the interval was set based on the partial skull CMM-V-4815 from Shattuck Zone 11 of the Plum Point Member (Godfrey & Lambert, 2023), giving 14.5–16.1 Mya. Total group Delphinidae.

***Brujadelphis ankylorostris* Lambert et al. 2017** Holotype (MUSM 1400) from the southern Cerro la Bruja hills, allomember P1 of the Pisco Formation (Lambert et al., 2017b), dated to the Tortonian by the P1 allomember age (8.9–9.5 Mya) (Di Celma et al., 2017). Crown Inioidea: the Lambert et al. (2017b) recovers it as an inioid more closely related to *Inia* than to *Pontoporia*, i.e., descended from the *Inia*–*Pontoporia* group (Lambert et al., 2017b).

***Brygmophyseter shigensis* (Hirota & Barnes 1995)** Holotype (SFM-0001) from the upper member of the Bessho Formation at Shiga, Nagano (Hirota & Barnes, 1995), of middle Miocene (Langhian) age, giving 13.82–15.98 Mya. Stem Physeteroidea.

***Cammackacetus hazenorum* Godfrey & Lambert 2023** Holotype (USNM PAL 639715), a partial skull and the only known specimen of the genus, recovered offshore from Governor Run, Calvert Cliffs, Calvert County, Maryland; its horizon is constrained only to the St. Marys Formation (Godfrey & Lambert, 2023), spanning the late Miocene of the Calvert Cliffs (Kidwell et al., 2015; Weems et al., 2017), giving 7.5–11.9 Mya. Total group Delphinida.

***Caolodelphis milleri* Godfrey & Lambert 2023** Holotype from low in the Popes Creek Sand of the Fairhaven Member of the Calvert Formation, about 30 cm above its contact with the Eocene Nanjemoy Formation, Maryland, a horizon correlated with dinoflagellate zone DN2C of the late Early Miocene (Burdigalian) (Godfrey & Lambert, 2023), giving 19.0–19.5 Mya. Total group Odontoceti.

***Carolinacetus gingerichi* Geisler et al. 2005** Holotype (ChM PV5401) from the lower Cross Member of the Tupelo Bay Formation, Berkeley County, South Carolina (Geisler et al., 2005), dated to the Bartonian by calcareous-nannoplankton Zone NP17 (Geisler et al., 2005), giving 37.0–40.2 Mya. Stem Cetacea.

***Casatia thermophila* Bianucci et al. 2019** Holotype from lower Pliocene marginal-marine deposits of Tuscany (Bianucci et al., 2019), giving 4.50–5.10 Mya. Crown Monodontidae.

***Caviziphius altirostris* Bianucci & Post 2005** Holotype (NNML 447230, an isolated cranium) from the "Post Mioceen Basisgrind", at Steendorp, Belgium (Bianucci & Post, 2005); the lag mixes Late Oligocene to latest Miocene bone, and Bianucci and Post (2005) assigns a Late Miocene origin of the cranium most plausible, giving 5.33–11.63 Mya. Total group Ziphiidae.

***Cephalotropis coronatus* Cope 1896** Holotype (USNM 9352), a braincase fragment without recorded locality (Cope, 1896); the horizon is supplied by the referred skull USNM 489194, from the Little Cove Point Member of the St. Marys Formation, Calvert County, Maryland (Marx & Fordyce, 2015), dated to the Tortonian by the Calvert Cliffs dinocyst stratigraphy (Kidwell et al., 2015; Weems et al., 2017), giving 8.7–11.9 Mya. Total group Mysticeti.

***Cephalotropis nectus* Kellogg 1941** Holotype from Adiça beach (Costa de Caparica, Almada, Portugal), in Cotter's division VIIb of the Lisbon Miocene (Kellogg, 1941; Figueiredo et al., 2024), placed in the lower Tortonian at ca. 10 Mya (Figueiredo et al., 2024), giving 7.25–11.63 Mya. Total group Mysticeti: its cetotheriid affinities are contested (Marx et al., 2019) and it is recovered outside Cetotheriidae *sensu stricto*, with *Pinocetus* and Neobalaenidae, by Gol'din (2018).

***Ceterhinops longifrons* Leidy 1877** Holotype (ANSP 11420) from the "Miocene marl" of the Ashley River, a Charleston phosphate lag mixing Oligocene to Pleistocene units (Leidy, 1877; Boessenecker & Boessenecker, 2019); the only lower Miocene source in that lag is the Burdigalian Marks Head Formation, giving 15.98–20.45 Mya. Total group Odontoceti.

***Cetorhynchus atavus* Abel 1905** Holotype, a mandibular symphysis fragment from Deurne (Fort 1) in the Deurne Sand Member of the Diest Formation (Abel, 1905b), mid–late Tortonian (Louwye, 2002; Goolaerts et al., 2020), giving 7.6–9.5 Mya. Total group Ziphiidae.

***Cetorhynchus christoli* (Gervais 1861)** Holotype mandibular fragments from the Miocene of southern France (Gervais, 1861), giving 5.33–23.04 Mya. Total group Ziphiidae.

***Cetotheriophanes capellinii* (Brandt 1873)** Holotype from the Pliocene of Italy (Brandt, 1873b), giving 2.58–5.33 Mya. Total group Mysticeti.

***Cetotheriopsis lintianus* (Meyer 1849)** Holotype, a braincase from the late Oligocene Linz Sands (Linzer Sande) at Linz, Austria (von Meyer, 1849), giving 23.04–27.3 Mya. Total group Mysticeti.

***Cetotherium crassangulum* Cope 1895** Holotype from the Yorktown Formation at City Point, Virginia, with no member recorded (Cope, 1895), so the horizon spans the whole formation, dated to the Zanclean–Piacenzian by the improved Yorktown chronology (Dowsett et al., 2021), giving 2.74–4.9 Mya. Total group Mysticeti.

***Cetotherium furlongi* Kellogg 1925** Holotype (USNM 26521) from the Vaqueros Formation of California, of early Miocene age (Kellogg, 1925a), giving 15.98–20.45 Mya. Total group Mysticeti.

***Cetotherium gervaisii* (Van Beneden & Gervais 1868)** Holotype from the Miocene of France (Van Beneden & Gervais, 1880), giving 5.33–23.04 Mya. Total group Mysticeti.

***Cetotherium megalophysum* Cope 1895** Holotype (USNM 10593), a cranium retaining the right bulla, from the St. Marys Formation at Cove Point, Calvert County, Maryland (Cope, 1895; Case, 1904), from the Little Cove Point Member dated to the Tortonian by the Calvert Cliffs dinocyst stratigraphy (Kidwell et al., 2015; Weems et al., 2017), giving 8.7–11.9 Mya. Total group Mysticeti.

***Cetotherium polyporum* (Cope 1869)** Holotype (ANSP 12694) from the Pliocene Yorktown Formation at Bells Bridge, North Carolina (Cope, 1869a), from the range spanning the early to late Pliocene (Boessenecker et al., 2019), giving 2.74–4.9 Mya. Total group Mysticeti.

***Cetotherium rathkii* Brandt 1843** Holotype from the Sarmatian of the northern Black Sea region (Brandt, 1843), giving 7.65–9.6 Mya (Palcu et al., 2021). Crown Mysticeti.

***Cetotherium riabinini* Hofstein 1948** Holotype, a skull and partial skeleton from the Sarmatian near Mykolaiv, southern Ukraine (Hofstein, 1948; Gol'din et al., 2014), giving 7.65–9.6 Mya (Palcu et al., 2021). Crown Mysticeti: recovered as sister to *C. rathkii* within Cetotheriidae by Gol'din and Startsev (2017).

***Charadrobalaena valentinae* Bisconti et al. 2023** Holotype (MCRE 232834) from near San Valentino Castle, Castellarano (Emilia Romagna) (**bisconti2023**), dated to the late Zanclean by molluscs and foraminifera (**bisconti2023**), giving 3.60–3.80 Mya. Total group Balaenidae: placed in a stem balaenid clade, sister to the crown, by **bisconti2023**.

***Chavinziphius maxillocristatus* Bianucci et al. 2016** Holotype (MUSM 2538), an incomplete skull, from Member F of the Pisco Formation at Cerro los Quesos, ∼10 m above the Mono key bed (Bianucci et al., 2016), dated to the Messinian by $^{40}$Ar/$^{39}$Ar ages on the bracketing ashes (6.93±0.09 Mya below; 6.71±0.02 Mya above) (Bianucci et al., 2016), giving 6.69–7.02 Mya. Stem Ziphiidae: recovered as the earliest-diverging ziphiid by Bianucci et al. (2016) and as sister to the clade of extant subfamilies by Kawatani and Kohno (2021), i.e., outside crown Ziphiidae in both (Bianucci et al., 2016; Kawatani & Kohno, 2021).

***Chilcacetus cavirhinus* Lambert et al. 2015** Holotype (MNHN.F.PRU11) from the Chilcatay Formation of Peru (Lambert et al., 2015b); the holotype horizon within the formation is not dated, so we applied the whole-formation age. Dated to the early Miocene by $^{40}$Ar/$^{39}$Ar tephra ages of 19.25±0.05 Mya near the base at Zamaca and 18.02±0.07 Mya near the top at Ullujaya (Bianucci et al., 2020), giving 18.0–19.3 Mya. Total group Odontoceti: homodont "*Chilcacetus* grade" (Chilcacetidae), outside crown Platanistoidea (Bianucci et al., 2020).

***Chilcacetus ullujayensis* Lambert et al. 2025** Holotype (MUSM 2527) from the Ct1a facies of the Chilcatay Formation at Ullujaya (Lambert et al., 2025), dated to the early Miocene (Burdigalian) within the Chilcatay tephra/Sr framework (Bianucci et al., 2020), giving 18.3–18.9 Mya. Odontoceti (total): see *Chilcacetus cavirhinus*.

***Chimuziphius coloradensis* Bianucci et al. 2016** Holotype (MUSM 2548) from Cerro Colorado, 61.5 m above the basal unconformity in allomember P1 of the Pisco Formation (Bianucci et al., 2016), dated to the Tortonian by a $^{40}$Ar/$^{39}$Ar ash ∼30 m below the holotype at 9.10±0.04 Mya (Bianucci et al., 2016), giving 8.5–9.14 Mya. Total group Ziphiidae: recovered within the stem *Messapicetus* clade by Bianucci et al. (2016).

***Chonecetus sookensis* Russell 1968** Holotype from the late Oligocene Sooke Formation of Vancouver Island (Russell, 1968), giving 24.1–24.8 Mya. Stem Mysticeti: an aetiocetid, a family recovered among the basal toothed mysticetes outside crown Mysticeti by Marx and Fordyce (2015).

***Chonecetus tomitai* (Barnes et al. 1995)** Holotype from the Middle Hard Shale Member of the Morawan Formation, Hokkaido, assigned to the late Oligocene (Barnes et al., 1995), giving 23.04–27.3 Mya. Stem Mysticeti.

***Chonecetus yabukii* (Barnes et al. 1995)** Holotype (AMP 1, a skull) from the upper tuffaceous siltstone member of the Morawan Formation, Ashoro, Hokkaido, Japan (Barnes et al., 1995), of late Oligocene (Chattian) age, giving 23.04–27.3 Mya. Stem Mysticeti: an aetiocetid toothed baleen whale, recovered within Aetiocetidae by Peredo and Pyenson (2018).

***Choneziphius planirostris* (Cuvier 1823)** Holotype from the Miocene of the Antwerp area, Belgium (Cuvier, 1823), placed in the Serravallian–Tortonian of the southern North Sea Basin (Lambert, 2005c), giving 7.6–15.97 Mya. Total group Ziphiidae.

***Chrysocetus fouadassii* Gingerich & Zouhri 2015** Holotype (FSAC Bouj-1), a left dentary with P4–M1, from a sandstone bed ∼11 m above the base of the Aridal Formation at the Sabkha of Gueran, Western Sahara (Gingerich & Zouhri, 2015), of Bartonian age, giving 37.71–41.03 Mya. Stem Cetacea.

***Chrysocetus healyorum* Uhen & Gingerich 2001** Holotype (SCSM 87.195) from the Santee Cement Quarry at Holly Hill, Orangeburg County, in the Cross Formation (Uhen & Gingerich, 2001), dated to the late Bartonian (Uhen & Gingerich, 2001), giving 36.0–39.5 Mya. Stem Cetacea.

***Ciuciulea davidi* Gol'din 2018** Holotype (ZIRM V 28/1) from the late Badenian of the Fore-Carpathian Basin near Ciuciulea, Moldova (Gol'din, 2018), giving 12.65–13.82 Mya. Crown Mysticeti: recovered as the most basal cetotheriid in the analysis of Gol'din (2018).

***Cochimicetus convexus* Cedillo-Ávila et al. 2025** Holotype (MHN-UABCS-Sg6/71/208, a partial skull with mandibles) from the Mount "Pilón" locality of the San Gregorio Formation, Baja California Sur (Cedillo-Ávila et al., 2025), placed in the late Oligocene (Chattian) San Gregorio Formation, giving 23.04–27.3 Mya. Stem Mysticeti: an eomysticetid (Cedillo-Ávila et al., 2025), Eomysticetidae being the earliest-diverging clade of toothless mysticetes, recovered rootward of crown Mysticeti (Boessenecker & Fordyce, 2015c; Marx & Fordyce, 2015).

***Cophocetus oregonensis* Packard & Kellogg 1934** Holotype (UO 305) from the Astoria Formation near Newport, Lincoln County, Oregon, of early Miocene age (Packard & Kellogg, 1934), giving 15.98–20.45 Mya. Total group Mysticeti: with uncertain familial position (Marx et al., 2017).

***Coronodon havensteini* Geisler et al. 2017** Holotype (CCNHM 108), a skull with mandibles, vertebrae, and ribs from the Wando River, Berkeley County, Ashley Formation (Geisler et al., 2017), dated by $^{87}$Sr/$^{86}$Sr correlation of the Ashley Formation (Weems et al., 2016; Boessenecker & Geisler, 2018), giving 26.57–29.0 Mya. Stem Mysticeti: recovered among the most basal toothed mysticetes by Geisler et al. (2017) and Boessenecker et al. (2023).

***Coronodon newtonorum* Boessenecker et al. 2023** Holotype (ChM PV 2778) from Bed 3 of the Chandler Bridge Formation, South Carolina (Boessenecker et al., 2023), dated by $^{87}$Sr/$^{86}$Sr correlation of the Chandler Bridge Formation (Weems et al., 2016; Boessenecker & Geisler, 2018), giving 23.5–24.7 Mya. Stem Mysticeti: recovered among the basal toothed mysticetes within *Coronodon* by Boessenecker et al. (2023).

***Coronodon planifrons* Boessenecker et al. 2023** Holotype (CCNHM 166) from Bed 2 of the Chandler Bridge Formation, North Charleston, South Carolina (Boessenecker et al., 2023), dated by $^{87}$Sr/$^{86}$Sr correlation of the Chandler Bridge Formation (Weems et al., 2016; Boessenecker & Geisler, 2018), giving 23.5–24.7 Mya. Stem Mysticeti: see *Coronodon newtonorum*

***Cotylocara macei* Geisler et al. 2014** Holotype (CCNHM 101) from the College Park drainage ditch, Bed 2 of the Chandler Bridge Formation, South Carolina (Geisler et al., 2014), dated by $^{87}$Sr/$^{86}$Sr correlation of the Chandler Bridge Formation (Weems et al., 2016; Boessenecker & Geisler, 2018), giving 23.5–24.7 Mya. Stem Odontoceti: a xenorophid recovered in a basal odontocete clade by Geisler et al. (2014).

***Cozzuoliphyseter rionegrensis* (Gondar 1975)** Holotype (MLP 62-XII-19-1), a nearly complete skull, from the Gran Bajo del Gualicho Formation near San Antonio Oeste, Río Negro, Argentina (Gondar, 1975; Paolucci et al., 2021); the two type skulls cannot be assigned to a member, so a conservative whole Miocene age was assigned (Paolucci et al., 2021), giving 5.33–20.03 Mya. Total group Physeteroidea: recovered as a crown physeteroid within Physeteridae by Paolucci et al. (2021), but as a stem physeteroid (sister to the crown) by Peri et al. (2022) and Lambert et al. (2023).

***Crenatocetus rayi* McLeod & Barnes 2008** Holotype (USNM 392014), an undocumented quarry find from the Martin Marietta New Bern Quarry, Craven County, North Carolina, probably derived from the Comfort Member of the Castle Hayne Formation (McLeod & Barnes, 2008), the Comfort Member falling within calcareous-nannofossil Zone NP16 (Weems et al., 2011), giving 37.71–42.87 Mya. Stem Cetacea.

***Crisocetus lydekkeri* Gaetán et al. 2023** Holotype, a partial skull, from the Gaiman Formation at Estancia El Crisol, Chubut, Argentina (Gaetán et al., 2023); the Gaiman overlies the Colhuehuapian Trelew Member and does not predate late Aquitanian (Buono et al., 2017), giving 15.98–20.45 Mya. Total group Odontoceti: recovered within crown Odontoceti, rootward of delphinidans, ziphiids and physeteroids, with weakly supported squaloziphiid affinities (Gaetán et al., 2023).

***Cynthiacetus maxwelli* Uhen 2005** Holotype (MMNS VP-445) from the Cynthia Clay Pit in the Yazoo Clay (Jackson Group), Hinds County, Mississippi (Uhen, 2005), Priabonian, giving 33.90–37.71 Mya. Stem Cetacea.

***Cynthiacetus peruvianus* Martínez-Cáceres & Muizon 2011** Holotype (MNHN.F.PRU10), a nearly complete skull and skeleton, from the Otuma Formation at Paracas Bay, southern Peru (Martínez-Cáceres & de Muizon, 2011; Martínez-Cáceres et al., 2017); a $^{40}$Ar/$^{39}$Ar age of 36.61 ± 0.03 Mya on the ash at the base of the Otuma Formation bounds the maximum (Uhen et al., 2011), giving 33.9–36.64 Mya. Stem Cetacea.

***Dagonodum mojnum* Ramassamy 2016** Holotype (MSM1001x), a partial skull and skeleton from the Gram Brick Works in the Gram Clay of the Gram Formation, Denmark (Ramassamy, 2016), with an in-situ Tortonian assemblage giving 7.2–9.9 Mya. Total group Ziphiidae: recovered by Ramassamy (2016) as sister to *Messapicetus*, i.e. within the stem "*Messapicetus* clade" of Bianucci et al. (2016).

***Dalanistes ahmedi* Gingerich et al. 1995** Holotype (GSP-UM 3106), a partial skull, from the Domanda Formation at Basti Ahmed, Sulaiman Range, Punjab, Pakistan (Gingerich et al., 1995), of middle Eocene age, giving 37.71–46 Mya. Stem Cetacea.

***Dalpiazina ombonii* (Longhi 1898)** Holotype from the upper Libano Sandstone of the Belluno quarries (Longhi, 1898), dated to the late Aquitanian–early Burdigalian (Nelson et al., 2025c), giving 19.0–22.0 Mya. Total group Odontoceti.

***Delphinavus newhalli* Lull 1914** Holotype from the middle to upper Miocene Monterey Formation of California (Lull, 1914), giving 9.4–12.5 Mya. Total group Odontoceti.

***Delphinodon carniolicus* (Gorjanović-Kramberger 1892)** Holotype from the Badenian of Carniola (Central Paratethys) (Gorjanović-Kramberger, 1892), dated to the Central Paratethys Badenian stage (Hohenegger et al., 2014), giving 12.83–16.30 Mya. Total group Delphinida.

***Delphinus rikuzenensis* Hatai et al. 1963** Holotype from the Tatsunokuchi Formation (Hatai et al., 1963), dated to the early Pliocene, giving 3.6–5.33 Mya. Total group Delphinidae.

***Denebola brachycephala* Barnes 1984** Holotype (UCR 21245, a skull) from Locality 11 in the lower member of the Almejas Formation, Isla Cedros, Baja California (Barnes, 1984), dated to the late Miocene by correlation of the lower Almejas Formation to 8–6 Mya (Boessenecker, 2013), giving 6.0–8.0 Mya. Total group Monodontidae: recovered as a stem monodontid (with *Haborodelphis*) by Bianucci et al. (2019), but on a 21-character matrix with weak support, so the total group is assigned.

***Dhedacetus hyaeni* Bajpai & Thewissen 2014** Holotype (IITR-SB 2870), a partial skull and skeleton, from the Harudi Formation near Dhedadi, Kutch, western India (Bajpai & Thewissen, 2014), of late middle Eocene (Bartonian) age, giving 37.71–42 Mya. Stem Cetacea.

***Diaphorocetus ortegai* Lambert et al. 2023** Holotype (MUSM 3246) from the Ct1a facies of the Chilcatay Formation at Ullujaya (Lambert et al., 2023), 18.3–18.9 Mya (Bianucci et al., 2020). Total group Physeteroidea: recovered as a stem physeteroid in the original description (Lambert et al., 2023), though *Diaphorocetus* falls inside the crown in other analyses (Alfsen et al., 2021; Peri et al., 2022).

***Diaphorocetus poucheti* (Moreno 1892)** Holotype (MLP 5-6) from the Gaiman Formation in the Puerto Madryn area, Chubut, Argentina (Moreno, 1892; Paolucci et al., 2020); the Gaiman overlies the Colhuehuapian Trelew Member and does not predate late Aquitanian (Buono et al., 2017), giving 15.98–20.45 Mya. Total group Physeteroidea: one of the earliest sperm whales, recovered as a stem physeteroid in a polytomy with the crown (Physeteridae+Kogiidae) by Paolucci et al. (2020).

***Dilophodelphis fordycei* Boersma et al. 2017** Holotype (USNM 214911) from the Astoria Formation at Nye Beach, Oregon (Boersma et al., 2017), dated to the early Burdigalian (Peredo et al., 2018b), giving 19–20.7 Mya. Crown Odontoceti: a platanistid, recovered within Platanistoidea (Boersma et al., 2017).

***Diorocetus chichibuensis* Yoshida et al. 2003** Holotype from the Nagura Formation, Chichibu Basin, Saitama (Yoshida et al., 2003), of early to middle Miocene age from the marine fill of the basin, giving 15.0–17.0 Mya. Total group Mysticeti.

***Diorocetus hiatus* Kellogg 1968** Holotype (USNM 16783), a nearly complete skull with the right periotic, mandibles and vertebrae, from Shattuck Zone 14 of the Calvert Formation south of the mouth of Parker Creek, Calvert County, Maryland (Kellogg, 1968), dated to the middle Miocene by the Calvert Cliffs stratigraphy (Kidwell et al., 2015), giving 13.5–14.6 Mya. Total group Mysticeti.

***Diorocetus shobarensis* Otsuka & Ota 2008** Holotype from the Korematsu Formation at Nakahara, Bihoku Group, Hiroshima (Otsuka & Ota, 2008), referred to the middle Miocene upper Bihoku Group, whose overlying Itahashi Formation falls in calcareous nannofossil Zone NN4 (Tsai, 2017), giving 15.6–16.9 Mya. Total group Mysticeti.

***Diunatans luctoretemergo* Bosselaers & Post 2010** Holotype (NHG 22279), a neurocranium with cervical vertebrae from the Scheldt estuary in the Kattendijk Sand Member of the Kattendijk Formation (Bosselaers & Post, 2010), Zanclean by the Kattendijk dinocyst chronology (Louwye et al., 2004), giving 4.4–5.0 Mya. Total group Balaenopteroidea: recovered as a stem balaenopterid, sister to the *Balaenoptera* + *Megaptera* clade, by Bosselaers and Post (2010).

***Dolgopolis kinchikafiforo* Viglino et al. 2021** Holotype, an incomplete skull, from the Gaiman Formation at Estancia El Crisol, Chubut, Argentina (Viglino et al., 2021); the Gaiman overlies the Colhuehuapian Trelew Member and does not predate late Aquitanian (Buono et al., 2017), giving 15.98–20.45 Mya. Total group Odontoceti.

***Dorudon atrox* (Andrews 1906)** Holotype (CGM 9319, a juvenile skull and partial skeleton) from the Birket Qarun Formation of the Fayum, Egypt (Andrews, 1906), of late Eocene (Priabonian) age, giving 33.9–37.7 Mya. Stem Cetacea.

***Dorudon serratus* Gibbes 1845** Holotype from the Mazyck Plantation locality in the greensand near Moncks Corner, South Carolina (Gibbes, 1845), dated to the late Eocene (Priabonian) by the revision of Uhen (2004), giving 33.9–37.71 Mya. Stem Cetacea.

***Echericetus novellus* Hernández-Cisneros et al. 2023** Holotype (MRAHBCS Pal/V119) from bed 2 of the San Juan Member, El Cien Formation, at Cerro El Divisadero, Baja California Sur, Mexico, capped by a reworked tuff with a U–Pb zircon age of 27.95 Mya (Hernández-Cisneros et al., 2023), giving 27.8–28.6 Mya. Stem Mysticeti: an eomysticetid (Hernández-Cisneros et al., 2023), a clade lying rootward of crown Mysticeti (Marx & Fordyce, 2015).

***Echovenator sandersi* Churchill et al. 2016** Holotype from the basal bed of the Chandler Bridge Formation, Berkeley County, South Carolina (Churchill et al., 2016), dated by $^{87}Sr/^{86}Sr$ correlation of the Chandler Bridge Formation (Weems et al., 2016; Boessenecker & Geisler, 2018), giving 23.5–24.7 Mya. Stem Odontoceti: a xenorophid recovered in the most basal odontocete clade by Churchill et al. (2016).

***Ediscetus osbornei* Albright et al. 2018** Holotype from the Givhans Ferry Member of the Ashley Formation, South Carolina (Albright et al., 2018), dated by $^{87}Sr/^{86}Sr$ correlation of the Ashley Formation (Weems et al., 2016; Boessenecker & Geisler, 2018), giving 26.57–29.0 Mya. Total group Odontoceti: recovered slightly outside the odontocete crown group by Albright et al. (2018), but as this was the only analysis to include it, the total group is retained.

***Enigmatocetus posidoni* Godfrey & Lambert 2023** Holotype (USNM PAL 244524), a fragmentary skull with both periotics and bullae, from the base of Shattuck Zone 10 (Plum Point Member) of the Calvert Formation by the George Washington Birthplace National Monument, Westmoreland County, Virginia (Godfrey & Lambert, 2023), dated to the middle Miocene by the Calvert Cliffs stratigraphy (Kidwell et al., 2015; Weems et al., 2017), giving 15.0–16.6 Mya. Total group Odontoceti.

***Ensidelphis riveroi* Bianucci et al. 2020** Holotype (MUSM 3898) from the Chilcatay Formation at Zamaca, 10.7 m above the contact with the underlying Otuma Formation (top of the Ct1c facies) (Bianucci et al., 2020), dated to the early Miocene by $^{40}Ar/^{39}Ar$ ages constraining Ct1c to 25±0.08–19.00±0.28 Mya (Bianucci et al., 2020), giving 18.7–19.3 Mya. Crown Odontoceti.

***Eobalaenoptera harrisoni* Dooley et al. 2004** Holotype, the only known occurrence, from the Carmel Church Quarry bone bed of the Calvert Formation, Caroline County, Virginia (Dooley et al., 2004), dated to the middle Miocene, giving 12.7–14.9 Mya. Total group Balaenopteroidea.

***Eocetus drazindai* (Gingerich et al. 1997)** Holotype (GSP-UM 3193), a lumbar vertebra, from the Drazinda Formation at Bari Nadi, Sulaiman Range, Punjab, Pakistan (Gingerich et al., 1997), of late middle Eocene (Bartonian) age, giving 37.71–41.03 Mya. Stem Cetacea.

***Eocetus schweinfurthi* (Fraas 1904)** Holotype (SMNS 10986), a skull from Schweinfurth's locality IV in the Giushi Formation at Gebel Mokattam (Fraas, 1904), Bartonian, giving 37.71–41.03 Mya. Stem Cetacea.

***Eodelphinus kabatensis* (Horikawa 1977)** Holotype from the Lower Mudstone of the Mashike Member (Horikawa, 1977; Murakami et al., 2014), dated to the late Miocene, giving 8.5–13.0 Mya. Total group Delphinidae.

***Eolipotes japonicus* Kimura & Hasegawa 2024** Holotype (TPM7847-1) from the upper Ogane Formation; the paratype (GMNH-PV-1011) lies in the Haraichi Formation between the Baba tuff (11.26±0.09 / 11.29±0.12 Mya) and the Mizuguchi tuff (ca. 11.25 Mya) (Kimura & Hasegawa, 2024b), giving 11.25–11.29 Mya. Crown Odontoceti: recovered within Lipotidae, in a clade with *Parapontoporia* and closer to it than to the extant *Lipotes vexillifer*, by Kimura and Hasegawa (2024b).

***Eomysticetus carolinensis* Sanders & Barnes 2002** Holotype from the Chandler Bridge Formation, South Carolina (Sanders & Barnes, 2002b), dated by $^{87}$Sr/$^{86}$Sr correlation of the Chandler Bridge Formation (Weems et al., 2016; Boessenecker & Geisler, 2018), giving 23.5–24.7 Mya. Stem Mysticeti.

***Eomysticetus whitmorei* Sanders & Barnes 2002** Holotype from Bed 3 of the Chandler Bridge Formation, South Carolina (Sanders & Barnes, 2002b), dated by $^{87}$Sr/$^{86}$Sr correlation of the Chandler Bridge Formation (Weems et al., 2016; Boessenecker & Geisler, 2018), giving 23.5–24.7 Mya. Stem Mysticeti.

***Eophyseter damarcoi* Bisconti et al. 2025** Holotype (MGPT-PU 13864) from the Sabbie d'Asti Formation at Vigliano d'Asti, Piedmont (Bisconti et al., 2025b), placed in the early Piacenzian (Bisconti et al., 2025b), giving 3.00–3.20 Mya. Total group Physeteroidea: recovered within Physeteridae by Bisconti et al. (2025b), but on a holotype comprising only a partial vertebral column and forelimb, without cranium, ear bones, or teeth.

***Eoplatanista gresalensis* (Dal Piaz 1916)** Holotype from the upper Libano Sandstone of the Belluno quarries (Dal Piaz, 1916a), dated to the late Aquitanian–early Burdigalian (Nelson et al., 2025c), giving 19.0–22.0 Mya. Total group Odontoceti.

***Eoplatanista italica* Dal Piaz 1916** Holotype from the upper Libano Sandstone of the Belluno quarries (Dal Piaz, 1916b), dated to the late Aquitanian–early Burdigalian (Nelson et al., 2025c), giving 19.0–22.0 Mya. Total group Odontoceti.

***Eoplatanista miocaenus* (Pilleri 1989)** Holotype from the upper Libano Sandstone of the Belluno quarries (Pilleri, 1989), dated to the late Aquitanian–early Burdigalian (Nelson et al., 2025c), giving 19.0–22.0 Mya. Total group Odontoceti.

***Eoplatanista taurinensis* Pilleri 1988** Holotype from the lower–middle Miocene of Piedmont (Pilleri, 1988), giving 15.98–20.45 Mya. Total group Odontoceti.

***Eosqualodon langewieschei* Rothausen 1968** Holotype, a skull with associated vertebrae from the Doberg near Bünde, Germany (Rothausen, 1968; Nelson et al., 2025b), of Chattian (Late Oligocene) age, giving 23.04–27.3 Mya. Stem Odontoceti: a squalodontid, one of the archaic heterodont odontocetes falling outside the crown-group (Nelson et al., 2025b).

***Eosqualodon latirostris* (Capellini 1903)** Holotype (MGP 26237) from the Arenaria di Grumi dei Frati at Schio, Vicenza (Capellini, 1903), correlated to the late Aquitanian–early Burdigalian Libano Sandstone, the older Aquitanian assignment lacking supporting evidence (Nelson et al., 2025c), giving 15.97–23.04 Mya. Stem Odontoceti.

***Eschrichtioides gastaldii* (Strobel 1881)** Holotype from the Sabbie d'Asti at Cortandone, Asti (Strobel, 1881; Bisconti, 2008), a late Zanclean–early Piacenzian unit, giving 3.00–4.00 Mya. Total group Balaenopteroidea: recovered within Eschrichtiidae, sister to *Eschrichtius*, by Bisconti et al. (2024).

***Eschrichtius akishimaensis* Kimura et al. 2018** Holotype (GMNH-PV-3210) from the middle member of the Komiya Formation at Akishima, Tokyo (Kimura et al., 2018); its horizon is correlated with the Olduvai Subchron (C2n) by magnetostratigraphy (Kimura et al., 2018), giving 1.77–1.95 Mya. Total group Balaenopteroidea.

***Etruridelphis giulii* (Lawley 1876)** Holotype from the middle Pliocene of the Pisa hills (Poggetto dei Greppoli, Lorenzana), Tuscany (Lawley, 1876; Bianucci, 1996), giving 2.58–4.00 Mya. Crown Delphinidae: recovered within Delphininae by Bianucci (2013).

***Eubalaena ianitrix* Bisconti et al. 2017** Holotype (RBINS M.879a-f), a neurocranium from the Kruisschans Sand Member of the Lillo Formation, dated by dinoflagellate cysts and sequence stratigraphy to 2.8–3.2 Mya (Bisconti et al., 2017). Crown Balaenidae: recovered as the sister species of the extant *Eubalaena glacialis* within crown *Eubalaena* (Bisconti et al., 2017).

***Eubalaena shinshuensis* Kimura et al. 2007** Holotype (SFMCV-0024) from the Gonda Formation at Shinshushinmachi, Nagano (Kimura et al., 2007), its maximum set by a K–Ar age of 6.1 Mya on the Koso Tuff in the underlying Ogawa Formation (Kimura et al., 2007), giving 5.333–6.1 Mya. Crown Balaenidae: recovered as the earliest-diverging *Eubalaena* by Bisconti et al. (2017).

***Eucetotherium helmersenii* (Brandt 1871)** Holotype from the Sarmatian of the Eastern Paratethys (Brandt, 1871), giving 7.65–12.65 Mya (Palcu et al., 2021). Total group Mysticeti.

***Eudelphis mortezelensis* (Du Bus 1872)** Holotype (IRSNB M.523) from the Antwerpen Sands Member of the Berchem Formation at Vieux-Dieu, Antwerp (Du Bus, 1872; Lambert, 2008b), giving 12.8–15.97 Mya. Total group Physeteroidea: recovered at the base of Physeteroidea, outside the crown, by Peri et al. (2022).

***Eurhinodelphis cocheteuxi* du Bus 1867** Holotype (IRSNB 3252-M.294) from the Antwerpen Sands Member of the Berchem Formation at Vieux-Dieu (Oude-God), Antwerp, Belgium (Du Bus, 1867; Lambert, 2005a), of late Early–Middle Miocene age, giving 11.63–15.98 Mya. Total group Odontoceti: a long-snouted eurhinodelphinid, an archaic odontocete grade retained at the total group (Lambert, 2005a).

***Eurhinodelphis longirostris* du Bus 1872** Holotype (IRSNB 3249-M.342) from the Miocene of the Antwerp area, Belgium (Du Bus, 1872; Lambert, 2005a), placed in the late Early–Middle Miocene of the southern North Sea Basin, giving 7.6–15.97 Mya. Total group Odontoceti: see *Eurhinodelphis cocheteuxi*.

***Ferecetotherium kelloggi* Mchedlidze 1970** Holotype from the late Oligocene Maikop Group of Azerbaijan (Mchedlidze, 1970), giving 23.03–28.1 Mya. Total group Physeteroidea.

***Fordycetus taitonga* Tanaka et al. 2026** Holotype (OU 22670, a partial rostrum and cranium with a cervical vertebra) collected as a float at the mouth of Awamoa Creek, coastal North Otago, New Zealand, from the Mount Harris Formation (Tanaka et al., 2026), of Otaian–Altonian (latest Aquitanian–Burdigalian) age by Sr-isotope and planktic-foraminiferal dating, giving 15.9–21.7 Mya (Tanaka et al., 2026). Crown Odontoceti.

***Fragilicetus velponi* Bisconti & Bosselaers 2016** Holotype (NMR 999100007727), found in situ in the Kattendijk Sand Member of the Kattendijk Formation at the Deurganckdok, about 3 m above the basal gravel (Bisconti & Bosselaers, 2016), early Zanclean (Louwye et al., 2004), giving 4.4–5.0 Mya. Total group Balaenopteroidea: recovered as a transitional taxon at the origin of Balaenopteridae by Bisconti and Bosselaers (2016).

***Fucaia buelli* Marx et al. 2015** Holotype (UWBM 84024) from the Makah Formation, Clallam County, Washington, dated to the early Oligocene by magnetochron C12r and Refugian foraminifera (Marx et al., 2015), giving 31–33.2 Mya. Stem Mysticeti: an aetiocetid (Marx et al., 2015), one of the basal toothed mysticete clades outside Chaeomysticeti (Marx & Fordyce, 2015).

***Fucaia goedertorum* (Barnes et al. 1995)** Holotype from the Twin River Quarry, Pysht Formation, Clallam County, Washington (Barnes et al., 1995), dated to the late Oligocene by the Twin River Group magnetostratigraphy of Prothero et al. (2001), giving 26.5–28.1 Mya. Stem Mysticeti: see *Fucaia buelli*.

***Fucaia humilis* Tsai et al. 2024** Holotype from the lower part of the Lincoln Creek Formation of Washington, dated to the latest Eocene near the Eocene–Oligocene transition (Tsai et al., 2024a), giving 33.9–37.71 Mya. Stem Mysticeti: see *Fucaia buelli*.

***Furcacetus flexirostrum* Bianucci et al. 2020** Holotype (MUSM 487) from the Chilcatay Formation (Bianucci et al., 2020); the holotype horizon is unknown, so we assigned the age bracket for the entire formation (see *Chilcacetus cavirhinus*), giving 18.0–19.3 Mya. Crown Odontoceti.

***Gandakasia potens* Dehm & Oettingen-Spielberg 1958** Holotype (IPHG 1956 II 4), teeth, from the Kuldana Formation at Ganda Kas, northern Pakistan (Dehm & Oettingen-Spielberg, 1958), of middle Eocene age, giving 46–49 Mya. Stem Cetacea.

***Gaviacetus razai* Gingerich et al. 1995** Holotype (GSP-UM 3095), a partial skull, from the Domanda Formation at Basti Ahmed, Sulaiman Range, Punjab, Pakistan (Gingerich et al., 1995), of middle Eocene (Lutetian) age, giving 42.5–46 Mya. Stem Cetacea.

***Georgiacetus vogtlensis* Hulbert et al. 1998** Holotype, a partial skeleton, from the Blue Bluff unit (formerly part of the McBean Formation) of the Upper Coastal Plain of eastern Georgia, USA (Hulbert et al., 1998), dated by biostratigraphically diagnostic mollusks and calcareous nannoplankton to the latest Lutetian–earliest Bartonian, giving 40.0–41.0 Mya (Hulbert et al., 1998). Stem Cetacea.

***Glaucobalaena inopinata* Bisconti et al. 2024** Holotype (MGPT-PU 19512) from the Sabbie d'Asti Formation, Piedmont (Bisconti et al., 2024), a late Zanclean–early Piacenzian unit, giving 3.00–4.00 Mya. Total group Balaenopteroidea: recovered within Eschrichtiidae, in a clade with *Gricetoides*, by Bisconti et al. (2024).

***Globicephala etruriae* Pilleri 1987** Holotype (IGF 1675V) from the Pliocene of La Rocca, near Volterra (Pisa), Tuscany (Pilleri, 1987; Bianucci, 1996), giving 2.58–5.33 Mya. Total group Delphinidae.

***Goedertius oregonensis* Kimura & Barnes 2016** Holotype from the Nye Mudstone, Lincoln County, Oregon, of early Miocene (Aquitanian) age (Kimura & Barnes, 2016), giving 20.45–23.04 Mya. Crown Odontoceti: an allodelphinid (Kimura & Barnes, 2016), a clade nested within crown Odontoceti (Boersma & Pyenson, 2016).

***Goniodelphis hudsoni* Allen 1941** Holotype (MCZ 17881) from the Palmetto Fauna of the upper Bone Valley Member of the Peace River Formation, Polk County, Florida (Allen, 1941), of latest Miocene–earliest Pliocene (Hemphillian) age (Morgan, 1994), giving 4.75–5.7 Mya. Total group Inioidea.

***Graamocetus longicollis* Hoch 2004** Holotype from the Gram Clay of the Gram Formation at Gram Brick Works, South Jutland, Denmark (Hoch, 2004), an in-situ Tortonian assemblage dated by magnetostratigraphy and dinocysts (Rasmussen et al., 2010), giving 7.2–9.9 Mya. Total group Odontoceti.

***Gricetoides aurorae* Whitmore & Kaltenbach 2008** Holotype from the Yorktown Formation at the Lee Creek Mine, North Carolina (Whitmore & Kaltenbach, 2008), the Lee Creek Yorktown given a corrected early Pliocene age (Boessenecker et al., 2019), giving 3.92–4.9 Mya. Total group Balaenopteroidea.

***Grimadelphis spectorum* Godfrey & Lambert 2023** Holotype from the Popes Creek Sand of the Fairhaven Member of the Calvert Formation, Maryland, dated to the late Early Miocene (Burdigalian) (Godfrey & Lambert, 2023), giving 19.0–19.5 Mya. Crown Odontoceti.

***Haborodelphis japonicus* Ichishima et al. 2019** Holotype from the Embetsu Formation, Haboro, Hokkaido (Ichishima et al., 2019), dated to the early Pliocene, giving 3.6–5.5 Mya. Total group

Monodontidae.

***Haborophocoena minutus* Ichishima & Kimura 2009** Holotype from the Embetsu Formation, Hokkaido (Ichishima & Kimura, 2009), dated to the early Pliocene, giving 3.6–5.5 Mya. Total group Phocoenidae: recovered outside crown Phocoenidae by Colpaert et al. (2015).

***Haborophocoena toyoshimai* Ichishima & Kimura 2005** Holotype (HMNH 110-1) from the Embetsu Formation, Hokkaido (Ichishima & Kimura, 2005), dated to the early Pliocene, giving 3.6–5.5 Mya. Total group Phocoenidae: see *Haborophocoena minutus*.

***Hadrodelphis calvertense* Kellogg 1966** Holotype (USNM V 23408), a mandible with teeth from the Calvert Formation at Governors Run, Calvert County, Maryland, its horizon within the formation uncertain (Kellogg, 1966); the interval was set based on the lower jaw USNM PAL 182938 from Shattuck Zone 13 (Godfrey & Lambert, 2023), giving 14.0–14.9 Mya. Total group Delphinida.

***Halicetus ignotus* Kellogg 1969** Holotype (USNM 23636), a skull with ear bones and much of the postcranium collected in situ from the shell layer of Shattuck Zone 19, Boston Cliffs Member of the Choptank Formation, Calvert County, Maryland (Kellogg, 1969), dated to the late Middle Miocene (Godfrey & Lambert, 2023), giving 11.5–12.5 Mya. Total group Mysticeti.

***Hemisyntrachelus cortesii* (Fischer 1829)** Holotype (MC CF2) from the Pliocene of Colle della Terrazza, Piacenza (Fischer, 1829; Bianucci, 1996), giving 2.58–5.33 Mya. Crown Delphinidae: recovered within Orcininae, in a clade with the extant *Orcinus* and *Pseudorca*, by Bianucci (2005).

***Hemisyntrachelus oligodon* (Pilleri & Siber 1989)** Holotype (SMNK 3841), a skull, from the Sud-Sacaco West horizon of the Pisco Formation, southern Peru (Pilleri & Siber, 1989); the Messinian Sud-Sacaco interval is dated ∼6.6–5.9 Mya (Ochoa et al., 2021). Crown Delphinidae: a fossil relative of the extant bottlenose dolphins (Post & Bosselaers, 2005).

***Herbeinodelphis nancei* Godfrey & Lambert 2023** Holotype (CMM-V-8343), a partial skull with periotic, bulla, mandible, and vertebrae from Shattuck Zone 13 of the Plum Point Member of the Calvert Formation at Warriors' Rest Sanctuary, Calvert County, Maryland, dated to the early Serravallian (Godfrey & Lambert, 2023), giving 14.0–14.9 Mya. Total group Delphinida.

***Herentalia nigra* Bisconti 2014** Holotype (ZMA 5069) from the Deurne Sand Member of the Diest Formation (Bisconti, 2014), dated to the late Tortonian by the dinocyst chronology of the Deurne

Member (Goolaerts et al., 2020), giving 7.6–9.5 Mya. Crown Mysticeti: a cetotheriid (Cetotheriidae *sensu stricto*) sharing a within-cetotheriid morphotype group with *Metopocetus* and *Piscobalaena* (Bisconti, 2014; Marx et al., 2017, 2019), a family nested inside crown Mysticeti (Marx & Fordyce, 2015).

***Herpetocetus bramblei* Whitmore & Barnes 2008** Holotype (UCMP 82465) from the "Crab Marker" bed of the Purisima Formation (Whitmore & Barnes, 2008), placed near the Miocene–Pliocene boundary with a validly referred specimen extending the range lower in the formation (Boessenecker, 2013), giving 4.5–6.4 Mya. Crown Mysticeti: see *Herpetocetus scaldiensis*.

***Herpetocetus morrowi* El Adli et al. 2014** Holotype from the San Diego Formation of California (El Adli et al., 2014), dated to the Pliocene–early Pleistocene range of the formation (Boessenecker, 2013), giving 1.8–4.2 Mya. Crown Mysticeti: see *Herpetocetus scaldiensis*.

***Herpetocetus scaldiensis* Van Beneden 1872** Lectotype (IRScNB CtM 379a-b) from Stuyvenberg, Antwerp (Van Beneden, 1872; Steeman, 2010), whose material is assigned to the upper Zanclean–Piacenzian (Scaldisian, ca 2.5–4 Mya) (Steeman, 2010). Crown Mysticeti: a herpetocetine cetotheriid (Steeman, 2010), within the cetotheriid clade that lies inside crown Mysticeti (Marx & Fordyce, 2015).

***Herpetocetus transatlanticus* Whitmore & Barnes 2008** Holotype (USNM 182962), a partial braincase from the Yorktown Formation at the Lee Creek Mine, North Carolina (Whitmore & Barnes, 2008), the Lee Creek Yorktown given a corrected early Pliocene age (Boessenecker et al., 2019), giving 3.92–4.9 Mya. Crown Mysticeti: see *Herpetocetus scaldiensis*.

***Heterocetus major* De Stefano 1900** Holotype from the Pliocene of Italy (De Stefano, 1900), giving 2.58–5.33 Mya. Total group Mysticeti; also see *Heterocetus scriptus*.

***Heterocetus scriptus* Vanden Broeck 1874** Based on 19th-century material from the "sables moyens d'Anvers" fortification excavations, of uncertain stratigraphic level spanning the Antwerpen Sand and Deurne Sand Members (Vanden Broeck, 1874; Steeman, 2010), giving a wide 7.6–15.97 Mya bracket. Total group Mysticeti: the genus *Heterocetus* is a *nomen dubium* (Steeman, 2010), but the species itself is valid.

***Heterodelphis croatica* (Gorjanović-Kramberger 1892)** Holotype from the Sarmatian of Podsused, Medvednica (Central Paratethys) (Gorjanović-Kramberger, 1892), dated to the Central Paratethys

Sarmatian stage (Hohenegger et al., 2014), giving 11.6–12.83 Mya. Total group Odontoceti.

***Heterodelphis klinderi* Brandt 1873** Holotype from the Sarmatian of the Paratethys (Brandt, 1873b), giving 7.65–9.6 Mya (Palcu et al., 2021). Total group Odontoceti.

***Heterodelphis leiodontus* Papp 1905** Holotype from the middle Miocene (Badenian) of Sopron County, Hungary (Papp, 1905), giving 12.83–16.3 Mya (Hohenegger et al., 2014). Total group Odontoceti.

***Hibacetus hirosei* Otsuka & Ota 2008** Holotype from the Korematsu Formation at Nakahara, Bihoku Group, Hiroshima (Otsuka & Ota, 2008), referred to the middle Miocene upper Bihoku Group, whose overlying Itahashi Formation falls in calcareous nannofossil Zone NN4 (Tsai, 2017), giving 15.6–16.9 Mya. Total group Mysticeti.

***Himalayacetus subathuensis* Bajpai & Gingerich 1998** Holotype (RUSB 2003) from the Subathu Formation at Kuthar Nala, India (Bajpai & Gingerich, 1998), of early Eocene (Ypresian) age, giving 45.4–52.9 Mya. Stem Cetacea.

***Horopeta umarere* Tsai & Fordyce 2015** Holotype (OU 21982) from the middle Kokoamu Greensand, Hakataramea Valley (Tsai & Fordyce, 2015), of Duntroonian age (Robinson et al., 2024), giving 25.2–27.3 Mya. Total group Mysticeti: originally recovered as a stem mysticete but subsequently nested within crown Mysticeti as a sister taxon of the cetotheriids under several analyses, leaving its within-clade position unresolved (Tsai, 2023).

***Huaridelphis raimondii* Lambert et al. 2014** Holotype (MUSM 1396) from the Ct1 strata of the Chilcatay Formation at Ullujaya (Lambert et al., 2014), dated to the early Miocene by the Chilcatay tephra chronology (see *Chilcacetus cavirhinus*) (Bianucci et al., 2020), giving 18.0–19.3 Mya. Crown Odontoceti.

***Ichthyolestes pinfoldi* Dehm & Oettingen-Spielberg 1958** Holotype (IPHG 1956 II 7), teeth, from the Kuldana Formation at Ganda Kas, northern Pakistan (Dehm & Oettingen-Spielberg, 1958), of middle Eocene age, giving 46–49 Mya. Stem Cetacea.

***Idiocetus guicciardinii* Capellini 1876** Holotype from the Pliocene of Montopoli, Lower Valdarno, Tuscany (Capellini, 1876), giving 1.80–3.55 Mya. Total group Balaenidae.

***Idiophyseter merriami* Kellogg 1925** Holotype from near Templeton, San Luis Obispo County, California, of middle Miocene age (Kellogg, 1925a), giving 13.82–15.98 Mya. Crown Physeteroidea: a physeterid, grouped with the extant *Physeter* (Peri et al., 2022; Lambert et al., 2023).

***Idiorophus bolzanensis* (Dal Piaz 1916)** Holotype from the upper Libano Sandstone of the Belluno quarries (Dal Piaz, 1916c), dated to the late Aquitanian–early Burdigalian (Nelson et al., 2025c), giving 19.0–22.0 Mya. Total group Physeteroidea.

***Idiorophus patagonicus* (Lydekker 1894)** Holotype, an incomplete skull, from the lower Gaiman Formation at Bryn Gwyn, Chubut, Argentina (Lydekker, 1894; Paolucci et al., 2025); the Gaiman overlies the Colhuehuapian Trelew Member and does not predate late Aquitanian (Buono et al., 2017), giving 15.98–20.45 Mya. Total group Physeteroidea: placed within Physeteridae (with *Physeter*) by Peri et al. (2022) and Lambert et al. (2023), but recovered as the most basal physeterid or a stem physeteroid by Paolucci et al. (2025).

***Imerocetus karaganicus* Mchedlidze 1964** Holotype from the Karaganian of Georgia (Mchedlidze, 1964), giving 13.4–13.8 Mya. Total group Mysticeti.

***Imerodelphis thabagarii* Mchedlidze 1959** Holotype from the Sarmatian near Zestafoni, Georgia (Mchedlidze, 1959), giving 11.63–13.82 Mya (Palcu et al., 2021). Total group Odontoceti.

***Incacetus broggii* Colbert 1944** Holotype (AMNH 32656), originally described by Colbert (1944), from the “Santa Rosa vertebrate level” of the P0 allomember of the Pisco Formation, southern Peru (Lambert et al., 2026); the P0 allomember is dated 14.8–12.4 Mya (Langhian–Serravallian) (Lambert et al., 2026). Total group Odontoceti.

***Incakujira anillodefuego* Marx & Kohno 2016** Holotype (GNHM Fs-098-12), a cranium, from the Aguada de Lomas level of the Pisco Formation, southern Peru (Marx & Kohno, 2016), whose lowermost succession is bracketed to 8.85–7.93 Mya by K–Ar dating of two tuffs (Ochoa et al., 2021). Total group Balaenopteroidea: placed within crown Balaenopteridae in the morphology-only analysis but as sister to *Megaptera* versus a monophyletic *Balaenoptera* in the total-evidence analysis of Marx and Kohno (2016).

***Incakujira fordycei* Kimura & Hasegawa 2024** Holotype (GMNH-PV-159), an almost complete skeleton with a preserved baleen rack, from the Aguada de Lomas level of the Pisco Formation, southern

Peru (Kimura & Hasegawa, 2024a); K–Ar ages on two tuffs give 8.85–7.93 Mya for the lowermost Aguada de Lomas succession (Ochoa et al., 2021; Kimura & Hasegawa, 2024a), giving 7.9–8.8 Mya. Total group Balaenopteroidea: see *Incakujira anillodefuego*.

***Indocetus ramani* Sahni & Mishra 1975** Holotype (LUVP 11034), a skull, from the Harudi Formation at Rato Nala near Harudi, Kutch, western India (Sahni & Mishra, 1975), of late middle Eocene (Bartonian) age, giving 37.71–42 Mya. Stem Cetacea.

***Inermorostrum xenops* Boessenecker et al. 2017** Holotype (CCNHM 171), a partial skull from the Ashley Formation near Charleston, South Carolina (Boessenecker et al., 2017b), dated by $^{87}$Sr/$^{86}$Sr correlation of the Ashley Formation (Weems et al., 2016; Boessenecker & Geisler, 2018), giving 26.57–29.0 Mya. Stem Odontoceti: placed within Xenorophidae, outside crown Odontoceti, by Boessenecker et al. (2017b).

***Iniopsis caucasica* Lydekker 1892** Holotype from the Maikop Group at Perekishkyul, on the Sumgait River (Azerbaijan) (Lydekker, 1892), reported as Chattian (late Oligocene) on the Maikop succession (Popov et al., 2008), giving 23.04–27.3 Mya. Total group Odontoceti.

***Inticetus vertizi* Lambert et al. 2018** Holotype (MUSM 1980) from the Chilcatay Formation at Roca Negra (Lambert et al., 2018b), dated by the describers to the late early Miocene (Burdigalian, 18.0–18.8 Mya) (Lambert et al., 2018b). Total group Odontoceti: type of the family Inticetidae, recovered either as a stem odontocete or an early-branching platanistoid (Lambert et al., 2018b).

***Isanacetus laticephalus* Kimura & Ozawa 2002** Holotype from the Hiramatsu Formation, Awa Group, Mie (Kimura & Ozawa, 2002), the succession constrained by its foraminiferal zonation (Zones N.6–N.8) and the age of the overlying Makino Formation (Fujiwara et al., 2005), giving 16.4–18.5 Mya. Total group Mysticeti.

***Ischyrorhynchus vanbenedeni* Ameghino 1891** Holotype from the upper Miocene Ituzaingó Formation (‘Conglomerado osífero’) near Paraná, Entre Ríos, Argentina (Ameghino, 1891), giving 6.67–9.47 Mya. Total group Inioidea.

***Isoninia borealis* Godfrey et al. 2021** Holotype (CMM-V-4061), recovered without encasing sediment and inferred to derive from the Cobham Bay Member of the Eastover Formation, North Carolina. Thus, we assigned a conservative age bracket from the upper Tortonian to the lower Piacenzian (Godfrey et al., 2021), giving 2.58–11.63 Mya. Total group Inioidea: assigned to Iniidae with reservation by Godfrey et al. (2021).

***Isthminia panamensis* Pyenson et al. 2015** Holotype (USNM 546125) from the Piña facies of the Chagres Formation at STRI locality 650009, Piña, Caribbean coast of Panama (Pyenson et al., 2015), constrained by planktonic foraminifera and superposition above the Toro Member to 5.8–6.1 Mya (Pyenson et al., 2015). Crown Inioidea: recovered as the sister taxon of the extant *Inia* by Pyenson et al. (2015).

***Janjucetus dullardi* Duncan et al. 2025** Holotype (NMV P256471) from the Jan Juc Marl at Half Moon Bay (Bells Headland), Victoria (Duncan et al., 2025), with a horizon ∼3 m above the fossil-producing layer $^{87}$Sr/$^{86}$Sr-dated to 24.67 Mya (Duncan et al., 2025), giving 24.67–26.05 Mya. Stem Mysticeti.

***Janjucetus hunderi* Fitzgerald 2006** Holotype (NMV P216929) from the Jan Juc Marl of the Torquay area, Victoria (Fitzgerald, 2006), of late Chattian age, confined to 24.2–26.05 Mya by the strontium chronology of the fossiliferous Jan Juc Marl (Duncan et al., 2025). Stem Mysticeti.

***Jobancetus pacificus* Kimura et al. 2022** Holotype (I-418267) from the lower part of the Minamishirado Formation, Shirado Group, Fukushima (Kimura et al., 2022), dated by a fission-track age of 15.9±0.7 Mya on a tuff in the lower Minamishirado Formation and by diatom correlation to Zone NPD 3A (Kimura et al., 2022), giving 15.2–16.6 Mya. Total group Mysticeti: recovered as a stem plicogulan under equal weighting but within Balaenopteridae + Eschrichtiidae under implied weighting by Kimura et al. (2022).

***Joumocetus shimizui* Kimura & Hasegawa 2010** Holotype from the Haraichi Formation at Annaka, Gunma (Kimura & Hasegawa, 2010), of earliest late Miocene (Tortonian) age, giving 11.25–11.29 Mya. Total group Mysticeti.

***Kaaucetus thesaurus* Hernández-Cisneros 2022** Holotype from the San Juan Member of the El Cien Formation (Hernández-Cisneros, 2022), dated to the late Oligocene by a U–Pb zircon age on a tuff in the San Juan Member (Rochín-Bañaga et al., 2024), giving 27.79–28.11 Mya. Total group Mysticeti.

***Kalakocetus aurorae* Waqas et al. 2026** Holotype (GU/RJ/07) from the upper Subathu Formation at Kalakote, Rajouri District, Jammu and Kashmir, India (Waqas et al., 2026), of middle Eocene (Lutetian) age, giving 45.4–48.7 Mya. Stem Cetacea.

***Kampholophos serrulus* Rensberger 1969** Holotype from the Claremont Shale of the Monterey Formation, California (Rensberger, 1969), of middle Miocene age, giving 11.63–13.82 Mya. Total group Delphinida.

***Kekenodon onamata* Hector 1881** Holotype (NMNZ Ma 306) from the Kekenodon Beds at Wharekuri, a lateral equivalent of the Kokoamu Greensand (Hector, 1881; Corrie & Fordyce, 2022), of Duntroonian age (Corrie & Fordyce, 2022), giving 25.2–27.3 Mya. Stem Neoceti: the latest-surviving archaeocete, recovered as the sister taxon to Neoceti (Corrie & Fordyce, 2022).

***Kennedycetus pericorum* Solís-Añorve et al. 2021** Holotype from the Trinidad Formation of the San José del Cabo Basin, Baja California Sur, Mexico, dated to the late Miocene by calcareous nannofossil zone NN11 (Solís-Añorve et al., 2021), giving 5.33–11.63 Mya. Total group Balaenopteroidea.

***Kentriodon diusinus* Salinas-Márquez et al. 2014** Holotype (FCMM 2387, a skull with mandibles and partial postcranium) from locality FCMM 103 (capa G), Los Indios Member of the Rosarito Beach Formation, Baja California (Salinas-Márquez et al., 2014), bracketed by an underlying basalt at 16.1±2.1 Mya and the overlying Costa Azul Member at 14.3±2.6 Mya with Temblor-stage molluscs (Salinas-Márquez et al., 2014), giving 14.0–16.0 Mya. Total group Delphinida: a kentriodontid, an archaic delphinidan grade lying outside the crown delphinoid families (Guo & Kohno, 2021).

***Kentriodon fuchsii* (Brandt 1873)** Holotype from the middle Miocene of the Central Paratethys (Brandt, 1873b), giving 11.6–12.83 Mya (Hohenegger et al., 2014). Total group Delphinida: see *Kentriodon diusinus*.

***Kentriodon hobetsu* Ichishima 1995** Holotype from the Takinoue Formation, Hobetsu, Hokkaido, dated to the middle Miocene (Ichishima, 1995), giving 11.63–16.7 Mya. Total group Delphinida: see *Kentriodon diusinus*.

***Kentriodon hoepfneri* Kazár & Hampe 2014** Holotype from the “Oberer Glimmerton” mica-clay horizon at Groß Pampau, Schleswig-Holstein, Germany (Kazár & Hampe, 2014), dated to the late

Langenfeldian (*Bolboforma fragori/subfragoris* Zone) at ca. 10.6–11.8 Mya (Kazár & Hampe, 2014). Total group Delphinida: see *Kentriodon diusinus*.

***Kentriodon nakajimai* Kimura & Hasegawa 2019** Holotype from the Haraichi Formation on the Usui River, Annaka, Gunma (Kimura & Hasegawa, 2019), of earliest late Miocene (Tortonian) age, giving 11.25–11.29 Mya. Total group Delphinida: see *Kentriodon diusinus*.

***Kentriodon obscurus* (Kellogg 1931)** Holotype from the Sharktooth Hill bonebed of the Round Mountain Silt (Kellogg, 1931), correlated with magnetochron C5Br by the magnetostratigraphy of Pyenson et al. (2009), giving 15.2–16.0 Mya. Total group Delphinida: see *Kentriodon diusinus*.

***Kentriodon pernix* Kellogg 1927** Holotype (USNM V 8060), a nearly complete articulated skeleton from Shattuck Zone 5 of the Plum Point Member of the Calvert Formation (Kellogg, 1927a), placed in the late Early Miocene (Burdigalian) by the Calvert Cliffs zonal framework of Godfrey and Lambert (2023), giving 16.0–20.6 Mya. Total group Delphinida: see *Kentriodon diusinus*.

***Kentriodon schneideri* Whitmore & Kaltenbach 2008** Holotype (USNM 323772, a skull) from the Lee Creek Mine, Pungo River Formation, North Carolina (Whitmore & Kaltenbach, 2008); the Pungo River Formation is an early–middle Miocene Calvert-equivalent unit of poorly resolved age (Godfrey & Lambert, 2023), giving 13.82–20.44 Mya. Total group Delphinida: see *Kentriodon diusinus*.

***Kentriodon sugawarai* Guo & Kohno 2021** Holotype (NMHF 999) from the Shikonai Siltstone Member of the Kadonosawa Formation at the Mabechi River, Ninohe City, Iwate (Guo & Kohno, 2021), dated to diatom Zone NPD 3B (the *Denticulopsis praelauta* Zone) (Guo & Kohno, 2021), giving 15.9–16.3 Mya. Total group Delphinida: see *Kentriodon diusinus*.

***Kharodacetus sahnii* (Bajpai & Thewissen 1998)** Holotype (VPL 1021) from a shallow subtidal mudstone of the Harudi Formation at Harudi, Kachchh, western India (Bajpai & Thewissen, 1998, 2014), of middle Eocene (Bartonian) age, giving 37.71–42.0 Mya. Stem Cetacea.

***Kogia danomurai* Benites-Palomino et al. 2021** Holotype (MUSM; catalog number unverified) from the Sacaco (SAO) horizon of the Pisco Formation (Benites-Palomino et al., 2021), dated to the Messinian by U–Pb zircon geochronology (Ochoa et al., 2021), giving 5.7–5.9 Mya. Total group Kogiidae: described as the most basal representative of the extant genus *Kogia* (Benites-Palomino et al., 2021) and predating the crown-*Kogia* node, which post-dates the early Pliocene (Vélez-Juarbe et al., 2015), but not directly recovered outside the crown.

***Kogia pusilla* (Pilleri 1987)** Holotype (IGF 1540V), an incomplete skull, from the Monte Voltraio cetacean beds near Volterra (Tuscany, Italy) (Pilleri, 1987), originally the type of *Hyperoodon pusillus* and transferred to *Kogia* by Bianucci (1997) (Collareta et al., 2017), in the middle Pliocene *Globorotalia aemiliana–Discoaster tamalis* zones, giving 2.58–3.6 Mya. Total group Kogiidae: recovered as the sister of the extant *Kogia* pair, outside the crown genus, by Vélez-Juarbe et al. (2015) and Collareta et al. (2017).

***Kogiopsis floridana* Kellogg 1929** Holotype from the lower Bone Valley Member of the Peace River Formation, central Florida (Kellogg, 1929b), referred to the late Miocene (Clarendonian) lower Peace River sequence (Morgan, 1994; Missimer, 1999), giving 8.5–11.0 Mya. Total group Physeteroidea.

***Koristocetus pescei* Collareta et al. 2017** Holotype (MUSM 888), a partial cranium, from the Aguada de Lomas level of the Pisco Formation, southern Peru (Collareta et al., 2017), whose lowermost succession is bracketed to 8.85–7.93 Mya by K–Ar dating of two tuffs (Ochoa et al., 2021), giving 7.9–8.8 Mya. Total group Kogiidae: the earliest-branching member of Kogiinae, a kogiid lying rootward of the extant *Kogia* crown (Vélez-Juarbe et al., 2015; Collareta et al., 2017).

***Kurdalagonus maicopicus* (Spasskii 1951)** Lectotype (PIN 764/1) from the Middle Sarmatian Blinovskaya Formation on the Belaya River at Maykop, Adygea, Russia (Spasskii, 1951; Tarasenko & Lopatin, 2012a; Tarasenko, 2024), giving 7.65–11.8 Mya (Palcu et al., 2021). Total group Mysticeti.

***Kurdalagonus mchedlidzei* Tarasenko & Lopatin 2012** Holotype from the middle–upper Sarmatian of Adygea, northern Caucasus (Tarasenko & Lopatin, 2012a), giving 7.65–11.8 Mya (Palcu et al., 2021). Total group Mysticeti.

***Kutchicetus minimus* Bajpai & Thewissen 2000** Holotype (RUSB 2647) from a shallow subtidal limestone of the Harudi Formation at Godhatad, Kachchh, western India (Bajpai & Thewissen, 2000), of middle Eocene (Bartonian) age, giving 37.71–42.0 Mya. Stem Cetacea.

***Kwanzacetus khoisani* Lambert et al. 2018** Holotype (CZA 1-16, a partial skeleton) from upper shoreface sands about 10 m above the basal unconformity in the Kwanza Basin, south of the Cuanza estuary, Angola (Lambert et al., 2018a), of late Miocene (Tortonian–Messinian) age, giving 5.33–11.63

Mya. Crown Inioidea: an iniid recovered as the closest fossil relative of the extant *Inia geoffrensis* (Lambert et al., 2018a).

***Lagenorhynchus harmatuki* Whitmore & Kaltenbach 2008** Holotype from the Yorktown Formation at the Lee Creek Mine, North Carolina (Whitmore & Kaltenbach, 2008), with an early Pliocene age (Boessenecker et al., 2019), giving 3.92–4.9 Mya. Total group Delphinidae.

***Lamprolithax annectens* Kellogg 1931** Holotype from the Sharktooth Hill bonebed of the Round Mountain Silt (Kellogg, 1931), correlated with magnetochron C5Br by the magnetostratigraphy of Pyenson et al. (2009), giving 15.2–16.0 Mya. Total group Odontoceti.

***Lamprolithax simulans* Kellogg 1931** Holotype from the Sharktooth Hill bonebed of the Round Mountain Silt (Kellogg, 1931), correlated with magnetochron C5Br by the magnetostratigraphy of Pyenson et al. (2009), giving 15.2–16.0 Mya. Total group Odontoceti.

***Leptodelphis stavropolitanus* Kirpichnikov 1954** Holotype from the middle Sarmatian (Bessarabian) of Stavropol, Ciscaucasia (Russia), in the Eastern Paratethys (Kirpichnikov, 1954), dated to 9.6–12.1 Mya by the Eastern Paratethys stage framework (Palcu et al., 2017, 2021). Total group Odontoceti.

***Liolithax kernensis* Kellogg 1931** Holotype (CAS 4370) from the Sharktooth Hill bonebed of the Round Mountain Silt (Kellogg, 1931), correlated with magnetochron C5Br by the magnetostratigraphy of Pyenson et al. (2009), giving 15.2–16.0 Mya. Total group Delphinida.

***Lissodelphis fockii* (Brandt 1873)** Holotype from the middle Sarmatian (Bessarabian) of Stavropol, Ciscaucasia (Russia) (Brandt, 1873b), dated to 9.6–12.1 Mya by the Eastern Paratethys stage framework (Palcu et al., 2017, 2021). Total group Odontoceti.

***Lissodelphis nordmanni* (Brandt 1873)** Holotype from the middle Sarmatian (Bessarabian) of Chişinău (Moldova) (Brandt, 1873b), dated to 9.6–12.1 Mya by the Eastern Paratethys stage framework (Palcu et al., 2017, 2021). Total group Odontoceti.

***Livyatan melvillei* Lambert et al. 2010** Holotype (MUSM 1676) from Cerro Colorado, allomember P1 of the Pisco Formation (Lambert et al., 2010), dated to the Tortonian by the P1 allomember age (Di Celma et al., 2017), giving 8.5–9.5 Mya. Total group Physeteroidea: recovered among the macroraptorial stem sperm whales in most studies (Lambert et al., 2010; Collareta et al., 2017), but sister to Kogiidae inside the crown in the two most recent analyses (Peri et al., 2022; Lambert et al., 2023).

***Llanocetus denticrenatus* Mitchell 1989** Holotype (USNM 183022) from the La Meseta Formation, Seymour Island, Antarctic Peninsula (Mitchell, 1989), of late Eocene (Priabonian) age, giving 33.9–35.07 Mya. Stem Mysticeti: a large toothed baleen whale placed in Llanocetidae as an early-diverging mysticete (Fordyce & Marx, 2018).

***Lomacetus ginsburgi* Muizon 1988** Holotype (MNHN PPI 104) from Aguada de Lomas in the Pisco Formation, Sacaco Basin, southern Peru (de Muizon, 1988), dated by the Aguada de Lomas vertebrate level of Ochoa et al. (2021), giving 7.9–8.8 Mya. Total group Phocoenidae.

***Lophocetus calvertensis* (Harlan 1842)** Holotype (USNM V 16314), an incomplete skull with both periotics and cervical vertebrae, from the St. Marys Formation at Cove Point, Calvert County, Maryland (Harlan, 1842; Godfrey & Lambert, 2023), from the Little Cove Point Member dated to the Tortonian by the Calvert Cliffs dinocyst stratigraphy (Kidwell et al., 2015; Weems et al., 2017), giving 8.7–11.9 Mya. Total group Delphinida.

***Lophocetus repenningi* Barnes 1978** Holotype from the lower Santa Margarita Sandstone (Stanford Locality) of the Santa Cruz Mountains (Barnes, 1978), of late Miocene age, giving 10.0–12.0 Mya. Total group Delphinida: placed in the family Lophocetidae within a kentriodontoid clade recovered as sister to Delphinoidea by Guo and Kohno (2023).

***Loxolithax sinuosa* Kellogg 1931** Holotype from the Sharktooth Hill bonebed of the Round Mountain Silt (Kellogg, 1931), correlated with magnetochron C5Br by the magnetostratigraphy of Pyenson et al. (2009), giving 15.2–16.0 Mya. Total group Odontoceti.

***Macrodelphinus kelloggi* Wilson 1935** Holotype from the Pyramid Hill Sand Member of the Jewett Sand, Kern County, California (Wilson, 1935), dated by the San Joaquin Basin chronostratigraphy of Scheirer and Magoon (2008), giving 24–25 Mya. Total group Odontoceti.

***Macrodelphinus valenciennesii* (Brandt 1873)** Holotype, a rostrum fragment (Brandt, 1873b), from Burdigalian marine molasse near Sort (Landes, Aquitaine, France), giving 15.98–20.45 Mya (Aguirre-Fernández et al., 2022). Total group Odontoceti.

***Macrokentriodon morani* Dawson 1996** Holotype (CMM-V-15), a skull with teeth and periotic from the contact between the Drumcliff Member (Shattuck Zone 17) and the St. Leonard Member (Shattuck Zone 18) of the Choptank Formation at Moran's Beach, Calvert County, Maryland (Dawson, 1996), dated to the late Middle Miocene (Serravallian) (Godfrey & Lambert, 2023), giving 12.0–13.0 Mya. Total group Delphinida.

***Macrosqualodelphis ukupachai* Bianucci et al. 2018** Holotype (MUSM 2545) from the Chilcatay Formation (Bianucci et al., 2018), dated to the early Miocene (Burdigalian) by a $^{40}Ar/^{39}Ar$ age of 18.78±0.08 Mya on a volcanic ash sampled near the fossil (Bianucci et al., 2018), giving 18.7–18.86 Mya. Crown Odontoceti.

***Maiabalaena nesbittae* Peredo et al. 2018** Holotype from the Alsea Formation of Oregon, of early Oligocene age (Peredo et al., 2018a), giving 27.82–33.9 Mya. Stem Mysticeti: a toothless mysticete recovered rootward of the crown by Peredo et al. (2018a).

***Maiacetus inuus* Gingerich et al. 2009** Holotype (GSP-UM 3475a) from the upper Habib Rahi Formation at Kunvit, Kohlu District, eastern Balochistan, Pakistan (Gingerich et al., 2009), of early middle Eocene (Lutetian) age, giving 45.5–48.0 Mya. Stem Cetacea.

***Makaracetus bidens* Gingerich et al. 2005** Holotype (GSP-UM 3570) from the Domanda Formation at Kunvit, Sulaiman Range, Balochistan, Pakistan (Gingerich et al., 2005), of middle Eocene (Lutetian) age, giving 41.03–44.0 Mya. Stem Cetacea.

***Mamaziphius reyesi* Bianucci et al. 2024** Holotype (MUSM 4687), a mostly complete skull from the Cerros la Mama y la Hija locality (allomember P1) of the Pisco Formation (Bianucci et al., 2024), dated to the mid-Tortonian (9.0–9.1 Mya) (Bianucci et al., 2024). Crown Ziphiidae: recovered nested within the crown ziphiids as sister to the berardiines (Bianucci et al., 2024).

***Mammalodon colliveri* Pritchard 1939** Holotype (NMV P199986) from the upper Jan Juc Marl at Bird Rock, Victoria, below the Bird Rock cap (Pritchard, 1939; Duncan et al., 2025), the horizon above which is $^{87}Sr/^{86}Sr$-dated to 24.2 Mya (Duncan et al., 2025), giving 24.2–26.05 Mya. Stem Mysticeti: a mammalodontid, an archaic toothed mysticete outside crown Mysticeti (Marx & Fordyce, 2015; Duncan et al., 2025).

***Mammalodon hakataramea* Fordyce & Marx 2016** Holotype (OU 22026) from the Kokoamu Greensand at Haughs Quarry, Hakataramea Valley (Fordyce & Marx, 2016), of Duntroonian age, ∼27 Mya (Robinson et al., 2024), giving 25.2–27.3 Mya. Stem Mysticeti: see *Mammalodon colliveri*.

***Marzanoptera bertae* (Boessenecker 2013)** Holotype (UCMP 219078) from the uppermost San Gregorio section of the Purisima Formation, bracketed between ash beds correlated with the Putah Tuff and the Ishi Tuff (Boessenecker, 2013), giving 2.5–3.35 Mya. Total group Balaenopteroidea.

***Marzanoptera tersillae* Bisconti et al. 2021** Holotype (207.13307, EGPPA-MPTA, Asti), a partial skull with periotic, seven vertebrae and nine ribs, from the higher part of the Argille Azzurre Formation (close to the overlying Sabbie d'Asti Formation) at San Marzanotto d'Asti (Piedmont, Italy), in the earliest Piacenzian foraminiferal zone MPL4a, giving 3.2–3.6 Mya (Bisconti et al., 2021b). Total group Balaenopteroidea.

***Masracetus markgrafi* Gingerich 2007** Holotype (SMNS 11414), a lumbar vertebral column collected near Dimeh, from the Birket Qarun Formation of the Fayum (Gingerich, 2007), Priabonian, giving 33.90–37.71 Mya. Stem Cetacea.

***Matapanui waihao* (Boessenecker & Fordyce 2017)** Holotype from the Kokoamu Greensand of the Waitaki region, South Island, New Zealand (Boessenecker & Fordyce, 2017), of late Oligocene (early Chattian, upper Whaingaroan) age, giving 27.3–28.1 Mya. Stem Mysticeti.

***Mauicetus parki* (Benham 1937)** Holotype (OU 11573) from the upper Oligocene of the Waitaki region, South Island, New Zealand (Benham, 1937; Robinson et al., 2024), giving 21.7–25.2 Mya. Total group Mysticeti.

***Medocinia tetragorhina* (Delfortrie 1875)** Holotype, a fragmentary skull originally *Delphinus tetragorhinus* (Delfortrie, 1875) and made the type of *Medocinia* by de Muizon (1988), from the Burdigalian molasse of Saint-Médard-en-Jalle (Gironde, France) (Aguirre-Fernández et al., 2022), giving 15.98–20.45 Mya. Crown Odontoceti: a squalodelphinid within Platanistoidea (de Muizon, 1988; Aguirre-Fernández et al., 2022).

***Megabalaena sapporoensis* Tanaka et al. 2025** Holotype from the upper Toyama Formation, Sapporo, Hokkaido (Tanaka et al., 2025), dated to the late Miocene by a fission-track age of 9.0±0.4 Mya on the associated tuff (Tanaka et al., 2025), giving 8.6–9.4 Mya. Total group Balaenidae.

***Meherrinia isoni* Geisler et al. 2012** Holotype from the late Miocene Eastover Formation of North Carolina (Geisler et al., 2012), its precise horizon unconstrained and conservatively bracketed with the co-occurring *Isoninia* (Godfrey et al., 2021), giving 2.58–11.63 Mya. Total group Inioidea: a total evidence analysis recovered *Meherrinia* as sister to *Inia*, tentatively within Iniidae (Geisler et al., 2012).

***Mesocetus agrami* Van Beneden 1884** Holotype from the Sarmatian marine marls of Podsused near Zagreb (Van Beneden, 1884), dated to the Sarmatian of the Central Paratethys (Hohenegger et al., 2014; Gol'din, 2018), giving 11.6–12.8 Mya. Total group Mysticeti.

***Mesocetus aquitanicus* Flot 1896** Holotype from the marine molasse near Dax (Aquitaine, France) (Flot, 1896), referred to the Burdigalian on the associated Mont-de-Marsan material (Flot, 1896), giving 15.98–20.45 Mya. Total group Mysticeti.

***Mesocetus siphunculus* (Cope 1895)** Holotype (AMNH 22665, a mandible) from the Pamunkey River near Hanover, Virginia, in the Calvert Formation (Cope, 1895); the type carries no measured horizon and Kellogg (1968) inferred it to lie near the base of the Calvert Formation, so the bracket spans the formation from its base to the top, giving 13.0–22.0 Mya (Carnevale & Godfrey, 2018). Total group Mysticeti.

***Mesokentriodon protohumboldti* Lambert et al. 2026** Holotype (MNHN.F.PPI282) from the Santa Rosa locality (allomember P0) of the Pisco Formation (Lambert et al., 2026), dated to the Langhian–Serravallian (Middle Miocene) by the P0 allomember age (Lambert et al., 2026), giving 12.4–14.8 Mya. Total group Delphinida: a kentriodontid dolphin whose relationships are unclear (Lambert et al., 2026).

***Mesoplodon blasiusi* (Brandt 1873)** Holotype from the Neogene of the Taman Peninsula (northern Caucasus, Russia) (Brandt, 1873b), referred to the Eastern Paratethys Sarmatian *sensu lato* (Palcu et al., 2017, 2021), giving 7.65–12.65 Mya. Total group Ziphiidae.

***Mesoplodon posti* Lambert & Louwye 2016** Holotype, a 19th-century cranium from the "Sable supérieur d'Anvers", referred to the Kattendijk and Lillo Formations (Zanclean) (Lambert & Louwye, 2016), giving 3.6–5.333 Mya; palynology of sediment in the holotype indicates 3.9–4.86 Mya (Lambert & Louwye, 2016). Total group *Mesoplodon*: a beaked whale referred to the extant genus and the oldest record of *Mesoplodon* (Bianucci et al., 2016; Lambert & Louwye, 2016); crown *Mesoplodon* is not bracketed on the backbone, so the total group is assigned.

***Messapicetus gregarius* Bianucci et al. 2010** Holotype (MUSM 1037) from Cerro Colorado, allomember P1 of the Pisco Formation (Bianucci et al., 2010), dated to the Tortonian, younger than the 9.10±0.04 Mya Cerro Colorado ash (Bianucci et al., 2016), giving 8.5–9.14 Mya. Stem Ziphiidae: the type of the stem *Messapicetus* clade, outside crown Ziphiidae (Lambert et al., 2015a; Bianucci et al., 2016).

***Messapicetus longirostris* Bianucci et al. 1992** Holotype, a nearly complete skull, from the Cisterna quarry near Lecce, in the upper part of the Pietra Leccese (Apulia, Italy) (Bianucci et al., 1992), of early Tortonian age (*Neogloboquadrina acostaensis* Zone), giving 8.14–10.5 Mya (Bianucci & Landini, 2006). Stem Ziphiidae: the basal member of the *Messapicetus* clade, recovered outside the ziphiid crown (Bianucci et al., 2016).

***Metasqualodon symmetricus* Okazaki 1982** Holotype from Umashima Island, Fukuoka (Okazaki, 1982), whose only exposed unit is the Jinnobaru Formation of the Ashiya Group, its base dated to 28.91 ± 0.2 Mya by SHRIMP zircon U–Pb (Tsai et al., 2024b), giving 27.82–29.11 Mya. Stem Mysticeti.

***Metopocetus durinasus* Cope 1896** Holotype (USNM 8518), a fragmentary cranium with the right periotic in situ, from a Miocene marl near the mouth of the Potomac River at Nomini Cliffs, Westmoreland County, Virginia (Cope, 1896; Marx et al., 2016); the type formation is disputed between the Calvert and St. Marys Formations, so set the age bracket to span both formations, dated to the middle–late Miocene by the Nomini Cliffs stratigraphy (Weems et al., 2017; Godfrey & Lambert, 2023), giving 7.5–14.4 Mya. Crown Mysticeti: placed in Cetotheriidae by Marx et al. (2016).

***Metopocetus hunteri* Marx et al. 2016** Holotype (NMR 9991-07729) from the Hoogdonk sand pit at Liessel, North Brabant, the Netherlands (Diessen Formation, formerly Breda Formation), dinocyst Zone SNSM14 (Marx et al., 2016), late Tortonian, giving 7.5–8.1 Mya. Crown Mysticeti: recovered within Cetotheriidae as sister to *M. durinasus* (Marx et al., 2016), a family nested inside crown Mysticeti (Marx & Fordyce, 2015).

***Micromysticetus rothauseni* Sanders & Barnes 2002** Holotype (ChM PV4844), a skull from the Ashley Formation near Charleston, South Carolina (Sanders & Barnes, 2002a), dated by $^{87}Sr/^{86}Sr$ correlation of the Ashley Formation (Weems et al., 2016; Boessenecker & Geisler, 2018), giving 26.57–29.0 Mya. Stem Mysticeti.

***Micromysticetus tobieni* (Rothausen 1971)** Holotype from the “upper Chattian A” beds (calcareous nannoplankton Zone NP24) at Lank-Latum, near Krefeld, Germany (Rothausen, 1971), giving 23.04–27.3 Mya. Stem Mysticeti.

***Microphocaena podolica* Kudrin & Tatarinov 1966** Holotype from the Volhynian ("Volyn’ horizon") of Brykiv (Ukraine), in the lower Eastern Paratethys Sarmatian s.l. (Kudrin & Tatarinov, 1966), dated to 12.1–12.65 Mya by the Eastern Paratethys stage framework (Palcu et al., 2017, 2021). Total group Odontoceti.

***Miminiacetus pappus* (Kellogg 1955)** Holotype (USNM V 15985, a skull with mandibles and teeth) from Shattuck Zone 11 (Plum Point Member) of the Calvert Formation, Calvert County, Maryland (Kellogg, 1955; Godfrey & Lambert, 2023), of middle Miocene (Langhian) age, giving 13.82–15.98 Mya. Total group Delphinida.

***Miobalaenoptera numataensis* Tanaka & Watanabe 2019** Holotype from the lower Horokaoshirarika Formation, Numata, Hokkaido (Tanaka & Watanabe, 2019), dated to the late Miocene (diatom Zone NPD 7A), giving 6.5–6.8 Mya. Total group Balaenopteroidea.

***Miocaperea pulchra* Bisconti 2012** Holotype (SMNS 46978) from Aguada de Lomas in the Pisco Formation, Sacaco Basin, southern Peru (Bisconti, 2012), dated by the Aguada de Lomas vertebrate level of Ochoa et al. (2021), giving 7.9–8.8 Mya. Crown Mysticeti: recovered as a pygmy right whale (Neobalaenidae) close to the extant *Caperea* by Bisconti (2012).

***Miodelphinus miensis* Tanaka & Nakagawa 2024** Holotype from the Iu Mudstone Member of the Haze Formation, Mie (Tanaka & Nakagawa, 2024), of early Miocene age, giving 18.5–18.7 Mya. Crown Odontoceti: a squalodelphinid (Tanaka & Nakagawa, 2024) within Platanistoidea (Bianucci et al., 2018).

***Miodelphis californicus* Wilson 1935** Holotype from the Pyramid Hill Sand Member of the Jewett Sand, Kern County, California (Wilson, 1935), dated by the San Joaquin Basin chronostratigraphy of Scheirer and Magoon (2008), giving 24–25 Mya. Total group Odontoceti.

***Miophocaena nishinoi* Murakami et al. 2012** Holotype from the upper Miocene Koetoi Formation, Hokkaido, dated to diatom Zone NPD 7A (Murakami et al., 2012b), giving 5.5–6.4 Mya. Total group Phocoenidae: recovered outside crown Phocoenidae by Colpaert et al. (2015).

***Miophyseter chitaensis* Kimura & Hasegawa 2022** Holotype from the middle part of the Toyohama Formation on the Chita Peninsula, Aichi (Kimura & Hasegawa, 2022), placed in the early Miocene by the magnetostratigraphic correlation of its horizon to polarity chron C5Dr (Kimura & Hasegawa, 2022), giving 17.53–18.06 Mya. Total group Physeteroidea.

***Mirocetus riabinini* Mchedlidze 1970** Holotype from the Oligocene Maikop Group of Azerbaijan (Mchedlidze, 1970), giving 27.82–33.9 Mya. Total group Odontoceti.

***Mithridatocetus adygeicus* (Tarasenko & Lopatin 2012)** Holotype from the lower Blinovo Formation (Blinovskaya beds) of Adygea, NW Caucasus (Russia) (Tarasenko & Lopatin, 2012a), of middle–late Sarmatian (Bessarabian–Khersonian) age, giving 7.65–11.8 Mya (Palcu et al., 2021). Total group Mysticeti.

***Mithridatocetus eichwaldi* Gol'din & Startsev 2017** Holotype from the Kamysh-Burun (Arshintsevo) quarries of the Kerch Peninsula (Ukraine), late Sarmatian s.l. (Khersonian) (Gol'din & Startsev, 2017), giving 7.65–9.6 Mya (Palcu et al., 2021). Total group Mysticeti.

***Mixocetus elysius* Kellogg 1934** Holotype from the Modelo Formation at Los Angeles (Kellogg, 1934), of poorly constrained late Miocene age, giving 7.246–11.63 Mya. Crown Mysticeti.

***Morenocetus parvus* Cabrera 1926** Holotype (MLP 5-11) from the lower Miocene Gaiman Formation of Chubut, Argentina (Cabrera, 1926), constrained to the early Miocene by its position above the Colhuehuapian Trelew Member (Buono et al., 2017), giving 15.98–20.45 Mya. Stem Balaenidae: recovered as the earliest-diverging balaenid by Buono et al. (2017).

***Mycteriacetus bellunensis* (Pilleri 1985)** Holotype (MGPD 26404), from the upper part of the Libano Sandstone at Libano (Belluno, Italy), originally *Eurhinodelphis bellunensis* (Pilleri, 1985) but later revised as the type of *Mycteriacetus* by Lambert (2004), of late Aquitanian–early Burdigalian age, giving 19.0–22.0 Mya (Nelson et al., 2025c). Total group Odontoceti.

***Mystacodon selenensis* Lambert et al. 2017** Holotype (MUSM 1917) from the upper Eocene Yumaque Member of the Paracas Formation at Playa Media Luna, southern Peru (Lambert et al., 2017c), giving 36.61–37.71 Mya. Stem Mysticeti: recovered as the earliest-diverging toothed mysticete, outside crown Mysticeti, by Lambert et al. (2017c).

***Nalacetus ratimitus* Thewissen & Hussain 1998** Holotype (H-GSP 18521) from a channel-fill conglomerate of the Kuldana Formation at locality H-GSP 62, northern Pakistan (Thewissen & Hussain, 1998), of early middle Eocene age, giving 46.0–49.0 Mya. Stem Cetacea.

***Nannocetus eremus* Kellogg 1929** Holotype (UCMP 26502) from the Towsley Formation, Los Angeles County, California (Kellogg, 1929a; Whitmore & Barnes, 2008), of late Miocene (Delmontian) age (Whitmore & Barnes, 2008), giving 5.2–7.7 Mya. Total group Mysticeti.

***Nannolithax gracilis* Kellogg 1931** Holotype from the Sharktooth Hill bonebed in the Round Mountain Silt, Kern County, California (Kellogg, 1931), dated by the bonebed to the middle Miocene (Pyenson et al., 2009), giving 15.2–15.98 Mya. Total group Odontoceti.

***Nanokogia isthmia* Vélez-Juarbe et al. 2015** Holotype (UF 280000) from the Piña facies of the Chagres Formation at Piña 1, Colón Province, Panama (Vélez-Juarbe et al., 2015), constrained to the Messinian by planktonic foraminifera and superposition above the Toro Member (Pyenson et al., 2015), giving 5.8–6.4 Mya. Total group Kogiidae: recovered as sister to *Praekogia* and *Kogia* by Vélez-Juarbe et al. (2015).

***Natchitochia jonesi* Uhen 1998** Holotype (USNM 16805) from the late middle Eocene (Bartonian) Cook Mountain Formation of Louisiana (Uhen, 1998), giving 37.71–41.03 Mya. Stem Cetacea.

***Nazcacetus urbinai* Lambert et al. 2009** Holotype (MUSM 949) from Member F of the Pisco Formation at Cerro los Quesos (Lambert et al., 2009), dated to the late Miocene by diatom biostratigraphy of the Member F base and the overlying 6.71±0.02 Mya ash (Di Celma et al., 2016), giving 6.69–7.35 Mya. Total group Ziphiidae: morphology places it at the stem/crown ziphiid boundary (Lambert et al., 2009; Bianucci et al., 2016), but the crown position depends on *Tasmacetus* being a basal crown ziphiid, which molecular phylogenetics contradicts (McGowen et al., 2020).

***Nehalaennia devossi* Bisconti et al. 2019** Holotype (NMR 9991-00014035) dredged from area 6d of the Westerschelde estuary, the Netherlands (Breda Formation), the dinocyst of the adhering matrix dated to the late Tortonian at 8.1–8.7 Mya (Bisconti et al., 2019). Total group Balaenopteroidea: a balaenopterid recovered as sister to *Archaebalaenoptera* (Bisconti et al., 2019).

***Neosqualodon assenzae* Dal Piaz 1904** Holotype from the lower–middle Miocene of Sicily (Dal Piaz, 1904), giving 13.82–20.45 Mya. Total group Odontoceti.

***Nihohae matakoi* Coste et al. 2023** Holotype (OU 22397) from the Kokoamu Greensand/Otekaike Limestone contact at Island Cliff, Tokarahi (Coste et al., 2023), assigned to the uppermost Duntroonian at ∼25.2 Mya (Robinson et al., 2024), giving 25.2–25.5 Mya. Total group Odontoceti.

***Nihoroa reimaea* Coste et al. 2024** Holotype (OU 22126) from the Otekaike Limestone at Awamoko Creek, Waitaki Valley (Coste et al., 2024), whose matrix foraminifera include *Globoquadrina dehiscens* (first appearance datum at the base of the Waitakian), placing it no older than 25.2 Mya and likely 23–24 Mya (Coste et al., 2024), giving 23.15–25.2 Mya. Total group Odontoceti: a waipatiid-grade dolphin recovered as the closest relative of *Nihohae matakoi* (Coste et al., 2024).

***Ninjadelphis ujiharai* Kimura & Barnes 2016** Holotype from the Hiramatsu Formation, Awa Group, Mie (Kimura & Barnes, 2016), its horizon correlated with foraminiferal Zone N7 and capped by the overlying Makino Formation (Fujiwara et al., 2005; Kimura & Barnes, 2016), giving 16.4–18.5 Mya. Crown Odontoceti: an allodelphinid (Kimura & Barnes, 2016), a subclade of Platanistoidea within crown Odontoceti (Boersma & Pyenson, 2016).

***Ninoziphius platyrostris* Muizon 1983** Holotype (MNHN SAS 941) from the Sud-Sacaco (SAS) horizon of the Pisco Formation (de Muizon, 1983c; de Muizon & DeVries, 1985), dated to the Messinian by $^{87}Sr/^{86}Sr$ geochronology of the SAS horizon (Ochoa et al., 2021), giving 5.9–6.6 Mya. Stem Ziphiidae: recovered as the earliest-diverging stem beaked whale (Lambert et al., 2013; Bianucci et al., 2016).

***Norisdelphis annakaensis* Kimura & Hasegawa 2020** Holotype (GMNH-PV-3463) from the Haraichi Formation on the Usui River, Annaka, Gunma (Kimura & Hasegawa, 2020), of earliest late Miocene (Tortonian) age, giving 11.25–11.29 Mya. Total group Delphinidae: recovered as the earliest-diverging member of Delphinidae by Kimura and Hasegawa (2020).

***Norrisanima miocaena* (Kellogg 1922)** Holotype (USNM 10300) from the main Celite quarry of the Lompoc diatomite mines, Monterey Formation, California (Kellogg, 1922), dated to the latest Tortonian by diatoms of the *Rouxia californica* Subzone (NPD7A) in the cranial matrix (Leslie et al., 2019), giving 7.3–7.6 Mya. Stem Balaenopteroidea: recovered outside crown Balaenopteroidea by Leslie et al. (2019).

***Notocetus vanbenedeni* Moreno 1892** Holotype from the lower Miocene of Patagonia, Argentina (Monte León Formation) (Moreno, 1892), with Peruvian Chilcatay occurrences (Bianucci et al., 2015,

2020). The Argentine type with the Patagonian and Peruvian material spans the early Miocene, giving the 17.5–22.6 Mya age bracket. Crown Odontoceti: squalodelphinid, within Platanistoidea (Bianucci et al., 2018).

***Notoziphius bruneti* Buono & Cozzuol 2013** Holotype from the upper Miocene Puerto Madryn Formation at Playa El Doradillo, north of Puerto Madryn, Chubut, Argentina (Buono & Cozzuol, 2013), giving 10.4–11.9 Mya. Stem Ziphiidae: recovered within the stem-ziphiid *Messapicetus* clade by Buono and Cozzuol (2013).

***Numataphocoena yamashitai* Ichishima & Kimura 2000** Holotype from the upper part of the Horokaoshirarika Formation, Hokkaido (Ichishima & Kimura, 2000), dated to the early Pliocene by a fission-track age of 4.5±0.7 Mya on the underlying tuff and by the *Thalassiosira oestrupii* diatom zone (Ichishima & Kimura, 2000), giving 3.5–5.2 Mya. Stem Phocoenidae.

***Ocucajea picklingi* Uhen et al. 2011** Holotype (MUSM 1442), a partial skeleton with an incomplete skull from locality AV-19 in the Yumaque Member of the Paracas Formation, Peru (Uhen et al., 2011), Bartonian; a $^{40}$Ar/$^{39}$Ar plateau age of 36.61 Mya on ash at the base of the overlying Otuma Formation sets a hard minimum, giving 36.61–41.03 Mya (Uhen et al., 2011). Stem Cetacea.

***Odobenocetops leptodon* Muizon et al. 1999** Holotype (SMNK 2492 Pal) from the Sacaco (SAO) horizon of the Pisco Formation (de Muizon & DeVries, 1985; de Muizon et al., 1999), dated to the Messinian by U–Pb zircon ages on tuffs (5.848±0.031 and 5.741±0.056 Mya) (Ochoa et al., 2021), giving 5.7–5.9 Mya. Total group Delphinoidea: placed in Delphinoidea with affinities to Monodontidae but cannot be confidently claimed as crown (de Muizon & Domning, 2002).

***Odobenocetops peruvianus* Muizon 1993** Holotype (incomplete skull) from the Sud-Sacaco (SAS) horizon of the Pisco Formation (de Muizon & DeVries, 1985; de Muizon, 1993), dated to the Messinian by $^{87}$Sr/$^{86}$Sr geochronology of the SAS horizon (Ochoa et al., 2021), giving 5.9–6.6 Mya. Total group Delphinoidea: see *Odobenocetops leptodon*.

***Oedolithax mira* Kellogg 1931** Holotype from the Sharktooth Hill bonebed of the Round Mountain Silt (Kellogg, 1931), correlated with magnetochron C5Br by the magnetostratigraphy of Pyenson et al. (2009), giving 15.2–16.0 Mya. Total group Odontoceti.

***Oligodelphis azerbajdzanicus* Aslanova & Mchedlidze 1976** Holotype from the Maikop Group at Perekishkyul, on the Sumgait River (Azerbaijan) (Aslanova & Mchedlidze, 1976), reported as Chattian (late Oligocene) (Popov et al., 2008), giving 23.04–27.3 Mya. Total group Odontoceti.

***Olympicetus avitus* Vêlez-Juarbe 2017** Holotype (LACM 5412) from the Pysht Formation, Clallam County, Washington (Vélez-Juarbe, 2017), dated to the Oligocene by the Twin River Group magnetostratigraphy of Prothero et al. (2001), giving 26.5–30.5 Mya. Stem Odontoceti: a simocetid, among the earliest-diverging odontocetes (Vélez-Juarbe, 2023).

***Olympicetus thalassodon* Velez-Juarbe 2023** Holotype (LACM 158720) from the Pysht Formation at Murdock Creek, Clallam County, Washington, dated to the late Rupelian–early Chattian span of the formation (Vélez-Juarbe, 2023), giving 26.5–30.5 Mya. Stem Odontoceti: a simocetid, one of the earliest-diverging odontocete clades (Vélez-Juarbe, 2023).

***Orcinus citoniensis* (Capellini 1883)** Holotype from Podere Poltriciano near Cetona, Tuscany, Italy (Capellini, 1883), referred to synthems S3–S6 spanning the Piacenzian (Dominici et al., 2018), giving 2.58–3.6 Mya. Crown Delphinidae: referred to the extant killer-whale genus *Orcinus* (Capellini, 1883; Dominici et al., 2018), which nests within crown Delphinidae (McGowen et al., 2020).

***Orcinus paleorca* (Matsumoto 1937)** Holotype (a tooth) from the Sanuki Formation at Naganuma, Kazusa (Matsumoto, 1937), of basal Calabrian (Early Pleistocene) age (Matsumoto, 1937), giving 0.774–1.8 Mya. Crown Delphinidae.

***Orycterocetus crocodilinus* Cope 1867** Holotype (ANSP 9126), a single tooth from the Calvert Formation of Maryland for which no precise horizon is recorded (Cope, 1867); the interval was set based on the cranium USNM V 22926 from Shattuck Zone 13, placed in the Langhian above the Sr-dated top of Zone 10 (Zimmt et al., 2022; Godfrey & Lambert, 2023), giving 13.82–15.95 Mya. Crown Physeteroidea.

***Otekaikea huata* Tanaka & Fordyce 2015** Holotype (OU 22306) from the upper Otekaike Limestone (Maerewhenua Member), Hakataramea Valley, a few centimeters below the *Protula* shellbed (Tanaka & Fordyce, 2015a), whose top yields an $^{87}$Sr/$^{86}$Sr age of 22.28 ± 0.13 Mya on *Lentipecten* as a minimum for the whale (Tanaka & Fordyce, 2015a), giving 22.28–24.61 Mya. Total group Odontoceti.

***Otekaikea marplesi* (Dickson 1964)** Holotype (OM GL 421) from the upper Miller Member of the Otekaike Limestone at Trig Z, Waitaki Valley (Dickson, 1964; Tanaka & Fordyce, 2014), of Waitakian age (≥23.9 Mya), the upper Miller Member $^{87}$Sr/$^{86}$Sr-dated to 23.6–24.2 Mya (Graham et al., 2000; Tanaka & Fordyce, 2014), giving 23.9–25.2 Mya. Total group Odontoceti.

***Otradnocetus virodovi* Mchedlidze 1984** Holotype (GNM CO 1–90), a partial skeleton from the Tarkhanian–Konkian (middle Miocene) near Otradnaya, northwestern Caucasus (Mchedlidze, 1984), giving 12.8–16.2 Mya (Gol'din, 2018). Total group Mysticeti: diverging before Cetotheriidae in the analysis of Gol'din (2018).

***Pachyacanthus suessii* Brandt 1871** Lectotype from the middle Miocene of the Vienna Basin at Nussdorf-Heiligenstadt, Austria (Brandt, 1871; Kazár, 2010), giving 11.6–12.83 Mya (Hohenegger et al., 2014). Total group Odontoceti: tentatively referred to Platanistidae by Kazár (2010).

***Pachycetus humilis* Van Beneden 1883** Type material from the Eocene phosphatic beds between the Elbe and Weser rivers (Helmstedt region, Germany) (Van Beneden, 1883; van Vliet et al., 2020), of Bartonian–Priabonian age, giving 33.9–41.03 Mya. Stem Cetacea.

***Pachycetus paulsonii* (Brandt 1873)** Holotype, vertebral centra from the Bartonian (middle Eocene) near Chyhyryn, Ukraine (Brandt, 1873a), giving 37.7–41.0 Mya. Stem Cetacea.

***Pachycetus wardii* (Uhen 1999)** Holotype (USNM 310633) from the Comfort Member of the Castle Hayne Formation at Lanier's Pit, Maple Hill, North Carolina (Uhen, 1999), the Comfort Member assigned to calcareous-nannofossil Zone NP16 (Weems et al., 2011), giving 40.4–42.87 Mya. Stem Cetacea.

***Pakicetus attocki* (West 1980)** Holotype (H-GSP 1694) from a channel-fill conglomerate of the Kuldana Formation at locality H-GSP 62, Ganda Kas region, northern Pakistan (West, 1980), of early middle Eocene age, giving 46.0–49.0 Mya. Stem Cetacea.

***Pakicetus calcis* Cooper et al. 2009** Holotype (H-GSP 96334) from a coastal limestone of the Kuldana Formation at Shepherd's Lake, Kala Chitta Hills, northern Pakistan (Cooper et al., 2009), of middle Eocene (Lutetian) age, giving 46.0–49.0 Mya. Stem Cetacea.

***Pakicetus chittas* Cooper et al. 2009** Holotype (H-GSP 97254) from a coastal limestone of the Kuldana Formation at Shepherd's Lake, Kala Chitta Hills, northern Pakistan (Cooper et al., 2009), of middle Eocene (Lutetian) age, giving 46.0–49.0 Mya. Stem Cetacea.

***Pakicetus inachus* Gingerich & Russell 1981** Holotype (GSP-UM 084) from a fluvial conglomerate of the Kuldana Formation at Chorlakki, Kohat District, northern Pakistan (Gingerich & Russell, 1981), of early middle Eocene age, giving 46.0–49.0 Mya. Stem Cetacea.

***Palaeophocaena andrussowi* Abel 1905** Holotype from the Taman Peninsula (northern Caucasus, Russia) (Abel, 1905a), referred to the Eastern Paratethys Sarmatian *sensu lato* (Palcu et al., 2017, 2021), giving 7.65–12.65 Mya. Total group Odontoceti.

***Papahu taitapu* Aguirre-Fernández & Fordyce 2014** Holotype (OU 22066) from the Kaipuke Siltstone at Hansen Creek, northwest Nelson (Aguirre-Fernández & Fordyce, 2014), directly $^{87}$Sr/$^{86}$Sr-dated on the holotype to 22.73 $\pm$ 0.53 Mya (Marx et al., 2024) and assigned an Early Miocene (Waitakian–Otaian) age of 22.2–23.26 Mya (Robinson et al., 2024). Total group Odontoceti: recovered as an early Miocene stem odontocete by Aguirre-Fernández and Fordyce (2014).

***Pappocetus lugardi* Andrews 1920** Holotype (NHMUK PV M 11414, a partial mandible) from the Ameki Formation, southern Nigeria (Andrews, 1920), of middle Eocene (Bartonian) age, giving 37.71–41.03 Mya. Stem Cetacea.

***Parabalaenoptera baulinensis* Zeigler et al. 1997** Holotype from the Santa Cruz Mudstone at Bolinas Point, Point Reyes Peninsula, California, dated to the late Miocene by diatoms of subzone a of the *Nitzschia reinholdii* Zone in the type stratum (Zeigler et al., 1997), giving 6.0–6.8 Mya. Total group Balaenopteroidea.

***Parapontoporia pacifica* Barnes 1984** Holotype (UCR 21244, a skull) from Locality 11 in the lower member of the Almejas Formation, Isla Cedros, Baja California (Barnes, 1984), dated to the late Miocene by correlation of the lower Almejas Formation to 6–8 Mya (Boessenecker, 2013). Crown Odontoceti: a long-snouted river-dolphin relative of lipotoid/pontoporioid grade (Barnes, 1984).

***Parapontoporia sternbergi* (Gregory & Kellogg 1927)** Holotype from the San Diego Formation of California (Gregory & Kellogg, 1927), dated to the Pliocene–early Pleistocene range of the formation (Boessenecker, 2013), giving 1.8–4.2 Mya. Crown Odontoceti: recovered as the sister taxon to *Lipotes* by Geisler et al. (2011).

***Parapontoporia wilsoni* Barnes 1985** Holotype from the Purisima Formation of Santa Cruz, California (Barnes, 1985a), its type horizon now identified as the "Crab Marker" bed at the 5.33 Mya Miocene–Pliocene boundary (Boessenecker, 2013), giving 5.2–5.5 Mya. Crown Odontoceti: recovered as the sister taxon to *Lipotes* by Geisler et al. (2011).

***Parietobalaena affinis* (Van Beneden 1880)** Holotype from the Miocene of the Antwerp area, Belgium (Van Beneden, 1880), of late Miocene (Tortonian) age, giving 7.6–9.5 Mya. Total group Mysticeti.

***Parietobalaena campiniana* Bisconti et al. 2013** Holotype (RBINS M.399-R.4018), a partial skeleton with skull and ear bones from Kessel in the Antwerpen Sand Member of the Berchem Formation (Bisconti et al., 2013), Langhian–Serravallian (Louwye et al., 2000, 2020), giving 12.8–15.98 Mya. Total group Mysticeti.

***Parietobalaena laxata* (Van Beneden 1880)** Lectotype (IRScNB CtM 727) from the Antwerpen Sand Member of the Berchem Formation (Van Beneden, 1880; Steeman, 2010), dated to the Langhian–mid-Serravallian by dinocysts (Louwye et al., 2020), giving 12.8–15.97 Mya. Total group Mysticeti.

***Parietobalaena palmeri* Kellogg 1924** Holotype (USNM 10668, a skull) from Shattuck Zone 11 (Plum Point Member) of the Calvert Formation, well above the Zone 10 shell bed (Kellogg, 1924); material spans Shattuck Zones 10–16A, whose base is fixed by the $^{87}Sr/^{86}Sr$ age of Zone 10 at 16.60 Mya (Zimmt et al., 2022) and whose top is dated to 13.0 Mya (Godfrey & Lambert, 2023), giving 13.0–16.6 Mya. Total group Mysticeti.

***Parietobalaena securis* Kellogg 1931** Holotype from the Sharktooth Hill bonebed of the Round Mountain Silt (Kellogg, 1931), correlated with magnetochron C5Br by the magnetostratigraphy of Pyenson et al. (2009), giving 15.2–16.0 Mya. Total group Mysticeti.

***Parietobalaena yamaokai* Otsuka & Ota 2008** Holotype from the Itahashi Formation, Bihoku Group, at Shobara, Hiroshima, its type horizon assigned to calcareous nannofossil Zone NN4 (Otsuka & Ota, 2008; Tsai, 2017), giving 15.6–16.9 Mya. Total group Mysticeti.

***Patriocetus denggi* Abel 1914** Holotype from the Linz Sands (Linzer Sande) at Linz, Upper Austria (Abel, 1914), of late Oligocene (Chattian) age, giving 23.04–27.30 Mya (Dubrovo & Sanders, 2000). Total group Odontoceti: see *Patriocetus ehrlichii*.

***Patriocetus ehrlichii* (Van Beneden 1865)** Holotype (OL Cet. 18) from the Linz Sands (Linzer Sande) near Linz, Upper Austria, originally described as *Squalodon ehrlichii* (Van Beneden, 1865) and designated the type species of *Patriocetus* (Abel, 1914), its identity and type horizon clarified by Marx (2011); late Oligocene (Chattian), giving 23.04–27.30 Mya (Dubrovo & Sanders, 2000). Total group Odontoceti: an archaic heterodont, squalodont-grade odontocete of the *Patriocetus* group (Dubrovo & Sanders, 2000; Marx, 2011).

***Patriocetus kazakhstanicus* Dubrovo & Sanders 2000** Holotype, a skull from the Segendyk unit of the Karagiye (Karaginskaya) Formation at Karagiye, Mangyshlak Peninsula, Kazakhstan (Dubrovo & Sanders, 2000), of late Oligocene (Chattian) age, giving 23.04–27.30 Mya (Dubrovo & Sanders, 2000). Total group Odontoceti: see *Patriocetus ehrlichii*.

***Pebanista yacuruna* Benites-Palomino et al. 2024** Holotype (MUSM 4017) from the middle Miocene Pebas Formation along the Napo River, Loreto, Peruvian Amazonia (Benites-Palomino et al., 2024), giving 14.0–17.0 Mya. Crown Odontoceti: recovered as the sister taxon of the extant *Platanista* by Benites-Palomino et al. (2024).

***Pelocetus calvertensis* Kellogg 1965** Holotype (USNM 11976), a partial skeleton from Shattuck Zone 13 of the Plum Point Member of the Calvert Formation north of Governor Run, Calvert County, Maryland (Kellogg, 1965), placed in the Langhian above the Sr-dated top of Zone 10 (Zimmt et al., 2022; Godfrey & Lambert, 2023), giving 13.82–15.95 Mya. Total group Mysticeti.

***Pelocetus mirabilis* Ginsburg & Janvier 1971** Holotype from the Miocene faluns of the Anjou, Maine-et-Loire (Ginsburg & Janvier, 1971), giving 7.246–15.98 Mya. Total group Mysticeti: a "cetothere" *sensu lato* whose family-level position is unresolved (Marx et al., 2017).

***Perditicetus yaconensis* Nelson & Uhen 2020** Holotype (USNM 335224) from the Nye Formation, Oregon, of Chattian–Aquitanian (latest Oligocene to early Miocene) age (Nelson & Uhen, 2020), giving 20.44–27.82 Mya. Total group Odontoceti.

***Peregocetus pacificus* Lambert et al. 2019** Holotype (MUSM 3580), an associated skeleton from 1.95 m above the base of the Yumaque Member of the Paracas Formation at Playa Media Luna, Peru, in the upper part of calcareous nannofossil Zone CNE13 (Lambert et al., 2019a), giving 42.37–43.06 Mya. Stem Cetacea.

***Peripolocetus vexillifer* Kellogg 1931** Holotype from the Sharktooth Hill bonebed of the Round Mountain Silt (Kellogg, 1931), correlated with magnetochron C5Br by the magnetostratigraphy of Pyenson et al. (2009), giving 15.2–16.0 Mya. Total group Mysticeti: recovered as the earliest-diverging balaenid by Buono et al. (2017).

***Persufflatius renefraaijeni* Bosselaers & Munsterman 2022** Holotype (MAB 010293), a partial neurocranium from Liessel (De Brink pit), the Netherlands (Diessen Formation, dinocyst Zone SNS M14) (Bosselaers & Munsterman, 2022), late Tortonian, giving 7.6–8.2 Mya. Total group Mysticeti: a balaenomorph recovered near the base of the clade leading to modern rorquals (Bosselaers & Munsterman, 2022).

***Perucetus colossus* Bianucci et al. 2023** Holotype (MUSM 3248) from the upper Yumaque Member of the Paracas Formation south of Zamaca, Ica Province, southern Peru (Bianucci et al., 2023), dated by biostratigraphy and $^{40}$Ar/$^{39}$Ar tephrochronology to 37.84–39.8 Mya (Bianucci et al., 2023). Stem Cetacea.

***Phiomicetus anubis* Gohar et al. 2021** Holotype (MUVP 500), a partial skull and skeleton from Al Amaim in the middle Lutetian part of the Midawara Formation (Mokattamian MK4–MK5), Fayum (Gohar et al., 2021), giving 42.00–43.00 Mya. Stem Cetacea.

***Phoberodon arctirostris* Cabrera 1926** Holotype (MLP 5-4) from the lower Miocene Gaiman Formation of Chubut, Argentina (Cabrera, 1926), constrained to the early Miocene (Buono et al., 2017), giving 15.98–20.45 Mya. Total group Odontoceti: recovered as an early-diverging odontocete distant from *Squalodon* by Viglino et al. (2018a).

***Phocageneus venustus* Leidy 1869** Holotype (ANSP 11227), a single tooth from the diatomaceous deposits of the Calvert Formation near Richmond, Henrico County, Virginia (Leidy, 1869), its exact horizon and source rock unrecorded, so we set the interval span the Burdigalian Calvert Formation to the Serravallian Choptank Formation (Godfrey & Lambert, 2023), giving 11.63–20.44 Mya. Crown Odontoceti.

***Phococetus vasconum* (Delfortrie 1873)** Holotype tooth from the molasse of Saint-Médard-en-Jalle (Aquitaine, France), originally described as *Zeuglodon vasconum* (Delfortrie, 1873), of Burdigalian age (Aguirre-Fernández et al., 2022), giving 15.98–20.45 Mya. Total group Cetacea: an archaic single-tooth taxon variously interpreted as a basilosaurid archaeocete, a kekenodontid, or an early odontocete, most recently reviewed by Boessenecker (2019).

***Physeter antiquus* Gervais 1849** Holotype from the Sables à Gryphaea virleti at Montpellier, southern France (Gervais, 1849), of early Pliocene (Zanclean, mammal zone MN14) age, giving 3.60–5.33 Mya. Total group Physeteroidea.

***Physeterula dubusi* (Van Beneden 1877)** Holotype (IRSNB M.527), a skull from the Deurne / Borgerhout commune boundary, Antwerp; no precise horizon was recorded, but its bone preservation matches the late Miocene Deurne Sand Member material (Van Beneden, 1877; Lambert, 2008b), giving 7.246–11.63 Mya. Crown Physeteroidea: recovered as the most rootward member of the family Physeteridae (hence within crown Physeteroidea) by Lambert (2008b), and as a fossil physeterid alongside *Physeter* by Lambert et al. (2023) and Peri et al. (2022).

***Pictodelphis kidwellae* Godfrey & Lambert 2023** Holotype (CMM-V-8776), a partial skull from Shattuck Zone 11 of the Plum Point Member of the Calvert Formation north of Parkers Creek, Calvert County, Maryland, dated to the Langhian (Godfrey & Lambert, 2023), giving 14.5–16.1 Mya. Total group Delphinida.

***Pinocetus polonicus* Czyżewska & Ryziewicz 1976** Holotype from Nowa Wieś in the Pińczów Limestone, Carpathian Foredeep, Poland (Czyżewska & Ryziewicz, 1976), early Badenian on its *Orbulina suturalis* + Blow Zone N10 assemblage (Czyżewska & Ryziewicz, 1976), whose 15.10 Mya first-appearance datum of *O. suturalis* brackets the maximum age (Hohenegger et al., 2014), giving 13.82–15.10 Mya. Total group Mysticeti.

***Piscobalaena nana* Pilleri & Siber 1989** Holotype (SMNK PAL 4050, incomplete skull) from the Sud-Sacaco (SAS) horizon of the Pisco Formation (Bouetel & de Muizon, 2006), dated to the Messinian by $^{87}$Sr/$^{86}$Sr geochronology of the SAS horizon (Ochoa et al., 2021), giving 5.9–6.6 Mya. Crown Mysticeti: a member of Cetotheriidae *sensu stricto* (the *Cetotherium*-like group), nested within crown Mysticeti (Marx & Kohno, 2016; Bisconti et al., 2025a).

***Piscolithax boreios* Barnes 1984** Holotype (UCR 15975, a skull) from Locality 11 in the lower member of the Almejas Formation, Isla Cedros, Baja California (Barnes, 1984), dated to the late Miocene by correlation of the lower Almejas Formation to 6–8 Mya (Boessenecker, 2013). Stem Phocoenidae: an archaic porpoise recovered outside the extant porpoise in the phylogeny of Racicot et al. (2016).

***Piscolithax longirostris* Muizon 1983** Holotype (MNHN SAS 933, skull and mandible) from the Sud-Sacaco (SAS) horizon of the Pisco Formation (de Muizon, 1983b; de Muizon & DeVries, 1985), dated to the Messinian by $^{87}$Sr/$^{86}$Sr geochronology of the SAS horizon (Ochoa et al., 2021), giving 5.9–6.6 Mya. Stem Phocoenidae: recovered outside crown Phocoenidae in all recent analyses (Fajardo-Mellor et al., 2006; Colpaert et al., 2015).

***Piscolithax tedfordi* Barnes 1984** Holotype (UCR 15972, a skull) from Locality 11 in the lower member of the Almejas Formation, Isla Cedros, Baja California (Barnes, 1984) (same locality and horizon as *P. boreios*), 6.0–8.0 Mya. Stem Phocoenidae: see *Piscolithax boreios* and *Piscolithax longirostris*.

***Piscorhynchus aenigmaticus* Pilleri & Siber 1989** Holotype (partial skeleton, H. J. Siber collection) from the Aguada de Lomas (AGL) horizon of the Pisco Formation (Pilleri & Siber, 1989), dated to the Tortonian by K–Ar ages on tuffs within the AGL level (Ochoa et al., 2021), giving 7.9–8.8 Mya. Total group Odontoceti.

***Pithanodelphis cornutus* (Du Bus 1872)** Lectotype from the Miocene of Antwerp, Belgium, without stratigraphic data (Du Bus, 1872); the species is dated on the referred partial skull USNM PAL 187575 from Shattuck Zone 14 of the Plum Point Member of the Calvert Formation, St. Mary's County, Maryland, of Serravallian age (Godfrey & Lambert, 2023), giving 13.0–14.0 Mya. Total group Delphinida.

***Placoziphius duboisii* Van Beneden 1869** Holotype (IRSNB M.530) from Edegem in the Edegem Sand Member of the Berchem Formation, early Burdigalian (latest Aquitanian–early Burdigalian) (Van Beneden, 1869; Lambert, 2008b; Louwye et al., 2020); with fragments tentatively referred from the Antwerpen Sand Member, the range spans 12.8–20.44 Mya. Total group Physeteroidea: provisionally kept outside the family Physeteridae by Lambert (2008b).

***Platylithax robusta* Kellogg 1931** Holotype from the Sharktooth Hill bonebed of the Round Mountain Silt (Kellogg, 1931), correlated with magnetochron C5Br by the magnetostratigraphy of Pyenson et al. (2009), giving 15.2–16.0 Mya. Total group Odontoceti.

***Platyscaphokogia landinii* Collareta et al. 2020** Holotype (MUSM 3405) from the upper part of allomember P2 at Cerro Hueco la Zorra, Pisco Formation (Collareta et al., 2020), dated to the Messinian by a 7.084±0.044 Mya tephra below the holotype and the overlying 6.71±0.02 Mya ash (Collareta et al., 2020), giving 6.69–7.02 Mya. Total group Kogiidae: recovered in the extinct Scaphokogiinae as sister to *Scaphokogia* (Collareta et al., 2020), defined as "the most inclusive clade including *Scaphokogia* but not *Kogia*" (Collareta et al., 2017), hence rootward of the crown-*Kogia* node, which post-dates the early Pliocene (Vélez-Juarbe et al., 2015).

***Platysvercus ugonis* Guo & Kohno 2023** Holotype (UTHFM 00034) from the Sugota Formation at Tashiro, Ugo Town, Akita (Guo & Kohno, 2023), placed in the late Burdigalian (late early Miocene) (Guo & Kohno, 2023), giving 16.29–16.70 Mya. Total group Delphinida.

***Plesiobalaenoptera quarantellii* Bisconti 2010** Holotype from the Tortonian marine succession of the Stirone River near Parma, northern Italy (Bisconti, 2010), dated by macrofossil biostratigraphy to the Tortonian, giving 7.25–11.63 Mya (Bisconti, 2010). Total group Balaenopteroidea.

***Pliodelphis doelensis* Belluzzo & Lambert 2021** Holotype (IRSNB M.2330) found in situ in the Kattendijk Formation along the walls of the Deurganck Dock at Doel, Belgium (Belluzzo & Lambert, 2021), early Zanclean by dinocysts (De Schepper & Head, 2009; Belluzzo & Lambert, 2021), giving 4.4–5.0 Mya. Total group Delphinidae: an early-diverging true dolphin that could not be attributed to any delphinid subfamily (Belluzzo & Lambert, 2021).

***Pliokogia apenninica* Collareta et al. 2019** Holotype from the Sant'Andrea Bagni horizon of the Argille Azzurre (Parma, northern Italy) (Collareta et al., 2019), constrained by integrated foraminiferal, calcareous-nannofossil, and magnetostratigraphy to between 5.08 and 5.04 Mya (Collareta et al., 2019). Total group Kogiidae: a member of Kogiinae recovered as the sister group of *Praekogia* + *Kogia*, rootward of the crown-*Kogia* node (Vélez-Juarbe et al., 2015; Collareta et al., 2019).

***Pliopontos littoralis* Muizon 1983** Holotype (MNHN SAS 931, partial skull) from the Sud-Sacaco (SAS) horizon of the Pisco Formation (de Muizon, 1983a; de Muizon & DeVries, 1985), dated to the Messinian by $^{87}$Sr/$^{86}$Sr geochronology of the SAS horizon (Ochoa et al., 2021), giving 5.9–6.6 Mya. Total group Inioidea: recovered within a monophyletic Pontoporiidae (crown) by one matrix but in an unresolved basal polytomy by the other (Pyenson et al., 2015; Lambert et al., 2018a).

***Pomatodelphis bobengi* (Case 1934)** Holotype (UMMP 15117) from the lower Bone Valley Member of the Peace River Formation, Polk County, Florida (Case, 1934), of middle Clarendonian (late Miocene) age (Morgan, 1994), giving 10.0–12.0 Mya. Crown Odontoceti.

***Pomatodelphis inaequalis* Allen 1921** Holotype (MCZ 15750) from the lower (Bradley-fauna) beds of the Bone Valley Member of the Peace River Formation at Brewster Mine, Polk County, Florida (Allen, 1921b), whose age spans the late Barstovian marine-mammal biochronology to the base of the lower Peace River sequence (Morgan, 1994; Missimer, 1999), giving 8.5–13.5 Mya. Crown Odontoceti.

***Pomatodelphis santamaria* Godfrey & Lambert 2023** Holotype (CMM-V-4669), a partial skull with both periotics in situ, from the Calvert Cliffs between Seahorse Beach and Driftwood Beach, Calvert County, Maryland; *Turritella*-bearing sediment infilling the skull ties it to the Little Cove Point Member (Shattuck Zones 21–23) of the St. Marys Formation (Godfrey & Lambert, 2023), dated to the Tortonian by the Calvert Cliffs stratigraphy (Kidwell et al., 2015), giving 8.7–11.9 Mya. Crown Odontoceti: see *Pomatodelphis stenorhynchus*.

***Pomatodelphis stenorhynchus* (Holl 1829)** Holotype from the falun deposits of the Anjou, Maine-et-Loire (Holl, 1829), an undiagnosable early-nineteenth-century name applied across the Anjou–Touraine falun record with no recorded type horizon, giving 7.25–20.45 Mya. Crown Odontoceti: the genus is recovered within Platanistidae, in a clade with *Zarhachis*, by Benites-Palomino et al. (2024).

***Pontistes rectifrons* (Bravard 1858)** Holotype from the upper Miocene Ituzaingó Formation (‘Conglomerado osífero’) near Paraná, Entre Ríos, Argentina (Bravard, 1858), giving 6.67–9.47 Mya. Total group Inioidea.

***Praekogia cedrosensis* Barnes 1973** Holotype (UCR 15229) from the Arroyo Delphín locality (unit O of Tedford’s section, below the lowest prominent pebble conglomerate) in the lower member of the Almejas Formation, Isla Cedros, Baja California (Barnes, 1973), dated to the late Miocene by correlation of the lower Almejas Formation to 6–8 Mya (Boessenecker, 2013). Total group Kogiidae: a fossil pygmy sperm whale recovered as the sister taxon of *Nanokogia* among stem kogiids, outside the crown of extant *Kogia*, by Vélez-Juarbe et al. (2015).

***Praemegaptera pampauensis* Behrmann 1995** Holotype from the “Oberer Glimmerton” mica-clay at Groß Pampau, Schleswig-Holstein, Germany (Behrmann, 1995), late Langenfeldian (*Bolboforma fragori/subfragoris* Zone), ca 10.6–11.8 Mya (Kazár & Hampe, 2014). Total group Mysticeti.

***Prepomatodelphis korneuburgensis* Barnes 2002** Holotype from Teiritzberg in the Korneuburg Basin, Lower Austria (Barnes, 2002), Karpatian (nannofossil Zone NN4, *Globigerinoides bisphericus* level; basin fill ∼16.5–16.7 Mya) (Harzhauser et al., 2002), giving 16.30–17.20 Mya. Total group

Odontoceti.

***Proinia patagonica* True 1909** Holotype from the lower Miocene of the Santa Cruz Territory, Patagonia, Argentina (True, 1909), giving 17.5–22.6 Mya. Total group Odontoceti.

***Prosqualodon australis* Lydekker 1894** Holotype (MLP 5-8) from the lower Miocene Gaiman Formation at Cerro Castillo, Trelew, Chubut, Argentina (Lydekker, 1894; Gaetán et al., 2018), constrained to the early Miocene (Buono et al., 2017), giving 15.98–20.45 Mya. Total group Odontoceti: recovered as a stem odontocete by Gaetán et al. (2018).

***Protocetus atavus* Fraas 1904** Holotype (SMNS 11084), a skull with associated vertebrae and ribs from the Lower Building Stone Member of the Mokattam Formation at Gebel Mokattam (Fraas, 1904), most likely middle Lutetian (Gingerich, 1992), giving 41.03–47.80 Mya. Stem Cetacea.

***Protodelphinus capellinii* Dal Piaz 1916** Holotype from the Libano Sandstone at Libano, Belluno, northern Italy (Dal Piaz, 1916d), with a further specimen from Torrente Gresal (Pilleri, 1985); the cetacean-bearing upper Libano Sandstone is late Aquitanian to early Burdigalian (Nelson et al., 2025c), giving 19.00–22.00 Mya. Total group Odontoceti.

***Protoglobicephala mexicana* Aguirre-Fernández et al. 2009** Holotype (UABCS/ISJ/3008) from the "Los Placeritos" locality, Punta Colorada Basin, Isla San José, Baja California Sur, in the lower bedded shelly sandstone of an unnamed late Pliocene deposit (Aguirre-Fernández et al., 2009), dated by the describers to approximately 2.5–3.6 Mya (Aguirre-Fernández et al., 2009), giving 2.58–3.6 Mya. Crown Delphinidae: recovered within Globicephalinae, alongside *Globicephala*, *Pseudorca*, *Grampus* and *Orcinus*, by Aguirre-Fernández et al. (2009).

***Protophocaena minima* Abel 1905** Holotype (IRSNB 3917-M.172), a fragmentary cranium from the 19th-century Antwerp fortification excavations with no recorded member-level horizon; Abel's "Boldérien" equates to the Antwerpen Sand Member (max 15.98 Mya), and the only formation-level records are Dutch (Breda Formation, Tortonian; min 7.246 Mya) (Abel, 1905b; Lambert & Post, 2005), giving wide 7.246–15.98 Mya bracket. Total group Inioidea.

***Protororqualus cuvieri* (Fischer 1829)** Holotype from the Piacenzian of Monte Pulgnasco (Val d'Arda), Monte Giogo Formation (Fischer, 1829; Bisconti, 2007b; Bisconti & Bosselaers, 2020), dated by the *Discoaster tamalis* nannofossil zone (Bisconti, 2007b), giving 2.58–3.60 Mya. Total group Balaenopteroidea.

***Protororqualus dyticus* (Cabrera 1926)** Holotype from the lower Miocene Gaiman Formation of Chubut, Argentina (Cabrera, 1926), constrained to the early Miocene (Buono et al., 2017), giving 15.98–20.45 Mya. Total group Balaenopteroidea: an early balaenopterid more derived than *Protororqualus cuvieri* but outside the crown (Bisconti & Bosselaers, 2020).

***Protororqualus wilfriedneesi* Bisconti & Bosselaers 2020** Holotype from the Kattendijk Sands at Wommelgem, Antwerp, Belgium (Bisconti & Bosselaers, 2020), early Zanclean, ca 4.4–5 Mya (De Schepper & Head, 2009; Bisconti & Bosselaers, 2020). Total group Balaenopteroidea: see *Protororqualus dyticus*

***Pseudorca yokoyamai* Matsumoto 1926** Holotype (two teeth) from the lower part of the Sanuki Formation at Hommoku, Yokohama, Kanagawa (Matsumoto, 1926), of Calabrian (Early Pleistocene) age, giving 0.774–2.58 Mya. Crown Delphinidae.

***Pterophocaena nishinoi* Murakami et al. 2012** Holotype from the upper Miocene Wakkanai Formation, Homare District, Hokkaido, dated to the late Miocene (Murakami et al., 2012a), giving 9.2–9.3 Mya. Total group Phocoenidae: recovered as the most rootward phocoenid, outside the crown, by Colpaert et al. (2015).

***Qaisracetus arifi* Gingerich et al. 2001** Holotype (GSP-UM 3410) from a coastal shale of the Domanda Formation at Ander Dabh Shumali, eastern Sulaiman Range, Balochistan, Pakistan (Gingerich et al., 2001a), of middle Eocene (Lutetian) age, giving 41.03–44.0 Mya. Stem Cetacea.

***Rayanistes afer* Bebej et al. 2015** Holotype (CGM 42190), a partial skeleton from Qusour el Arab, Wadi El Rayan, in a middle-to-late Lutetian interval of the Midawara Formation (Bebej et al., 2015), giving 41.03–43.00 Mya. Stem Cetacea.

***Remingtonocetus domandaensis* Gingerich et al. 2001** Holotype (GSP-UM 3225) from a coastal shale of the Domanda Formation at Ander Dabh Shumali, eastern Sulaiman Range, Balochistan, Pakistan (Gingerich et al., 2001a), of middle Eocene (Lutetian) age, giving 42.0–45.5 Mya. Stem Cetacea.

***Remingtonocetus harudiensis* (Sahni & Mishra 1975)** Holotype (LUVP 11037) from a shallow subtidal mudstone of the Harudi Formation at Harudi, Kachchh, western India (Sahni & Mishra, 1975), of middle Eocene (Bartonian) age, giving 37.71–42.0 Mya. Stem Cetacea.

***Rhaphicetus valenciae* Lambert et al. 2020** Holotype (MUSM 2543) from Zamaca (locality ZM 27), Chilcatay Formation (Lambert et al., 2020), dated to the early Miocene (early Burdigalian, 18.02–19.0 Mya) (Bianucci et al., 2020; Lambert et al., 2020). Stem Physeteroidea: one of the early branching stem physeteroids, outside the crown (Lambert et al., 2020).

***Rodhocetus balochistanensis* Gingerich et al. 2001** Holotype (GSP-UM 3485) from the Habib Rahi Formation at Kunvit, eastern Balochistan, Pakistan (Gingerich et al., 2001b), of early middle Eocene (Lutetian) age, giving 45.5–48.0 Mya. Stem Cetacea.

***Rodhocetus kasranii* Gingerich et al. 1994** Holotype (GSP-UM 3012) from a coastal shale of the Domanda Formation at Bozmar Nadi, Sulaiman Range, Pakistan (Gingerich et al., 1994), of middle Eocene (Lutetian) age, giving 44.0–47.5 Mya. Stem Cetacea.

***Rododelphis stamatiadisi* Bianucci et al. 2022** Holotype (MMPS 544) from the Lindos Formation at Pefka, Rhodes, Greece (Bianucci et al., 2022), lower Calabrian (1.3–1.8 Mya, with associated *Micromesistius poutassou* otoliths further restricting the horizon to 1.3–1.5 Mya) (Bianucci et al., 2022), giving 1.30–1.80 Mya. Crown Delphinidae: a true dolphin recovered within crown Delphinidae in the analysis of Bianucci et al. (2022).

***Romaleodelphis pollerspoecki* Sanchez-Posada et al. 2024** Holotype from Traun-Pucking in the Ebelsberg Formation near Linz, Upper Austria (Sanchez-Posada et al., 2024), Aquitanian (upper Egerian; calcareous-nannofossil control NN1–NN2), giving 22.00–22.50 Mya (Sanchez-Posada et al., 2024). Total group Odontoceti.

***Rudicetus squalodontoides* (Capellini 1878)** Holotype from the Pietra Leccese near Lecce, Apulia (Capellini, 1878), recovered as a sawn building block and not tied to a dated horizon within the heterochronous formation (Bianucci, 2001), giving 6.0–18.0 Mya. Total group Delphinida: recovered within Kentriodontidae by Guo and Kohno (2023), whose competing hypotheses (sister to Delphinoidea, or stem delphinidan (Lambert et al., 2026)) both fall within Delphinida.

***Sachalinocetus cholmicus* Dubrovo 1971** Holotype from the Kholmsk Formation at the Grebenskaya locality, western Sakhalin, Russia (Dubrovo, 1971), of Oligocene–Miocene age, giving 11.63–27.30 Mya. Total group Odontoceti: a Sakhalin odontocete listed among putative waipatiid platanistoids but treated as *incertae sedis* pending redescription (Tanaka & Fordyce, 2014).

***Saghacetus osiris* (Dames 1894)** Holotype, a dentary, from the Qasr el-Sagha Formation near Garet el-Esh, Fayum (Dames, 1894; Gingerich, 1992), Priabonian, giving 33.90–37.71 Mya. Stem Cetacea.

***Salishicetus meadi* Peredo & Pyenson 2018** Holotype (UWBM 50004) from the upper Oligocene of Washington State (Peredo & Pyenson, 2018), of Chattian age, giving 23.04–27.3 Mya. Stem Mysticeti: an aetiocetid recovered crownward of Mammalodontidae on the stem to crown Mysticeti (Peredo & Pyenson, 2018).

***Salumiphocaena stocktoni* (Wilson 1973)** Holotype (UCMP 34576) from the Valmonte Diatomite Member of the Monterey Formation, Palos Verdes Hills, California (Wilson, 1973), dated to the middle–late Miocene (Racicot et al., 2016), giving 9.0–12.6 Mya. Total group Phocoenidae.

***Samaydelphis chacaltanae* Lambert et al. 2021** Holotype (MUSM 566) from Cerro Colorado, allomember P1 of the Pisco Formation (Lambert et al., 2021), dated to the Tortonian (P1, above the 9.10±0.04 Mya ash) (Bianucci et al., 2016; Lambert et al., 2021), giving 8.5–9.14 Mya. Total group Inioidea: recovered as a pontoporiid sister to *Meherrinia isoni*, whose affinity flips between Iniidae and stem Inioidea, so the crown status cannot be affirmed (Lambert et al., 2021).

***Sarmatodelphis moldavicus* Kirpichnikov 1954** Holotype (UBFG.628) from Kishinev (Chişinău), Republic of Moldova (Kirpichnikov, 1954), revised by Kazár and Grigorescu (2005), from the Bessarabian (middle Sarmatian of the Eastern Paratethys) (Kazár & Grigorescu, 2005), dated to 9.6–12.1 Mya (Palcu et al., 2017, 2021). Total group Odontoceti.

***Saurocetes argentinus* Burmeister 1871** Holotype from the upper Miocene Ituzaingó Formation (‘Conglomerado osífero’) near Paraná, Entre Ríos, Argentina (Burmeister, 1871), giving 6.0–9.47 Mya. Total group Odontoceti.

***Saurocetes gigas* (Cozzuol 1988)** Holotype from the upper Miocene Ituzaingó Formation (‘Mesopotamiense’) near Paraná, Entre Ríos, Argentina (Cozzuol, 1988), giving 6.67–9.47 Mya. Total group Odontoceti.

***Scaldicetus caretti* du Bus 1867** Holotype from the upper Miocene (Tortonian) of the Antwerp area, Belgium (Du Bus, 1867), giving 7.25–11.63 Mya. Total group Physeteroidea (Snoodijk et al., 2024).

***Scaldiporia vandokkumi* Post et al. 2017** Holotype (NMR 9991-12018) dredged from area 6D of the Westerschelde estuary, the Netherlands (Breda Formation), the adhering matrix palynologically dated to 5–7.6 Mya (Post et al., 2017). Total group Inioidea: a pontoporiid, closer to *Pontoporia* than to *Inia* (Post et al., 2017; Lambert et al., 2018a), but Pontoporiidae collapses to a basal polytomy in broader sampling (Pyenson et al., 2015), so total group is assigned.

***Scaphokogia cochlearis* Muizon 1988** Holotype from Aguada de Lomas in the Pisco Formation, Sacaco Basin, southern Peru (de Muizon, 1988), dated by the Aguada de Lomas vertebrate level of Ochoa et al. (2021), giving 7.9–8.8 Mya. Total group Kogiidae: recovered in a *Scaphokogia* clade outside crown *Kogia* by Benites-Palomino et al. (2020).

***Scaphokogia totajpe* Benites-Palomino et al. 2020** Holotype from the upper Miocene Pisco Formation, Sacaco Basin, southern Peru (Benites-Palomino et al., 2020), dated by the Sacaco-Basin chronology of Ochoa et al. (2021), giving 6.6–7.9 Mya. Total group Kogiidae: recovered as the sister taxon of *S. cochlearis* by Benites-Palomino et al. (2020).

***Schizodelphis barnesi* (Myrick 1979)** Lectotype (USNM 244403), designated by Godfrey and Lambert (2023), from Shattuck Zone 12–13 (Plum Point Member) of the Calvert Formation; the species is confined to Calvert Cliffs and ranges Shattuck Zones 4–14, which was bracketed by Godfrey and Lambert (2023) as the base of dinocyst zone DN3 to the top of Zone 14, giving 13.0–19.0 Mya. Total group Odontoceti.

***Schizodelphis bogatshowi* (Mchedlidze 1964)** Holotype from Perekishkyul (Azerbaijan), of late Sarmatian (Khersonian) age in the Eastern Paratethys (Mchedlidze, 1964), dated by the Khersonian chronology of Palcu et al. (2021) to 7.65–9.6 Mya. Total group Odontoceti.

***Schizodelphis morckhoviensis* (Du Bus 1872)** Holotype (IRSNB 3235-M.343) from the Antwerp area, Belgium (Du Bus, 1872), for which no stratigraphic horizon is recorded (Godfrey & Lambert, 2023); the age bracket is therefore set using the Calvert Cliffs record (Shattuck Zones 3B–13), which Godfrey and Lambert (2023) place from the base of dinocyst zone DN3 to the top of Zone 13, giving 14.0–19.0 Mya. Total group Odontoceti.

***Schizodelphis planus* (Gervais 1859)** Holotype from the lower Miocene molasse of Chamaret (Drôme, France) (Gervais, 1859), of Burdigalian age, giving 15.98–20.45 Mya. Total group Odontoceti.

***Schizodelphis squalodontoides* Longhi 1897** Holotype from the Belluno whetstone ("pietra da coti") of the Belluno molasse (Veneto, Italy) (Longhi, 1897), of Burdigalian age, giving 15.98–20.45 Mya. Total group Odontoceti.

***Schizodelphis sulcatus* Gervais 1861** Holotype (MNHN RL12) from Cournonsec, Hérault (Gervais, 1861), re-diagnosed on a skull from the Burdigalian of Pignan, Hérault (Benoit et al., 2011), giving 16.0–20.4 Mya. Total group Odontoceti.

***Semirostrum ceruttii* Racicot et al. 2014** Holotype (SDSNH 65276) from the San Diego Formation at Rancho del Rey, Chula Vista, California, dated to the Pliocene–early Pleistocene range of the formation (Racicot et al., 2014), giving 1.6–4.2 Mya. Total group Phocoenidae.

***Septemtriocetus bosselaersi* Lambert 2008** Holotype an isolated skull from the Verrebroek dock, Kruisschans Sand Member of the Lillo Formation, Belgium (Lambert, 2008a), dated to 2.8–3.2 Mya by dinocysts and sequence stratigraphy (Deckers et al., 2020). Stem Phocoenidae: a porpoise recovered as sister to a clade of stem phocoenids (*Haborophocoena*, *Archaeophocaena*, *Miophocaena*, *Semirostrum*) (Colpaert et al., 2015).

***Septidelphis morii* Bianucci 2013** Holotype from Settime (Asti, Italy), Sabbie d'Asti Formation, of late Zanclean–early Piacenzian age (Bianucci, 2013), giving 3.0–4.0 Mya. Crown Delphinidae: a true dolphin recovered within the crown-group radiation of Delphinidae by Bianucci (2013).

***Simocetus rayi* Fordyce 2002** Holotype (USNM 256517) from the Alsea Formation of Oregon (Fordyce, 2002), placed in the early Oligocene (Vélez-Juarbe, 2023), giving 27.82–33.9 Mya. Stem Odontoceti: type species of Simocetidae, recovered among the earliest-diverging odontocete clades by Vélez-Juarbe (2023).

***Sinanodelphis izumidaensis* Makiyama 1936** Holotype from the Bessho Formation (Makiyama, 1936), dated to the middle Miocene, giving 13.82–15.98 Mya. Total group Odontoceti.

***Sitsqwayk cornishorum* Peredo & Uhen 2016** Holotype from the Twin River Quarry, Pysht Formation, Clallam County, Washington (Peredo & Uhen, 2016), dated to the late Oligocene by the Twin River Group magnetostratigraphy of Prothero et al. (2001), giving 26.5–28.1 Mya. Stem Mysticeti: recovered within Eomysticetidae as a latest-diverging stem mysticete by Boessenecker and Richards (2024).

***Sophianaecetus commenticius* (Kazár 2005)** Holotype from Kovácsszénája (Hungary), Kozárd Formation (Kazár, 2005), of Central Paratethys Sarmatian *sensu stricto* age, dated by the Badenian–Sarmatian chronology of Hohenegger et al. (2014) to 11.6–12.83 Mya; the genus is a replacement name for the preoccupied *Mediocris* (Kazár, 2006). Total group Delphinida: a kentriodontid, recovered within Kentriodontoidea near Delphinoidea by Guo and Kohno (2023) and as a stem delphinidan by Lambert et al. (2026).

***Squalodelphis fabianii* Dal Piaz 1917** Holotype from the upper Libano Sandstone of the Belluno quarries (Dal Piaz, 1917), dated to the late Aquitanian–early Burdigalian (Nelson et al., 2025c), giving 19.0–22.0 Mya. Crown Odontoceti: recovered as a basal member of Platanistoidea by Boersma and Pyenson (2016).

***Squalodon barbarus* Mchedlidze & Aslanova 1968** Holotype from Perekishkyul (Azerbaijan), Maikop Group, of late Oligocene (Chattian) age (Mchedlidze, 1976), giving 23.04–27.3 Mya. Stem Odontoceti: a heterodont squalodontid (Rothausen, 1968), assigned as a valid, diagnostic species by the revision of Nelson et al. (2025c).

***Squalodon bellunensis* Dal Piaz 1901** Holotype (MGP 17715) from the upper Libano Sandstone of the Belluno quarries, dated to the late Aquitanian–early Burdigalian (Dal Piaz, 1901; Nelson et al., 2025c), giving 19.0–22.0 Mya. Stem Odontoceti: see *Squalodon barbarus*.

***Squalodon calvertensis* Kellogg 1923** Holotype (USNM V 10484), a nearly complete skull with periotic, teeth, and postcrania from Shattuck Zone 6 of the Calvert Formation on the western shore of Chesapeake Bay south of Chesapeake Beach, Calvert County, Maryland (Kellogg, 1923b), placed in the late Early Miocene (Burdigalian) by the zonal framework of Godfrey and Lambert (2023), giving 16.0–20.6 Mya. Stem Odontoceti: see *Squalodon barbarus*.

***Squalodon grateloupii* von Meyer 1843** Holotype from Léognan, Gironde (von Meyer, 1843); the neotype (MHNL Dr.15) is from the early Burdigalian molasse near Léognan (Nelson et al., 2025a), giving 15.98–20.44 Mya. Stem Odontoceti: see *Squalodon barbarus*.

***Squalodon murdochi* Godfrey & Lambert 2023** Holotype from Shattuck Zone 10 of the Plum Point Member of the Calvert Formation at Plum Point, Maryland, dated by Sr isotopes on Zone 10 (Zimmt et al., 2022; Godfrey & Lambert, 2023), giving 15.95–16.6 Mya. Stem Odontoceti: see *Squalodon barbarus*.

***Squalodon peregrinus* Dal Piaz 1971** Holotype from Libano, upper Libano Sandstone (Belluno, Italy) (Dal Piaz, 1971), late Aquitanian to early Burdigalian in age (Nelson et al., 2025c), giving 19.0–22.0 Mya. Stem Odontoceti: see *Squalodon barbarus*.

***Squalodon servatus* (Meyer 1841)** Type from the Upper Marine Molasse at Baltringen, Württemberg, Germany, of Burdigalian age (von Meyer, 1841), giving 15.98–20.45 Mya. Stem Odontoceti: see *Squalodon barbarus*.

***Squalodon whitmorei* Dooley 2005** Holotype from the Fairhaven Member of the Calvert Formation, Virginia (Dooley, 2005), the member correlative with the Pungo River Formation of North Carolina (Godfrey & Lambert, 2023), dated to the early Miocene, giving 19.0–22.0 Mya. Stem Odontoceti: see *Squalodon barbarus*.

***Squaloziphius emlongi* Muizon 1991** Holotype from the Clallam Formation of Washington State (Muizon, 1991), of early Miocene (Aquitanian) age, giving 20.45–23.04 Mya. Stem Odontoceti: recovered among the stem odontocetes by Geisler et al. (2011).

***Stenasodelphis russellae* Godfrey & Barnes 2008** Holotype (CMM-V-2234), the facial region of a skull with the rostrum base (Godfrey & Barnes, 2008), encased in gray clay typical of Shattuck Zones 22–23 of the Little Cove Point Member of the St. Marys Formation, Calvert County, Maryland, dated to the Tortonian by the Calvert Cliffs stratigraphy (Kidwell et al., 2015; Godfrey & Lambert, 2023), giving 8.7–11.9 Mya. Total group Inioidea.

***Stenella rayi* Whitmore & Kaltenbach 2008** Holotype (USNM 182930), a partial cranium from the Yorktown Formation at the Lee Creek Mine, North Carolina (Whitmore & Kaltenbach, 2008), early Pliocene age (Boessenecker et al., 2019), giving 3.92–4.9 Mya. Crown Delphinidae.

***Stromerius nidensis* Gingerich 2007** Holotype (CGM 42192), a series of thoracic, lumbar and caudal vertebrae from Garet el-Esh in the Temple Member of the Qasr el-Sagha Formation, Fayum (Gingerich, 2007), Priabonian, giving 33.90–37.71 Mya. Stem Cetacea.

***Sulakocetus dagestanicus* Mchedlidze 1976** Holotype from Miatla (Dagestan, Russia), of late Oligocene (Chattian) age (Mchedlidze, 1976), giving 23.04–27.3 Mya. Total group Odontoceti.

***Supayacetus muizoni* Uhen et al. 2011** Holotype (MUSM 1465) from the middle Eocene Paracas Formation, Ica Province, southern Peru, which underlies the basal Otuma Formation ash dated $^{40}$Ar/$^{39}$Ar at 36.61 Mya (Uhen et al., 2011), giving 37.71–41.03 Mya. Stem Cetacea.

***Tagicetus joneti* Lambert et al. 2005** Holotype from Penedo Norte (Lower Tagus Basin, Portugal), strontium-dated to ~11.3 and ~12.5 Mya and correlated with planktonic-foraminiferal Zones N11–N13 (Serravallian) (Lambert et al., 2005), giving 11.63–13.82 Mya. Total group Delphinida: a kentriodontine dolphin (Lambert et al., 2005), a grade regarded as stem-group Delphinida by Lambert et al. (2026).

***Taikicetus inouei* Tanaka et al. 2018** Holotype (AMP 35) from the Hikatagawa Formation, Nupinai Group, Hokkaido (Tanaka et al., 2018), dated to the middle Miocene (Langhian–Serravallian), giving 11.5–15.2 Mya. Total group Mysticeti: recovered in a clade with *Isanacetus* and *Parietobalaena*, basal to the Balaenopteroidea + Cetotheriidae clade, by Tanaka et al. (2018).

***Takracetus simus* Gingerich et al. 1995** Holotype (GSP-UM 3041) from the Domanda Formation at Takra Daab, Sulaiman Range, Punjab, Pakistan (Gingerich et al., 1995), of middle Eocene (Lutetian) age, giving 43.5–47.0 Mya. Stem Cetacea.

***Thalassocetus antwerpiensis* Abel 1905** Lectotype (IRSNB M.525); the bone color and preservation indicate an origin in the lower-to-middle Miocene Berchem Formation (probably the Antwerpen Sand Member) (Lambert, 2008b), giving 12.8–19.0 Mya. Crown Physeteroidea: an archaic kogiid (within crown Physeteroidea) by Lambert (2008b).

***Thinocetus arthritus* Kellogg 1969** Holotype (USNM 23794), a posterior cranium with ear bones and much of the postcranium from the Choptank Formation of Westmoreland County, Virginia, in a transitional layer straddling the Shattuck Zone 16B/17 contact below the Zone 17 shell bed (Kellogg, 1969; Marx & Fordyce, 2015), dated to the Serravallian (Godfrey & Lambert, 2023), giving 12.0–13.2 Mya. Total group Mysticeti.

***Tiphyocetus temblorensis* Kellogg 1931** Holotype from the Sharktooth Hill bonebed of the Round Mountain Silt (Kellogg, 1931), correlated with magnetochron C5Br by the magnetostratigraphy of Pyenson et al. (2009), giving 15.2–16.0 Mya. Total group Mysticeti.

***Titanocetus sammarinensis* (Capellini 1901)** Holotype from the Monte Fumaiolo Sandstone of San Marino (Capellini, 1901), dated to the Langhian–Serravallian (middle Miocene) (Bisconti, 2006), giving 11.63–15.98 Mya. Total group Mysticeti.

***Tiucetus rosae* Marx et al. 2017** Holotype (MNHN.F.PPI261) from the lowest allomember (P0) of the Pisco Formation, Sacaco Basin, southern Peru (Marx et al., 2017), dated to 12.4–14.8 Mya by the P0 chronology of Lambert et al. (2026). Total group Mysticeti.

***Tlaxcallicetus guaycurae* Hernández-Cisneros 2018** Holotype (MHN-UABCS EcSj5/06/31), a partial skull from the El Saladito locality, San Juan Member of the El Cien Formation, Baja California Sur (Hernández-Cisneros, 2018), dated by a U–Pb zircon age of 27.95±0.16 Mya on a tuff at the top of bed 2 of the San Juan Member (Hernández-Cisneros et al., 2023), giving 27.79–28.11 Mya. Total group Mysticeti.

***Togocetus traversei* Gingerich & Cappetta 2014** Holotype (KPG-M 1), a left dentary with M3 from the "bone-bed reposant" near the top of mining level Couche-1 in the Kpogamé–Hahotoé phosphates, Togo (Gingerich & Cappetta, 2014), planktonic-foraminiferal Zone P11 (middle Lutetian), giving 43.00–47.00 Mya. Stem Cetacea.

***Tohoraata raekohao* Boessenecker & Fordyce 2015** Holotype (OU 22178) from the Otekaike Limestone (Maerewhenua Member) at Waipati Creek, Hakataramea Valley (Boessenecker & Fordyce, 2015a), directly $^{87}Sr/^{86}Sr$-dated on the holotype to 24.20 ± 0.30 Mya (Marx et al., 2024), giving 23.9–24.5 Mya. Stem Mysticeti: an eomysticetid (Boessenecker & Fordyce, 2015a; Marx & Fordyce, 2015; Boessenecker & Richards, 2024).

***Tohoraata waitakiensis* (Marples 1956)** Holotype (OM GL 402) from the Kokoamu Greensand at Duntroon, Waitaki Valley (Marples, 1956; Boessenecker & Fordyce, 2015a), spanning the full Whaingaroan–Duntroonian Kokoamu Greensand, as the holotype level is not established (Boessenecker & Fordyce, 2015a; Tsai & Fordyce, 2018; Boessenecker & Richards, 2024), giving 25.2–27.82 Mya. Stem Mysticeti: see *Tohoraata raekohao.*

***Tohoraonepu nihokaiwaiu* Corrie & Fordyce 2024** Holotype (OU 22394) from the Kokoamu Greensand, Awamoko Valley, Waitaki (Corrie & Fordyce, 2024), placed low in the Duntroonian (∼26

Mya) by its brachiopod–pectinid horizon (Corrie & Fordyce, 2024; Robinson et al., 2024), giving 25.2–27.3 Mya. Stem Neoceti: a kekenodontid (Corrie & Fordyce, 2022, 2024).

***Toipahautea waitaki* Tsai & Fordyce 2018** Holotype (OU 21981) from the lower Kokoamu Greensand of the Waitaki Valley, South Island, New Zealand (Tsai & Fordyce, 2018), of late Oligocene (upper Whaingaroan, ∼27.5 Mya) age, giving 27.3–28.1 Mya. Total group Mysticeti: an archaic toothless baleen whale placed near the base of crown Mysticeti (Tsai & Fordyce, 2018).

***Tokarahia kauaeroa* Boessenecker & Fordyce 2015** Holotype (OU 22235) from the Otekaike Limestone at Island Cliff, Waitaki Valley (Boessenecker & Fordyce, 2015b), directly $^{87}Sr/^{86}Sr$-dated on the holotype to 24.43 ± 0.32 Mya (Marx et al., 2024), giving 24.11–24.75 Mya. Stem Mysticeti: an eomysticetid rootward of crown Mysticeti (Boessenecker & Fordyce, 2015b; Marx & Fordyce, 2015; Boessenecker & Richards, 2024).

***Tokarahia lophocephalus* (Marples 1956)** Holotype (OM GL 412) from the Kokoamu Greensand at Duntroon, Waitaki Valley (Boessenecker & Fordyce, 2015b), of Duntroonian age (Boessenecker & Fordyce, 2015b), giving 25.2–27.3 Mya. Stem Mysticeti: see *Tokarahia kauaeroa*.

***Tranatocetus argillarius* (Roth 1978)** Holotype (MGUH VP 2319) from the Gram Clay of the Gram Formation, 1.5 km north of Gram, South Jutland, Denmark (Roth, 1978; Gol'din & Steeman, 2015), mid-Tortonian by magnetostratigraphy, dinocysts and malacofauna (Rasmussen et al., 2010; Gol'din & Steeman, 2015), giving 7.2–9.9 Mya. Crown Mysticeti: type of Tranatocetidae (Gol'din & Steeman, 2015), recovered deeply nested within Cetotheriidae as sister to *Metopocetus* (Marx et al., 2019), a family lying inside crown Mysticeti (Marx & Fordyce, 2015).

***Tranatocetus maregermanicum* Marx et al. 2019** Holotype (NMR 9991-16680) dredged from site 6d of the Western Scheldt, the Netherlands (Breda Formation, dinocyst Zone SNSM14) (Marx et al., 2019), late Tortonian, giving 7.5–8.1 Mya. Crown Mysticeti: see *Tranatocetus argillarius*.

***Tupelocetus palmeri* Gibson et al. 2019** Holotype from the Cross Member of the Tupelo Bay Formation, South Carolina (Gibson et al., 2019), dated to the Bartonian by calcareous nannoplankton Zone NP17 of the Cross Member (Geisler et al., 2005; Gibson et al., 2019), giving 37.71–41.03 Mya. Stem Cetacea.

***Tusciziphius crispus* Bianucci 1997** Holotype from Case il Poggio, Val di Pugna (Siena, Italy) (Bianucci, 1997), placed at the MPl3–MPl4 transition (nannofossil Zone MNN14–15) by Bianucci et al. (2001), giving 3.84–4.12 Mya. Total group Ziphiidae.

***Tutcetus rayanensis* Antar et al. 2023** Holotype (MUVP 501, a skull with mandibles, hyoid, and atlas) from the Sath El-Hadid Formation, Wadi El-Rayan, Fayum, Egypt (Antar et al., 2023), of middle Eocene (early Bartonian) age, giving 37.71–41.03 Mya. Stem Cetacea.

***Uranocetus gramensis* Steeman 2009** Holotype (MSM P813) from the Gram Formation, South-West Jutland, Denmark (Steeman, 2009), an in-situ Tortonian assemblage (Rasmussen et al., 2010), giving 7.2–9.9 Mya. Total group Mysticeti.

***Urkudelphis chawpipacha* Tanaka et al. 2017** Holotype (MO-1) from a coastal boulder at Montañita/Olón, Santa Elena Province, Ecuador (Tanaka et al., 2017), assigned a probable Chattian age, giving 23.04–27.3 Mya. Total group Odontoceti: recovered near the base of Platanistoidea by Tanaka et al. (2017).

***Vampalus sayasanicus* Tarasenko & Lopatin 2012** Holotype (PIN 5341/1) from the Aksai River valley (Chechnya, Russia), of Middle–Late Miocene age (Tarasenko & Lopatin, 2012b), giving 5.333–15.98 Mya. Total group Mysticeti.

***Vanbreenia trigonia* Bianucci & Landini 2002** Holotype, an incomplete cranium from the Miste Bed at the base of the Aalten Member near Winterswijk, the eastern Netherlands (Bianucci & Landini, 2002), of Langhian (middle Miocene) age, giving 13.82–15.98 Mya. Total group Odontoceti.

***Waharoa ruwhenua* Boessenecker & Fordyce 2015** Holotype (OU 22044) from the Otekaike Limestone (Maerewhenua Member) at The Earthquakes, Waitaki Valley (Boessenecker & Fordyce, 2015c), directly $^{87}$Sr/$^{86}$Sr-dated on the holotype to 25.30 $\pm$ 0.35 Mya (Marx et al., 2024), giving 24.95–25.65 Mya. Stem Mysticeti.

***Waipatia hectori* (Benham 1935)** Holotype (NMNZ Ma 653) from the Maerewhenua Member of the Otekaike Limestone, Otekaike River, Waitaki Valley (Benham, 1935; Tanaka & Fordyce, 2015b), of upper Duntroonian age; matrix bearing foraminifera *Turborotalia euapertura* but lack *Globoquadrina dehiscens*, so older than the 25.2 Mya Waitakian base (Tanaka & Fordyce, 2015b), giving 25.2–27.3 Mya. Total group Odontoceti.

***Waipatia maerewhenua* Fordyce 1994** Holotype (OU 22095) from the Maerewhenua Member of the Otekaike Limestone at The Earthquakes, Waitaki Valley (Fordyce, 1994), of Duntroonian age (Robinson et al., 2024), giving 25.2–27.3 Mya. Total group Odontoceti.

***Westmorelandelphis tacheroni* Godfrey & Lambert 2023** Holotype from the Choptank Formation of Virginia (Godfrey & Lambert, 2023), dated to the Serravallian, giving 11.5–13.0 Mya. Total group Delphinida.

***Whakakai waipata* Tsai & Fordyce 2016** Holotype (OU 21927) from the Kokoamu Greensand at The Earthquakes, Waitaki Valley (Tsai & Fordyce, 2016), of Duntroonian age (Robinson et al., 2024), giving 25.2–27.3 Mya. Total group Mysticeti.

***Wimahl chinookensis* Peredo et al. 2018** Holotype from the lower Miocene Astoria Formation of Washington State (Peredo et al., 2018b), giving 15.98–23.04 Mya. Total group Delphinida.

***Xenorophus simplicidens* Boessenecker & Geisler 2023** Holotype (CCNHM 8720) from Bed 2 of the Chandler Bridge Formation near North Charleston, South Carolina (Boessenecker & Geisler, 2023), dated by $^{87}$Sr/$^{86}$Sr correlation of the Chandler Bridge Formation (Weems et al., 2016; Boessenecker & Geisler, 2018), giving 23.5–24.7 Mya. Stem Odontoceti: recovered as an early-diverging xenorophid by Boessenecker and Geisler (2023).

***Xenorophus sloanii* Kellogg 1923** Holotype (USNM 11049) from the Ashley Formation near Ladson, South Carolina (Kellogg, 1923a; Boessenecker & Geisler, 2023), dated by $^{87}$Sr/$^{86}$Sr correlation of the Ashley Formation (Weems et al., 2016; Boessenecker & Geisler, 2018), giving 26.57–29.0 Mya. Stem Odontoceti: see *Xenorophus simplicidens*.

***Xiphiacetus bossi* (Kellogg 1925)** Holotype (USNM 8842, a partial skeleton), described as *Eurhinodelphis bossi* (Kellogg, 1925b) and transferred to *Xiphiacetus* by Lambert (2005b), from Shattuck Zones 3–6 of the Calvert Formation, Maryland; along Calvert Cliffs the species ranges Shattuck Zones 3B–14, which was bracketed by Godfrey and Lambert (2023) from the base of dinocyst zone DN3 to the top of Zone 14, giving 13.0–19.0 Mya. Total group Odontoceti: a long-snouted eurhinodelphinid, an archaic odontocete grade retained at the total group (Lambert, 2005b; Godfrey & Lambert, 2023).

***Xiphiacetus cristatus* (du Bus 1872)** Holotype (IRSNB 3234-M.361) from the Berchem Formation, Antwerp, Belgium (Du Bus, 1872; Lambert, 2005b), placed in the Middle Miocene, giving

14.0–14.9 Mya. Total group Odontoceti: see *Xiphiacetus bossi*.

***Yamatocetus canaliculatus* Okazaki 2012** Holotype from the Jinnobaru Formation of the Ashiya Group at the Wakamatsu coast, Kitakyushu (Okazaki, 2012), its upper Oligocene horizon lying above a formation base dated to 28.91 ± 0.2 Mya by SHRIMP zircon U–Pb (Tsai et al., 2024b), giving 27.82–29.11 Mya. Stem Mysticeti: recovered within a monophyletic Eomysticetidae, among the latest-diverging stem mysticetes, by Boessenecker and Richards (2024).

***Yaquinacetus meadi* Lambert et al. 2019** Holotype from the upper Nye Mudstone north of Thiel Creek, Lincoln County, Oregon, of latest Chattian to earliest Burdigalian age (Lambert et al., 2019b), giving 19.2–24 Mya. Stem Odontoceti.

***Zarhachis crassangulum* (Case 1904)** Holotype, a partial skull from Shattuck Zone 13 of the Calvert Formation at Governor Run, Calvert County, Maryland (Case, 1904), dated to the late Middle Miocene by the Calvert Cliffs zonal framework of Godfrey and Lambert (2023), giving 14.0–14.9 Mya. Crown Odontoceti.

***Zarhachis flagellator* Cope 1868** Holotype from the Calvert Beach Member (Shattuck Zones 14–16A) of the Calvert Formation on the James T. Thomas estate, Maryland (Cope, 1868), dated to the late Middle Miocene by the Calvert Cliffs zonal framework of Godfrey and Lambert (2023), giving 13.0–14.6 Mya. Crown Odontoceti.

***Zarhachis tysonii* Cope 1869** Holotype, a single posterior lumbar vertebra (ANSP), from the St. Marys Formation near the mouth of the Patuxent River, Calvert County, Maryland (Cope, 1869b; Case, 1904), from the Little Cove Point Member dated to the Tortonian by the Calvert Cliffs dinocyst stratigraphy (Kidwell et al., 2015; Weems et al., 2017), giving 8.7–11.9 Mya. Crown Odontoceti.

***Zarhachis velox* Cope 1869** Holotype (ANSP) from the Shiloh Marl of the Kirkwood Formation, southern New Jersey (Cope, 1869b), dated to the early Miocene by Sr-isotope and diatom biostratigraphy of the Shiloh Marl (Sugarman et al., 1993), giving 19.5–20.8 Mya. Crown Odontoceti.

***Zarhinocetus donnamatsonae* Kimura & Barnes 2016** Holotype (UCMP 86139) from an undetermined level of the Astoria Formation near Elma, Grays Harbor County, Washington (Kimura & Barnes, 2016), spanning the formation's magnetostratigraphic range (Prothero et al., 2008), giving

15.2–19.2 Mya. Crown Odontoceti: an allodelphinid, recovered within crown Odontoceti (Boersma & Pyenson, 2016).

***Zarhinocetus errabundus* (Kellogg 1931)** Holotype (CAS 905) from the Sharktooth Hill bonebed of the Round Mountain Silt (Kellogg, 1931), correlated with magnetochron C5Br by the magnetostratigraphy of Pyenson et al. (2009), giving 15.2–16.0 Mya. Crown Odontoceti: recovered within Allodelphinidae, and thus within Platanistoidea, by Boersma and Pyenson (2016).

***Ziphiodelphis abeli* Dal Piaz 1908** Holotype from the upper Libano Sandstone of the Belluno quarries (Dal Piaz, 1908), of late Aquitanian–early Burdigalian age, giving 19.0–22.0 Mya. Total group Odontoceti: a eurhinodelphinid (Tosetto et al., 2023).

***Ziphiodelphis sigmoideus* (Pilleri 1985)** Holotype from the upper Libano Sandstone of the Belluno quarries (Pilleri, 1985), its lower bound set by a Burdigalian Pietra da Cantoni skull from Rosignano Monferrato referred to the species (Tosetto et al., 2023), giving 15.98–22.0 Mya. Total group Odontoceti: a eurhinodelphinid (Tosetto et al., 2023).

***Ziphirostrum marginatum* Du Bus 1868** Holotype (IRSNB 3783-M.1878), a rostrum from Antwerp; the species is characteristic of the Deurne Sand Member of the Diest Formation, from which the well-preserved referred skull IRSNB M.1877 derives (Du Bus, 1868; Lambert, 2005c), mid–late Tortonian (Louwye, 2002; Goolaerts et al., 2020), giving 7.6–9.5 Mya. Total group Ziphiidae: placed in the stem "*Messapicetus* clade" by Bianucci et al. (2016) but inside crown Ziphiinae by Lambert (2005c).

***Ziphirostrum recurvus* (Du Bus 1868)** Holotype (IRSNB 3805-M.544), an incomplete rostrum from Antwerp with no recorded type horizon (Du Bus, 1868; Lambert, 2005c); the bracket spans the Antwerpen Sand Member base to the top of the Kattendijk Formation, giving a wide 4.4–15.98 Mya. Total group Ziphiidae: see *Ziphirostrum marginatum*.

***Ziphirostrum turniense* Du Bus 1868** Lectotype (IRSNB 3785-M.539), a partial skull from Antwerp; its type horizon "Crag gris" denotes a type of alteration rather than a stratigraphic level, and similar coloring occurs in the upper Miocene, so the species may be late Miocene or early Pliocene (Lambert, 2005c), the bracket spanning the Diest Formation to the top of the Kattendijk Formation, giving a wide age bracket of 4.4–11.63 Mya. Total group Ziphiidae: see *Ziphirostrum marginatum*.

***Zygiocetus nartorum* Tarasenko 2014** Holotype from Polevoe 1 (Adygea, Russia), Krasnooktyabr'sk Formation, of Middle Sarmatian (Bessarabian) age (Tarasenko, 2014), dated by the Eastern Paratethys Bessarabian chronology of Palcu et al. (2017, 2021) to 9.6–12.1 Mya. Total group Mysticeti.

***Zygophyseter varolai* Bianucci & Landini 2006** Holotype from the Cisterna quarry, upper Pietra Leccese (Salento, Italy), of early Tortonian age (*Neogloboquadrina acostaensis* Zone), giving 8.14–10.5 Mya (Bianucci & Landini, 2006). Stem Physeteroidea: a macroraptorial sperm whale recovered outside the crown by all recent analyses (Bianucci & Landini, 2006; Peri et al., 2022; Lambert et al., 2023).

***Zygorhiza kochii* (Reichenbach 1847)** Holotype (MB.Ma.43248, Museum für Naturkunde Berlin) from the upper Eocene Pachuta Marl Member of the Yazoo Formation, Alabama (Reichenbach, 1847), of Priabonian age (Uhen, 2013; Gingerich, 2015), giving 33.9–37.71 Mya. Stem Cetacea.

**Unsampled extants** Twenty-two extant cetacean species recognized by the Society for Marine Mammalogy (Committee on Taxonomy, 2026) are not sampled in the McGowen et al. (2020) backbone. Two rorquals, *Balaenoptera omurai* Wada et al. 2003 and *B. ricei* Rosel et al. 2021, were assigned to the total group of Balaenopteroidea, as *B. omurai* was shown to be an early-diverging balaenopterid lineage (Sasaki et al., 2006). Four beaked whales fall within crown Ziphiidae: on the McGowen et al. (2020) backbone, *Berardius* is the most basal ziphiid genus and *Hyperoodon* nests inside the ziphiid crown as sister to *Mesoplodon* (McGowen et al., 2020), placing *Berardius arnuxii* Duvernoy 1851, *B. minimus* Yamada et al. 2019 and *Hyperoodon planifrons* Flower 1882 in the crown, while the monotypic *Indopacetus pacificus* (Longman 1926), whose position within Ziphiidae is only weakly resolved (Dalebout et al., 2004), is assigned at the family crown. The three unsampled *Mesoplodon* species (*M. eueu* Carroll et al. 2021, *M. hotaula* Deraniyagala 1963 and *M. traversii* (Gray 1874)) were placed in the total group of *Mesoplodon*, as this genus was shown to be monophyletic (Dalebout et al., 2008). Eight oceanic dolphins (*Cephalorhynchus cruciger* (Quoy & Gaimard 1824), *C. eutropia* Gray 1846, *C. hectori* (Van Beneden 1881), *Sotalia fluviatilis* (Gervais & Deville 1853), *Sousa plumbea* (G. Cuvier 1829), *S. sahulensis* Jefferson & Rosenbaum 2014, *S. teuszii* (Kükenthal 1892) and *Tursiops erebennus* (Cope 1865)) were assigned to crown Delphinidae rather than to a genus crown, as delphinid genera are known to be widely non-monophyletic: e.g., the McGowen et al. (2020) backbone recovers *Stenella* as polyphyletic, with *Tursiops*, *Delphinus* and *Lagenodelphis* nested within it (McGowen et al., 2020). Three porpoises, *Neophocaena asiaeorientalis* Pilleri & Gihr 1972, *Phocoena sinus* Norris & McFarland 1958 and *Phocoenoides dalli* (True 1885), were assigned to the total group of Phocoenidae, the genus *Phocoena* being paraphyletic with respect to *Phocoenoides* (Ben Chehida et al., 2020). Finally, the

South Asian river dolphins *Platanista gangetica* (Lebeck 1801) and *P. minor* Owen 1853 were assigned to crown Odontoceti, since no *Platanista* was sampled, but the deepest-diverging odontocete lineage, Physeteroidea, was (McGowen et al., 2020).

#### 1.2 Emydidae

***Acherontemys heckmani* Hay 1899** Holotype (USNM 4213), an almost complete carapace, from the Roslyn Formation near Roslyn, Kittitas County, Washington (Hay, 1899), whose deposition is constrained by U–Pb zircon geochronology to the middle Eocene (Eddy et al., 2016), giving 45.9–48.8 Mya. Stem Emydidae: placed in an unresolved basal polytomy at the base of the emydid total group by Vlachos (2018).

***Chrysemys corniculata* Jasinski 2022** Holotype (ETMNH-12491), a complete carapace and nearly complete plastron, from the lacustrine sinkhole fill of the Gray Fossil Site, Washington County, Tennessee, a latest Hemphillian–early Blancan local fauna dated by biostratigraphy to 4.5–4.9 Mya (Jasinski, 2022). Total group *Chrysemys*: recovered in a basal position within Deirochelyinae, leaving its position relative to the extant *Chrysemys* unresolved (Jasinski, 2022).

***Chrysemys platymarginata* Weaver & Robertson 1967** Holotype (UF 11046), a partial shell, from Haile 15A, Alachua County, Florida (Weaver & Robertson, 1967), a late Blancan local fauna (Vlachos, 2018), giving 2.2–2.6 Mya. Total group Deirochelyinae: of uncertain generic position within Deirochelyinae (Vlachos, 2018).

***Chrysemys timida* Hay 1908** Holotype (YPM VPPU 010853), a partial carapace, from a river-lag deposit on the Niobrara River, Sheridan County, Nebraska (Hay, 1908), assigned to the Irvingtonian by faunal association (Vlachos, 2018), giving 0.21–1.8 Mya. Total group Emydinae: reassigned from the deirochelyine *Chrysemys* to a basal polytomy within Emydinae by Vlachos (2018).

***Deirochelys carri* Jackson 1978** Holotype (UF 20908), a partial shell, from the Love Bone Bed at the Love Site near Archer, Alachua County, Florida (Jackson, 1978), a latest Clarendonian local fauna (Vlachos, 2018), giving 9.0–9.5 Mya. Total group Deirochelyinae: referred to Deirochelyinae but of uncertain generic position (Vlachos, 2018).

***Emys wermuthi* Młynarski 1956** Holotype from the Pliocene of Wężе, Poland (Młynarski, 1956), dated to the Polish Pliocene (Seidel & Ernst, 2017), giving 2.5–4.5 Mya. Total group *Emys*: of uncertain position within *Emys* relative to the extant *E. orbicularis* (Seidel & Ernst, 2017).

***Glyptemys valentinensis* Holman & Fritz 2001** Holotype (USNM 76564), a partial carapace, from the Crookston Bridge Member of the Valentine Formation at Valentine Railway Quarry A, Cherry County, Nebraska (Holman & Fritz, 2001), a late Barstovian (middle Miocene) unit (Korth, 2008), giving 12.5–13.6 Mya. Total group *Glyptemys*: proposed as the last common ancestor of the two extant *Glyptemys* species but without a corroborating phylogenetic analysis (Vlachos, 2018).

***Graptemys kerneri* Ehret & Bourque 2011** Holotype (UF 239000), a nearly complete skull, from Suwannee River bed sediments at the Gilchrist–Dixie county boundary, Florida, a presumed Rancholabrean occurrence (Ehret & Bourque, 2011), giving 0.0117–0.3 Mya. Crown *Graptemys*: diagnosed as a member of the extant genus *Graptemys* by Vlachos (2018).

***Pseudemys carri* (Rose & Weaver 1966)** Holotype (UF 9427), an almost complete shell, from the McGehee site, Alachua County, Florida (Rose & Weaver, 1966), an early Hemphillian local fauna (Vlachos, 2018), giving 8.0–9.0 Mya. Total group *Pseudemys*.

***Pseudemys idahoensis* Gilmore 1933** Holotype (USNM 12059), an almost complete skeleton, from the Plesippus (Hagerman Horse) Quarry in the Glenns Ferry Formation, Twin Falls County, Idaho (Gilmore, 1933), whose Hagerman beds are dated to 3.11–4.18 Mya (Bernor et al., 2019). Total group Deirochelyinae: recovered in a basal polytomy within Deirochelyinae (Vlachos, 2018).

***Pseudemys williamsi* (Rose & Weaver 1966)** Holotype (UF 11561), an almost complete shell, from the McGehee site, Alachua County, Florida (Rose & Weaver, 1966), an early Hemphillian local fauna (Vlachos, 2018), giving 8.0–9.0 Mya. Total group *Pseudemys*.

***Psilosemys wyomingensis* Hutchison 2013** Holotype (UCMP 179440), carapacial fragments, from the Willwood Formation at Fifteen Mile Creek, Washakie County, Wyoming (Hutchison, 2013), a Graybullian (early Wasatchian, early Eocene) horizon dated by magnetostratigraphic correlation to the geomagnetic polarity time scale (Clyde et al., 1994), giving 53.0–54.8 Mya. Stem Emydidae: regarded as the sister taxon of crown Emydidae by Hutchison (2013) and placed on the emydid stem by Vlachos (2018).

***Terrapene corneri* Holman & Fritz 2005** Holotype (USNM 21618), a near-complete anterior plastral lobe, from the Myers Farm local fauna at USNM locality Wt-15A, Webster County, Nebraska (Holman & Fritz, 2005), a late Barstovian (middle Miocene) fauna (Korth, 2008), giving 12.5–13.6 Mya. Stem *Terrapene*: placed in a basal polytomy along the stem of *Terrapene* by Vlachos (2018).

***Terrapene eurypygia* (Cope 1870)** Holotype (AMNH 1484), carapacial fragments, from cave sediments at Oxford Neck, Talbot County, Maryland (Cope, 1870), poorly dated but assigned to the Irvingtonian on the presence of *Mammuthus* (Vlachos, 2018), giving 0.21–1.8 Mya. Crown *Terrapene*: recovered within crown *Terrapene* in all analyses of Vlachos (2018).

***Terrapene parornata* Joyce et al. 2012** Holotype (OMNH 58158), an almost complete shell, from the Buis Ranch local fauna at Buckshot Arroyo (OMNH locality V671), Beaver County, Oklahoma, a Hemphillian 4 fauna placed at the Miocene/Pliocene boundary (Joyce et al., 2012), giving 4.9–5.33 Mya. Crown *Terrapene*: recovered as sister to the extant *Terrapene ornata* within crown *Terrapene* by Vlachos (2018).

***Terrapene putnami* Hay 1906** Neotype (UF 3066), a complete shell and associated skeleton that formally replaces the undiagnostic holotype (AMNH 6097) of Hay (1906), from the red zone of the Haile 8A deposits, Alachua County, Florida (Ehret et al., 2013), a late Rancholabrean (late Pleistocene) horizon (Vlachos, 2018), giving 0.0117–0.129 Mya. Crown *Terrapene*: recovered within crown *Terrapene* in all analyses of Vlachos (2018).

***Terrapene singletoni* Gilmore 1927** Holotype (USNM 11181) from the Melbourne Bone Bed near Melbourne, Brevard County, Florida (Gilmore, 1927), a terminal-Pleistocene (Rancholabrean) local fauna dated to ∼11–25 ka (Vlachos, 2018), giving 0.011–0.025 Mya. Crown *Terrapene*: recovered within crown *Terrapene* in all analyses of Vlachos (2018).

***Trachemys antiqua* Clark 1937** Holotype (YPM VPPU 013839), an almost complete plastron, from the Chadron Formation 3 km southeast of Cedar Butte, Pennington County, South Dakota (Clark, 1937), a Chadronian (late Eocene) horizon (Vlachos, 2018), giving 33.9–37.0 Mya. Total group Deirochelyinae: the oldest crown emydid, recovered as a basal deirochelyine of uncertain generic position by Vlachos (2018).

***Trachemys haugrudi* Jasinski 2018** Holotype (ETMNH-8549), a near-complete shell and associated skeleton, from the lacustrine sinkhole fill of the Gray Fossil Site, Washington County, Tennessee (Jasinski, 2018), a late Hemphillian deposit whose biostratigraphy dates it to 4.9–4.5 Mya (Jasinski, 2022), giving 4.5–4.9 Mya. Total group *Trachemys*: recovered in a fossil clade of uncertain position relative to the extant *Trachemys* lineages by Jasinski (2018).

***Trachemys sculpta* Hay 1908** Holotype (USNM 16681), a nuchal plate, from a lag over the Peace River Formation, Hillsborough County, Florida (Hay, 1908), a reworked river deposit mixing Miocene and Pleistocene material (Vlachos, 2018), giving 0.0117–5.3 Mya. Total group *Trachemys*: recovered with moderate support as sister to the extant *Trachemys* by Vlachos (2018).

***Wilburemys yakimensis* Martin & Mallory 2011** Holotype (UWBM 58775), a complete carapace and plastron, from a diatomite lens in the upper Ellensburg Formation at the Wilbur 1 Locality, Yakima County, Washington, the diatomite radiometrically dated to 7.41±0.36 Mya (Martin & Mallory, 2011), giving 7.05–7.77 Mya. Total group Emydinae: placed at an ancestral position near the emydine–deirochelyine divergence, closest to the extant *Actinemys*, by Martin and Mallory (2011).

**Unsampled extants** Seven Turtle Turtle Taxonomy Working Group (2025)-valid extant species are not sampled in the Thomson et al. (2021) backbone and were placed within their respective genus. Three box turtles: *Terrapene triunguis* Agassiz 1857, *T. mexicana* Gray 1849 and *T. yucatana* Boulenger 1895, belong to the sampled *T. carolina* group, and thus fall within crown *Terrapene*. *Graptemys sabinensis* Cagle 1953 nests among the non-*geographica* map turtles, and because the deepest-diverging *G. geographica* is sampled (Vogt et al., 2018), it is assigned to the genus crown. Three Mexican sliders: *Trachemys hartwegi* Legler 1990, *T. ornata* Gray 1831 and *T. yaquia* Legler & Webb 1970 were not sampled in the Thomson et al. (2021) tree. Despite being studied only using multigene fragments, multiple studies place them within the sampled extent of *Trachemys* in the backbone tree (Parham et al., 2015; Fritz et al., 2024; Cuadrado-Ríos et al., 2025), and thus were assigned to the crown *Trachemys*.

#### 1.3 Palaeodictyopterida

***Abaptilon sibiricum* Zalessky 1946** Holotype from the Kemerovo Formation (Balakhonka Group) at Prokopyevsk, Kuznetsk Basin (Zalessky, 1946), of early Kungurian age (Silantiev et al., 2025), giving 277.0–283.5 Mya. A Spilapteridae (order Palaeodictyoptera).

***Actinohymen russelli* Carpenter 1962** Holotype from the Petrolia Formation (Wichita Group) at the Emily Irish Land Grant locality, north-central Texas (Carpenter, 1962), of Early Permian (Cisuralian) age (Carpenter, 1962), giving 273.0–298.9 Mya. A Bardohymenidae (order Megasecoptera).

***Adiaphtharsia ferrea* Handlirsch 1906** Holotype from the Francis Creek Shale (Carbondale Formation) at Mazon Creek, Illinois (Handlirsch, 1906b), of Moscovian (Westphalian D) age (Peppers, 1996), giving 306.0–311.0 Mya. A megasecopteran of uncertain family (order Megasecoptera).

***Adolarryia bairdi* Kukalová-Peck and Richardson 1983** Holotype from the Francis Creek Shale (Carbondale Formation) at Benson Farm, Grundy County, Illinois (Kukalová-Peck & Richardson, 1983), of Moscovian (Westphalian D) age (Peppers, 1996), giving 306.0–311.0 Mya. A Homoiopteridae (order Palaeodictyoptera).

***Aenigmatidia kaltanica* Rohdendorf 1961** Holotype from bed 7 of outcrop 1300 in the Mitina Formation (Kolchugino Group) at Kaltan, Kuznetsk Basin (PIN collection 504) (Rohdendorf et al., 1961), of latest Kungurian to Roadian age (Silantiev et al., 2025), giving 266.9–277.0 Mya. A Aenigmatidiidae (order Diaphanopterodea).

***Aenigmatopsis regularis* (Bolton 1921)** Holotype from the Coal Measures at Monckton Main Colliery, Yorkshire, England (Bolton, 1921), of Duckmantian (Westphalian B) age (Gradstein et al., 2020; Schneider et al., 2020), giving 313.7–316.5 Mya. A palaeodictyopteran of uncertain family (order Palaeodictyoptera).

***Agaeoleptoptera uniotempla* Beckemeyer and Engel 2011** Holotype from the Mary Lee coal zone of the Pottsville Formation at the Union Chapel Mine, Alabama (Beckemeyer & Engel, 2011), of Langsettian (Westphalian A) age (Hunt et al., 2005), giving 315.2–319.0 Mya. An Ancopteridae (order Megasecoptera).

***Alectoneura europaea* Kukalová-Peck 1975** Holotype from the upper Letovice Formation at Obora, southern Moravia (Kukalová-Peck, 1975), of Sakmarian–Artinskian (Early Permian) age (Opluštil et al., 2017), giving 283.30–293.52 Mya. An Alectoneuridae (order Megasecoptera).

***Alexahymen aestatis* (Brauckmann 1991)** Holotype from the Flöz Zweibänke at the Piesberg quarry near Osnabrück, Lower Saxony, Germany (Brauckmann, 1991), of Westphalian D (Asturian) age (Gradstein et al., 2020; Schneider et al., 2020), giving 307.50–310.70 Mya. A Bardohymenidae (order Megasecoptera).

***Alexahymen maruska* Kukalová-Peck 1972** Holotype from the upper Letovice Formation at Obora, southern Moravia (Kukalová-Peck, 1972), of Sakmarian–Artinskian (Early Permian) age (Opluštil et al., 2017), giving 283.30–293.52 Mya. A Bardohymenidae (order Megasecoptera).

***Alexrasnitsynia permiana*** **Prokop & Nel 2011** Holotype from the Mérifons Member of the Salagou Formation in the Lodève Basin, southern France (Prokop & Nel, 2011), of late Kungurian age, younger than the 282.86 Mya tuff dated in the underlying Octon Member (Michel et al., 2015), giving 272.95–282.86 Mya. An Alexrasnitsyniidae (order Diaphanopterodea).

***Althansia sahneri*** **Guthörl 1934** Holotype from the Friedrichsthal railway cutting, Saarbrücken district, Saarland, Germany (Guthörl, 1934), of Westphalian age (Gradstein et al., 2020; Schneider et al., 2020), giving 307.5–319.2 Mya. A palaeodictyopteran of uncertain family (order Palaeodictyoptera).

***Ametretus laevis*** **Handlirsch 1911** Holotype from the Francis Creek Shale (Carbondale Formation) at Mazon Creek, Illinois (Handlirsch, 1911), of Moscovian (Westphalian D) age (Peppers, 1996), giving 306.0–311.0 Mya. A Homoiopteridae (order Palaeodictyoptera).

***Amousus mazonus*** **Handlirsch 1911** Holotype from the Francis Creek Shale (Carbondale Formation) at Mazon Creek, Illinois (Handlirsch, 1911), of Moscovian (Westphalian D) age (Peppers, 1996), giving 306.0–311.0 Mya. A Homoiopteridae (order Palaeodictyoptera).

***Anagesthes affinis*** **(Goldenberg 1854)** Holotype from the Friedrichsthal railway cutting, Saarbrücken district, Saarland, Germany (Goldenberg, 1854), of Westphalian age (Gradstein et al., 2020; Schneider et al., 2020), giving 307.5–319.2 Mya. A palaeodictyopteran of uncertain family (order Palaeodictyoptera).

***Anchineura hispanica*** **Carpenter 1963** Holotype (IRSNB 97.587), a forewing, from North of La Magdalena in the La Magdalena Coalfield, León Province, northwestern Spain (Carpenter, 1963), of Gzhelian age (Bashforth et al., 2010; Wagner & Paz Castro, 2011), giving 300.0–304.0 Mya. An Anchineuridae (order Megasecoptera).

***Anconeura havlatai*** **Kukalová-Peck 1975** Holotype from the upper Letovice Formation at Obora, southern Moravia (Kukalová-Peck, 1975), of Sakmarian–Artinskian (Early Permian) age (Opluštil et al., 2017), giving 283.30–293.52 Mya. An Arcioneuridae (order Megasecoptera).

***Ancoptera permiana*** **Kukalová-Peck 1975** Holotype from the upper Letovice Formation at Obora, southern Moravia (Kukalová-Peck, 1975), of Sakmarian–Artinskian (Early Permian) age (Opluštil et al., 2017), giving 283.30–293.52 Mya. An Ancopteridae (order Megasecoptera).

***Anglopterum magnificum* Prokop et al. 2006** Holotype from the Wigan Four-Foot Coal at Crock Hey opencast pit, near Wigan, Lancashire, England (Prokop et al., 2006), of Langsettian (Westphalian A) age (Gradstein et al., 2020; Schneider et al., 2020), giving 316.5–319.2 Mya. A Homoiopteridae (order Palaeodictyoptera).

***Anomalohymen dochmus* Beckemeyer & Engel 2009** Holotype from the Midco Member of the Wellington Formation at the Billings 3 locality, Noble County, Oklahoma (Beckemeyer & Engel, 2009), of Leonardian (late Artinskian–early Kungurian) age (Tasch & Zimmerman, 1962; Schneider et al., 2020), giving 279–286 Mya. A Biarmohymenidae (order Diaphanopterodea).

***Anthohymen hejinensis* Hong 1985** Holotype from the Xiashihezi (Lower Shihezi) Formation at Meidigou, Hejin County, Shanxi Province, China (Hong, 1985b), of Kungurian age (Shen et al., 2022), giving 274.4–283.3 Mya. A Bardohymenidae (order Megasecoptera).

***Anthracosta duboisi* Pruvost 1930** Holotype from the roof of the Sainte-Barbe vein at the Courcelles-Nord coal mine, Courcelles, Hainaut, Belgium (Pruvost, 1930), of Langsettian (Westphalian A) age (Gradstein et al., 2020; Schneider et al., 2020), giving 316.5–319.2 Mya. A palaeodictyopteran of uncertain family (order Palaeodictyoptera).

***Apopappus guernei* (Brongniart 1894)** Holotype from the bituminous shale of the Commentry coal basin at Commentry, Allier, France (Brongniart, 1894), of Gzhelian (Stephanian B) age (Gradstein et al., 2020), giving 298.9–303.7 Mya. A Lycocercidae (order Palaeodictyoptera).

***Archaemegaptilus blakelyi* Beckemeyer and Engel 2018** Holotype from the Pottsville Formation at the Cahaba Living River Retreat, Bibb County, Alabama, of Langsettian (Westphalian A) age (Beckemeyer & Engel, 2018), giving 315.2–319.0 Mya. An Archaemegaptilidae (order Palaeodictyoptera).

***Archaemegaptilus kiefferi* Meunier 1908** Holotype from the Commentry Shales at Commentry, Allier, France (Meunier, 1908a), of Gzhelian (Stephanian B) age (Gradstein et al., 2020), giving 298.9–303.7 Mya. An Archaemegaptilidae (order Palaeodictyoptera).

***Archaemegaptilus schloesseri* Brauckmann et al. 2003** Holotype from the Vorhalle Beds at the Hagen-Vorhalle quarry, North Rhine-Westphalia, Germany (Brauckmann et al., 2003), of late

Marsdenian (Namurian B, *Bilinguites metabilinguis* Marine Band, Ziegelschiefer Formation) age (Ilger & Brauckmann, 2011; Opluštil et al., 2022), giving 319.0–320.5 Mya. An Archaemegaptilidae (order Palaeodictyoptera).

***Archaeophasma grandis* Matthew 1910** Holotype from Plant Bed 2 of the Fern Ledges (Lancaster Formation) at Carleton, Saint John, New Brunswick, Canada (Matthew, 1910), of Langsettian (Westphalian A) age (Gradstein et al., 2020; Schneider et al., 2020), giving 316.5–319.2 Mya. A Lithomanteidae (order Palaeodictyoptera).

***Arcioneura juveniles* Kukalová-Peck 1975** Holotype from the upper Letovice Formation at Obora, southern Moravia (Kukalová-Peck, 1975), of Sakmarian–Artinskian (Early Permian) age (Opluštil et al., 2017), giving 283.30–293.52 Mya. An Arcioneuridae (order Megasecoptera).

***Arltia carnica* Brauckmann & Hahn 1983** Holotype, a wing, from the Corona Formation (Auernig Group) at the Kronalpe (~1650 m, near the Italian border), Carnic Alps, Carinthia, Austria (Brauckmann & Hahn, 1983), of Late Carboniferous (Gzhelian) age (Brauckmann & Hahn, 1983), giving 300.5–303.7 Mya. A Dictyoneuridae (order Palaeodictyoptera).

***Arltia eurycubitalis* Guthörl 1940** Holotype from the Geisheck Formation at the Frankenholz-Saar colliery (shaft 5), Saarland, Germany (Guthörl, 1940), of Westphalian D (Asturian) age (Gradstein et al., 2020; Schneider et al., 2020), giving 307.5–310.7 Mya. A Dictyoneuridae (order Palaeodictyoptera).

***Arltia mioanali* Guthörl 1940** Holotype from the Geisheck Formation at the Frankenholz-Saar colliery (shaft 5), Saarland, Germany (Guthörl, 1940), of Westphalian D (Asturian) age (Gradstein et al., 2020; Schneider et al., 2020), giving 307.5–310.7 Mya. A Dictyoneuridae (order Palaeodictyoptera).

***Arltia obsoleta* (Goldenberg 1877)** Holotype from the Lower Saarbrücken layers at the Altenwald mine, Saarland, Germany (Goldenberg, 1877), of Westphalian C (Bolsovian) age (Gradstein et al., 2020; Schneider et al., 2020), giving 310.7–313.7 Mya. A Dictyoneuridae (order Palaeodictyoptera).

***Arltia schmitzi* (Goldenberg 1877)** Holotype from the Lower Saarbrücken layers at the Altenwald mine, Saarland, Germany (Goldenberg, 1877), of Westphalian C (Bolsovian) age (Gradstein et al., 2020; Schneider et al., 2020), giving 310.7–313.7 Mya. A Dictyoneuridae (order Palaeodictyoptera).

***Arltia willerti* (Guthörl 1930)** Holotype from the Friedrichsthal railway cutting, Saarbrücken district, Saarland, Germany (Guthörl, 1930), of Westphalian age (Gradstein et al., 2020; Schneider et al., 2020), giving 307.5–319.2 Mya. A Dictyoneuridae (order Palaeodictyoptera).

***Arroyohymen splendens* Prokop and Kukalová-Peck 2017** Holotype from the Red Tanks Member of the Bursum Formation at Carrizo Arroyo, New Mexico (Prokop & Kukalová-Peck, 2017), of latest Gzhelian–earliest Asselian (Carboniferous–Permian boundary) age (Schneider et al., 2020), giving 296.5–299.5 Mya. A Protohymenidae (order Megasecoptera).

***Asiodictya rossica* Rohdendorf 1961** Holotype from the Alykaeva Formation at Zavyalovo, Kuznetsk Basin (PIN collection 742) (Rohdendorf et al., 1961), of Gzhelian age (Silantiev et al., 2025), giving 298.9–303.7 Mya. A palaeodictyopteran of uncertain family (order Palaeodictyoptera).

***Aspidohymen chopardi* Zalessky 1937** Holotype from the Baitugan Beds at Tikhie Gory, Volga region (Zalessky, 1937b), of early Kazanian (Roadian) age (Davydov et al., 2018), giving 266.9–277.0 Mya. A Aspidohymenidae (order Megasecoptera).

***Aspidohymen extensus* Martynov 1930** Holotype from the Baitugan Beds at Tikhie Gory, Volga region (PIN collection 2295) (Martynov, 1930a), of early Kazanian (Roadian) age (Davydov et al., 2018), giving 266.9–277.0 Mya. A Aspidohymenidae (order Megasecoptera).

***Aspidothorax hispanicus* Santos et al. 2025** Holotype from an outcrop near the Carmen mine in the La Magdalena Coalfield, León Province, northwestern Spain (Santos et al., 2025), of Gzhelian age (Bashforth et al., 2010; Wagner & Paz Castro, 2011), giving 300.0–304.0 Mya. An Aspidothoracidae (order Megasecoptera).

***Aspidothorax permianus* Sinitshenkova et al. 2020** Holotype from the Klyuchiki locality, Perm Region (PGNIU collection 6571) (Sinitshenkova et al., 2020), of Kazanian (Roadian) age (Davydov et al., 2018), giving 266.9–277.0 Mya. A Aspidothoracidae (order Megasecoptera).

***Aspidothorax triangularis* Brongniart 1894** Holotype from the bituminous shale of the Commentry coal basin at Commentry, Allier, France (Brongniart, 1894), of Gzhelian (Stephanian B) age (Gradstein et al., 2020), giving 298.9–303.7 Mya. An Aspidothoracidae (order Megasecoptera).

***Asthenohymen affinis* (Tillyard 1926)** Holotype from the Wellington Formation at Elmo, Kansas (Tillyard, 1926), of Leonardian (late Artinskian–early Kungurian) age (Tasch & Zimmerman, 1962;

Schneider et al., 2020), giving 279–286 Mya. An Asthenohymenidae (order Diaphanopterodea).

***Asthenohymen apicalis* Carpenter 1947** Holotype from the Midco Member of the Wellington Formation at Midco locality 15, Oklahoma (Carpenter, 1947), of Leonardian (late Artinskian–early Kungurian) age (Tasch & Zimmerman, 1962; Schneider et al., 2020), giving 279–286 Mya. An Asthenohymenidae (order Diaphanopterodea).

***Asthenohymen dunbari* Tillyard 1924** Holotype from the Wellington Formation at Elmo, Kansas (Tillyard, 1924), of Leonardian (late Artinskian–early Kungurian) age (Tasch & Zimmerman, 1962; Schneider et al., 2020), giving 279–286 Mya. An Asthenohymenidae (order Diaphanopterodea).

***Asthenohymen gracilis* Tillyard 1926** Holotype from the Wellington Formation at Elmo, Kansas (Tillyard, 1926), of Leonardian (late Artinskian–early Kungurian) age (Tasch & Zimmerman, 1962; Schneider et al., 2020), giving 279–286 Mya. An Asthenohymenidae (order Diaphanopterodea).

***Asthenohymen kansasensis* Tillyard 1926** Holotype from the Wellington Formation at Elmo, Kansas (Tillyard, 1926), of Leonardian (late Artinskian–early Kungurian) age (Tasch & Zimmerman, 1962; Schneider et al., 2020), giving 279–286 Mya. An Asthenohymenidae (order Diaphanopterodea).

***Asthenohymen latus* Carpenter 1947** Holotype from the Midco Member of the Wellington Formation at Midco locality 15, Oklahoma (Carpenter, 1947), of Leonardian (late Artinskian–early Kungurian) age (Tasch & Zimmerman, 1962; Schneider et al., 2020), giving 279–286 Mya. An Asthenohymenidae (order Diaphanopterodea).

***Asthenohymen micron* Sinitshenkova 2004** Holotype from the Red Tanks Member of the Bursum Formation at Carrizo Arroyo, New Mexico (Rasnitsyn et al., 2004), of latest Gzhelian–earliest Asselian (Carboniferous–Permian boundary) age (Schneider et al., 2020), giving 296.5–299.5 Mya. An Asthenohymenidae (order Diaphanopterodea).

***Asthenohymen minutum* Zalessky 1939** Holotype from the Koshelevka Formation in an outcrop above Yulaevo on the Sylva River, Perm region, Russia (Zalessky, 1939), of Kungurian (Irenian substage) age (Lucas & Shen, 2018; Khramov et al., 2023), giving 273.0–283.5 Mya. An Asthenohymenidae (order Diaphanopterodea).

***Asthenohymen minutus* Zimmerman 1962** Holotype from the *Asthenohymen–Delopterum* bed of the Wellington Formation at Tasch's Nobel V locality (bed 7), Oklahoma, of Leonardian (late

Artinskian–early Kungurian) age (Tasch & Zimmerman, 1962; Schneider et al., 2020), giving 279–286 Mya. An Asthenohymenidae (order Diaphanopterodea).

***Asthenohymen parvulus* Carpenter 1947** Holotype from the Midco Member of the Wellington Formation at Midco locality 15, Oklahoma (Carpenter, 1947), of Leonardian (late Artinskian–early Kungurian) age (Tasch & Zimmerman, 1962; Schneider et al., 2020), giving 279–286 Mya. An Asthenohymenidae (order Diaphanopterodea).

***Asthenohymen pusillus* (Tillyard 1926)** Holotype from the Wellington Formation at Elmo, Kansas (Tillyard, 1926), of Leonardian (late Artinskian–early Kungurian) age (Tasch & Zimmerman, 1962; Schneider et al., 2020), giving 279–286 Mya. An Asthenohymenidae (order Diaphanopterodea).

***Asthenohymen stenobasis* Tillyard 1926** Holotype from the Wellington Formation at Elmo, Kansas (Tillyard, 1926), of Leonardian (late Artinskian–early Kungurian) age (Tasch & Zimmerman, 1962; Schneider et al., 2020), giving 279–286 Mya. An Asthenohymenidae (order Diaphanopterodea).

***Asthenohymen stigmatizans* Tillyard 1926** Holotype from the Wellington Formation at Elmo, Kansas (Tillyard, 1926), of Leonardian (late Artinskian–early Kungurian) age (Tasch & Zimmerman, 1962; Schneider et al., 2020), giving 279–286 Mya. An Asthenohymenidae (order Diaphanopterodea).

***Asthenohymen triangularis* Carpenter 1947** Holotype from the Midco Member of the Wellington Formation at Midco locality 15, Oklahoma (Carpenter, 1947), of Leonardian (late Artinskian–early Kungurian) age (Tasch & Zimmerman, 1962; Schneider et al., 2020), giving 279–286 Mya. An Asthenohymenidae (order Diaphanopterodea).

***Asthenohymen uralicum* Zalessky 1939** Holotype from the Koshelevka Formation in an outcrop above Yulaevo on the Sylva River, Perm region, Russia (Zalessky, 1939), of Kungurian (Irenian substage) age (Lucas & Shen, 2018; Khramov et al., 2023), giving 273.0–283.5 Mya. An Asthenohymenidae (order Diaphanopterodea).

***Asthenohymen zonatus* Sinitshenkova 2004** Holotype from the Red Tanks Member of the Bursum Formation at Carrizo Arroyo, New Mexico (Rasnitsyn et al., 2004), of latest Gzhelian–earliest Asselian (Carboniferous–Permian boundary) age (Schneider et al., 2020), giving 296.5–299.5 Mya. An Asthenohymenidae (order Diaphanopterodea).

***Aviobreyeria gracilis* Prokop et al. 2013** Holotype from the spoil heap of Terril No. 7 at Avion, Pas-de-Calais, France (Prokop et al., 2013), of Bolsovian (Westphalian C) age (Gradstein et al., 2020; Schneider et al., 2020), giving 310.7–313.7 Mya. A Breyeriidae (order Palaeodictyoptera).

***Avionbrodia incompleta* Schubnel et al. 2020** Holotype from the spoil heap of Terril No. 7 at Avion, Pas-de-Calais, France (Schubnel et al., 2020), of Bolsovian (Westphalian C) age (Gradstein et al., 2020; Schneider et al., 2020), giving 310.7–313.7 Mya. A diaphanopterodean of uncertain family (order Diaphanopterodea).

***Aykhal helenae* Sinitshenkova 1993** Holotype from borehole 4756 at Aykhal on the Markha River, Yakutia (PIN collection 3838) (Sinitshenkova, 1993), of Gzhelian age (Budnikov et al., 2020), giving 298.9–303.7 Mya. A Aykhalidae (order Megasecoptera).

***Baeoneura desperata* Sinitshenkova 2020** Holotype from a depth of 626.0 m in borehole K-1304 of the Vorkuta Field, in the Inta Formation of the Pechora (Vorkuta) Basin (Sinitshenkova, 2020), of Kungurian age (Davydov et al., 2018), giving 273.0–283.5 Mya. A Spilapteridae (order Palaeodictyoptera).

***Baeoneura obscura* Sinitshenkova 1977** Holotype from the Kata Formation on the Chunya River, Tunguska Basin (PIN collection 3115) (Sharov & Sinitshenkova, 1977), of Gzhelian age (Budnikov et al., 2020), giving 298.9–303.7 Mya. A Spilapteridae (order Palaeodictyoptera).

***Bardohymen magnipennifer* Zalessky 1937** Holotype from the Permian of the Sylva River basin, Perm region, Russia (Zalessky, 1937a), of Kungurian (Irenian substage) age (Lucas & Shen, 2018; Khramov et al., 2023), giving 273.0–283.5 Mya. A Bardohymenidae (order Megasecoptera).

***Bathytaptus falcipennis* Handlirsch 1906** Holotype from the Pottsville Formation at Coalburg, near Birmingham, Alabama (Handlirsch, 1906b), of Westphalian A–B age (Beckemeyer & Engel, 2011), giving 313.0–318.7 Mya. A palaeodictyopteran of uncertain family (order Palaeodictyoptera).

***Becquerelia elegans* (Brongniart 1894)** Holotype from the bituminous shale of the Commentry coal basin at Commentry, Allier, France (Brongniart, 1894), of Gzhelian (Stephanian B) age (Gradstein et al., 2020), giving 298.9–303.7 Mya. A Spilapteridae (order Palaeodictyoptera).

***Becquerelia superba* Brongniart 1894** Holotype from the bituminous shale of the Commentry coal basin at Commentry, Allier, France (Brongniart, 1894), of Gzhelian (Stephanian B) age (Gradstein et al., 2020), giving 298.9–303.7 Mya. A Spilapteridae (order Palaeodictyoptera).

***Becquerelia tincta* (Brongniart 1894)** Holotype from the bituminous shale of the Commentry coal basin at Commentry, Allier, France (Brongniart, 1894), of Gzhelian (Stephanian B) age (Gradstein et al., 2020), giving 298.9–303.7 Mya. A Spilapteridae (order Palaeodictyoptera).

***Biarmohymen bardense* Zalessky 1937** Holotype from the Permian of the Sylva River basin, Perm region, Russia (Zalessky, 1937a), of Kungurian (Irenian substage) age (Lucas & Shen, 2018; Khramov et al., 2023), giving 273.0–283.5 Mya. A Biarmohymenidae (order Diaphanopterodea).

***Bizarrea obscura* Prokop et al. 2016** Holotype from the Francis Creek Shale (Carbondale Formation) at Mazon Creek, Illinois (Prokop et al., 2016), of Moscovian (Westphalian D) age (Peppers, 1996), giving 306.0–311.0 Mya. A Spilapteridae (order Palaeodictyoptera).

***Bojoptera colorata* Kukalová 1958** Holotype from the Radnice Member of the Kladno Formation at Rakovník (Rakonitz), Bohemia (Czech Republic) (Kukalová, 1958), of Moscovian age (Gradstein et al., 2020), giving 307.0–315.2 Mya. A palaeodictyopteran of uncertain family (order Palaeodictyoptera).

***Boltonia robusta* (Pruvost 1919)** Holotype from the Thiers pit spoil heap of the Anzin mines at Anzin, Nord, France (Pruvost, 1919), of Duckmantian (Westphalian B) age (Gradstein et al., 2020; Schneider et al., 2020), giving 313.7–316.5 Mya. A Homoiopteridae (order Palaeodictyoptera).

***Boltoniella uniformis* Bolton 1934** Holotype from the Coal Measures at Ystradgynlais Colliery, South Wales (Bolton, 1934), of Duckmantian (Westphalian B) age (Gradstein et al., 2020; Schneider et al., 2020), giving 313.7–316.5 Mya. A Spilapteridae (order Palaeodictyoptera).

***Boskovicia celtai* Dostál & Prokop 2009** Holotype from the upper Letovice Formation at Obora, southern Moravia (Dostál & Prokop, 2009), of Sakmarian–Artinskian (Early Permian) age (Opluštil et al., 2017), giving 283.30–293.52 Mya. A Martynoviidae (order Diaphanopterodea).

***Breyeria barborae* Kukalová 1959** Holotype from the roof of coal seam 31 at the Barbora mine, Karviná, Upper Silesian Coal Basin, Czech Republic (Kukalová, 1959), of Westphalian A (Langsettian) age (Gradstein et al., 2020; Schneider et al., 2020), giving 316.50–319.20 Mya. A Breyeriidae (order Palaeodictyoptera).

***Breyeria bistrata* Brauckmann 1995** Holotype from the Flöz Zweibänke at the Piesberg quarry near Osnabrück, Lower Saxony, Germany (Brauckmann, 1995), of Westphalian D (Asturian) age (Gradstein et al., 2020; Schneider et al., 2020), giving 307.50–310.70 Mya. A Breyeriidae (order Palaeodictyoptera).

***Breyeria borinensis* (de Borre 1875)** Holotype from the Flénu Formation at the Belle-et-Bonne colliery, Jemappes, Hainaut, Belgium (Preudhomme de Borre, 1875), of Bolsovian (Westphalian C) age (Gradstein et al., 2020; Schneider et al., 2020), giving 310.7–313.7 Mya. A Breyeriidae (order Palaeodictyoptera).

***Breyeria boulei* (Meunier 1910)** Holotype from the Commentry Shales at Commentry, Allier, France (Meunier, 1910), of Gzhelian (Stephanian B) age (Gradstein et al., 2020), giving 298.9–303.7 Mya. A Breyeriidae (order Palaeodictyoptera).

***Breyeria brauckmanni* Pinto 1992** Holotype from profile NFC, bed A, La Casilda section, Piedra Shotel Formation, Chubut Province, Argentina (Pinto, 1992), of Cisuralian (Sakmarian–Artinskian) age (Archangelsky et al., 1981), giving 283.5–293.5 Mya. A Breyeriidae (order Palaeodictyoptera).

***Breyeria delruei* (Laurentiaux 1949)** Holotype from the spoil heap of pit 11 at Noeux-les-Mines (Verquigneul), Pas-de-Calais, France (Laurentiaux, 1949b), of Duckmantian (Westphalian B) age (Gradstein et al., 2020; Schneider et al., 2020), giving 313.7–316.5 Mya. A Breyeriidae (order Palaeodictyoptera).

***Breyeria janseni* (Zessin 2006)** Holotype from the Flöz Dreibänke at the Piesberg quarry near Osnabrück, Lower Saxony, Germany (Zessin, 2006), of Westphalian D (Asturian) age (Gradstein et al., 2020; Schneider et al., 2020), giving 307.50–310.70 Mya. A Breyeriidae (order Palaeodictyoptera).

***Breyeria lachlani* (Brongniart 1894)** Holotype from the Commentry Shales at Commentry, Allier, France (Brongniart, 1894), of Gzhelian (Stephanian B) age (Gradstein et al., 2020), giving 298.9–303.7 Mya. A Breyeriidae (order Palaeodictyoptera).

***Breyeria limburgica* Laurentiaux 1950** Holotype from the roof of vein A at the Julia Mine, South Limburg, Netherlands (Laurentiaux, 1950), of Westphalian A (Langsettian) age (Gradstein et al., 2020; Schneider et al., 2020), giving 316.5–319.2 Mya. A Breyeriidae (order Palaeodictyoptera).

***Breyeria rappi* Carpenter 1967** Holotype (a forewing) from the Pewee Coal at Peach Tree Gap, Anderson County, Tennessee (Carpenter, 1967), of late Kasimovian age (Schneider et al., 2020), giving 303.9–306.0 Mya. A Breyeriidae (order Palaeodictyoptera).

***Breyeria sharovi* (Sinitshenkova 1979)** Holotype from the Kata Formation on the Paymbu River, Tunguska Basin (PIN collection 2293) (Sinitshenkova, 1979), of Gzhelian age (Budnikov et al., 2020), giving 298.9–303.7 Mya. A Breyeriidae (order Palaeodictyoptera).

***Breyeria solida* Brauckmann & Herd 2000** Holotype from the Flöz Zweibänke at the Piesberg quarry near Osnabrück, Lower Saxony, Germany (Brauckmann & Herd, 2000), of Westphalian D (Asturian) age (Gradstein et al., 2020; Schneider et al., 2020), giving 307.50–310.70 Mya. A Breyeriidae (order Palaeodictyoptera).

***Breyeria triramosa* (Sinitshenkova 1979)** Holotype from the Kata Formation on the Vilyuy River, Tunguska Basin (PIN collection 3628) (Sinitshenkova, 1979), of Gzhelian age (Budnikov et al., 2020), giving 298.9–303.7 Mya. A Breyeriidae (order Palaeodictyoptera).

***Breyeria vrankeni* Laurentiaux-Vieira & Laurentiaux 1964** Holotype from the Mauritz mine, Ruhr, Germany (Laurentiaux-Vieira & Laurentiaux, 1964), of Westphalian B (Duckmantian) age (Gradstein et al., 2020; Schneider et al., 2020), giving 313.7–316.5 Mya. A Breyeriidae (order Palaeodictyoptera).

***Brodia jogginsensis* Prokop et al. 2017** Holotype from the Joggins Formation at Joggins, Nova Scotia (Prokop et al., 2017), of Langsettian (Westphalian A) age (Davies et al., 2005), giving 315.0–318.7 Mya. A Brodiidae (order Megasecoptera).

***Brodia priscotincta* Scudder 1881** Holotype from the Coal Measures at Coseley, near Dudley, South Staffordshire, England (Scudder, 1881), of Duckmantian (Westphalian B) age (Gradstein et al., 2020; Schneider et al., 2020), giving 313.7–316.5 Mya. A Brodiidae (order Megasecoptera).

***Brodioptera cumberlandensis* Copeland 1957** Holotype from the Joggins Formation at West Bay, Parrsboro, Nova Scotia (Copeland, 1957), of Langsettian (Westphalian A) age (Davies et al., 2005), giving 315.0–318.7 Mya. A Brodiopteridae (order Megasecoptera).

***Brodioptera pintoi* (Brauckmann et al. 2003)** Holotype from the Vorhalle Beds at the Hagen-Vorhalle quarry, North Rhine-Westphalia, Germany (Brauckmann et al., 2003), of late Marsdenian

(Namurian B, *Bilinguites metabilinguis* Marine Band, Ziegelschiefer Formation) age (Ilger & Brauckmann, 2011; Opluštil et al., 2022), giving 319.0–320.5 Mya. A Brodiopteridae (order Megasecoptera).

***Brodioptera sinensis* Pecharová et al. 2015** Holotype from the Yanghugou Formation at Xiaheyan, Zhongwei, Ningxia, China (Pecharová et al., 2015b), of latest Bashkirian to middle Moscovian age (Trümper et al., 2020), giving 309.0–316.0 Mya. A Brodiopteridae (order Megasecoptera).

***Brodioptera stricklani* Nelson and Tidwell 1987** Holotype from the upper Manning Canyon Shale at the Manning Canyon clay pits, Utah, of earliest Bashkirian (Kinderscoutian) age (Nelson & Tidwell, 1987), giving 320.5–322.5 Mya. A Brodiopteridae (order Megasecoptera).

***Brongniartula rastrata* Handlirsch 1919** Holotype from the Commentry Shales at Commentry, Allier, France (Handlirsch, 1919), of Gzhelian (Stephanian B) age (Gradstein et al., 2020), giving 298.9–303.7 Mya. A palaeodictyopteran of uncertain family (order Palaeodictyoptera).

***Calohymen permianus* Carpenter 1947** Holotype from the Midco Member of the Wellington Formation at Midco locality 15, Oklahoma (Carpenter, 1947), of Leonardian (late Artinskian–early Kungurian) age (Tasch & Zimmerman, 1962; Schneider et al., 2020), giving 279–286 Mya. A Bardohymenidae (order Megasecoptera).

***Calvertiella permiana* Tillyard 1925** Holotype from the Wellington Formation at Elmo, Kansas (Tillyard, 1925), of Leonardian (late Artinskian–early Kungurian) age (Tasch & Zimmerman, 1962; Schneider et al., 2020), giving 279–286 Mya. A Calvertiellidae (order Palaeodictyoptera).

***Camptodiapha atkinsoni* Beckemeyer and Engel 2011** Holotype from the Mary Lee coal zone of the Pottsville Formation at the Union Chapel Mine, Alabama (Beckemeyer & Engel, 2011), of Langsettian (Westphalian A) age (Hunt et al., 2005), giving 315.2–319.0 Mya. A Namurodiaphidae (order Diaphanopterodea).

***Carbohymen testai* Pecharová and Prokop 2018** Holotype from the Francis Creek Shale (Carbondale Formation) at Pit 11, Mazon Creek, Illinois (Pecharová & Prokop, 2018), of Moscovian (Westphalian D) age (Peppers, 1996), giving 306.0–311.0 Mya. A Protohymenidae (order Megasecoptera).

***Carrizodiaphanoptera permiana* Prokop and Kukalová-Peck 2017** Holotype from the Red Tanks Member of the Bursum Formation at Carrizo Arroyo, New Mexico (Prokop & Kukalová-Peck,

2017), of latest Gzhelian–earliest Asselian (Carboniferous–Permian boundary) age (Schneider et al., 2020), giving 296.5–299.5 Mya. A Diaphanopteridae (order Diaphanopterodea).

***Carrizopteryx arroyo* Kukalová-Peck 1976** Holotype from the Red Tanks Member of the Bursum Formation at Carrizo Arroyo, New Mexico (Kukalová-Peck & Peck, 1976), of latest Gzhelian–earliest Asselian (Carboniferous–Permian boundary) age (Schneider et al., 2020), giving 296.5–299.5 Mya. A Calvertiellidae (order Palaeodictyoptera).

***Carrizopteryx lucasi* Sinitshenkova 2004** Holotype from the Red Tanks Member of the Bursum Formation at Carrizo Arroyo, New Mexico (Rasnitsyn et al., 2004), of latest Gzhelian–earliest Asselian (Carboniferous–Permian boundary) age (Schneider et al., 2020), giving 296.5–299.5 Mya. A Calvertiellidae (order Palaeodictyoptera).

***Catadyesthus priscus* (Andrée 1864)** Holotype from the Upper Carboniferous of Stradonitz (Stradonice), Bohemia (Czech Republic) (Andrée, 1864), of Moscovian age (Gradstein et al., 2020), giving 307.0–315.2 Mya. A Lithomanteidae (order Palaeodictyoptera).

***Cauloptera colorata* Kukalová-Peck 1975** Holotype from the upper Letovice Formation at Obora, southern Moravia (Kukalová-Peck, 1975), of Sakmarian–Artinskian (Early Permian) age (Opluštil et al., 2017), giving 283.30–293.52 Mya. A Caulopteridae (order Megasecoptera).

***Cleffia palatina* (Guthörl 1940)** Holotype from the Geisheck Formation at the Frankenholz-Saar colliery (shaft 5), Saarland, Germany (Guthörl, 1940), of Westphalian D (Asturian) age (Gradstein et al., 2020; Schneider et al., 2020), giving 307.5–310.7 Mya. A Dictyoneuridae (order Palaeodictyoptera).

***Cleffia sarana* Guthörl 1931** Holotype from the Sulzbach Beds at the Grube Friedrichsthal-Saar (Erkershöhe shaft), Saarbrücken, Saarland, Germany (Guthörl, 1931), of Westphalian C (Bolsovian) age (Gradstein et al., 2020; Schneider et al., 2020), giving 310.7–313.7 Mya. A Dictyoneuridae (order Palaeodictyoptera).

***Cockerelliella sepulta* Meunier 1910** Holotype from the bituminous shale of the Commentry coal basin at Commentry, Allier, France (Meunier, 1910), of Gzhelian (Stephanian B) age (Gradstein et al., 2020), giving 298.9–303.7 Mya. An Eugereonidae (order Palaeodictyoptera).

***Compsoneura formosa* Brongniart 1894** Holotype from the bituminous shale of the Commentry coal basin at Commentry, Allier, France (Brongniart, 1894), of Gzhelian (Stephanian B) age (Gradstein et al., 2020), giving 298.9–303.7 Mya. A palaeodictyopteran of uncertain family (order Palaeodictyoptera).

***Compsoneura fusca* (Brongniart 1885)** Holotype from the bituminous shale of the Commentry coal basin at Commentry, Allier, France (Brongniart, 1885), of Gzhelian (Stephanian B) age (Gradstein et al., 2020), giving 298.9–303.7 Mya. A palaeodictyopteran of uncertain family (order Palaeodictyoptera).

***Corydaloides scudderi* Brongniart 1885** Holotype from the Commentry Shales at Commentry, Allier, France (Brongniart, 1885), of Gzhelian (Stephanian B) age (Gradstein et al., 2020), giving 298.9–303.7 Mya. A Corydaloididae (order Megasecoptera).

***Cryptovenia moyseyi* Bolton 1912** Holotype from the Middle Coal Measures at Shipley, Derbyshire, England (Bolton, 1912), of Duckmantian (Westphalian B) age (Gradstein et al., 2020; Schneider et al., 2020), giving 313.7–316.5 Mya. A Cryptoveniidae (order Palaeodictyoptera).

***Cyclocelis chatini* (Brongniart 1894)** Holotype from the Commentry Shales at Commentry, Allier, France (Brongniart, 1894), of Gzhelian (Stephanian B) age (Gradstein et al., 2020), giving 298.9–303.7 Mya. A Sphecopteridae (order Megasecoptera).

***Cyclocelis devexa* Sinitshenkova 2019** Holotype from a depth of 111.3 m in borehole VK-1571 at Upper Syr-Yaga, in the Rudnitskaya Member of the Lekvorkuta Formation of the Pechora (Vorkuta) Basin (Sinitshenkova, 2019), of Kungurian age (Davydov et al., 2018), giving 273.0–283.5 Mya. A Sphecopteridae (order Megasecoptera).

***Cyclocelis elegantissima* (Meunier 1908)** Holotype from the Commentry Shales at Commentry, Allier, France (Meunier, 1908c), of Gzhelian (Stephanian B) age (Gradstein et al., 2020), giving 298.9–303.7 Mya. A Sphecopteridae (order Megasecoptera).

***Cyclocelis obscura* Brongniart 1894** Holotype from the Commentry Shales at Commentry, Allier, France (Brongniart, 1894), of Gzhelian (Stephanian B) age (Gradstein et al., 2020), giving 298.9–303.7 Mya. A Sphecopteridae (order Megasecoptera).

***Delafondia granulata* Langiaux & Parriat 1975** Holotype from La Sorme at Montceau-les-Mines, Saône-et-Loire, France (Langiaux & Parriat, 1975), of Duckmantian (Westphalian B) age

(Gradstein et al., 2020; Schneider et al., 2020), giving 313.7–316.5 Mya. A palaeodictyopteran of uncertain family (order Palaeodictyoptera).

***Delitzschala bitterfeldensis* Brauckmann & Schneider 1996** Holotype from a borehole (WIS-BAW 1315, 517.6–517.78 m) in the Bitterfeld/Delitzsch area, Germany (Brauckmann & Schneider, 1996), of Serpukhovian (Arnsbergian) age (Gradstein et al., 2020), giving 323.6–326.9 Mya. A Spilapteridae (order Palaeodictyoptera).

***Diapha candida* Kukalová-Peck 1974** Holotype from the upper Letovice Formation at Obora, southern Moravia (Kukalová-Peck, 1974), of Sakmarian–Artinskian (Early Permian) age (Opluštil et al., 2017), giving 283.30–293.52 Mya. A diaphanopterodean of uncertain family (order Diaphanopterodea).

***Diaphanoptera munieri* Brongniart 1894** Holotype from the Commentry Shales at Commentry, Allier, France (Brongniart, 1894), of Gzhelian (Stephanian B) age (Gradstein et al., 2020), giving 298.9–303.7 Mya. A Diaphanopteridae (order Diaphanopterodea).

***Diaphanoptera vetusta* Brongniart 1894** Holotype from the Commentry Shales at Commentry, Allier, France (Brongniart, 1894), of Gzhelian (Stephanian B) age (Gradstein et al., 2020), giving 298.9–303.7 Mya. A Diaphanopteridae (order Diaphanopterodea).

***Diaphpterum rasnitsyni* Pinto 1999** Holotype from the Pallero Member of the Bajo de Véliz Formation at Bajo de Véliz, San Luis Province, Argentina (Pinto, 1999), of Gzhelian–Asselian age (Césari & Chiesa, 2017), giving 295.0–303.7 Mya. A Diaphanopteridae (order Diaphanopterodea).

***Diathema concinnum* Sinitshenkova 1980** Holotype from the Koshelevka Formation at Chekarda on the Sylva River, Perm region, Russia (Sinitshenkova, 1980), of Kungurian (Irenian substage) age (Lucas & Shen, 2018; Khramov et al., 2023), giving 273.0–283.5 Mya. A Diathemidae (order Permothemistida).

***Diathema tenerum* Sinitshenkova 1980** Holotype from the Koshelevka Formation at Chekarda on the Sylva River, Perm region, Russia (Sinitshenkova, 1980), of Kungurian (Irenian substage) age (Lucas & Shen, 2018; Khramov et al., 2023), giving 273.0–283.5 Mya. A Diathemidae (order Permothemistida).

***Diathemidia monstruosa* Sinitshenkova 1980** Holotype from the Koshelevka Formation at Chekarda on the Sylva River, Perm region, Russia (Sinitshenkova, 1980), of Kungurian (Irenian substage) age (Lucas & Shen, 2018; Khramov et al., 2023), giving 273.0–283.5 Mya. A Diathemidae (order Permothemistida).

***Dictyoneura higginsii* (Handlirsch 1906)** Holotype from the Middle Coal Measures at the Ravenhead railway cutting, near St Helens, Lancashire, England (Handlirsch, 1906a), of Langsettian (Westphalian A) age (Gradstein et al., 2020; Schneider et al., 2020), giving 316.5–319.2 Mya. A Dictyoneuridae (order Palaeodictyoptera).

***Dictyoneura kemperi* Brauckmann & Koch 1983** Holotype from the Vorhalle Beds at the Hagen-Vorhalle quarry, North Rhine-Westphalia, Germany (Brauckmann & Koch, 1983), of late Marsdenian (Namurian B, *Bilinguites metabilinguis* Marine Band, Ziegelschiefer Formation) age (Ilger & Brauckmann, 2011; Opluštil et al., 2022), giving 319.0–320.5 Mya. A Dictyoneuridae (order Palaeodictyoptera).

***Dictyoneura libelluloides* (Goldenberg 1854)** Holotype from the Auerswald coal seam at Gersweiler (Saarbrücken), Saarland, Germany (Goldenberg, 1854), of Westphalian D (Asturian) age (Gradstein et al., 2020; Schneider et al., 2020), giving 307.5–310.7 Mya. A Dictyoneuridae (order Palaeodictyoptera).

***Dictyoneura nigra* (Kliver 1883)** Holotype from the Lower Saarbrücken layers at Frankenholz, Saarland, Germany (Kliver, 1883), of Westphalian C (Bolsovian) age (Gradstein et al., 2020; Schneider et al., 2020), giving 310.7–313.7 Mya. A Dictyoneuridae (order Palaeodictyoptera).

***Dictyoneurella perfecta* Laurentiaux 1949** Holotype from the pit 9 spoil heap at Béthune, Pas-de-Calais, France (Laurentiaux, 1949a), of Duckmantian (Westphalian B) age (Gradstein et al., 2020; Schneider et al., 2020), giving 313.7–316.5 Mya. A Dictyoneurellidae (order Megasecoptera).

***Dictyoneurula gracilis* (Kliver 1886)** Holotype from the Friedrichsthal railway cutting, Saarbrücken district, Saarland, Germany (Kliver, 1886), of Westphalian age (Gradstein et al., 2020; Schneider et al., 2020), giving 307.5–319.2 Mya. A Dictyoneuridae (order Palaeodictyoptera).

***Dictyoptilus peromapteroides* (Meunier 1908)** Holotype from the bituminous shale of the Commentry coal basin at Commentry, Allier, France (Meunier, 1908e), of Gzhelian (Stephanian B) age (Gradstein et al., 2020), giving 298.9–303.7 Mya. An Eugereonidae (order Palaeodictyoptera).

***Dictyoptilus renaulti* Brongniart 1894** Holotype from the bituminous shale of the Commentry coal basin at Commentry, Allier, France (Brongniart, 1894), of Gzhelian (Stephanian B) age (Gradstein et al., 2020), giving 298.9–303.7 Mya. An Eugereonidae (order Palaeodictyoptera).

***Diexodus debilis* Handlirsch 1911** Holotype from the Francis Creek Shale (Carbondale Formation) at Mazon Creek, Illinois (Handlirsch, 1911), of Moscovian (Westphalian D) age (Peppers, 1996), giving 306.0–311.0 Mya. A palaeodictyopteran of uncertain family (order Palaeodictyoptera).

***Dunbaria borealis* Sharov & Sinitshenkova 1977** Holotype from a depth of 360.1 m in borehole K-384 of the Vorkuta Deposit, in the Rudnitskaya Member of the Lekvorkuta Formation of the Pechora (Vorkuta) Basin (Sharov & Sinitshenkova, 1977), of Kungurian age (Davydov et al., 2018), giving 273.0–283.5 Mya. A Spilapteridae (order Palaeodictyoptera).

***Dunbaria elkunensis* Sinitshenkova 2025** Holotype from the Baitugan Beds at Golyusherma, Udmurtia (PIN collection 5909) (Sinitshenkova, 2025), of Kazanian (Roadian) age (Davydov et al., 2018), giving 266.9–277.0 Mya. A Spilapteridae (order Palaeodictyoptera).

***Dunbaria fasciipennis* Tillyard 1924** Holotype from the Wellington Formation at Elmo, Kansas (Dunbar & Tillyard, 1924), of Leonardian (late Artinskian–early Kungurian) age (Tasch & Zimmerman, 1962; Schneider et al., 2020), giving 279–286 Mya. A Spilapteridae (order Palaeodictyoptera).

***Dunbaria mira* (Zalessky 1946)** Holotype from the Bardian beds of the Cis-Urals (Zalessky, 1946), of Kungurian (Irenian substage) age (Lucas & Shen, 2018; Khramov et al., 2023), giving 273.0–283.5 Mya. A Spilapteridae (order Palaeodictyoptera).

***Dyscritus vetustus* Scudder 1880** Holotype from Plant Bed 8 of the Fern Ledges (Lancaster Formation) at Carleton, Saint John, New Brunswick, Canada (Scudder, 1880), of Langsettian (Westphalian A) age (Gradstein et al., 2020; Schneider et al., 2020), giving 316.5–319.2 Mya. A Lithomanteidae (order Palaeodictyoptera).

***Elmoa trisecta* Tillyard 1937** Holotype from the Wellington Formation at Elmo, Kansas (Tillyard, 1937), of Leonardian (late Artinskian–early Kungurian) age (Tasch & Zimmerman, 1962; Schneider et al., 2020), giving 279–286 Mya. An Elmoidae (order Diaphanopterodea).

***Elmoboria piperi* Carpenter 1976** Holotype from the Wellington Formation at Elmo, Kansas (Carpenter, 1976), of Leonardian (late Artinskian–early Kungurian) age (Tasch & Zimmerman, 1962;

Schneider et al., 2020), giving 279–286 Mya. An Elmoboriidae (order Palaeodictyoptera).

***Elmodiapha ovata* Kukalová-Peck 1974** Holotype from the upper Letovice Formation at Obora, southern Moravia (Kukalová-Peck, 1974), of Sakmarian–Artinskian (Early Permian) age (Opluštil et al., 2017), giving 283.30–293.52 Mya. A diaphanopterodean of uncertain family (order Diaphanopterodea).

***Engisoptera ivanae* Rýznar 2021** Holotype from the upper Letovice Formation at Obora, southern Moravia (Rýznar, 2021), of Sakmarian–Artinskian (Early Permian) age (Opluštil et al., 2017), giving 283.30–293.52 Mya. An Engisopteridae (order Megasecoptera).

***Engisoptera simplices* Kukalová-Peck 1975** Holotype from the upper Letovice Formation at Obora, southern Moravia (Kukalová-Peck, 1975), of Sakmarian–Artinskian (Early Permian) age (Opluštil et al., 2017), giving 283.30–293.52 Mya. An Engisopteridae (order Megasecoptera).

***Epitethe meunieri* (Brongniart 1894)** Holotype from the bituminous shale of the Commentry coal basin at Commentry, Allier, France (Brongniart, 1894), of Gzhelian (Stephanian B) age (Gradstein et al., 2020), giving 298.9–303.7 Mya. A Spilapteridae (order Palaeodictyoptera).

***Eubleptus danielsi* Handlirsch 1906** Holotype from the Francis Creek Shale (Carbondale Formation) at Mazon Creek, Illinois (Handlirsch, 1906b), of Moscovian (Westphalian D) age (Peppers, 1996), giving 306.0–311.0 Mya. A Eubleptidae (order Palaeodictyoptera).

***Eubleptus maculosus* Carpenter 1992** Holotype from the Francis Creek Shale (Carbondale Formation) at Pit 11, Braidwood, Mazon Creek, Illinois (Carpenter, 1992), of Moscovian (Westphalian D) age (Peppers, 1996), giving 306.0–311.0 Mya. A Eubleptidae (order Palaeodictyoptera).

***Eubrodia dabasinskasi* Carpenter 1967** Holotype from the Francis Creek Shale (Carbondale Formation) at Pit 11, Braidwood, Mazon Creek, Illinois (Carpenter, 1967), of Moscovian (Westphalian D) age (Peppers, 1996), giving 306.0–311.0 Mya. A Brodiidae (order Megasecoptera).

***Euchoroptera longipennis* Carpenter 1940** Holotype from the Rock Lake Shale Member of the Stanton Formation at the Bradford Chandler farm, Anderson County, Kansas (Carpenter, 1940), of late Kasimovian age (Schneider et al., 2020), giving 303.9–306.0 Mya. A Prochoropteridae (order Diaphanopterodea).

***Eugereon boeckingi* Dohrn 1866** Holotype from the Lower Permian of Birkenfeld, Germany (Dohrn, 1866), of Asselian age (Lucas & Shen, 2018), giving 293.5–298.9 Mya. An Eugereonidae (order Palaeodictyoptera).

***Eumartynovia raaschi* Carpenter 1947** Holotype from the Midco Member of the Wellington Formation at Midco locality 15, Oklahoma (Carpenter, 1947), of Leonardian (late Artinskian–early Kungurian) age (Tasch & Zimmerman, 1962; Schneider et al., 2020), giving 279–286 Mya. A Martynoviidae (order Diaphanopterodea).

***Eumecoptera laxa* (Goldenberg 1877)** Holotype from the Lower Saarbrücken layers at the mine dump near Dudweiler station, Saarland, Germany (Goldenberg, 1877), of Westphalian C (Bolsovian) age (Gradstein et al., 2020; Schneider et al., 2020), giving 310.7–313.7 Mya. A palaeodictyopteran of uncertain family (order Palaeodictyoptera).

***Eurydictyella richteri* (Guthörl 1934)** Holotype from the Friedrichsthal railway cutting, Saarbrücken district, Saarland, Germany (Guthörl, 1934), of Westphalian age (Gradstein et al., 2020; Schneider et al., 2020), giving 307.5–319.2 Mya. A palaeodictyopteran of uncertain family (order Palaeodictyoptera).

***Eurythmopteryx antiqua* Handlirsch 1906** Holotype from the Pottsville Formation at the Pratt mines, Birmingham, Alabama (Handlirsch, 1906b), of Westphalian A age (Beckemeyer & Engel, 2011), giving 315.2–319.0 Mya. A palaeodictyopteran of uncertain family (order Palaeodictyoptera).

***Forcynthia cynthiae* Sinitshenkova 2004** Holotype from the Red Tanks Member of the Bursum Formation at Carrizo Arroyo, New Mexico (Rasnitsyn et al., 2004), of latest Gzhelian–earliest Asselian (Carboniferous–Permian boundary) age (Schneider et al., 2020), giving 296.5–299.5 Mya. A Hanidae (order Palaeodictyoptera).

***Foriria maculata* Meunier 1908** Holotype from the Commentry Shales at Commentry, Allier, France (Meunier, 1908a), of Gzhelian (Stephanian B) age (Gradstein et al., 2020), giving 298.9–303.7 Mya. A Foririidae (order Megasecoptera).

***Fouquea lacroixi* Brongniart 1894** Holotype from the bituminous shale of the Commentry coal basin at Commentry, Allier, France (Brongniart, 1894), of Gzhelian (Stephanian B) age (Gradstein et al., 2020), giving 298.9–303.7 Mya. A Fouqueidae (order Palaeodictyoptera).

***Fouquea needhami* Lameere 1917** Holotype from the bituminous shale of the Commentry coal basin at Commentry, Allier, France (Lameere, 1917), of Gzhelian (Stephanian B) age (Gradstein et al., 2020), giving 298.9–303.7 Mya. A Fouqueidae (order Palaeodictyoptera).

***Fouquea sauvagei* Brongniart 1894** Holotype from the bituminous shale of the Commentry coal basin at Commentry, Allier, France (Brongniart, 1894), of Gzhelian (Stephanian B) age (Gradstein et al., 2020), giving 298.9–303.7 Mya. A Fouqueidae (order Palaeodictyoptera).

***Fouquea superba* (Meunier 1909)** Holotype from the bituminous shale of the Commentry coal basin at Commentry, Allier, France (Meunier, 1909), of Gzhelian (Stephanian B) age (Gradstein et al., 2020), giving 298.9–303.7 Mya. A Fouqueidae (order Palaeodictyoptera).

***Fragmohymen submissus* Novokshonov 1995** Holotype from the Koshelevka Formation at Chekarda on the Sylva River, Perm region, Russia (Novokshonov, 1995), of Kungurian (Irenian substage) age (Lucas & Shen, 2018; Khramov et al., 2023), giving 273.0–283.5 Mya. A Vorkutiidae (order Megasecoptera).

***Gegenemene sinuosa* (Kliver 1883)** Holotype from the Lower Saarbrücken layers at Dudweiler, Saarland, Germany (Kliver, 1883), of Westphalian C (Bolsovian) age (Gradstein et al., 2020; Schneider et al., 2020), giving 310.7–313.7 Mya. A palaeodictyopteran of uncertain family (order Palaeodictyoptera).

***Goldenbergia elongata* (Goldenberg 1877)** Holotype from the Lower Saarbrücken layers at the mine dump near Dudweiler station, Saarland, Germany (Goldenberg, 1877), of Westphalian C (Bolsovian) age (Gradstein et al., 2020; Schneider et al., 2020), giving 310.7–313.7 Mya. A Dictyoneuridae (order Palaeodictyoptera).

***Goldenbergia formosa* (Goldenberg 1854)** Holotype from the Burguklinskaya Formation on the Fat'yanikha River, Tunguska Basin (Goldenberg, 1854; Sharov & Sinitshenkova, 1977), of Sakmarian age (Budnikov et al., 2020), giving 290.1–293.52 Mya. A Dictyoneuridae (order Palaeodictyoptera).

***Grandelmoa jingqii* Xu et al. 2025** Holotype from the Shanxi Formation at Zhiyu, Shanxi Province, China (Xu et al., 2025), of mid-Asselian age (Wu et al., 2021; Santos et al., 2024), giving 294.0–297.0 Mya. A Parelmoidae (order Diaphanopterodea).

***Graphiptilus heeri*** **Brongniart 1894** Holotype from the bituminous shale of the Commentry coal basin at Commentry, Allier, France (Brongniart, 1894), of Gzhelian (Stephanian B) age (Gradstein et al., 2020), giving 298.9–303.7 Mya. A Graphiptilidae (order Palaeodictyoptera).

***Hadroneuria heidei*** **Laurentiaux 1950** Holotype (G.B. no. 17) from the roof of vein X at the Wilhelmina Mine, Terwinselen, South Limburg, Netherlands (Laurentiaux, 1950), of Westphalian A (Langsettian) age (Gradstein et al., 2020; Schneider et al., 2020), giving 316.5–319.2 Mya. A Lithomanteidae (order Palaeodictyoptera).

***Hana filia*** **Kukalová-Peck 1975** Holotype from the upper Letovice Formation at Obora, southern Moravia (Kukalová-Peck, 1975), of Sakmarian–Artinskian (Early Permian) age (Opluštil et al., 2017), giving 283.30–293.52 Mya. A Hanidae (order Megasecoptera).

***Hana lineata*** **Kukalová-Peck 1975** Holotype from the upper Letovice Formation at Obora, southern Moravia (Kukalová-Peck, 1975), of Sakmarian–Artinskian (Early Permian) age (Opluštil et al., 2017), giving 283.30–293.52 Mya. A Hanidae (order Megasecoptera).

***Haplophlebium barnesii*** **Scudder 1867** Holotype from the Sydney Coalfield succession (Morien Group) at Schooner Pond, Cape Breton, Nova Scotia (Scudder, 1867), of Westphalian D–Cantabrian age (Rust et al., 1987), giving 303.4–312.0 Mya. A palaeodictyopteran of uncertain family (order Palaeodictyoptera).

***Hasala inferiorsaxonica*** **Brauckmann 1995** Holotype from the Flöz Zweibänke at the Piesberg quarry near Osnabrück, Lower Saxony, Germany (Brauckmann, 1995), of Westphalian D (Asturian) age (Gradstein et al., 2020; Schneider et al., 2020), giving 307.50–310.70 Mya. A Breyeriidae (order Palaeodictyoptera).

***Haseneura jarmilae*** **Rosová et al. 2024** Holotype from the Flöz Dreibänke at the Piesberg quarry near Osnabrück, Lower Saxony, Germany (Rosová et al., 2024), of Westphalian D (Asturian) age (Gradstein et al., 2020; Schneider et al., 2020), giving 307.50–310.70 Mya. An Archaemegaptilidae (order Palaeodictyoptera).

***Haseneura meyeri*** **(Brauckmann et al. 2009)** Holotype from the Flöz Dreibänke at the Piesberg quarry near Osnabrück, Lower Saxony, Germany (Brauckmann et al., 2009), of Westphalian D

(Asturian) age (Gradstein et al., 2020; Schneider et al., 2020), giving 307.50–310.70 Mya. An Archaemegaptilidae (order Palaeodictyoptera).

***Heolus providentiae* Handlirsch 1906** Holotype from the Upper Carboniferous coal measures at East Providence, Rhode Island (USA) (Handlirsch, 1906b), of Moscovian age (Gradstein et al., 2020), giving 310.7–313.7 Mya. A palaeodictyopteran of uncertain family (order Palaeodictyoptera).

***Homaloneura berenice* Brauckmann & Gröning 1998** Holotype from the Vorhalle Beds at the Hagen-Vorhalle quarry, North Rhine-Westphalia, Germany (Brauckmann & Gröning, 1998), of late Marsdenian (Namurian B, *Bilinguites metabilinguis* Marine Band, Ziegelschiefer Formation) age (Ilger & Brauckmann, 2011; Opluštil et al., 2022), giving 319.0–320.5 Mya. A Spilapteridae (order Palaeodictyoptera).

***Homaloneura bonnieri* (Brongniart 1894)** Holotype from the bituminous shale of the Commentry coal basin at Commentry, Allier, France (Brongniart, 1894), of Gzhelian (Stephanian B) age (Gradstein et al., 2020), giving 298.9–303.7 Mya. A Spilapteridae (order Palaeodictyoptera).

***Homaloneura brauckmanni* Sinitshenkova 2020** Holotype from a depth of 355.4 m in borehole HK-966 of the Halmeryu coalfield, in the Rudnitskaya Member of the Lekvorkuta Formation of the Pechora (Vorkuta) Basin (Sinitshenkova, 2020), of Kungurian age (Davydov et al., 2018), giving 273.0–283.5 Mya. A Spilapteridae (order Palaeodictyoptera).

***Homaloneura bucklandi* Brongniart 1894** Holotype from the bituminous shale of the Commentry coal basin at Commentry, Allier, France (Brongniart, 1894), of Gzhelian (Stephanian B) age (Gradstein et al., 2020), giving 298.9–303.7 Mya. A Spilapteridae (order Palaeodictyoptera).

***Homaloneura dabasinskasi* Carpenter 1964** Holotype from the Francis Creek Shale (Carbondale Formation) at Will County, Illinois (Carpenter, 1964), of Moscovian (Westphalian D) age (Peppers, 1996), giving 306.0–311.0 Mya. A Spilapteridae (order Palaeodictyoptera).

***Homaloneura elegans* Brongniart 1885** Holotype from the bituminous shale of the Commentry coal basin at Commentry, Allier, France (Brongniart, 1885), of Gzhelian (Stephanian B) age (Gradstein et al., 2020), giving 298.9–303.7 Mya. A Spilapteridae (order Palaeodictyoptera).

***Homaloneura joannae* (Brongniart 1894)** Holotype from the bituminous shale of the Commentry coal basin at Commentry, Allier, France (Brongniart, 1894), of Gzhelian (Stephanian B) age

(Gradstein et al., 2020), giving 298.9–303.7 Mya. A Spilapteridae (order Palaeodictyoptera).

***Homaloneura kiliani* Brauckmann et al. 2009** Holotype from the Flöz Dreibänke at the Piesberg quarry near Osnabrück, Lower Saxony, Germany (Brauckmann et al., 2009), of Westphalian D (Asturian) age (Gradstein et al., 2020; Schneider et al., 2020), giving 307.50–310.70 Mya. A Spilapteridae (order Palaeodictyoptera).

***Homaloneura lehmani* Kukalová 1969** Holotype from the bituminous shale of the Commentry coal basin at Commentry, Allier, France (Kukalová, 1969), of Gzhelian (Stephanian B) age (Gradstein et al., 2020), giving 298.9–303.7 Mya. A Spilapteridae (order Palaeodictyoptera).

***Homaloneura ligeia* Brauckmann 1986** Holotype from the Vorhalle Beds at the Hagen-Vorhalle quarry, North Rhine-Westphalia, Germany (Brauckmann, 1986), of late Marsdenian (Namurian B, *Bilinguites metabilinguis* Marine Band, Ziegelschiefer Formation) age (Ilger & Brauckmann, 2011; Opluštil et al., 2022), giving 319.0–320.5 Mya. A Spilapteridae (order Palaeodictyoptera).

***Homaloneura ornata* Brongniart 1894** Holotype from the bituminous shale of the Commentry coal basin at Commentry, Allier, France (Brongniart, 1894), of Gzhelian (Stephanian B) age (Gradstein et al., 2020), giving 298.9–303.7 Mya. A Spilapteridae (order Palaeodictyoptera).

***Homaloneura punctata* Brongniart 1894** Holotype from the bituminous shale of the Commentry coal basin at Commentry, Allier, France (Brongniart, 1894), of Gzhelian (Stephanian B) age (Gradstein et al., 2020), giving 298.9–303.7 Mya. A Spilapteridae (order Palaeodictyoptera).

***Homaloneura redbirdi* (Brauckmann et al. 1993)** Holotype from the Whitesburg coal zone of the Breathitt Group at the Daniel Boone Parkway roadcut, Kentucky, of Duckmantian (Westphalian B) age (Brauckmann et al., 1993), giving 313.5–316.5 Mya. A Spilapteridae (order Palaeodictyoptera).

***Homaloneura ribeiroi* Teixeira 1946** Holotype from the Santa Bárbara works at São Pedro da Cova, Douro Carboniferous Basin, northern Portugal (Teixeira, 1946), of early Gzhelian age (Correia et al., 2019), giving 300.5–303.7 Mya. A Spilapteridae (order Palaeodictyoptera).

***Homoioptera gigantea* Agnus 1902** Holotype from the Commentry Shales at Commentry, Allier, France (Agnus, 1902), of Gzhelian (Stephanian B) age (Gradstein et al., 2020), giving 298.9–303.7 Mya. A Homoiopteridae (order Palaeodictyoptera).

***Homoioptera kortumi* Brauckmann & Herd 2002** Holotype from the Flöz Zweibänke at the Piesberg quarry near Osnabrück, Lower Saxony, Germany (Brauckmann & Herd, 2002), of Westphalian D (Asturian) age (Gradstein et al., 2020; Schneider et al., 2020), giving 307.50–310.70 Mya. A Homoiopteridae (order Palaeodictyoptera).

***Homoioptera latipenne* (Handlirsch 1904)** Holotype from the Flénu Formation at the Belle-et-Bonne colliery (Avaleresse pit), Frameries, Hainaut, Belgium (Handlirsch, 1904), of Bolsovian (Westphalian C) age (Gradstein et al., 2020; Schneider et al., 2020), giving 310.7–313.7 Mya. A Homoiopteridae (order Palaeodictyoptera).

***Homoioptera vorhallensis* Brauckmann & Koch 1982** Holotype from the Vorhalle Beds at the Hagen-Vorhalle quarry, North Rhine-Westphalia, Germany (Brauckmann & Koch, 1982), of late Marsdenian (Namurian B, *Bilinguites metabilinguis* Marine Band, Ziegelschiefer Formation) age (Ilger & Brauckmann, 2011; Opluštil et al., 2022), giving 319.0–320.5 Mya. A Homoiopteridae (order Palaeodictyoptera).

***Homothetus fossilis* Scudder 1867** Holotype from Plant Bed 8 of the Fern Ledges (Lancaster Formation) at Carleton, Saint John, New Brunswick, Canada (Scudder, 1867), of Langsettian (Westphalian A) age (Gradstein et al., 2020; Schneider et al., 2020), giving 316.5–319.2 Mya. A Homothetidae (order Palaeodictyoptera).

***Hongius qiligouensis* (Hong 1985)** Holotype from the Shanxi Formation at Qiligou, Xishan of Taiyuan, Shanxi Province, China (Hong, 1985a), of mid-Asselian age (Wu et al., 2021; Santos et al., 2024), giving 294.0–297.0 Mya. A Dictyoneuridae (order Palaeodictyoptera).

***Ideliella decora* Zalessky 1937** Holotype from the Baitugan Beds at Tikhie Gory, Volga region (Zalessky, 1937b), of early Kazanian (Roadian) age (Davydov et al., 2018), giving 266.9–277.0 Mya. A Permothemistidae (order Permothemistida).

***Idoptilus onisciformis* Wootton 1972** Holotype from the Coal Measures at Round Green opencast mine, Barnsley, Yorkshire, England (Wootton, 1972), of Duckmantian (Westphalian B) age (Gradstein et al., 2020; Schneider et al., 2020), giving 313.7–316.5 Mya. A palaeodictyopteran of uncertain family (order Palaeodictyoptera).

***Idoptilus peachii* (Woodward 1887)** Holotype from the Coal Measures at Greenhill Pit, Kilmaurs, Ayrshire, Scotland (Woodward, 1887), of Duckmantian (Westphalian B) age (Gradstein et al., 2020; Schneider et al., 2020), giving 313.7–316.5 Mya. A palaeodictyopteran of uncertain family (order Palaeodictyoptera).

***Ischnoptilus elegans* Brongniart 1894** Holotype from the bituminous shale of the Commentry coal basin at Commentry, Allier, France (Brongniart, 1894), of Gzhelian (Stephanian B) age (Gradstein et al., 2020), giving 298.9–303.7 Mya. An Ischnoptilidae (order Megasecoptera).

***Issadohymen ponomarenkoi* Sinitshenkova & Aristov 2013** Holotype from the Kichuga Member of the Poldarsa Formation at Isady, Vologda region (PIN collection 3840) (Sinitshenkova & Aristov, 2013), of latest Severodvinian (early Wuchiapingian) age (Gradstein et al., 2020), giving 256.0–259.51 Mya. A Moravohymenidae (order Megasecoptera).

***Ivahymen constrictus* Martynov 1932** Holotype from the Iva-Gora Beds at Soyana on the Soyana River, Arkhangelsk Region, Russia (Martynov, 1932), of Roadian (Kazanian) age (Naugolnykh, 2015; Davydov et al., 2018), giving 266.9–277.0 Mya. A Protohymenidae (order Megasecoptera).

***Jongmansia tuberculata* (Bolton 1921)** Holotype from the Middle Coal Measures at Sparth Bottoms, Rochdale, Lancashire, England (Bolton, 1921), of Langsettian (Westphalian A) age (Gradstein et al., 2020; Schneider et al., 2020), giving 316.5–319.2 Mya. A Jongmansiidae (order Palaeodictyoptera).

***Jugobreyeria sippelorum* Brauckmann et al. 1985** Holotype from the Vorhalle Beds at the Hagen-Vorhalle quarry, North Rhine-Westphalia, Germany (Brauckmann et al., 1985), of late Marsdenian (Namurian B, *Bilinguites metabilinguis* Marine Band, Ziegelschiefer Formation) age (Ilger & Brauckmann, 2011; Opluštil et al., 2022), giving 319.0–320.5 Mya. A Breyeriidae (order Palaeodictyoptera).

***Kallenbergia handlirschi* (Guthörl 1930)** Holotype from the Friedrichsthal railway cutting, Saarbrücken district, Saarland, Germany (Guthörl, 1930), of Westphalian age (Gradstein et al., 2020; Schneider et al., 2020), giving 307.5–319.2 Mya. A Dictyoneuridae (order Palaeodictyoptera).

***Kansasia pulchra* Tillyard 1937** Holotype from the Wellington Formation at Elmo, Kansas (Tillyard, 1937), of Leonardian (late Artinskian–early Kungurian) age (Tasch & Zimmerman, 1962; Schneider et al., 2020), giving 279–286 Mya. A palaeodictyopteran of uncertain family (order Palaeodictyoptera).

***Karoohymen delicatulus* Riek 1976** Holotype from the Mooi River locality (Daptocephalus Assemblage Zone, Beaufort Group), KwaZulu-Natal, South Africa (Riek, 1976b), of Changhsingian age (Gastaldo et al., 2020), giving 252.0–254.14 Mya. A Scytohymenidae (order Megasecoptera).

***Karoohymen minutus* van Dijk & Geertsema 1999** Holotype from the Mooi River locality (Daptocephalus Assemblage Zone, Beaufort Group), KwaZulu-Natal, South Africa (van Dijk & Geertsema, 1999), of Changhsingian age (Gastaldo et al., 2020), giving 252.0–254.14 Mya. A Scytohymenidae (order Megasecoptera).

***Katosaxoniapteron brauneri* Prokop et al. 2023** Holotype (Pal1242ab) from the claystone above the Flöz Dreibänke at the Piesberg quarry near Osnabrück, Lower Saxony, Germany (Prokop et al., 2023), of Westphalian D (Asturian) age (Gradstein et al., 2020; Schneider et al., 2020), giving 307.50–310.70 Mya. A palaeodictyopteran of uncertain family (order Palaeodictyoptera).

***Lameereites curvipennis* Handlirsch 1911** Holotype from the Francis Creek Shale (Carbondale Formation) at Mazon Creek, Illinois (Handlirsch, 1911), of Moscovian (Westphalian D) age (Peppers, 1996), giving 306.0–311.0 Mya. A megasecopteran of uncertain family (order Megasecoptera).

***Lamproptilia grandeuryi* Brongniart 1885** Holotype from the Commentry Shales at Commentry, Allier, France (Brongniart, 1885), of Gzhelian (Stephanian B) age (Gradstein et al., 2020), giving 298.9–303.7 Mya. A Spilapteridae (order Palaeodictyoptera).

***Lamproptilia tenuitegminata* Bolton 1911** Holotype from the Coal Measures at Resolven Station, near Neath, South Wales (Bolton, 1911), of Bolsovian (Westphalian C) age (Gradstein et al., 2020; Schneider et al., 2020), giving 310.7–313.7 Mya. A Spilapteridae (order Palaeodictyoptera).

***Larryia osterbergi* Kukalová-Peck and Richardson 1983** Holotype from the Francis Creek Shale (Carbondale Formation) at Mazon Creek, Illinois (Kukalová-Peck & Richardson, 1983), of Moscovian (Westphalian D) age (Peppers, 1996), giving 306.0–311.0 Mya. A Homoiopteridae (order Palaeodictyoptera).

***Lithomantis bohemica* (Novák 1880)** Holotype from the Upper Carboniferous of Stradonitz (Stradonice), Bohemia (Czech Republic) (Novák, 1880), of Moscovian age (Gradstein et al., 2020), giving 307.0–315.2 Mya. A Lithomanteidae (order Palaeodictyoptera).

***Lithomantis carbonarius* Woodward 1876** Holotype from the Coal Measures on the coast of Ayrshire, near Ayr, Scotland (Woodward, 1876), of Duckmantian (Westphalian B) age (Gradstein et al., 2020; Schneider et al., 2020), giving 313.7–316.5 Mya. A Lithomanteidae (order Palaeodictyoptera).

***Lithomantis varius* Brauckmann et al. 1985** Holotype from the Vorhalle Beds at the Hagen-Vorhalle quarry, North Rhine-Westphalia, Germany (Brauckmann et al., 1985), of late Marsdenian (Namurian B, *Bilinguites metabilinguis* Marine Band, Ziegelschiefer Formation) age (Ilger & Brauckmann, 2011; Opluštil et al., 2022), giving 319.0–320.5 Mya. A Lithomantidae (order Palaeodictyoptera).

***Lithomantis woodwardi* Brongniart 1890** Holotype from the bituminous shale of the Commentry coal basin at Commentry, Allier, France (Brongniart, 1890), of Gzhelian (Stephanian B) age (Gradstein et al., 2020), giving 298.9–303.7 Mya. A Lithomanteidae (order Palaeodictyoptera).

***Lithoptilus boulei* (Meunier 1908)** Holotype from the Commentry Shales at Commentry, Allier, France (Meunier, 1908b), of Gzhelian (Stephanian B) age (Gradstein et al., 2020), giving 298.9–303.7 Mya. A Megaptilidae (order Palaeodictyoptera).

***Lithosialis brongniarti* (Mantell 1844)** Holotype from the Pennystone ironstone at Coalbrookdale, Shropshire, England (Mantell, 1844), of Bolsovian (Westphalian C) age (Gradstein et al., 2020; Schneider et al., 2020), giving 310.7–313.7 Mya. A Lithomanteidae (order Palaeodictyoptera).

***Lithosialis daviesii* Bolton 1930** Holotype from the Coal Measures at Cambrian Colliery, Clydach Vale, Rhondda, South Wales (Bolton, 1930), of Duckmantian (Westphalian B) age (Gradstein et al., 2020; Schneider et al., 2020), giving 313.7–316.5 Mya. A Lithomanteidae (order Palaeodictyoptera).

***Lithosialis truemani* Bolton 1930** Holotype from the Coal Measures at Port Henry Colliery, South Wales (Bolton, 1930), of Langsettian (Westphalian A) age (Gradstein et al., 2020; Schneider et al., 2020), giving 316.5–319.2 Mya. A Lithomanteidae (order Palaeodictyoptera).

***Lodetiella magnifica* Béthoux et al. 2007** Holotype from the Mérifons Member of the Salagou Formation at the La Lieude farm, Lodève Basin, southern France (Béthoux et al., 2007), of late Kungurian age, younger than the 282.86 Mya tuff dated in the underlying Octon Member (Michel et al., 2015), giving 272.95–282.86 Mya. A Calvertiellidae (order Palaeodictyoptera).

***Lodevohymen lapeyriei* Prokop & Nel 2011** Holotype from locality F23, Les Canals, Mérifons Member of the Salagou Formation, Lodève Basin, southern France (Prokop & Nel, 2011), of late Kungurian age, younger than the 282.86 Mya tuff dated in the underlying Octon Member (Michel et al., 2015), giving 272.95–282.86 Mya. A diaphanopterodean of uncertain family (order Diaphanopterodea).

***Lusiella fariai* (Laurentiaux & Teixeira 1958)** Holotype from Alto do Pejão (Baixo-Douro), Douro Carboniferous Basin, northern Portugal (Laurentiaux & Teixeira, 1958), of early Gzhelian age (Correia et al., 2019), giving 300.5–303.7 Mya. A Lithomanteidae (order Palaeodictyoptera).

***Lusiella greberi* (Laurentiaux 1949)** Holotype from the Cuvinot pit (360 m level) at Valenciennes, Nord, France (Laurentiaux, 1949a), of Duckmantian (Westphalian B) age (Gradstein et al., 2020; Schneider et al., 2020), giving 313.7–316.5 Mya. A Lithomanteidae (order Palaeodictyoptera).

***Lycocercus goldenbergi* (Brongniart 1885)** Holotype from the Commentry Shales at Commentry, Allier, France (Brongniart, 1885), of Gzhelian (Stephanian B) age (Gradstein et al., 2020), giving 298.9–303.7 Mya. A Lycocercidae (order Palaeodictyoptera).

***Lycocercus pictus* Handlirsch 1919** Holotype from the Commentry Shales at Commentry, Allier, France (Handlirsch, 1919), of Gzhelian (Stephanian B) age (Gradstein et al., 2020), giving 298.9–303.7 Mya. A Lycocercidae (order Palaeodictyoptera).

***Lycodemas adolescens* Carpenter and Richardson 1971** Holotype from the Francis Creek Shale (Carbondale Formation) at Pit 11, Braidwood, Mazon Creek, Illinois (Carpenter & Richardson, 1971), of Moscovian (Westphalian D) age (Peppers, 1996), giving 306.0–311.0 Mya. A Lycocercidae (order Palaeodictyoptera).

***Lycodus garretti* Carpenter 1992** Holotype from the Lawrence Shale at the Lone Star Lake spillway, Kansas (Carpenter, 1992), of latest Kasimovian age (Schneider et al., 2020), giving 303.7–304.5 Mya. A Lycocercidae (order Palaeodictyoptera).

***Macrodictya stenomediali* Guthörl 1940** Holotype from the Geisheck Formation at the Frankenholz-Saar colliery (shaft 5), Saarland, Germany (Guthörl, 1940), of Westphalian D (Asturian) age (Gradstein et al., 2020; Schneider et al., 2020), giving 307.5–310.7 Mya. A Dictyoneuridae (order Palaeodictyoptera).

***Madera mamayi* Carpenter 1970** Holotype from the Tinajas Member of the Atrasado Formation (formerly Madera Group) at the Kinney Brick Quarry, New Mexico (Carpenter, 1970), of middle Kasimovian age (Schneider et al., 2020), giving 304.0–306.5 Mya. A Lycocercidae (order Palaeodictyoptera).

***Mammia alutacea* Handlirsch 1906** Holotype from the Francis Creek Shale (Carbondale Formation) at Mazon Creek, Illinois (Handlirsch, 1906b), of Moscovian (Westphalian D) age (Peppers, 1996), giving 306.0–311.0 Mya. A palaeodictyopteran of uncertain family (order Palaeodictyoptera).

***Martynovia halli* Beckemeyer 2004** Holotype from the Midco Member of the Wellington Formation at Midco locality 16, Noble County, Oklahoma (Beckemeyer, 2004), of Leonardian (late Artinskian–early Kungurian) age (Tasch & Zimmerman, 1962; Schneider et al., 2020), giving 279–286 Mya. A Martynoviidae (order Diaphanopterodea).

***Martynovia insignis* Tillyard 1932** Holotype from the Wellington Formation at Elmo, Kansas (Tillyard, 1932), of Leonardian (late Artinskian–early Kungurian) age (Tasch & Zimmerman, 1962; Schneider et al., 2020), giving 279–286 Mya. A Martynoviidae (order Diaphanopterodea).

***Martynovia longipennis* Carpenter 1947** Holotype from the Midco Member of the Wellington Formation at Midco locality 16, Oklahoma (Carpenter, 1947), of Leonardian (late Artinskian–early Kungurian) age (Tasch & Zimmerman, 1962; Schneider et al., 2020), giving 279–286 Mya. A Martynoviidae (order Diaphanopterodea).

***Martynovia protohymenoides* (Tillyard 1932)** Holotype from the Wellington Formation at Elmo, Kansas (Tillyard, 1932), of Leonardian (late Artinskian–early Kungurian) age (Tasch & Zimmerman, 1962; Schneider et al., 2020), giving 279–286 Mya. A Martynoviidae (order Diaphanopterodea).

***Mazonopterum cooperi* Prokop et al. 2018** Holotype from the Farrington Formation at the Writhlington Geological Nature Reserve, near Radstock, Somerset, England (Prokop et al., 2018), of Asturian (Westphalian D) age (Gradstein et al., 2020; Schneider et al., 2020), giving 307.5–310.7 Mya. A Homoiopteridae (order Palaeodictyoptera).

***Mazonopterum wolfforum* Kukalová-Peck and Richardson 1983** Holotype (FMNH PE 15138) from the Francis Creek Shale (Carbondale Formation) at Worthington Township, Will County, Illinois (Kukalová-Peck & Richardson, 1983), of Moscovian (Westphalian D) age (Peppers, 1996), giving

306.0–311.0 Mya. A Homoiopteridae (order Palaeodictyoptera).

***Mazothairos enormis* Kukalová-Peck and Richardson 1983** Holotype from the Francis Creek Shale (Carbondale Formation) at Mazon Creek, Illinois (Kukalová-Peck & Richardson, 1983), of Moscovian (Westphalian D) age (Peppers, 1996), giving 306.0–311.0 Mya. A Homoiopteridae (order Palaeodictyoptera).

***Mcluckiepteron luciae* Richardson 1956** Holotype from the Francis Creek Shale (Carbondale Formation) at the Coal City–Braidwood strip mines, Illinois (Richardson, 1956), of Moscovian (Westphalian D) age (Peppers, 1996), giving 306.0–311.0 Mya. A Spilapteridae (order Palaeodictyoptera).

***Mecynoptera splendida* Handlirsch 1904** Holotype from the Sainte-Félicité pit (no. 23) of the Produits colliery at Flénu, Hainaut, Belgium (Handlirsch, 1904), of Bolsovian (Westphalian C) age (Gradstein et al., 2020; Schneider et al., 2020), giving 310.7–313.7 Mya. A palaeodictyopteran of uncertain family (order Palaeodictyoptera).

***Mecynostomata dohrni* (Brongniart 1894)** Holotype from the bituminous shale of the Commentry coal basin at Commentry, Allier, France (Brongniart, 1894), of Gzhelian (Stephanian B) age (Gradstein et al., 2020), giving 298.9–303.7 Mya. A Mecynostomatidae (order Palaeodictyoptera).

***Mecynostomites brongniarti* Handlirsch 1919** Holotype from the Commentry Shales at Commentry, Allier, France (Handlirsch, 1919), of Gzhelian (Stephanian B) age (Gradstein et al., 2020), giving 298.9–303.7 Mya. A palaeodictyopteran of uncertain family (order Palaeodictyoptera).

***Megaptiloides brodiei* (Brongniart 1894)** Holotype from the bituminous shale of the Commentry coal basin at Commentry, Allier, France (Brongniart, 1894), of Gzhelian (Stephanian B) age (Gradstein et al., 2020), giving 298.9–303.7 Mya. A Breyeriidae (order Palaeodictyoptera).

***Megaptilus blanchardi* Brongniart 1885** Holotype from the Commentry Shales at Commentry, Allier, France (Brongniart, 1885), of Gzhelian (Stephanian B) age (Gradstein et al., 2020), giving 298.9–303.7 Mya. A Megaptilidae (order Palaeodictyoptera).

***Microdictya hamyi* (Brongniart 1894)** Holotype from the bituminous shale of the Commentry coal basin at Commentry, Allier, France (Brongniart, 1894), of Gzhelian (Stephanian B) age (Gradstein et al., 2020), giving 298.9–303.7 Mya. A Dictyoneuridae (order Palaeodictyoptera).

***Microdictya handlirschi* (Bolton 1925)** Holotype from the bituminous shale of the Commentry coal basin at Commentry, Allier, France (Bolton, 1925), of Gzhelian (Stephanian B) age (Gradstein et al., 2020), giving 298.9–303.7 Mya. A Dictyoneuridae (order Palaeodictyoptera).

***Microdictya lacroixi* Meunier 1910** Holotype from the bituminous shale of the Commentry coal basin at Commentry, Allier, France (Meunier, 1910), of Gzhelian (Stephanian B) age (Gradstein et al., 2020), giving 298.9–303.7 Mya. A Dictyoneuridae (order Palaeodictyoptera).

***Microdictya latipennis* Handlirsch 1919** Holotype from the bituminous shale of the Commentry coal basin at Commentry, Allier, France (Handlirsch, 1919), of Gzhelian (Stephanian B) age (Gradstein et al., 2020), giving 298.9–303.7 Mya. A Dictyoneuridae (order Palaeodictyoptera).

***Microdictya vaillanti* (Brongniart 1894)** Holotype from the bituminous shale of the Commentry coal basin at Commentry, Allier, France (Brongniart, 1894), of Gzhelian (Stephanian B) age (Gradstein et al., 2020), giving 298.9–303.7 Mya. A Dictyoneuridae (order Palaeodictyoptera).

***Microdictya villeneuvei* Meunier 1908** Holotype from the bituminous shale of the Commentry coal basin at Commentry, Allier, France (Meunier, 1908c), of Gzhelian (Stephanian B) age (Gradstein et al., 2020), giving 298.9–303.7 Mya. A Dictyoneuridae (order Palaeodictyoptera).

***Microhymen inconspicuus* Yang et al. 2023** Holotype from the Yanghugou Formation at Xiaheyan, Zhongwei, Ningxia, China (Yang et al., 2023), of latest Bashkirian to middle Moscovian age (Trümper et al., 2020), giving 309.0–316.0 Mya. A Bardohymenidae (order Megasecoptera).

***Mischoptera douglassi* Carpenter and Richardson 1968** Holotype from the Francis Creek Shale (Carbondale Formation) at Pit 6, Coal City, Mazon Creek, Illinois (Carpenter & Richardson, 1968), of Moscovian (Westphalian D) age (Peppers, 1996), giving 306.0–311.0 Mya. A Mischopteridae (order Megasecoptera).

***Mischoptera nigra* (Brongniart 1885)** Holotype from the bituminous shale of the Commentry coal basin at Commentry, Allier, France (Brongniart, 1885), of Gzhelian (Stephanian B) age (Gradstein et al., 2020), giving 298.9–303.7 Mya. A Mischopteridae (order Megasecoptera).

***Mongolianus callidus* (Sinitshenkova 1992)** Holotype from the Tsankhin Formation at Bor-Tologoi, Mongolia (PIN collection 4305) (Sinitshenkova, 1992), of Wuchiapingian age (Gradstein et al., 2020), giving 254.14–259.51 Mya. A Calvertiellidae (order Palaeodictyoptera).

***Monsteropterum moravicum* Kukalová-Peck 1972** Holotype from the upper Letovice Formation at Obora, southern Moravia (Kukalová-Peck, 1972), of Sakmarian–Artinskian (Early Permian) age (Opluštil et al., 2017), giving 283.30–293.52 Mya. A palaeodictyopteran of uncertain family (order Palaeodictyoptera).

***Moravia convergens* Kukalová 1964** Holotype from the upper Letovice Formation at Obora, southern Moravia (Kukalová, 1964), of Sakmarian–Artinskian (Early Permian) age (Opluštil et al., 2017), giving 283.30–293.52 Mya. A Calvertiellidae (order Palaeodictyoptera).

***Moravia grandis* Carpenter 1979** Holotype from the Midco Member of the Wellington Formation at Midco locality 1, Oklahoma (Carpenter, 1979), of Leonardian (late Artinskian–early Kungurian) age (Tasch & Zimmerman, 1962; Schneider et al., 2020), giving 279–286 Mya. A Calvertiellidae (order Palaeodictyoptera).

***Moraviptera reticulata* Kukalová 1955** Holotype from the Padochov Formation at the Svatý Antonín mine sawmill exposure, Zbýšov, Boskovice Basin (Rosice–Oslavany Coalfield), Moravia, Czech Republic (Kukalová, 1955), of Late Carboniferous (Gzhelian) age (Kukalová, 1955), giving 300.0–304.0 Mya. A Calvertiellidae (order Palaeodictyoptera).

***Moravohymen vitreus* Kukalová-Peck 1972** Holotype from the upper Letovice Formation at Obora, southern Moravia (Kukalová-Peck, 1972), of Sakmarian–Artinskian (Early Permian) age (Opluštil et al., 2017), giving 283.30–293.52 Mya. A Moravohymenidae (order Megasecoptera).

***Moravoptera hladilovae* Dostál & Prokop 2009** Holotype from the upper Letovice Formation at Obora, southern Moravia (Dostál & Prokop, 2009), of Sakmarian–Artinskian (Early Permian) age (Opluštil et al., 2017), giving 283.30–293.52 Mya. A Martynoviidae (order Diaphanopterodea).

***Namurodiapha sippelorum* Kukalová-Peck & Brauckmann 1990** Holotype from the Vorhalle Beds at the Hagen-Vorhalle quarry, North Rhine-Westphalia, Germany (Kukalová-Peck & Brauckmann, 1990), of late Marsdenian (Namurian B, *Bilinguites metabilinguis* Marine Band, Ziegelschiefer Formation) age (Ilger & Brauckmann, 2011; Opluštil et al., 2022), giving 319.0–320.5 Mya. A Namurodiaphidae (order Diaphanopterodea).

***Namuroningxia elegans* Prokop & Ren 2007** Holotype from the Yanghugou Formation at Xiaheyan, Zhongwei, Ningxia, China (Prokop & Ren, 2007), of latest Bashkirian to middle Moscovian age (Trümper et al., 2020), giving 309.0–316.0 Mya. A Namuroningxiidae (order Palaeodictyoptera).

***Namuroptera minuta* Pecharová et al. 2015** Holotype from the Yanghugou Formation at Xiaheyan, Zhongwei, Ningxia, China (Pecharová et al., 2015a), of latest Bashkirian to middle Moscovian age (Trümper et al., 2020), giving 309.0–316.0 Mya. An Aykhalidae (order Megasecoptera).

***Neofouquea suzanneae* Carpenter 1967** Holotype from the Francis Creek Shale (Carbondale Formation) between Braidwood and Coal City, Illinois (Carpenter, 1967), of Moscovian (Westphalian D) age (Peppers, 1996), giving 306.0–311.0 Mya. A Fouqueidae (order Palaeodictyoptera).

***Neuburgia altaica* Martynov 1930** Holotype from the Alykaeva Formation at Verkhotomskoye, Kuznetsk Basin (PIN collection 1079) (Martynov, 1930b; Rohdendorf et al., 1961), of Gzhelian age (Silantiev et al., 2025), giving 298.9–303.7 Mya. A Spilapteridae (order Palaeodictyoptera).

***Oboria longa* Kukalová 1960** Holotype from the upper Letovice Formation at Obora, southern Moravia (Kukalová, 1960), of Sakmarian–Artinskian (Early Permian) age (Opluštil et al., 2017), giving 283.30–293.52 Mya. An Elmoboriidae (order Palaeodictyoptera).

***Oceanoptera elenae* Shcherbakov 2009** Holotype from the Pospelovo Formation at Cape Novosilsky, Russky Island, South Primorye (PIN collection 5306) (Shcherbakov et al., 2009), of Kungurian age (Gradstein et al., 2020), giving 274.4–283.3 Mya. A Scytohymenidae (order Megasecoptera).

***Osnabrugapteron rasnitsyni* Rosová et al. 2022** Holotype from the Flöz Mittel at the Piesberg quarry near Osnabrück, Lower Saxony, Germany (Rosová et al., 2022), of Westphalian D (Asturian) age (Gradstein et al., 2020; Schneider et al., 2020), giving 307.50–310.70 Mya. An Archaemegaptilidae (order Palaeodictyoptera).

***Ostrava nigra* Kukalová 1960** Holotype from coal seam 31 (Upper Sucha Beds) at the Gottwald (František) Mine, Horní Suchá, Karviná, Czech Republic (Kukalová, 1960), of Westphalian A (Langsettian) age (Gradstein et al., 2020; Schneider et al., 2020), giving 316.5–319.2 Mya. A Homoiopteridae (order Palaeodictyoptera).

***Paimbia fenestrata* Sinitshenkova 1979** Holotype from the Kata Formation on the Paymbu River, Tunguska Basin (PIN collection 2293) (Sinitshenkova, 1979), of Gzhelian age (Budnikov et al., 2020), giving 298.9–303.7 Mya. A Tchirkovaeidae (order Palaeodictyoptera).

***Paimbia ultima* Sinitshenkova 1981** Holotype from the Kata Formation on the Chunya River, Tunguska Basin (PIN collection 3115) (Sinitshenkova, 1981), of Gzhelian age (Budnikov et al., 2020), giving 298.9–303.7 Mya. A Tchirkovaeidae (order Palaeodictyoptera).

***Palaeoptilus brullei* Brongniart 1894** Holotype from the Commentry Shales at Commentry, Allier, France (Brongniart, 1894), of Gzhelian (Stephanian B) age (Gradstein et al., 2020), giving 298.9–303.7 Mya. A Spilapteridae (order Palaeodictyoptera).

***Palaiotaptus mazonus* Handlirsch 1906** Holotype from the Francis Creek Shale (Carbondale Formation) at Mazon Creek, Illinois (Handlirsch, 1906b), of Moscovian (Westphalian D) age (Peppers, 1996), giving 306.0–311.0 Mya. A palaeodictyopteran of uncertain family (order Palaeodictyoptera).

***Palapteris stenodictyus* Guthörl 1940** Holotype from the Geisheck Formation at the Frankenholz-Saar colliery (shaft 5), Saarland, Germany (Guthörl, 1940), of Westphalian D (Asturian) age (Gradstein et al., 2020; Schneider et al., 2020), giving 307.5–310.7 Mya. A palaeodictyopteran of uncertain family (order Palaeodictyoptera).

***Paleohymen renmazhuangensis* Hong 1985** Holotype from the Shanxi Formation at Renmazhuang, Xiangning County, Shanxi Province, China (Hong, 1985b), of mid-Asselian age (Wu et al., 2021; Santos et al., 2024), giving 294.0–297.0 Mya. A Bardohymenidae (order Megasecoptera).

***Paleohymen shanxiensis* Hong 1985** Holotype from the Shanxi Formation at Renmazhuang, Xiangning County, Shanxi Province, China (Hong, 1985b), of mid-Asselian age (Wu et al., 2021; Santos et al., 2024), giving 294.0–297.0 Mya. A Bardohymenidae (order Megasecoptera).

***Paraboltonia macroptera* (Bolton 1916)** Holotype from the Middle Coal Measures at the Ravenhead railway cutting, near St Helens, Lancashire, England (Bolton, 1916), of Langsettian (Westphalian A) age (Gradstein et al., 2020; Schneider et al., 2020), giving 316.5–319.2 Mya. A palaeodictyopteran of uncertain family (order Palaeodictyoptera).

***Parabrodia carbonaria* Carpenter 1934** Holotype from the Rock Lake Shale Member of the Stanton Formation near Garnett, Anderson County, Kansas (Carpenter, 1934), of late Kasimovian age (Schneider et al., 2020), giving 303.9–306.0 Mya. A Parabrodiidae (order Diaphanopterodea).

***Paradiapha delicatula* Kukalová-Peck 1974** Holotype from the upper Letovice Formation at Obora, southern Moravia (Kukalová-Peck, 1974), of Sakmarian–Artinskian (Early Permian) age (Opluštil et al., 2017), giving 283.30–293.52 Mya. A diaphanopterodean of uncertain family (order Diaphanopterodea).

***Paradunbaria pectinata* Sharov & Sinitshenkova 1977** Holotype from the Koshelevka Formation at Chekarda on the Sylva River, Perm region, Russia (Sharov & Sinitshenkova, 1977), of Kungurian (Irenian substage) age (Lucas & Shen, 2018; Khramov et al., 2023), giving 273.0–283.5 Mya. A Spilapteridae (order Palaeodictyoptera).

***Parahaplophlebium longipennis* (Scudder 1885)** Holotype from the Llewellyn Formation at Campbells Ledge, near Pittston, Pennsylvania, USA (Scudder, 1885), of Late Pennsylvanian (Gzhelian) age (Scudder, 1885), giving 300.5–303.7 Mya. A palaeodictyopteran of uncertain family (order Palaeodictyoptera).

***Paramecynostoma dohrnianum* Handlirsch 1919** Holotype from the bituminous shale of the Commentry coal basin at Commentry, Allier, France (Handlirsch, 1919), of Gzhelian (Stephanian B) age (Gradstein et al., 2020), giving 298.9–303.7 Mya. A palaeodictyopteran of uncertain family (order Palaeodictyoptera).

***Paramegaptilus scudderi* (Brongniart 1894)** Holotype from the Commentry Shales at Commentry, Allier, France (Brongniart, 1894), of Gzhelian (Stephanian B) age (Gradstein et al., 2020), giving 298.9–303.7 Mya. A palaeodictyopteran of uncertain family (order Palaeodictyoptera).

***Paraostrava stanislavi* Prokop & Nel 2004** Holotype from the Žacléř (Schatzlar) Formation at the Jan Šverma (Julia) mine, Žacléř, Intra-Sudetic Basin, Czech Republic (Prokop & Nel, 2004), of late Westphalian B (Duckmantian) age (Gradstein et al., 2020; Schneider et al., 2020), giving 313.70–314.50 Mya. A Homoiopteridae (order Palaeodictyoptera).

***Parathesoneura anfractuosa* Sinitshenkova 1977** Holotype from Kata-equivalent strata at Razreznyy spring on the Chulyshman River, Baskhaus River basin (Sharov & Sinitshenkova, 1977), of Gzhelian age (Budnikov et al., 2020), giving 298.9–303.7 Mya. A Homoiopteridae (order Palaeodictyoptera).

***Parathesoneura carpenteri* Sinitshenkova 1977** Holotype from the Kata Formation on the Chunya River, Tunguska Basin (PIN collection 3115) (Sharov & Sinitshenkova, 1977), of Gzhelian age (Budnikov et al., 2020), giving 298.9–303.7 Mya. A Homoiopteridae (order Palaeodictyoptera).

***Parelmoa obtusa* Carpenter 1947** Holotype from the Midco Member of the Wellington Formation at Midco locality 15, Oklahoma (Carpenter, 1947), of Leonardian (late Artinskian–early Kungurian) age (Tasch & Zimmerman, 1962; Schneider et al., 2020), giving 279–286 Mya. A Parelmoidae (order Diaphanopterodea).

***Parelmoa radialis* Carpenter 1947** Holotype from the Midco Member of the Wellington Formation at Midco locality 15, Oklahoma (Carpenter, 1947), of Leonardian (late Artinskian–early Kungurian) age (Tasch & Zimmerman, 1962; Schneider et al., 2020), giving 279–286 Mya. A Parelmoidae (order Diaphanopterodea).

***Parelmoa revelata* Carpenter 1947** Holotype from the Midco Member of the Wellington Formation at Midco locality 15, Oklahoma (Carpenter, 1947), of Leonardian (late Artinskian–early Kungurian) age (Tasch & Zimmerman, 1962; Schneider et al., 2020), giving 279–286 Mya. A Parelmoidae (order Diaphanopterodea).

***Paruralia carpenteri* Kukalová-Peck & Sinichenkova 1992** Holotype from the Koshelevka Formation at Chekarda on the Sylva River, Perm region, Russia (Kukalová-Peck & Sinichenkova, 1992), of Kungurian (Irenian substage) age (Lucas & Shen, 2018; Khramov et al., 2023), giving 273.0–283.5 Mya. A Paruraliidae (order Diaphanopterodea).

***Paruralia rohdendorfi* Kukalová-Peck & Sinichenkova 1992** Holotype from the Koshelevka Formation at Chekarda on the Sylva River, Perm region, Russia (Kukalová-Peck & Sinichenkova, 1992), of Kungurian (Irenian substage) age (Lucas & Shen, 2018; Khramov et al., 2023), giving 273.0–283.5 Mya. A Paruraliidae (order Diaphanopterodea).

***Patteiskya bouckaerti* Laurentiaux 1958** Holotype from the Upper Hagen Beds at the Schmiedestraße quarry, Haßlinghausen, North Rhine-Westphalia, Germany (Laurentiaux, 1958), of late Namurian (Marsdenian) age (Gradstein et al., 2020; Opluštil et al., 2022), giving 319.50–320.50 Mya. A Graphiptilidae (order Palaeodictyoptera).

***Patteiskya volmensis* (Brauckmann 1984)** Holotype from the Vorhalle Beds at the Hagen-Vorhalle quarry, North Rhine-Westphalia, Germany (Brauckmann, 1984), of late Marsdenian (Namurian B, *Bilinguites metabilinguis* Marine Band, Ziegelschiefer Formation) age (Ilger & Brauckmann, 2011; Opluštil et al., 2022), giving 319.0–320.5 Mya. A Graphiptilidae (order Palaeodictyoptera).

***Pauciramus demoulini* Sinitshenkova 1980** Holotype from the Koshelevka Formation at Chekarda on the Sylva River, Perm region, Russia (Sinitshenkova, 1980), of Kungurian (Irenian substage) age

(Lucas & Shen, 2018; Khramov et al., 2023), giving 273.0–283.5 Mya. A Permothemistidae (order Permothemistida).

***Permelmoa magnifica* Prokop & Nel 2011** Holotype from locality F34, Les Vignasses, Mérifons Member of the Salagou Formation, Lodève Basin, southern France (Prokop & Nel, 2011), of late Kungurian age, younger than the 282.86 Mya tuff dated in the underlying Octon Member (Michel et al., 2015), giving 272.95–282.86 Mya. A Parelmoidae (order Diaphanopterodea).

***Permiakovia quinquefasciata* (Martynov 1940)** Holotype from the Koshelevka Formation at Chekarda on the Sylva River, Perm region, Russia (Martynov, 1940), of Kungurian (Irenian substage) age (Lucas & Shen, 2018; Khramov et al., 2023), giving 273.0–283.5 Mya. A Spilapteridae (order Palaeodictyoptera).

***Permodiapha carpenteri* Kukalová-Peck 1974** Holotype from the upper Letovice Formation at Obora, southern Moravia (Kukalová-Peck, 1974), of Sakmarian–Artinskian (Early Permian) age (Opluštil et al., 2017), giving 283.30–293.52 Mya. A diaphanopterodean of uncertain family (order Diaphanopterodea).

***Permodiapha lata* Kukalová-Peck 1974** Holotype from the upper Letovice Formation at Obora, southern Moravia (Kukalová-Peck, 1974), of Sakmarian–Artinskian (Early Permian) age (Opluštil et al., 2017), giving 283.30–293.52 Mya. A diaphanopterodean of uncertain family (order Diaphanopterodea).

***Permodiapha specula* Kukalová-Peck 1974** Holotype from the upper Letovice Formation at Obora, southern Moravia (Kukalová-Peck, 1974), of Sakmarian–Artinskian (Early Permian) age (Opluštil et al., 2017), giving 283.30–293.52 Mya. A diaphanopterodean of uncertain family (order Diaphanopterodea).

***Permohymen schucherti* Tillyard 1924** Holotype from the Wellington Formation at Elmo, Kansas (Tillyard, 1924), of Leonardian (late Artinskian–early Kungurian) age (Tasch & Zimmerman, 1962; Schneider et al., 2020), giving 279–286 Mya. A Protohymenidae (order Megasecoptera).

***Permothemidia caudata* Rohdendorf 1940** Holotype from the Koshelevka Formation at Chekarda on the Sylva River, Perm region, Russia (Rohdendorf, 1940b), of Kungurian (Irenian substage) age (Lucas & Shen, 2018; Khramov et al., 2023), giving 273.0–283.5 Mya. A Permothemistidae (order Permothemistida).

***Permothemis flexus* Sinitshenkova 1980** Holotype from the Iva-Gora Beds at Soyana on the Soyana River, Arkhangelsk Region, Russia (Sinitshenkova, 1980), of Roadian (Kazanian) age (Naugolnykh, 2015; Davydov et al., 2018), giving 266.9–277.0 Mya. A Permothemistidae (order Permothemistida).

***Permothemis intermedius* Sinitshenkova 1980** Holotype from the Iva-Gora Beds at Soyana on the Soyana River, Arkhangelsk Region, Russia (Sinitshenkova, 1980), of Roadian (Kazanian) age (Naugolnykh, 2015; Davydov et al., 2018), giving 266.9–277.0 Mya. A Permothemistidae (order Permothemistida).

***Permothemis lepidus* Sinitshenkova 1980** Holotype from the Iva-Gora Beds at Soyana on the Soyana River, Arkhangelsk Region, Russia (Sinitshenkova, 1980), of Roadian (Kazanian) age (Naugolnykh, 2015; Davydov et al., 2018), giving 266.9–277.0 Mya. A Permothemistidae (order Permothemistida).

***Permothemis libelluloides* (Martynov 1932)** Holotype from the Iva-Gora Beds at Soyana on the Soyana River, Arkhangelsk Region, Russia (Martynov, 1932), of Roadian (Kazanian) age (Naugolnykh, 2015; Davydov et al., 2018), giving 266.9–277.0 Mya. A Permothemistidae (order Permothemistida).

***Permothemis retrocristatus* Sinitshenkova 1980** Holotype from the Iva-Gora Beds at Soyana on the Soyana River, Arkhangelsk Region, Russia (Sinitshenkova, 1980), of Roadian (Kazanian) age (Naugolnykh, 2015; Davydov et al., 2018), giving 266.9–277.0 Mya. A Permothemistidae (order Permothemistida).

***Permuralia maculata* (Kukalová-Peck & Sinichenkova 1992)** Holotype from the Koshelevka Formation at Chekarda on the Sylva River, Perm region, Russia (Kukalová-Peck & Sinichenkova, 1992), of Kungurian (Irenian substage) age (Lucas & Shen, 2018; Khramov et al., 2023), giving 273.0–283.5 Mya. A Parelmoidae (order Diaphanopterodea).

***Permuralia sharovi* (Kukalová-Peck & Sinichenkova 1992)** Holotype from the Koshelevka Formation at Chekarda on the Sylva River, Perm region, Russia (Kukalová-Peck & Sinichenkova, 1992), of Kungurian (Irenian substage) age (Lucas & Shen, 2018; Khramov et al., 2023), giving 273.0–283.5 Mya. A Parelmoidae (order Diaphanopterodea).

***Peromaptera filholi* Brongniart 1894** Holotype from the bituminous shale of the Commentry coal basin at Commentry, Allier, France (Brongniart, 1894), of Gzhelian (Stephanian B) age (Gradstein et al., 2020), giving 298.9–303.7 Mya. An Eugereonidae (order Palaeodictyoptera).

***Phaneroneura martynovae* Carpenter 1947** Holotype from the Midco Member of the Wellington Formation at Midco locality 15, Oklahoma (Carpenter, 1947), of Leonardian (late Artinskian–early Kungurian) age (Tasch & Zimmerman, 1962; Schneider et al., 2020), giving 279–286 Mya. A Martynoviidae (order Diaphanopterodea).

***Phaneroneura minuta* Béthoux et al. 2003** Holotype from locality F28, Notre Dame des Clans, Octon Member of the Salagou Formation, Lodève Basin, southern France (Béthoux et al., 2003), of latest Artinskian to earliest Kungurian age (Michel et al., 2015), giving 282.9–284.6 Mya. A Martynoviidae (order Diaphanopterodea).

***Phaneroneura reducta* Carpenter 1947** Holotype from the Midco Member of the Wellington Formation at Midco locality 15, Oklahoma (Carpenter, 1947), of Leonardian (late Artinskian–early Kungurian) age (Tasch & Zimmerman, 1962; Schneider et al., 2020), giving 279–286 Mya. A Martynoviidae (order Diaphanopterodea).

***Phaneroneura rineharti* Sinitshenkova 2004** Holotype from the Red Tanks Member of the Bursum Formation at Carrizo Arroyo, New Mexico (Rasnitsyn et al., 2004), of latest Gzhelian–earliest Asselian (Carboniferous–Permian boundary) age (Schneider et al., 2020), giving 296.5–299.5 Mya. A Martynoviidae (order Diaphanopterodea).

***Pharciphyzelus lacefieldi* Beckemeyer and Engel 2011** Holotype from the Pottsville Formation at the Haley Brothers and Barbour coal mine, Alabama, of Langsettian (Westphalian A) age (Beckemeyer & Engel, 2011), giving 315.2–319.0 Mya. A Homoiopteridae (order Palaeodictyoptera).

***Philiasptilon maculosum* Zalessky 1931** Holotype from the Alykaeva Formation at Cheremichkino, Kuznetsk Basin (Zalessky, 1931), of Gzhelian age (Silantiev et al., 2025), giving 298.9–303.7 Mya. A Diaphanopteridae (order Diaphanopterodea).

***Piesbergala leipnerae* Brauckmann & Herd 2002** Holotype from the Flöz Dreibänke at the Piesberg quarry near Osnabrück, Lower Saxony, Germany (Brauckmann & Herd, 2002), of Westphalian D (Asturian) age (Gradstein et al., 2020; Schneider et al., 2020), giving 307.50–310.70 Mya. A Parabrodiidae (order Diaphanopterodea).

***Piesbergbrodia tristrata*** **(Brauckmann & Herd 2002)** Holotype from the Flöz Dreibänke at the Piesberg quarry near Osnabrück, Lower Saxony, Germany (Brauckmann & Herd, 2002), of Westphalian D (Asturian) age (Gradstein et al., 2020; Schneider et al., 2020), giving 307.50–310.70 Mya. A Brodiidae (order Megasecoptera).

***Piesbergptera niesseae*** **Rosová et al. 2022** Holotype from a lacustrine horizon at the Piesberg quarry near Osnabrück, Lower Saxony, Germany (Rosová et al., 2022), of Westphalian D (Asturian) age (Gradstein et al., 2020; Schneider et al., 2020), giving 307.50–310.70 Mya. A Piesbergpteridae (order Megasecoptera).

***Platephemera antiqua*** **Scudder 1867** Holotype from Plant Bed 7 of the Fern Ledges (Lancaster Formation) at Carleton, Saint John, New Brunswick, Canada (Scudder, 1867), of Langsettian (Westphalian A) age (Gradstein et al., 2020; Schneider et al., 2020), giving 316.5–319.2 Mya. A palaeodictyopteran of uncertain family (order Palaeodictyoptera).

***Polioptenus bertrandi*** **Waterlot 1934** Holotype from the Wendel pit (vein 8, étage 432) of the Petite-Rosselle colliery, Moselle, France (Waterlot, 1934), of Westphalian D (Asturian) age (Gradstein et al., 2020; Schneider et al., 2020), giving 307.5–310.7 Mya. A Dictyoneuridae (order Palaeodictyoptera).

***Polioptenus chanderisi*** **(Waterlot 1934)** Holotype from the Geisheck Formation at Jägersfreude (vein d), Saarbrücken, Saarland, Germany (Waterlot, 1934), of Westphalian D (Asturian) age (Gradstein et al., 2020; Schneider et al., 2020), giving 307.5–310.7 Mya. A Dictyoneuridae (order Palaeodictyoptera).

***Polioptenus decheni*** **(Goldenberg 1854)** Holotype from the Lower Saarbrücken layers at the Altenwald mine, Saarland, Germany (Goldenberg, 1854), of Westphalian C (Bolsovian) age (Gradstein et al., 2020; Schneider et al., 2020), giving 310.7–313.7 Mya. A Dictyoneuridae (order Palaeodictyoptera).

***Polioptenus elegans*** **(Goldenberg 1877)** Holotype from the Lower Saarbrücken layers at the mine dump near Dudweiler station, Saarland, Germany (Goldenberg, 1877), of Westphalian C (Bolsovian) age (Gradstein et al., 2020; Schneider et al., 2020), giving 310.7–313.7 Mya. A Dictyoneuridae (order Palaeodictyoptera).

***Polioptenus mancomediali* Guthörl 1940** Holotype from the Geisheck Formation at the Frankenholz-Saar colliery (shaft 5), Saarland, Germany (Guthörl, 1940), of Westphalian D (Asturian) age (Gradstein et al., 2020; Schneider et al., 2020), giving 307.5–310.7 Mya. A Dictyoneuridae (order Palaeodictyoptera).

***Polycreagra elegans* Handlirsch 1906** Holotype from the Rhode Island Formation at Cranston, Rhode Island (Handlirsch, 1906b), of Bolsovian–Stephanian (Westphalian C to Stephanian B/C) age (Getty et al., 2017), giving 300.0–315.0 Mya. A Polycreagridae (order Palaeodictyoptera).

***Prochoroptera calopteryx* Handlirsch 1911** Holotype from the Francis Creek Shale (Carbondale Formation) at Mazon Creek, Illinois (Handlirsch, 1911), of Moscovian (Westphalian D) age (Peppers, 1996), giving 306.0–311.0 Mya. A Prochoropteridae (order Diaphanopterodea).

***Propalingenia feistmanteli* (Frič 1880)** Holotype (Me 119) from the Radnice Member of the Kladno Formation on the spoil heap near Otvovice, Kladno–Rakovník Basin, central Bohemia, Czech Republic (Fritsch, 1880), of Bolsovian (Duckmantian) age (Opluštil et al., 2016), giving 313.6–314.5 Mya. A palaeodictyopteran of uncertain family (order Palaeodictyoptera).

***Protagrion audouini* Brongniart 1894** Holotype from the Commentry Shales at Commentry, Allier, France (Brongniart, 1894), of Gzhelian (Stephanian B) age (Gradstein et al., 2020), giving 298.9–303.7 Mya. A Protagriidae (order Palaeodictyoptera).

***Protodiapha lineata* Kukalová-Peck 1974** Holotype from the upper Letovice Formation at Obora, southern Moravia (Kukalová-Peck, 1974), of Sakmarian–Artinskian (Early Permian) age (Opluštil et al., 2017), giving 283.30–293.52 Mya. A diaphanopterodean of uncertain family (order Diaphanopterodea).

***Protodiapha maculifera* Kukalová-Peck 1974** Holotype from the upper Letovice Formation at Obora, southern Moravia (Kukalová-Peck, 1974), of Sakmarian–Artinskian (Early Permian) age (Opluštil et al., 2017), giving 283.30–293.52 Mya. A diaphanopterodean of uncertain family (order Diaphanopterodea).

***Protohymen angustipennis* (Martynov 1932)** Holotype from the Koshelevka Formation at Posadskii Log on the Sylva River, Perm region, Russia (Martynov, 1932), of Kungurian (Irenian substage) age (Lucas & Shen, 2018; Khramov et al., 2023), giving 273.0–283.5 Mya. A Protohymenidae (order Megasecoptera).

***Protohymen anomalus* Zimmerman 1962** Holotype from the *Asthenohymen–Delopterum* bed of the Wellington Formation at Tasch's Kay II locality (bed 6), Oklahoma, of Leonardian (late Artinskian–early Kungurian) age (Tasch & Zimmerman, 1962; Schneider et al., 2020), giving 279–286 Mya. A Protohymenidae (order Megasecoptera).

***Protohymen bifurcatus* Zimmerman 1962** Holotype from the *Asthenohymen–Delopterum* bed of the Wellington Formation at Tasch's Nobel V locality (bed 7), Oklahoma, of Leonardian (late Artinskian–early Kungurian) age (Tasch & Zimmerman, 1962; Schneider et al., 2020), giving 279–286 Mya. A Protohymenidae (order Megasecoptera).

***Protohymen carpenteri* (Novokshonov 1995)** Holotype from the Koshelevka Formation at Chekarda on the Sylva River, Perm region, Russia (Novokshonov, 1995), of Kungurian (Irenian substage) age (Lucas & Shen, 2018; Khramov et al., 2023), giving 273.0–283.5 Mya. A Protohymenidae (order Megasecoptera).

***Protohymen concinna* (Zalessky 1956)** Holotype from the terrigenous beds of the Barda basin at Krasnaya Glinka, Perm region, Russia (Zalessky, 1956), of Kungurian (Irenian substage) age (Lucas & Shen, 2018; Khramov et al., 2023), giving 273.0–283.5 Mya. A Protohymenidae (order Megasecoptera).

***Protohymen curvatus* Carpenter 1947** Holotype from the Midco Member of the Wellington Formation at Midco locality 8, Oklahoma (Carpenter, 1947), of Leonardian (late Artinskian–early Kungurian) age (Tasch & Zimmerman, 1962; Schneider et al., 2020), giving 279–286 Mya. A Protohymenidae (order Megasecoptera).

***Protohymen elongatus* Carpenter 1930** Holotype from the Wellington Formation at Elmo, Kansas (Carpenter, 1930), of Leonardian (late Artinskian–early Kungurian) age (Tasch & Zimmerman, 1962; Schneider et al., 2020), giving 279–286 Mya. A Protohymenidae (order Megasecoptera).

***Protohymen largus* Carpenter 1947** Holotype from the Midco Member of the Wellington Formation at Midco locality 15, Oklahoma (Carpenter, 1947), of Leonardian (late Artinskian–early Kungurian) age (Tasch & Zimmerman, 1962; Schneider et al., 2020), giving 279–286 Mya. A Protohymenidae (order Megasecoptera).

***Protohymen latus* Carpenter 1947** Holotype from the Midco Member of the Wellington Formation at Midco locality 8, Oklahoma (Carpenter, 1947), of Leonardian (late Artinskian–early Kungurian) age (Tasch & Zimmerman, 1962; Schneider et al., 2020), giving 279–286 Mya. A Protohymenidae (order Megasecoptera).

***Protohymen novokshonovi* Pecharová & Prokop 2018** Holotype from the Koshelevka Formation at Chekarda on the Sylva River, Perm region, Russia (Pecharová & Prokop, 2018), of Kungurian (Irenian substage) age (Lucas & Shen, 2018; Khramov et al., 2023), giving 273.0–283.5 Mya. A Protohymenidae (order Megasecoptera).

***Protohymen permianus* Tillyard 1924** Holotype from the Wellington Formation at Elmo, Kansas (Tillyard, 1924), of Leonardian (late Artinskian–early Kungurian) age (Tasch & Zimmerman, 1962; Schneider et al., 2020), giving 279–286 Mya. A Protohymenidae (order Megasecoptera).

***Protohymen pictus* Zimmerman 1962** Holotype from the *Asthenohymen–Delopterum* bed of the Wellington Formation at Tasch's Kay II locality (bed 6), Oklahoma, of Leonardian (late Artinskian–early Kungurian) age (Tasch & Zimmerman, 1962; Schneider et al., 2020), giving 279–286 Mya. A Protohymenidae (order Megasecoptera).

***Protohymen readi* Carpenter 1933** Holotype from the upper limestone of the Wellington Formation at Elmo, Kansas (Carpenter, 1933), of Leonardian (late Artinskian–early Kungurian) age (Tasch & Zimmerman, 1962; Schneider et al., 2020), giving 279–286 Mya. A Protohymenidae (order Megasecoptera).

***Protohymen shafferi* Zimmerman 1962** Holotype from the *Asthenohymen–Delopterum* bed of the Wellington Formation at Tasch's Kay II locality (bed 6), Oklahoma, of Leonardian (late Artinskian–early Kungurian) age (Tasch & Zimmerman, 1962; Schneider et al., 2020), giving 279–286 Mya. A Protohymenidae (order Megasecoptera).

***Protohymen tenuis* Carpenter 1939** Holotype from the lower limestone of the Wellington Formation at Elmo, Kansas (Carpenter, 1939), of Leonardian (late Artinskian–early Kungurian) age (Tasch & Zimmerman, 1962; Schneider et al., 2020), giving 279–286 Mya. A Protohymenidae (order Megasecoptera).

***Protohymen venustus* Carpenter 1947** Holotype from the Midco Member of the Wellington Formation at Midco locality 15, Oklahoma (Carpenter, 1947), of Leonardian (late Artinskian–early

Kungurian) age (Tasch & Zimmerman, 1962; Schneider et al., 2020), giving 279–286 Mya. A Protohymenidae (order Megasecoptera).

***Pseudelmoa ampla* Carpenter 1947** Holotype from the Midco Member of the Wellington Formation at Midco locality 15, Oklahoma (Carpenter, 1947), of Leonardian (late Artinskian–early Kungurian) age (Tasch & Zimmerman, 1962; Schneider et al., 2020), giving 279–286 Mya. A Parelmoidae (order Diaphanopterodea).

***Pseudohymen minor* (Zalessky 1956)** Holotype from the Koshelevka Formation at Chekarda on the Sylva River, Perm region, Russia (Zalessky, 1956), of Kungurian (Irenian substage) age (Lucas & Shen, 2018; Khramov et al., 2023), giving 273.0–283.5 Mya. A Protohymenidae (order Megasecoptera).

***Pseudohymen sylvaensis* (Martynov 1940)** Holotype from the Koshelevka Formation at Chekarda on the Sylva River, Perm region, Russia (Martynov, 1940), of Kungurian (Irenian substage) age (Lucas & Shen, 2018; Khramov et al., 2023), giving 273.0–283.5 Mya. A Protohymenidae (order Megasecoptera).

***Pseudomecynostoma dubium* Handlirsch 1919** Holotype from the bituminous shale of the Commentry coal basin at Commentry, Allier, France (Handlirsch, 1919), of Gzhelian (Stephanian B) age (Gradstein et al., 2020), giving 298.9–303.7 Mya. A palaeodictyopteran of uncertain family (order Palaeodictyoptera).

***Psilothorax longicauda* (Brongniart 1894)** Holotype from the Commentry Shales at Commentry, Allier, France (Brongniart, 1894), of Gzhelian (Stephanian B) age (Gradstein et al., 2020), giving 298.9–303.7 Mya. A Mischopteridae (order Megasecoptera).

***Psychroptilus burrettae* Riek 1976** Holotype from the Wynyard Formation at Hellyer Gorge, Tasmania, Australia (Riek, 1976a), of latest Carboniferous to Early Permian age (Ives et al., 2022), giving 295.0–303.7 Mya. A Psychroptilidae (order Palaeodictyoptera).

***Pteronidia plicatula* Bolton 1912** Holotype from the Middle Coal Measures at Shipley, Derbyshire, England (Bolton, 1912), of Duckmantian (Westphalian B) age (Gradstein et al., 2020; Schneider et al., 2020), giving 313.7–316.5 Mya. A palaeodictyopteran of uncertain family (order Palaeodictyoptera).

***Pyebrodia martinsnetoi* Brauckmann & Herd 2002** Holotype from the Flöz Dreibänke at the Piesberg quarry near Osnabrück, Lower Saxony, Germany (Brauckmann & Herd, 2002), of Westphalian D (Asturian) age (Gradstein et al., 2020; Schneider et al., 2020), giving 307.50–310.70 Mya. A palaeodictyopteran of uncertain family (order Palaeodictyoptera).

***Rhabdoptilus edwardsi* Brongniart 1894** Holotype from the bituminous shale of the Commentry coal basin at Commentry, Allier, France (Brongniart, 1894), of Gzhelian (Stephanian B) age (Gradstein et al., 2020), giving 298.9–303.7 Mya. A Graphiptilidae (order Palaeodictyoptera).

***Rhaphidiopsis diversipenna* Scudder 1893** Holotype from the Rhode Island Formation at the Lockonosett Mine, Cranston, Rhode Island (Scudder, 1893), of Bolsovian–Stephanian (Westphalian C to Stephanian B/C) age (Getty et al., 2017), giving 300.0–315.0 Mya. A Rhaphidiopsidae (order Diaphanopterodea).

***Rochdalia parkeri* Woodward 1913** Holotype from the Middle Coal Measures at Sparth Bottoms, Rochdale, Lancashire, England (Woodward, 1913), of Langsettian (Westphalian A) age (Gradstein et al., 2020; Schneider et al., 2020), giving 316.5–319.2 Mya. A palaeodictyopteran of uncertain family (order Palaeodictyoptera).

***Rotundopteris multimediali* Guthörl 1940** Holotype from the Geisheck Formation at the Frankenholz-Saar colliery (shaft 5), Saarland, Germany (Guthörl, 1940), of Westphalian D (Asturian) age (Gradstein et al., 2020; Schneider et al., 2020), giving 307.5–310.7 Mya. A Dictyoneuridae (order Palaeodictyoptera).

***Saarlandia flexisubcostata* Guthörl 1930** Holotype from the Friedrichsthal railway cutting, Saarbrücken district, Saarland, Germany (Guthörl, 1930), of Westphalian age (Gradstein et al., 2020; Schneider et al., 2020), giving 307.5–319.2 Mya. A palaeodictyopteran of uncertain family (order Palaeodictyoptera).

***Saarodictyum abnormis* Guthörl 1939** Holotype from the Sulzbach Beds at the Grube Friedrichsthal-Saar (Erkershöhe shaft), Saarbrücken, Saarland, Germany (Guthörl, 1939), of Westphalian C (Bolsovian) age (Gradstein et al., 2020; Schneider et al., 2020), giving 310.7–313.7 Mya. A Dictyoneuridae (order Palaeodictyoptera).

***Sabitaptus lagagei* Pruvost 1930** Holotype from the roof of vein A at the Fontaine-l'Évêque coal mine, Fontaine-l'Évêque, Hainaut, Belgium (Pruvost, 1930), of Duckmantian (Westphalian B) age (Gradstein et al., 2020; Schneider et al., 2020), giving 313.7–316.5 Mya. A palaeodictyopteran of uncertain family (order Palaeodictyoptera).

***Salagouneura chimaira* Béthoux et al. 2003** Holotype from locality F28, Notre Dame des Clans, Octon Member of the Salagou Formation, Lodève Basin, southern France (Béthoux et al., 2003), of latest Artinskian to earliest Kungurian age (Michel et al., 2015), giving 282.9–284.6 Mya. A Martynoviidae (order Diaphanopterodea).

***Sandiella herbigi* Brauckmann et al. 2023** Holotype from a lacustrine horizon at the Piesberg quarry near Osnabrück, Lower Saxony, Germany (Brauckmann et al., 2023), of Westphalian D (Asturian) age (Gradstein et al., 2020; Schneider et al., 2020), giving 307.50–310.70 Mya. An Eugereonidae (order Palaeodictyoptera).

***Sandiella readi* Carpenter 1970** Holotype from the Sandia Formation at Santa Fe Creek (USGS locality 8941), New Mexico (Carpenter, 1970), of Atokan (latest Bashkirian–early Moscovian) age (Krainer et al., 2011), giving 311.0–316.0 Mya. A Eugereonidae (order Palaeodictyoptera).

***Saxonyptilus sowiaki* Rosová et al. 2022** Holotype from the Flöz Dreibänke at the Piesberg quarry near Osnabrück, Lower Saxony, Germany (Rosová et al., 2022), of Westphalian D (Asturian) age (Gradstein et al., 2020; Schneider et al., 2020), giving 307.50–310.70 Mya. A Spilapteridae (order Palaeodictyoptera).

***Scepasma gigas* Handlirsch 1911** Holotype (Peabody Museum, Yale University, no. 22) from the Francis Creek Shale (Carbondale Formation) at Mazon Creek, Illinois (Handlirsch, 1911), of Moscovian (Westphalian D) age (Gradstein et al., 2020; Schneider et al., 2020), giving 307.5–310.7 Mya. A Homoiopteridae (order Palaeodictyoptera).

***Scepasma mediomatricorum* Brauckmann & Becker 1992** Holotype from the Sulzbach Beds of the Saarland, Germany (Brauckmann & Becker, 1992), of Westphalian C (Bolsovian) age (Gradstein et al., 2020; Schneider et al., 2020), giving 310.7–313.7 Mya. A Homoiopteridae (order Palaeodictyoptera).

***Schedoneura amii* (Copeland 1957)** Holotype from Howard's Mills, River Wallace, Cumberland County, Nova Scotia (Copeland, 1957), of Langsettian (Westphalian A) age (Davies et al., 2005), giving

315.2–318.7 Mya. A palaeodictyopteran of uncertain family (order Palaeodictyoptera).

***Schmidtopteron adictyon* Brauckmann & Hahn 1978** Holotype from the Upper Hagen Beds at the Schmiedestraße quarry, Haßlinghausen, North Rhine-Westphalia, Germany (Brauckmann & Hahn, 1978), of late Namurian (Marsdenian) age (Gradstein et al., 2020; Opluštil et al., 2022), giving 319.50–320.50 Mya. A Dictyoneuridae (order Palaeodictyoptera).

***Scytohymen extremus* Martynov 1937** Holotype from the Savinovski gully near the village of Savino-Potukhovo (Martynov, 1937), of Kungurian age (Gradstein et al., 2020), giving 273.0–283.3 Mya. A Scytohymenidae (order Megasecoptera).

***Severinopsis vetusta* Kukalová 1958** Holotype from coal seam 31 (Upper Sucha Beds) at the Gottwald (František) Mine, Horní Suchá, Karviná, Czech Republic (Kukalová, 1958), of Westphalian A (Langsettian) age (Gradstein et al., 2020; Schneider et al., 2020), giving 316.5–319.2 Mya. A palaeodictyopteran of uncertain family (order Palaeodictyoptera).

***Sharovia sojanica* Sinitshenkova 1977** Holotype from the Iva-Gora Beds at Soyana on the Soyana River, Arkhangelsk Region, Russia (Sharov & Sinitshenkova, 1977), of Roadian (Kazanian) age (Naugolnykh, 2015; Davydov et al., 2018), giving 266.9–277.0 Mya. A Calvertiellidae (order Palaeodictyoptera).

***Sharovia tshekardica* Ivanov 2000** Holotype from the Koshelevka Formation at Chekarda on the Sylva River, Perm region, Russia (Ivanov, 2000), of Kungurian (Irenian substage) age (Lucas & Shen, 2018; Khramov et al., 2023), giving 273.0–283.5 Mya. A Calvertiellidae (order Palaeodictyoptera).

***Siberiodictya gigas* Sharov & Sinitshenkova 1977** Holotype from the Burguklinskaya Formation on the left bank of the Lower Tunguska River, Tunguska Basin (Sharov & Sinitshenkova, 1977), of Sakmarian age (Budnikov et al., 2020), giving 290.1–293.52 Mya. A Dictyoneuridae (order Palaeodictyoptera).

***Siberiohymen asiaticus* Rohdendorf 1961** Holotype from the Alykaeva Formation at Verkhotomskoye, Kuznetsk Basin (PIN collection 742) (Rohdendorf et al., 1961), of Gzhelian age (Silantiev et al., 2025), giving 298.9–303.7 Mya. A Vorkutiidae (order Megasecoptera).

***Sinitshenkovae gallica* Nel et al. 2019** Holotype from locality F23, Les Canals, Mérifons Member of the Salagou Formation, Lodève Basin, southern France (Nel et al., 2019), of late Kungurian age, younger than the 282.86 Mya tuff dated in the underlying Octon Member (Michel et al., 2015), giving 272.95–282.86 Mya. A Xenopteraidae (order Megasecoptera).

***Sinitshenkovae huenickeni* (Pinto & de Ornellas 1978)** Holotype from the Pallero Member of the Bajo de Véliz Formation at Bajo de Véliz, San Luis Province, Argentina (Pinto & de Ornellas, 1978), of Gzhelian–Asselian age (Césari & Chiesa, 2017), giving 295.0–303.7 Mya. A Xenopteraidae (order Megasecoptera).

***Sinodiapha ramosa* Yang et al. 2020** Holotype from the Yanghugou Formation at Xiaheyan, Zhongwei, Ningxia, China (Yang et al., 2020), of latest Bashkirian to middle Moscovian age (Trümper et al., 2020), giving 309.0–316.0 Mya. A Sinodiaphidae (order Diaphanopterodea).

***Sinodunbaria jarmilae* Li et al. 2013** Holotype from the Yanghugou Formation at Xiaheyan, Zhongwei, Ningxia, China (Li et al., 2013), of latest Bashkirian to middle Moscovian age (Trümper et al., 2020), giving 309.0–316.0 Mya. A Spilapteridae (order Palaeodictyoptera).

***Sinoelmoa yangquanensis* Yang et al. 2024** Holotype from the Shanxi Formation at Shuiquan Gully, Yangquan City, Shanxi Province, China (Yang et al., 2024), of mid-Asselian age (Wu et al., 2021; Santos et al., 2024), giving 294.0–297.0 Mya. A Parelmoidae (order Diaphanopterodea).

***Sinohymen gancaoshanensis* Hong 1985** Holotype from the Shanxi Formation at Renmazhuang, Xiangning County, Shanxi Province, China (Hong, 1985b), of mid-Asselian age (Wu et al., 2021; Santos et al., 2024), giving 294.0–297.0 Mya. A Bardohymenidae (order Megasecoptera).

***Sinopalaeopteryx olivieri* Pecharová et al. 2015** Holotype from the Yanghugou Formation at Xiaheyan, Zhongwei, Ningxia, China (Pecharová et al., 2015a), of latest Bashkirian to middle Moscovian age (Trümper et al., 2020), giving 309.0–316.0 Mya. An Aykhalidae (order Megasecoptera).

***Sinopalaeopteryx splendens* Pecharová et al. 2015** Holotype from the Yanghugou Formation at Xiaheyan, Zhongwei, Ningxia, China (Pecharová et al., 2015a), of latest Bashkirian to middle Moscovian age (Trümper et al., 2020), giving 309.0–316.0 Mya. An Aykhalidae (order Megasecoptera).

***Sphecoptera brongniarti* Meunier 1908** Holotype from the Commentry Shales at Commentry, Allier, France (Meunier, 1908c), of Gzhelian (Stephanian B) age (Gradstein et al., 2020), giving 298.9–303.7 Mya. A Sphecopteridae (order Megasecoptera).

***Sphecoptera gracilis* Brongniart 1894** Holotype from the Commentry Shales at Commentry, Allier, France (Brongniart, 1894), of Gzhelian (Stephanian B) age (Gradstein et al., 2020), giving 298.9–303.7 Mya. A Sphecopteridae (order Megasecoptera).

***Sphecoptera minor* (Brongniart 1894)** Holotype from the Commentry Shales at Commentry, Allier, France (Brongniart, 1894), of Gzhelian (Stephanian B) age (Gradstein et al., 2020), giving 298.9–303.7 Mya. A Sphecopteridae (order Megasecoptera).

***Sphecoptera pulchra* Brongniart 1894** Holotype from the Commentry Shales at Commentry, Allier, France (Brongniart, 1894), of Gzhelian (Stephanian B) age (Gradstein et al., 2020), giving 298.9–303.7 Mya. A Sphecopteridae (order Megasecoptera).

***Sphecorydaloides lucchesei* Pinto 1994** Holotype from the Pallero Member of the Bajo de Véliz Formation at Bajo de Véliz, San Luis Province, Argentina (Pinto, 1994), of Gzhelian–Asselian age (Césari & Chiesa, 2017), giving 295.0–303.7 Mya. A Sphecorydaloididae (order Megasecoptera).

***Spilaptera americana* Carpenter and Richardson 1971** Holotype from the Francis Creek Shale (Carbondale Formation) at Pit 6, Coal City, Mazon Creek, Illinois (Carpenter & Richardson, 1971), of Moscovian (Westphalian D) age (Peppers, 1996), giving 306.0–311.0 Mya. A Spilapteridae (order Palaeodictyoptera).

***Spilaptera libelluloides* Brongniart 1894** Holotype from the bituminous shale of the Commentry coal basin at Commentry, Allier, France (Brongniart, 1894), of Gzhelian (Stephanian B) age (Gradstein et al., 2020), giving 298.9–303.7 Mya. A Spilapteridae (order Palaeodictyoptera).

***Spilaptera packardi* Brongniart 1885** Holotype from the bituminous shale of the Commentry coal basin at Commentry, Allier, France (Brongniart, 1885), of Gzhelian (Stephanian B) age (Gradstein et al., 2020), giving 298.9–303.7 Mya. A Spilapteridae (order Palaeodictyoptera).

***Spilaptera peckae* Rosová et al. 2022** Holotype from the Flöz Dreibänke at the Piesberg quarry near Osnabrück, Lower Saxony, Germany (Rosová et al., 2022), of Westphalian D (Asturian) age (Gradstein et al., 2020; Schneider et al., 2020), giving 307.50–310.70 Mya. A Spilapteridae (order Palaeodictyoptera).

***Spilaptera splendens* Prokop et al. 2014** Holotype from the spoil heap of Terril No. 7 at Avion, Pas-de-Calais, France (Prokop et al., 2014), of Bolsovian (Westphalian C) age (Gradstein et al., 2020;

Schneider et al., 2020), giving 310.7–313.7 Mya. A Spilapteridae (order Palaeodictyoptera).

***Spilaptera tanaica* Sharov & Sinitshenkova 1977** Holotype from the Isaeva Formation at Alymaznaya on the Lomovatka River, Tunguska Basin (PIN collection 1866) (Sharov & Sinitshenkova, 1977), of Kasimovian age (Gradstein et al., 2020), giving 303.7–307.0 Mya. A Spilapteridae (order Palaeodictyoptera).

***Spilaptera venusta* (Brongniart 1894)** Holotype from the bituminous shale of the Commentry coal basin at Commentry, Allier, France (Brongniart, 1894), of Gzhelian (Stephanian B) age (Gradstein et al., 2020), giving 298.9–303.7 Mya. A Spilapteridae (order Palaeodictyoptera).

***Spiloptilus ramondi* (Brongniart 1894)** Holotype from the Commentry Shales at Commentry, Allier, France (Brongniart, 1894), of Gzhelian (Stephanian B) age (Gradstein et al., 2020), giving 298.9–303.7 Mya. A Spilapteridae (order Palaeodictyoptera).

***Stenodiapha angusta* Kukalová-Peck 1974** Holotype from the upper Letovice Formation at Obora, southern Moravia (Kukalová-Peck, 1974), of Sakmarian–Artinskian (Early Permian) age (Opluštil et al., 2017), giving 283.30–293.52 Mya. A diaphanopterodean of uncertain family (order Diaphanopterodea).

***Stenodiapha moravica* Kukalová-Peck 1974** Holotype from the upper Letovice Formation at Obora, southern Moravia (Kukalová-Peck, 1974), of Sakmarian–Artinskian (Early Permian) age (Opluštil et al., 2017), giving 283.30–293.52 Mya. A diaphanopterodean of uncertain family (order Diaphanopterodea).

***Stenodictya agnita* (Meunier 1908)** Holotype from the Commentry Shales at Commentry, Allier, France (Meunier, 1908d), of Gzhelian (Stephanian B) age (Gradstein et al., 2020), giving 298.9–303.7 Mya. A Dictyoneuridae (order Palaeodictyoptera).

***Stenodictya arnaudi* Brongniart 1894** Holotype from the Commentry Shales at Commentry, Allier, France (Brongniart, 1894), of Gzhelian (Stephanian B) age (Gradstein et al., 2020), giving 298.9–303.7 Mya. A Dictyoneuridae (order Palaeodictyoptera).

***Stenodictya fritschi* Brongniart 1894** Holotype from the Commentry Shales at Commentry, Allier, France (Brongniart, 1894), of Gzhelian (Stephanian B) age (Gradstein et al., 2020), giving 298.9–303.7 Mya. A Dictyoneuridae (order Palaeodictyoptera).

***Stenodictya grandissima* (Meunier 1911)** Holotype from the Commentry Shales at Commentry, Allier, France (Meunier, 1911), of Gzhelian (Stephanian B) age (Gradstein et al., 2020), giving 298.9–303.7 Mya. A Dictyoneuridae (order Palaeodictyoptera).

***Stenodictya klebsi* (Meunier 1908)** Holotype from the Commentry Shales at Commentry, Allier, France (Meunier, 1908d), of Gzhelian (Stephanian B) age (Gradstein et al., 2020), giving 298.9–303.7 Mya. A Dictyoneuridae (order Palaeodictyoptera).

***Stenodictya laurentiauxi* Kukalová 1970** Holotype from the Commentry Shales at Commentry, Allier, France (Kukalová, 1970), of Gzhelian (Stephanian B) age (Gradstein et al., 2020), giving 298.9–303.7 Mya. A Dictyoneuridae (order Palaeodictyoptera).

***Stenodictya lobata* (Brongniart 1890)** Holotype from the Commentry Shales at Commentry, Allier, France (Brongniart, 1890), of Gzhelian (Stephanian B) age (Gradstein et al., 2020), giving 298.9–303.7 Mya. A Dictyoneuridae (order Palaeodictyoptera).

***Stenodictya lusitanica* Correia et al. 2014** Holotype from São Pedro da Cova, Douro Carboniferous Basin, northern Portugal (Correia et al., 2014), of early Gzhelian age (Correia et al., 2019), giving 300.5–303.7 Mya. A Dictyoneuridae (order Palaeodictyoptera).

***Stenodictya minima* Brongniart 1894** Holotype from the Commentry Shales at Commentry, Allier, France (Brongniart, 1894), of Gzhelian (Stephanian B) age (Gradstein et al., 2020), giving 298.9–303.7 Mya. A Dictyoneuridae (order Palaeodictyoptera).

***Stenodictya oustaleti* Brongniart 1894** Holotype from the Commentry Shales at Commentry, Allier, France (Brongniart, 1894), of Gzhelian (Stephanian B) age (Gradstein et al., 2020), giving 298.9–303.7 Mya. A Dictyoneuridae (order Palaeodictyoptera).

***Stenodictya parisiana* Kukalová 1970** Holotype from the Commentry Shales at Commentry, Allier, France (Kukalová, 1970), of Gzhelian (Stephanian B) age (Gradstein et al., 2020), giving 298.9–303.7 Mya. A Dictyoneuridae (order Palaeodictyoptera).

***Stenodictya perrieri* Brongniart 1894** Holotype from the Commentry Shales at Commentry, Allier, France (Brongniart, 1894), of Gzhelian (Stephanian B) age (Gradstein et al., 2020), giving 298.9–303.7 Mya. A Dictyoneuridae (order Palaeodictyoptera).

***Stenodictya pygmaea* Meunier 1911** Holotype from the Commentry Shales at Commentry, Allier, France (Meunier, 1911), of Gzhelian (Stephanian B) age (Gradstein et al., 2020), giving 298.9–303.7 Mya. A Dictyoneuridae (order Palaeodictyoptera).

***Stenodictya spinosa* (Brongniart 1890)** Holotype from the Commentry Shales at Commentry, Allier, France (Brongniart, 1890), of Gzhelian (Stephanian B) age (Gradstein et al., 2020), giving 298.9–303.7 Mya. A Dictyoneuridae (order Palaeodictyoptera).

***Stenodictya thevenini* Meunier 1908** Holotype from the Commentry Shales at Commentry, Allier, France (Meunier, 1908b), of Gzhelian (Stephanian B) age (Gradstein et al., 2020), giving 298.9–303.7 Mya. A Dictyoneuridae (order Palaeodictyoptera).

***Stenodictya vasseuri* Meunier 1914** Holotype from the Commentry Shales at Commentry, Allier, France (Meunier, 1914), of Gzhelian (Stephanian B) age (Gradstein et al., 2020), giving 298.9–303.7 Mya. A Dictyoneuridae (order Palaeodictyoptera).

***Stenodictyoneura belgica* Leriche 1911** Holotype from the Flénu Formation at the Grand-Hornu colliery, Saint-Ghislain, Hainaut, Belgium (Leriche, 1911), of Bolsovian (Westphalian C) age (Gradstein et al., 2020; Schneider et al., 2020), giving 310.7–313.7 Mya. A Dictyoneuridae (order Palaeodictyoptera).

***Stilbocrocis heeri* (Goldenberg 1854)** Holotype from the Lower Saarbrücken layers at the Altenwald mine, Saarland, Germany (Goldenberg, 1854), of Westphalian C (Bolsovian) age (Gradstein et al., 2020; Schneider et al., 2020), giving 310.7–313.7 Mya. A Dictyoneuridae (order Palaeodictyoptera).

***Stilbocrocis lanceolata* Guthörl 1940** Holotype from the Geisheck Formation at the Frankenholz-Saar colliery (shaft 5), Saarland, Germany (Guthörl, 1940), of Westphalian D (Asturian) age (Gradstein et al., 2020; Schneider et al., 2020), giving 307.5–310.7 Mya. A Dictyoneuridae (order Palaeodictyoptera).

***Stilbocrocis pulchra* (Guthörl 1940)** Holotype from the Geisheck Formation at the Frankenholz-Saar colliery (shaft 5), Saarland, Germany (Guthörl, 1940), of Westphalian D (Asturian) age (Gradstein et al., 2020; Schneider et al., 2020), giving 307.5–310.7 Mya. A Dictyoneuridae (order Palaeodictyoptera).

***Stobbsia woodwardiana* (Handlirsch 1908)** Holotype from the Peacock Marls at Foley, near Longton, North Staffordshire, England (Handlirsch, 1908), of Langsettian (Westphalian A) age (Gradstein et al., 2020; Schneider et al., 2020), giving 316.5–319.2 Mya. A Breyeriidae (order Palaeodictyoptera).

***Straeleniella namurensis* Laurentiaux-Vieira & Laurentiaux 1986** Holotype from the Yeadonian coal measures at the Amercoeur Collieries, Belgium (Laurentiaux-Vieira & Laurentiaux, 1986), of Bashkirian (Yeadonian) age (Gradstein et al., 2020), giving 318.1–319.2 Mya. A Straeleniellidae (order Palaeodictyoptera).

***Sunohymen xishanensis* Hong 1985** Holotype from the Shanxi Formation at Qiligou, Xishan of Taiyuan, Shanxi Province, China (Hong, 1985a), of mid-Asselian age (Wu et al., 2021; Santos et al., 2024), giving 294.0–297.0 Mya. A Bardohymenidae (order Megasecoptera).

***Sylvohymen ingens* Carpenter 1947** Holotype from the Midco Member of the Wellington Formation at Midco locality 3, Oklahoma (Carpenter, 1947), of Leonardian (late Artinskian–early Kungurian) age (Tasch & Zimmerman, 1962; Schneider et al., 2020), giving 279–286 Mya. A Bardohymenidae (order Megasecoptera).

***Sylvohymen marginatus* Pecharová et al. 2020** Holotype from the Koshelevka Formation at Chekarda on the Sylva River, Perm region, Russia (Pecharová et al., 2020), of Kungurian (Irenian substage) age (Lucas & Shen, 2018; Khramov et al., 2023), giving 273.0–283.5 Mya. A Bardohymenidae (order Megasecoptera).

***Sylvohymen peckae* Brauckmann 1988** Holotype from the Vorhalle Beds at the Hagen-Vorhalle quarry, North Rhine-Westphalia, Germany (Brauckmann, 1988), of late Marsdenian (Namurian B, *Bilinguites metabilinguis* Marine Band, Ziegelschiefer Formation) age (Ilger & Brauckmann, 2011; Opluštil et al., 2022), giving 319.0–320.5 Mya. A Bardohymenidae (order Megasecoptera).

***Sylvohymen robustus* Martynov 1940** Holotype from the Koshelevka Formation at Chekarda on the Sylva River, Perm region, Russia (Martynov, 1940), of Kungurian (Irenian substage) age (Lucas & Shen, 2018; Khramov et al., 2023), giving 273.0–283.5 Mya. A Bardohymenidae (order Megasecoptera).

***Sylvohymen sibiricus* Kukalová-Peck 1972** Holotype from the Lower Permian of Siberia (Kukalová-Peck, 1972), of Early Permian (Cisuralian) age (Lucas & Shen, 2018), giving 273.0–298.9 Mya. A

Bardohymenidae (order Megasecoptera).

***Synaptoneura champyi* Pruvost 1919** Holotype from the Cuvinot pit (Faisceau de Cuvinot) at Anzin, Nord, France (Pruvost, 1919), of Duckmantian (Westphalian B) age (Gradstein et al., 2020; Schneider et al., 2020), giving 313.7–316.5 Mya. A palaeodictyopteran of uncertain family (order Palaeodictyoptera).

***Synarmoge ferrarii* Handlirsch 1910** Holotype from the Wendel colliery near Hamm, Westphalia, Germany (Handlirsch, 1910), of Duckmantian (Westphalian B) age (Gradstein et al., 2020; Schneider et al., 2020), giving 313.7–316.5 Mya. A Synarmogidae (order Palaeodictyoptera).

***Taigahymen borealis* Sinitshenkova 2019** Holotype from a depth of 194.7 m in borehole KhK-861 at Khalmer-Yu, in the Inta Formation of the Pechora (Vorkuta) Basin (Sinitshenkova, 2019), of Kungurian age (Davydov et al., 2018), giving 273.0–283.5 Mya. A Bardohymenidae (order Megasecoptera).

***Tchirkovaea guttata* Zalessky 1931** Holotype from the Alykaeva Formation at Cheremichkino, Kuznetsk Basin (Zalessky, 1931), of Gzhelian age (Silantiev et al., 2025), giving 298.9–303.7 Mya. A Tchirkovaeidae (order Palaeodictyoptera).

***Tectoptilus grehanti* (Brongniart 1894)** Holotype from the bituminous shale of the Commentry coal basin at Commentry, Allier, France (Brongniart, 1894), of Gzhelian (Stephanian B) age (Gradstein et al., 2020), giving 298.9–303.7 Mya. A Spilapteridae (order Palaeodictyoptera).

***Thesoneura americana* Carpenter 1943** Holotype from the Francis Creek Shale (Carbondale Formation) at Mazon Creek, Illinois (Carpenter, 1943), of Moscovian (Westphalian D) age (Peppers, 1996), giving 306.0–311.0 Mya. A Homoiopteridae (order Palaeodictyoptera).

***Titanodictya jucunda* (Scudder 1885)** Holotype from the Llewellyn Formation at Campbells Ledge, near Pittston, Pennsylvania, USA (Scudder, 1885), of Late Pennsylvanian (Gzhelian) age (Scudder, 1885), giving 300.5–303.7 Mya. A palaeodictyopteran of uncertain family (order Palaeodictyoptera).

***Titanodictya rowlandi* Sinitshenkova 2004** Holotype from the Red Tanks Member of the Bursum Formation at Carrizo Arroyo, New Mexico (Rasnitsyn et al., 2004), of latest Gzhelian–earliest

Asselian (Carboniferous–Permian boundary) age (Schneider et al., 2020), giving 296.5–299.5 Mya. A palaeodictyopteran of uncertain family (order Palaeodictyoptera).

***Titanophasma fayoli* (Brongniart 1882)** Holotype from the Commentry Shales at Commentry, Allier, France (Brongniart, 1882), of Gzhelian (Stephanian B) age (Gradstein et al., 2020), giving 298.9–303.7 Mya. A palaeodictyopteran of uncertain family (order Palaeodictyoptera).

***Tshekardohymen martynovi* Rohdendorf 1940** Holotype from the Koshelevka Formation at Chekarda on the Sylva River, Perm region, Russia (Rohdendorf, 1940a), of Kungurian (Irenian substage) age (Lucas & Shen, 2018; Khramov et al., 2023), giving 273.0–283.5 Mya. A Scytohymenidae (order Megasecoptera).

***Turnbullia priscillae* Richardson 1956** Holotype from the Francis Creek Shale (Carbondale Formation) near Coal City, Illinois (Richardson, 1956), of Moscovian (Westphalian D) age (Peppers, 1996), giving 306.0–311.0 Mya. A palaeodictyopteran of uncertain family (order Palaeodictyoptera).

***Turneropterum turneri* Kukalová-Peck and Richardson 1983** Holotype from the Francis Creek Shale (Carbondale Formation) near Worthington (Terre Haute locality), Indiana (Kukalová-Peck & Richardson, 1983), of Moscovian (Westphalian D) age (Peppers, 1996), giving 306.0–311.0 Mya. A Homoiopteridae (order Palaeodictyoptera).

***Tytthospilaptera wangae* Liu et al. 2015** Holotype from the Yanghugou Formation at Xiaheyan, Zhongwei, Ningxia, China (Liu et al., 2015), of latest Bashkirian to middle Moscovian age (Trümper et al., 2020), giving 309.0–316.0 Mya. A Spilapteridae (order Palaeodictyoptera).

***Valdeania medeirosi* Teixeira 1941** Holotype from Valdeão, near Valongo, Douro Carboniferous Basin, northern Portugal (Teixeira, 1941), of early Gzhelian age (Correia et al., 2019), giving 300.5–303.7 Mya. An Eugereonidae (order Palaeodictyoptera).

***Velisoptera taschi* Pinto & Adami-Rodrigues 1997** Holotype from the Pallero Member of the Bajo de Véliz Formation at Bajo de Véliz, San Luis Province, Argentina (Pinto & Adami-Rodrigues, 1997), of Gzhelian–Asselian age (Césari & Chiesa, 2017), giving 295.0–303.7 Mya. A Velisopteridae (order Diaphanopterodea).

***Vernooijia harlemensis* (Brauckmann & Gröning 1996)** Holotype from the Coal Measures of the former Laura Coalmine at Eygelshoven, South Limburg, the Netherlands (Brauckmann & Gröning,

1996), of Langsettian (Westphalian A) age (Gradstein et al., 2020; Schneider et al., 2020), giving 316.5–319.2 Mya. A Breyeriidae (order Palaeodictyoptera).

***Vernooijia sassoonae* Prokop et al. 2018** Holotype from the Farrington Formation at the Writhlington Geological Nature Reserve, near Radstock, Somerset, England (Prokop et al., 2018), of Asturian (Westphalian D) age (Gradstein et al., 2020; Schneider et al., 2020), giving 307.5–310.7 Mya. A Breyeriidae (order Palaeodictyoptera).

***Vorkutia dimina* Novokshonov 1998** Holotype from the Koshelevka Formation at Chekarda on the Sylva River, Perm region, Russia (Novokshonov, 1998), of Kungurian (Irenian substage) age (Lucas & Shen, 2018; Khramov et al., 2023), giving 273.0–283.5 Mya. A Vorkutiidae (order Megasecoptera).

***Vorkutia tshernovi* Rohdendorf 1947** Holotype from a depth of 278.5 m in borehole K-138 of the Vorkuta Deposit, in the Rudnitskaya Member of the Lekvorkuta Formation of the Pechora (Vorkuta) Basin (Rohdendorf, 1947), of Kungurian age (Davydov et al., 2018), giving 273.0–283.5 Mya. A Vorkutiidae (order Megasecoptera).

***Vorkutoneura variabilis* Sinitshenkova 1977** Holotype from a depth of 162.8 m in borehole NSK-83 of the Nizhne-Syryaginskoe deposit, in the Rudnitskaya Member of the Lekvorkuta Formation of the Pechora (Vorkuta) Basin (Sharov & Sinitshenkova, 1977), of Kungurian age (Davydov et al., 2018), giving 273.0–283.5 Mya. A Spilapteridae (order Palaeodictyoptera).

***Xenoptera latigra* Petrulevičius & Gutiérrez 2025** Holotype from the Guandacol Formation at Cerro Guandacol, Quebrada de las Libélulas, La Rioja Province, Argentina (Petrulevičius & Gutiérrez, 2025), of Serpukhovian age (Petrulevičius & Gutiérrez, 2025), giving 323.6–326.9 Mya. A Xenopteraidae (order Megasecoptera).

***Xenoptera riojaensis* Pinto 1986** Holotype from the La Divisoria Member of the Malanzán Formation at Malanzán (Cuesta de la Herradura), La Rioja Province, Argentina (Pinto, 1986), of Serpukhovian age (Pérez Loinaze, 2009), giving 323.6–326.9 Mya. A Xenopteraidae (order Megasecoptera).

***Xiaheyanella orta* Fu et al. 2015** Holotype from the Dragon Hill (Dragon 1) outcrop of the Yanghugou Formation at Xiaheyan, Zhongwei, Ningxia, China (Fu et al., 2015), of latest Bashkirian to middle Moscovian age (Trümper et al., 2020), giving 309.0–316.0 Mya. A Calvertiellidae (order Palaeodictyoptera).

### 2 Analysis Settings

#### 2.1 Cetacea

For each of the OU-smoothed $\lambda$ and $\mu$ skylines, we assigned $\theta \sim \text{Normal}(\log 0.2, 1.6^2)$, $\sigma_{\text{OU}} \sim \text{Exp}(10/3)$, and $\nu \sim \text{Exp}(2)$. The fossil sampling rates were independently assigned $\psi_i \sim \text{Exp}(2)$. Each of the six sequence partitions was assigned its own GBM clock partition. Under the gamma-Dirichlet prior, $\bar{r}_{\text{clock}} \sim \text{Gamma}(2, 2000)$ and $\overline{\sigma^2}_{\text{clock}} \sim \text{Gamma}(1, 10)$, while $(r_{\text{clock},j} / \sum_k r_{\text{clock},k})_j$ and $(\sigma^2_{\text{clock},j} / \sum_k \sigma^2_{\text{clock},k})_j$ each followed $\text{Dirichlet}(1, \ldots, 1)$.

We assigned $x_0 \sim \text{TruncNormal}(55, 2^2)$, truncated below at 50 Mya. The potentially oldest sampled cetacean, *Himalayacetus subathuensis*, was assigned a broad age interval of 45.4–52.9 Mya, and the joint model requires $x_0$ to predate its sampled age. Published estimates for the Cetacea–Hippopotamidae divergence, also termed crown Whippomorpha or Cetancodonta, cluster around 54–55 Mya: 53.6 Mya with a 95% HPD of 52.6–54.8 Mya (Zurano et al., 2019), 53.92 Mya (McGowen et al., 2020), and approximately 55 Mya (Meredith et al., 2011). We therefore centered the origin prior at 55 Mya. In a sensitivity analysis, we replaced this prior with $x_0 - 46 \sim \text{Exp}(0.1)$, offset at 46 Mya, the largest minimum age among the sampled cetaceans (*Pakicetus* spp. and *Nalacetus ratimitus*, 46.0–49.0 Mya), below which the origin cannot lie.

#### 2.2 Emydidae

The constant FBD rates were assigned mutually independent $\lambda, \mu, \psi \sim \text{Exp}(10)$ priors. Each of the 15 sequence partitions was assigned its own UCLN clock partition and used the same gamma-Dirichlet construction, with $\bar{r}_{\text{clock}} \sim \text{Gamma}(2, 2000)$, $\overline{\sigma^2}_{\text{clock}} \sim \text{Gamma}(1, 10)$, and 15-dimensional $\text{Dirichlet}(1, \ldots, 1)$ normalized mean-rate and variance vectors. Each sequence partition had its own GTR exchangeabilities, equilibrium base frequencies, and discrete-gamma shape parameter, assigned $\text{Dirichlet}(1, \ldots, 1)$ priors of dimensions six and four and $\alpha_j \sim \text{Exp}(1)$ for the discrete gamma shape parameter, respectively.

We assigned $x_0 - 54.8 \sim \text{Exp}(0.1)$. Its lower offset is the older bound of *Psilosemys wyomingensis*, which was assigned an age interval of 53.0–54.8 Mya and placed on the emydid stem (Hutchison, 2013; Vlachos, 2018). The prior has mean 64.8 Mya and a central 95% interval 55.1–91.7 Mya, reflecting the considerable disagreement among published estimates of the emydid stem age. These range from

52.49 Mya, with a 95% HPD of 32.83–88.69 Mya (Spinks et al., 2016), through approximately 74–75 Mya (Near et al., 2005; Spinks et al., 2009), to 82.43 Mya (Joyce et al., 2013), 87.6 Mya (Pereira et al., 2017), and approximately 90 Mya (Lourenço et al., 2012). We therefore used the oldest fossil bound as a strict lower offset while retaining a broad upper tail.

#### 2.3 Palaeodictyopterida

For each of the OU-smoothed $\lambda$ and $\mu$ skylines, we assigned $\theta \sim \text{Normal}(\log 0.2, 1.6^2)$, $\sigma_{\text{OU}} \sim \text{Exp}(10/3)$, and $\nu \sim \text{Exp}(2)$. Fossil-recovery rates were independently assigned $\psi_i \sim \text{Exp}(10)$.

We assigned $x_0 - 327 \sim \text{Exp}(0.1)$, giving a mode of 327 Mya, a mean of 337 Mya, and a central 95% interval of 327.3–363.9 Mya. The offset lies immediately beyond the older bounds of the oldest sampled records, *Delitzschala bitterfeldensis*, *Xenoptera latigra*, and *X. riojaensis*, whose age intervals extend to 326.9 Mya (Pinto, 1986; Brauckmann & Schneider, 1996; Pérez Loinaze, 2009; Petrulevičius & Gutiérrez, 2025). Because Palaeodictyopterida is nested within Pterygota, its origin must postdate the origin of winged insects. Older putative evidence for winged insects is uncertain; *Rhyniognatha hirsti* Tillyard, 1928, for example, has been reinterpreted as more likely a myriapod (Haug & Haug, 2017). Fossil-based analyses place the origin of crown Pterygota near the Serpukhovian (Schachat et al., 2023), whereas molecular-clock analyses have placed the origin of insect flight near 406 Mya (Misof et al., 2014; Wang et al., 2016). We therefore concentrated prior mass near the oldest verifiable Palaeodictyopterida fossils while retaining a diffuse tail toward older origins.

### References


Abel O. 1905a. Eine stammtype der Delphiniden aus dem Miocän der halbinsel Taman. *Jahrbuch der Kaiserlich-Königlichen Geologischen Reichsanstalt*. *55*:375–392.

Abel O. 1905b. Les odontocètes du Boldérien (Miocène supérieur) d'Anvers. *Mémoires du Musée royal d'histoire naturelle de Belgique*. *3*:1–155. https://doi.org/10.5962/bhl.title.15923

Abel O. 1914. Die vorfahren der Bartenwale. *Denkschriften der Kaiserlichen Akademie der Wissenschaften, Mathematisch-Naturwissenschaftliche Klasse*. *90*:155–224.

Agnus A. 1902. Description d'un névroptère fossile nouveau *Homoioptera gigantea*. *Bulletin de la Société entomologique de France*. *7*(16):259–261. https://doi.org/10.3406/bsef.1902.23126

Aguirre-Fernández G, Barnes LG, Aranda-Manteca FJ, Fernández-Rivera JR. 2009. *Protoglobicephala mexicana*, a new genus and species of Pliocene fossil dolphin (Cetacea; Odontoceti; Delphinidae) from the Gulf of California, Mexico. *Boletín de la Sociedad Geológica Mexicana*. *61*(2):245–265. https://doi.org/10.18268/bsgm2009v61n2a13

Aguirre-Fernández G, Fordyce RE. 2014. *Papahu taitapu*, gen. et sp. nov., an early Miocene stem odontocete (Cetacea) from New Zealand. *Journal of Vertebrate Paleontology*. *34*(1):195–210. https://doi.org/10.1080/02724634.2013.799069

Aguirre-Fernández G, Jost J, Hilfiker S. 2022. First records of extinct kentriodontid and squalodelphinid dolphins from the Upper Marine Molasse (Burdigalian age) of Switzerland and a reappraisal of the Swiss cetacean fauna. *PeerJ*. *10*:e13251. https://doi.org/10.7717/peerj.13251

Albright LB, Sanders AE, Geisler JH. 2018. An unexpectedly derived odontocete from the Ashley formation (upper Rupelian) of South Carolina, U.S.A. *Journal of Vertebrate Paleontology*. *38*(4):e1482555. https://doi.org/10.1080/02724634.2018.1482555

Alfsen A, Bosselaers M, Lambert O. 2021. New sperm whale remains from the late Miocene of the North Sea and a revised family attribution for the small crown physeteroid *Thalassocetus* Abel, 1905. *Comptes Rendus Palevol*. *20*(39):807–822. https://doi.org/10.5852/cr-palevol2021v20a39

Allen GM. 1921a. A new fossil cetacean. *Bulletin of the Museum of Comparative Zoology*. *65*(1):1–14.

Allen GM. 1921b. Fossil cetaceans from the Florida phosphate beds. *Journal of Mammalogy*. *2*(3):144–159. https://doi.org/10.2307/1373603

Allen GM. 1941. A fossil river dolphin from Florida. *Bulletin of the Museum of Comparative Zoology*. *89*(1):1–8.

Allen JA. 1887. Note on squalodont remains from Charleston, S.C. *Bulletin of the American Museum of Natural History*. *2*(2):35–39.

Ameghino F. 1891. Caracteres diagnósticos de cincuenta especies nuevas de mamíferos fósiles argentinos. *Revista Argentina de Historia Natural*. *1*(3):129–167.

Andrée R. 1864. Die Versteinerungen der Steinkohlenformation von Stradonic in Böhmen. *Neues Jahrbuch für Mineralogie, Geologie und Paläontologie*:160–176.

Andrews CW. 1906. *A Descriptive Catalogue of the Tertiary Vertebrata of the Fayûm, Egypt*. London: British Museum (Natural History). https://doi.org/10.5962/bhl.title.55134

Andrews CW. 1920. A description of new species of zeuglodont and of leathery turtle from the Eocene of southern Nigeria. *Proceedings of the Zoological Society of London*. *1919*:309–319.

Antar MS, Gohar AS, El-Desouky H, Seiffert ER, El-Sayed S, Claxton AG, Sallam HM. 2023. A diminutive new basilosaurid whale reveals the trajectory of the cetacean life histories during the Eocene. *Communications Biology*. *6*:707. https://doi.org/10.1038/s42003-023-04986-w

Archangelsky S, Azcuy CL, Wagner RH. 1981. Three dwarf lycophytes from the Carboniferous of Argentina. *Scripta Geologica*. *64*:1–35.

Aslanova SM. 1977. Noviy rod kitoobraznykh (*Atropatenocetus posteocenicus* gen. et sp. nov.) iz oligotsena azerbaidzhana. *Doklady Akademii Nauk Azerbaidzhanskoi SSR*. *33*(7):60–64.

Aslanova SM, Mchedlidze GA. 1976. Novyy rod semeystva Delphinidae na Kavkaze. *Voprosy Paleontologii i Stratigrafii Azerbaydzhana. 1*:17–39.

Bajpai S, Gingerich PD. 1998. A new Eocene archaeocete (Mammalia, Cetacea) from India and the time of origin of whales. *Proceedings of the National Academy of Sciences of the United States of America. 95*(26):15464–15468. https://doi.org/10.1073/pnas.95.26.15464

Bajpai S, Thewissen JGM. 1998. Middle Eocene cetaceans from the Harudi and Subathu formations of India. In: Thewissen JGM, editor. *The Emergence of Whales: Evolutionary Patterns in the Origin of Cetacea*. New York: Plenum Press. p. 213–233. https://doi.org/10.1007/978-1-4899-0159-0_7

Bajpai S, Thewissen JGM. 2000. A new, diminutive Eocene whale from Kachchh (Gujarat, India) and its implications for locomotor evolution of cetaceans. *Current Science. 79*(10):1478–1482.

Bajpai S, Thewissen JGM. 2014. Protocetid cetaceans (Mammalia) from the Eocene of India. *Palaeontologia Electronica. 17*(3):34A. https://doi.org/10.26879/459

Barnes LG. 1973. *Praekogia cedrosensis*, a new genus and species of fossil pygmy sperm whale from Isla Cedros, Baja California, Mexico. *Contributions in Science, Natural History Museum of Los Angeles County. 247*:1–20.

Barnes LG. 1978. A review of *Lophocetus* and *Liolithax* and their relationships to the delphinoid family Kentriodontidae (Cetacea: Odontoceti). *Bulletin of the Natural History Museum of Los Angeles County. 28*:1–35.

Barnes LG. 1984. Fossil odontocetes (Mammalia: Cetacea) from the Almejas Formation, Isla Cedros, Mexico. *PaleoBios. 42*:1–46.

Barnes LG. 1985a. Fossil pontoporiid dolphins (Mammalia: Cetacea) from the Pacific coast of North America. *Contributions in Science, Natural History Museum of Los Angeles County. 363*:1–34.

Barnes LG. 1985b. The late Miocene dolphin *Pithanodelphis* Abel, 1905 (Cetacea: Kentriodontidae) from California. *Contributions in Science, Natural History Museum of Los Angeles County. 367*:1–27.

Barnes LG. 2002. An early Miocene long-snouted marine platanistid dolphin (Mammalia, Cetacea, Odontoceti) from the Korneuburg Basin (Austria). *Beiträge zur Paläontologie. 27*:407–418.

Barnes LG. 2008. Miocene and Pliocene Albireonidae (Cetacea, Odontoceti), rare and unusual fossil dolphins from the eastern North Pacific Ocean. In: Wang X, Barnes LG, editors. *Geology and Vertebrate Paleontology of Western and Southern North America: Contributions in Honor of David P. Whistler*. Vol. 41. Natural History Museum of Los Angeles County. p. 99–152.

Barnes LG, Kimura M, Furusawa H, Sawamura H. 1995. Classification and distribution of Oligocene Aetiocetidae (Mammalia; Cetacea; Mysticeti) from western North America and Japan. *The Island Arc. 3*(4):392–431. https://doi.org/10.1111/j.1440-1738.1994.tb00122.x

Barnes LG, Reynolds RE. 2009. A New Species of Early Miocene Allodelphinid Dolphin (Cetacea, Odontoceti, Platanistoidea) from Cajon Pass, Southern California, U.S.A. In: Albright LB, editor. *Papers on Geology, Vertebrate Paleontology, and Biostratigraphy in Honor of Michael O. Woodburne*. Vol. 65. p. 483–507.

Bashforth AR, Falcon-Lang HJ, Gibling MR. 2010. Vegetation heterogeneity on a Late Pennsylvanian braided-river plain draining the Variscan Mountains, La Magdalena Coalfield, northwestern Spain. *Palaeogeography, Palaeoclimatology, Palaeoecology*. *292*:367–390. https://doi.org/10.1016/j.palaeo.2010.03.037

Bebej RM, Zalmout IS, Abd El-Aziz AA, Antar MSM, Gingerich PD. 2015. First remingtonocetid archaeocete (Mammalia, Cetacea) from the middle Eocene of Egypt with implications for biogeography and locomotion in early cetacean evolution. *Journal of Paleontology*. *89*(5):882–893. https://doi.org/10.1017/jpa.2015.57

Beckemeyer RJ. 2004. A new species of *Martynovia* Tillyard, 1932 (Insecta: Diaphanopterodea: Martynoviidae) from the Lower Permian Wellington Formation of Noble County, Oklahoma. *Journal of the Kansas Entomological Society*. *77*(2):127–131. https://doi.org/10.2317/0310.07.1

Beckemeyer RJ, Engel MS. 2009. An enigmatic new genus of biarmohymenid from the Early Permian Wellington Formation of Noble County, Oklahoma (Palaeodictyopterida: Diaphanopterodea). *Transactions of the Kansas Academy of Science*. *112*(1/2):103–108. https://doi.org/10.1660/062.112.0213

Beckemeyer RJ, Engel MS. 2011. Upper Carboniferous insects from the Pottsville Formation of northern Alabama. *Scientific Papers, Natural History Museum, University of Kansas*. *44*:1–19.

Beckemeyer RJ, Engel MS. 2018. First North American species of the European genus *Archaemegaptilus* from the Upper Carboniferous Pottsville Formation of northern Alabama (Palaeodictyoptera: Archaemegaptilidae). *Novitates Paleoentomologicae*. (21):1–7. https://doi.org/10.17161/np.v0i21.7164

Behrmann G. 1995. Der Bartenwal aus dem Miozän von Gr. Pampau (Schleswig-Holstein). *Geschiebekunde aktuell*. *11*:119–129.

Belluzzo A, Lambert O. 2021. A new delphinid from the lower Pliocene of the North Sea and the early radiations of true dolphins. *Fossil Record*. *24*(1):77–92. https://doi.org/10.5194/fr-24-77-2021

Ben Chehida Y, Thumloup J, Schumacher C, Harkins T, Aguilar A, Borrell A, Ferreira M, Rojas-Bracho L, Robertson KM, Taylor BL, et al. 2020. Mitochondrial genomics reveals the evolutionary history of the porpoises (Phocoenidae) across the speciation continuum. *Scientific Reports*. *10*(1):15190. https://doi.org/10.1038/s41598-020-71603-9

Benham W. 1935. The teeth of an extinct whale *Microcetus hectori*, n. sp. *Transactions of the Royal Society of New Zealand*. *65*:239–243.

Benham WB. 1937. Fossil Cetacea of New Zealand. II. *Lophocephalus*, a new genus of zeuglodont Cetacea. *Transactions of the Royal Society of New Zealand*. *67*:1–7.

Benites-Palomino A, Aguirre-Fernández G, Baby P, Ochoa D, Altamirano A, Flynn JJ, Sanchez-Villagra MR. 2024. The largest freshwater odontocete: A South Asian river dolphin relative from the proto-Amazonia. *Science Advances*. *10*(12):eadk6320. https://doi.org/10.1126/sciadv.adk6320

Benites-Palomino A, Vélez-Juarbe J, Salas-Gismondi R, Urbina M. 2020. *Scaphokogia totajpe*, sp. nov., a new bulky-faced pygmy sperm whale (Kogiidae) from the late Miocene of Peru. *Journal of Vertebrate Paleontology*. *39*(6):e1728538. https://doi.org/10.1080/02724634.2019.1728538

Benites-Palomino AM, Vélez-Juarbe J, Collareta A, Ochoa D, Altamirano A, Carré M, Laime MJ, Urbina M, Salas-Gismondi R. 2021. Nasal compartmentalization in Kogiidae (Cetacea, Physeteroidea): Insights from a new late Miocene dwarf sperm whale from the Pisco formation. *Papers in Palaeontology*. *7*(3):1507–1524. https://doi.org/10.1002/spp2.1351

Benoit J, Adnet S, Welcomme JL, Fabre PH. 2011. New skull of *Schizodelphis sulcatus* Gervais, 1861 (Mammalia, Odontoceti, Eurhinodelphinidae) from the lower Miocene of Pignan (Hérault, France) and its implications for systematics of Eurhinodelphinidae. *Geobios*. *44*(4):323–334. https://doi.org/10.1016/j.geobios.2010.10.006

Bernor RL, Cirilli O, Jukar AM, Potts R, Buskianidze M, Rook L. 2019. Evolution of early *Equus* in Italy, Georgia, the Indian subcontinent, East Africa, and the origins of African zebras. *Frontiers in Ecology and Evolution*. *7*:166. https://doi.org/10.3389/fevo.2019.00166

Béthoux O, Nel A, Lapeyrie J, Gand G, Galtier J. 2003. New Martynoviidae from the Permian of southern France (Lodève basin) (Insecta: Palaeoptera: Diaphanopterodea). *Geobios*. *36*(2):131–139. https://doi.org/10.1016/s0016-6995(03)00002-0

Béthoux O, Nel A, Schneider JW, Gand G. 2007. *Lodetiella magnifica* nov. gen. and nov. sp. (Insecta: Palaeodictyoptera; Permian), an extreme situation in wing morphology of palaeopterous insects. *Geobios*. *40*(2):181–189. https://doi.org/10.1016/j.geobios.2006.04.001

Bianucci G. 1996. The Odontoceti (Mammalia, Cetacea) from Italian Pliocene. Systematics and phylogenesis of Delphinidae. *Palaeontographia Italica*. *83*:73–167.

Bianucci G. 1997. The Odontoceti (Mammalia, Cetacea) from Italian Pliocene. The Ziphiidae. *Palaeontographia Italica*. *84*:163–192.

Bianucci G. 2001. A new genus of kentriodontid (Cetacea: Odontoceti) from the Miocene of south Italy. *Journal of Vertebrate Paleontology*. *21*(3):573–577. https://doi.org/10.1671/0272-4634(2001)021[0573:ANGOKC]2.0.CO;2

Bianucci G. 2005. *Arimidelphis sorbinii* a new small killer whale-like dolphin from the Pliocene of Marecchia River (Central Eastern Italy) and a phylogenetic analysis of the Orcininae (Cetacea: Odontoceti). *Rivista Italiana di Paleontologia e Stratigrafia*. *111*(2):329–344. https://doi.org/10.13130/2039-4942/6324

Bianucci G. 2013. *Septidelphis morii*, n. gen. et sp., from the Pliocene of Italy: New evidence of the explosive radiation of true dolphins (Odontoceti, Delphinidae). *Journal of Vertebrate Paleontology*. *33*(3):722–740. https://doi.org/10.1080/02724634.2013.744757

Bianucci G, Bosio G, Malinverno E, de Muizon C, Villa IM, Urbina M, Lambert O. 2018. A new large squalodelphinid (Cetacea, Odontoceti) from Peru sheds light on the early Miocene platanistoid disparity and ecology. *Royal Society Open Science*. *5*(4):172302. https://doi.org/10.1098/rsos.172302

Bianucci G, Collareta A, Bosio G, Di Celma C, Malinverno E, Villa IM, Urbina M, Lambert O. 2024. A new late Miocene beaked whale (Cetacea, Odontoceti) from the Pisco formation, and a revised age for the fossil Ziphiidae of Peru. *Bollettino della Società Paleontologica Italiana*. *63*(1):21–43. https://doi.org/10.4435/BSPI.2024.10

Bianucci G, de Muizon C, Urbina M, Lambert O. 2020. Extensive diversity and disparity of the early Miocene platanistoids (Cetacea, Odontoceti) in the southeastern Pacific (Chilcatay formation, Peru). *Life*. *10*(3):27. https://doi.org/10.3390/life10030027

Bianucci G, Di Celma C, Urbina M, Lambert O. 2016. New beaked whales from the late Miocene of Peru and evidence for convergent evolution in stem and crown Ziphiidae (Cetacea, Odontoceti). *PeerJ*. *4*:e2479. https://doi.org/10.7717/peerj.2479

Bianucci G, Geisler JH, Citron S, Collareta A. 2022. The origins of the killer whale ecomorph. *Current Biology*. *32*(8):1843–1851.e2. https://doi.org/10.1016/j.cub.2022.02.041

Bianucci G, Gingerich PD. 2011. *Aegyptocetus tarfa*, n. gen. et sp. (Mammalia, Cetacea), from the middle Eocene of Egypt: Clinorhynchy, olfaction, and hearing in a protocetid whale. *Journal of Vertebrate Paleontology*. *31*(6):1173–1188. https://doi.org/10.1080/02724634.2011.607985

Bianucci G, Lambert O, Post K. 2007. A high diversity in fossil beaked whales (Mammalia, Odontoceti, Ziphiidae) recovered by trawling from the sea floor off South Africa. *Geodiversitas*. *29*(4):561–618.

Bianucci G, Lambert O, Post K. 2010. High concentration of long-snouted beaked whales (genus *Messapicetus*) from the Miocene of Peru. *Palaeontology*. *53*(5):1077–1098. https://doi.org/10.1111/j.1475-4983.2010.00995.x

Bianucci G, Lambert O, Urbina M, Merella M, Collareta A, Bennion R, Salas-Gismondi R, Benites-Palomino A, Post K, de Muizon C, et al. 2023. A heavyweight early whale pushes the boundaries of vertebrate morphology. *Nature*. *620*:824–829. https://doi.org/10.1038/s41586-023-06381-1

Bianucci G, Landini W. 2002. A new short-rostrum odontocete (Mammalia: Cetacea) from the middle Miocene of the eastern Netherlands. *Beaufortia*. *52*(11):187–196.

Bianucci G, Landini W. 2006. Killer sperm whale: A new basal physeteroid (Mammalia, Cetacea) from the late Miocene of Italy. *Zoological Journal of the Linnean Society*. *148*(1):103–131. https://doi.org/10.1111/j.1096-3642.2006.00228.x

Bianucci G, Landini W, Varola A. 1992. *Messapicetus longirostris*, a new genus and species of Ziphiidae (Cetacea) from the late Miocene of "Pietra leccese" (Apulia, Italy). *Bollettino della Società Paleontologica Italiana*. *31*(2):261–264.

Bianucci G, Mazza P, Merola D, Sarti G, Cascella A. 2001. The early Pliocene mammal assemblage of Val di Pugna (Tuscany, Italy) in the light of calcareous plankton biostratigraphical data and paleoecological observations. *Rivista Italiana di Paleontologia e Stratigrafia*. *107*(3). https://doi.org/10.13130/2039-4942/5443

Bianucci G, Pesci F, Collareta A, Tinelli C. 2019. A new Monodontidae (Cetacea, Delphinoidea) from the lower Pliocene of Italy supports a warm-water origin for narwhals and white whales. *Journal of Vertebrate Paleontology*. *39*(3):e1645148. https://doi.org/10.1080/02724634.2019.1645148

Bianucci G, Post K. 2005. *Caviziphius altirostris*, a new beaked whale from the Miocene southern North Sea basin. *Deinsea*. *11*:1–6.

Bianucci G, Urbina M, Lambert O. 2015. A new record of *Notocetus vanbenedeni* (Squalodelphinidae, Odontoceti, Cetacea) from the early Miocene of Peru. *Comptes Rendus Palevol*. *14*(1):5–13. https://doi.org/10.1016/j.crpv.2014.08.003

Bisconti M. 2005. Skull morphology and phylogenetic relationships of a new diminutive balaenid from the Lower Pliocene of Belgium. *Palaeontology*. *48*(4):793–816. https://doi.org/10.1111/j.1475-4983.2005.00488.x

Bisconti M. 2006. *Titanocetus*, a new baleen whale from the Middle Miocene of northern Italy (Mammalia, Cetacea, Mysticeti). *Journal of Vertebrate Paleontology*. *26*(2):344–354. https://doi.org/10.1671/0272-4634(2006)26[344:TANBWF]2.0.CO;2

Bisconti M. 2007a. A new basal balaenopterid whale from the Pliocene of northern Italy. *Palaeontology*. *50*(5):1103–1122. https://doi.org/10.1111/j.1475-4983.2007.00696.x

Bisconti M. 2007b. Taxonomic revision and phylogenetic relationships of the rorqual-like mysticete from the Pliocene of Mount Pulgnasco, northern Italy (Cetacea, Mysticeti). *Palaeontographia Italica*. *91*:85–108.

Bisconti M. 2008. Morphology and phylogenetic relationships of a new eschrichtiid genus (Cetacea: Mysticeti) from the Early Pliocene of northern Italy. *Zoological Journal of the Linnean Society*. *153*(1):161–186. https://doi.org/10.1111/j.1096-3642.2008.00374.x

Bisconti M. 2010. A new balaenopterid whale from the late Miocene of the Stirone river, northern Italy (Mammalia, Cetacea, Mysticeti). *Journal of Vertebrate Paleontology*. *30*(3):943–958. https://doi.org/10.1080/02724631003762922

Bisconti M. 2012. Comparative osteology and phylogenetic relationships of *Miocaperea pulchra*, the first fossil pygmy right whale genus and species (Cetacea, Mysticeti, Neobalaenidae). *Zoological Journal of the Linnean Society*. *166*(4):876–911. https://doi.org/10.1111/j.1096-3642.2012.00862.x

Bisconti M. 2014. Anatomy of a new cetotheriid genus and species from the Miocene of Herentals, Belgium, and the phylogenetic and palaeobiogeographical relationships of Cetotheriidae. *Journal of Systematic Palaeontology*. *13*(5):377–395. https://doi.org/10.1080/14772019.2014.890136

Bisconti M, Bosselaers M. 2016. *Fragilicetus velponi*: A new mysticete genus and species and its implications for the origin of Balaenopteridae (Mammalia, Cetacea, Mysticeti). *Zoological Journal of the Linnean Society*. *177*(2):450–474. https://doi.org/10.1111/zoj.12370

Bisconti M, Bosselaers M. 2020. A new balaenopterid species from the Southern North Sea Basin informs about phylogeny and taxonomy of *Burtinopsis* and *Protororqualus* (Cetacea, Mysticeti, Balaenopteridae). *PeerJ*. *8*:e9570. https://doi.org/10.7717/peerj.9570

Bisconti M, Collareta A, Coletti G, Bosio G, Di Celma C, Bianucci G. 2025a. A new fossil from the late Miocene of Piedmont (north-western Italy) sheds light on the evolution of the Cetotheriidae (Mammalia: Cetacea: Mysticeti). *Journal of Systematic Palaeontology*. *23*(1):2540310. https://doi.org/10.1080/14772019.2025.2540310

Bisconti M, Damarco P, Mao S, Pavia M, Carnevale G. 2021a. The earliest baleen whale from the Mediterranean: Large-scale implications of an early Miocene thalassotherian mysticete from Piedmont, Italy. *Papers in Palaeontology*. *7*(2):1147–1166. https://doi.org/10.1002/spp2.1336

Bisconti M, Damarco P, Marengo L, Macagno M, Daniello R, Pavia M, Carnevale G. 2024. Anatomy and relationships of a new gray whale from the Pliocene of Piedmont, northwestern Italy. *Diversity*. *16*(9):547. https://doi.org/10.3390/d16090547

Bisconti M, Damarco P, Pavia M, Sorce B, Carnevale G. 2021b. *Marzanoptera tersillae*, a new balaenopterid genus and species from the Pliocene of Piedmont, north-west Italy. *Zoological Journal of the Linnean Society*. *192*(4):1253–1292. https://doi.org/10.1093/zoolinnean/zlaa131

Bisconti M, Daniello R, Stecca R, Carnevale G. 2025b. A new Pliocene sperm whale from Vigliano d'Asti, Piedmont, northwest Italy. *Rivista Italiana di Paleontologia e Stratigrafia*. *131*(1):139–175. https://doi.org/10.54103/2039-4942/22338

Bisconti M, Lambert O, Bosselaers M. 2013. Taxonomic revision of *Isocetus depauwi* (Mammalia, Cetacea, Mysticeti) and the phylogenetic relationships of archaic 'cetothere' mysticetes. *Palaeontology*. *56*(1):95–127. https://doi.org/10.1111/j.1475-4983.2012.01168.x

Bisconti M, Lambert O, Bosselaers M. 2017. Revision of "*Balaena*" *belgica* reveals a new right whale species, the possible ancestry of the northern right whale, *Eubalaena glacialis*, and the ages of divergence for the living right whale species. *PeerJ*. *5*:e3464. https://doi.org/10.7717/peerj.3464

Bisconti M, Munsterman DK, Fraaije RHB, Bosselaers M, Post K. 2020. A new species of rorqual whale (Cetacea, Mysticeti, Balaenopteridae) from the Late Miocene of the Southern North Sea Basin and the role of the North Atlantic in the paleobiogeography of *Archaebalaenoptera*. *PeerJ*. *8*:e8315. https://doi.org/10.7717/peerj.8315

Bisconti M, Munsterman DK, Post K. 2019. A new balaenopterid whale from the late Miocene of the Southern North Sea Basin and the evolution of balaenopterid diversity (Cetacea, Mysticeti). *PeerJ*. *7*:e6915. https://doi.org/10.7717/peerj.6915

Bisconti M, Ochoa D, Urbina M, Salas-Gismondi R. 2021c. *Archaebalaenoptera eusebioi*, a new rorqual from the late Miocene of Peru (Cetacea, Mysticeti, Balaenopteridae) and its impact in reconstructing body size evolution, ecomorphology and palaeobiogeography of Balaenopteridae. *Journal of Systematic Palaeontology*. *19*(16):1129–1160. https://doi.org/10.1080/14772019.2021.2017363

Bisconti M, Varola A. 2006. The oldest eschrichtiid mysticete and a new morphological diagnosis of Eschrichtiidae (gray whales). *Rivista Italiana di Paleontologia e Stratigrafia*. *112*(1):1–11.

Boersma AT, McCurry MR, Pyenson ND. 2017. A new fossil dolphin *Dilophodelphis fordycei* provides insight into the evolution of supraorbital crests in Platanistoidea (Mammalia, Cetacea). *Royal Society Open Science*. *4*(5):170022. https://doi.org/10.1098/rsos.170022

Boersma AT, Pyenson ND. 2015. *Albicetus oxymycterus*, a new generic name and redescription of a basal physeteroid (Mammalia, Cetacea) from the Miocene of California, and the evolution of body size in sperm whales. *PLoS One*. *10*(12):e0135551. https://doi.org/10.1371/journal.pone.0135551

Boersma AT, Pyenson ND. 2016. *Arktocara yakataga*, a new fossil odontocete (Mammalia, Cetacea) from the Oligocene of Alaska and the antiquity of Platanistoidea. *PeerJ*. *4*:e2321. https://doi.org/10.7717/peerj.2321

Boessenecker RW. 2013. A new marine vertebrate assemblage from the late Neogene Purisima Formation in central California, part II: Pinnipeds and cetaceans. *Geodiversitas*. *35*(4):815–940. https://doi.org/10.5252/g2013n4a5

Boessenecker RW. 2019. Problematic archaic whale *Phococetus* (Cetacea: Odontoceti) from the Lee Creek mine, North Carolina, USA, with comments on geochronology of the Pungo River formation. *PalZ*. *93*(1):93–103. https://doi.org/10.1007/s12542-018-0419-3

Boessenecker RW, Ahmed E, Geisler JH. 2017a. New records of the dolphin *Albertocetus meffordorum* (Odontoceti: Xenorophidae) from the lower Oligocene of South Carolina: Encephalization, sensory anatomy, postcranial morphology, and ontogeny of early odontocetes. *PLoS One*. *12*(11):e0186476. https://doi.org/10.1371/journal.pone.0186476

Boessenecker RW, Beatty BL, Geisler JH. 2023. New specimens and species of the Oligocene toothed baleen whale *Coronodon* from South Carolina and the origin of Neoceti. *PeerJ*. *11*:e14795. https://doi.org/10.7717/peerj.14795

Boessenecker RW, Boessenecker SJ. 2019. Paleontology of the "Ashley Phosphate Beds" of Charleston: Insights from Northbridge Park, Charleston, South Carolina. In: *Field Excursions in the Carolinas: Guides for the 2019 GSA Southeastern Section Meeting*. Vol. 53. Geological Society of America. p. 1–8.

Boessenecker RW, Churchill M, Buchholtz EA, Beatty BL, Geisler JH. 2020. Convergent evolution of swimming adaptations in modern whales revealed by a large macrophagous dolphin from the Oligocene of South Carolina. *Current Biology*. *30*(16):3267–3273.e2. https://doi.org/10.1016/j.cub.2020.06.012

Boessenecker RW, Ehret DJ, Long DJ, Churchill M, Martin E, Boessenecker SJ. 2019. The early Pliocene extinction of the mega-toothed shark *Otodus megalodon*: A view from the eastern North Pacific. *PeerJ*. *7*:e6088. https://doi.org/10.7717/peerj.6088

Boessenecker RW, Fordyce RE. 2015a. A new eomysticetid (Mammalia: Cetacea) from the late Oligocene of New Zealand and a re-evaluation of *'Mauicetus' waitakiensis*. *Papers in Palaeontology*. *1*(2):107–140. https://doi.org/10.1002/spp2.1005

Boessenecker RW, Fordyce RE. 2015b. A new genus and species of eomysticetid (Cetacea: Mysticeti) and a reinterpretation of *'Mauicetus' lophocephalus* Marples, 1956: Transitional baleen whales from the upper Oligocene of New Zealand. *Zoological Journal of the Linnean Society*. *175*(3):607–660. https://doi.org/10.1111/zoj.12297

Boessenecker RW, Fordyce RE. 2015c. Anatomy, feeding ecology, and ontogeny of a transitional baleen whale: A new genus and species of Eomysticetidae (Mammalia: Cetacea) from the Oligocene of New Zealand. *PeerJ*. *3*:e1129. https://doi.org/10.7717/peerj.1129

Boessenecker RW, Fordyce RE. 2017. A new eomysticetid from the Oligocene Kokoamu Greensand of New Zealand and a review of the Eomysticetidae (Mammalia, Cetacea). *Journal of Systematic Palaeontology*. *15*(6):429–469. https://doi.org/10.1080/14772019.2016.1191045

Boessenecker RW, Fraser D, Churchill M, Geisler JH. 2017b. A toothless dwarf dolphin (Odontoceti: Xenorophidae) points to explosive feeding diversification of modern whales (Neoceti). *Proceedings of the Royal Society B: Biological Sciences*. *284*(1861):20170531. https://doi.org/10.1098/rspb.2017.0531

Boessenecker RW, Geisler JH. 2018. New records of the archaic dolphin *Agorophius* (Mammalia: Cetacea) from the upper Oligocene Chandler Bridge formation of South Carolina, USA. *PeerJ*. *6*:e5290. https://doi.org/10.7717/peerj.5290

Boessenecker RW, Geisler JH. 2023. New skeletons of the ancient dolphin *Xenorophus sloanii* and *Xenorophus simplicidens* sp. nov. (Mammalia, Cetacea) from the Oligocene of South Carolina and the ontogeny, functional anatomy, asymmetry, pathology, and evolution of the earliest Odontoceti. *Diversity*. *15*(11):1154. https://doi.org/10.3390/d15111154

Boessenecker RW, Richards MD. 2024. A review of New Zealand Eomysticetidae (Mammalia, Cetacea) and implications for the evolution of baleen whales: New specimens, functional anatomy, and phylogeny. *Journal of the Royal Society of New Zealand*. *54*(5):696–710. https://doi.org/10.1080/03036758.2023.2277739

Bogachev VV. 1956. Sarmatskii del'fin *Anacharsis orbus* iz okrestnostei g. Stavropolya Kavkazskogo. *Materialy po izucheniyu Stavropol'skogo Kraya*. *8*:237–254.

Bolton H. 1911. On a collection of insect-remains from the South Wales Coalfield. *Quarterly Journal of the Geological Society of London*. *67*(1-4):149–174. https://doi.org/10.1144/gsl.jgs.1911.067.01-04.06

Bolton H. 1912. Insect-remains from the Midland and South-Eastern Coal Measures. *Quarterly Journal of the Geological Society of London*. *68*(1-4):310–323. https://doi.org/10.1144/gsl.jgs.1912.068.01-04.25

Bolton H. 1916. On some insects from the British Coal Measures. *Quarterly Journal of the Geological Society of London*. *72*(1-4):43–62. https://doi.org/10.1144/gsl.jgs.1916.072.01-04.06

Bolton H. 1921. A monograph of the fossil insects of the British Coal Measures. Part I. *Monographs of the Palaeontographical Society*. *73*(348):1–80. https://doi.org/10.1080/02693445.1921.12035583

Bolton H. 1925. *Insects from the Coal Measures of Commentry*. Fossil Insects 2. London: British Museum (Natural History).

Bolton H. 1930. Fossil insects of the South Wales Coalfield. *Quarterly Journal of the Geological Society of London*. *86*(1-4):9–49. https://doi.org/10.1144/gsl.jgs.1930.086.01-04.03

Bolton H. 1934. New forms from the insect fauna of the British Coal Measures. *Quarterly Journal of the Geological Society of London*. *90*(1-4):277–301. https://doi.org/10.1144/gsl.jgs.1934.090.01-04.10

Bosselaers M, Munsterman DK. 2022. The discovery of a Balaenomorpha (*Persufflatius renefraaijeni* n. gen., n. sp.) from the upper Miocene of the Netherlands sheds new light on the cranial anatomy of archaic rorqual relatives. *Geodiversitas*. *44*(30):933–973. https://doi.org/10.5252/geodiversitas2022v44a30

Bosselaers M, Post K. 2010. A new fossil rorqual (Mammalia, Cetacea, Balaenopteridae) from the Early Pliocene of the North Sea, with a review of the rorqual species described by Owen and Van Beneden. *Geodiversitas*. *32*(2):331–363. https://doi.org/10.5252/g2010n2a6

Bouetel V, de Muizon C. 2006. The anatomy and relationships of *Piscobalaena nana* (Cetacea, Mysticeti), a Cetotheriidae s.s. from the early Pliocene of Peru. *Geodiversitas*. *28*(2):319–395.

Brandt JF. 1843. De cetotherio, novo balaenarum familiae genere in rossia meridionali ante aliquot annos effosso. *Bulletin de la Classe Physico-Mathématique de l'Académie Impériale des Sciences de Saint-Pétersbourg*. *1*(10–12):145–148.

Brandt JF. 1871. Bericht über den fortgang meiner studien über die cetaceen. *Bulletin de l'Académie Impériale des Sciences de Saint-Pétersbourg*. *16*:563–566.

Brandt JF. 1873a. Über bisher in russland gefundene reste von zeuglodonten. *Mélanges biologiques tirés du Bulletin de l'Académie Impériale des Sciences de Saint-Pétersbourg*. *9*:111–112.

Brandt JF. 1873b. Untersuchungen über die fossilen und subfossilen Cetaceen Europa's. *Mémoires de l'Académie Impériale des Sciences de Saint-Pétersbourg, 7e série*. *20*(1):1–372. https://doi.org/10.5962/bhl.title.39524

Brandt JF. 1874. Ergänzungen zu den fossilen Cetaceen Europa's. *Mémoires de l'Académie Impériale des Sciences de Saint-Pétersbourg, série 7*. *21*:1–54.

Brauckmann C. 1984. Weitere neue Insekten (Palaeodictyoptera; Protorthoptera) aus dem Namurium B von Hagen-Vorhalle. *Jahresberichte des Naturwissenschaftlichen Vereins in Wuppertal*. *37*:108–115.

Brauckmann C. 1986. Eine neue Spilapteriden-Art aus dem Namurium B von Hagen-Vorhalle (Insecta: Palaeodictyoptera; Ober-Karbon, West-Deutschland). *Dortmunder Beiträge zur Landeskunde, Naturwissenschaftliche Mitteilungen*. *20*:57–64.

Brauckmann C. 1988. Zwei neue Insekten (Odonata, Megasecoptera) aus dem Namurium von Hagen-Vorhalle (West-Deutschland). *Dortmunder Beiträge zur Landeskunde, Naturwissenschaftliche Mitteilungen*. *22*:91–101.

Brauckmann C. 1991. Ein neuer Insekten-Rest (Megasecoptera) aus dem Ober-Karbon von Osnabrück. *Osnabrücker Naturwissenschaftliche Mitteilungen*. *17*:25–32.

Brauckmann C. 1995. Neue Insekten-Funde (Palaeodictyoptera: Breyeriidae) aus dem Ober-Karbon von Osnabrück (Deutschland). *Osnabrücker Naturwissenschaftliche Mitteilungen*. *20/21*:157–165.

Brauckmann C, Becker R. 1992. Ein neues Riesen-Insekt (*Scepasma mediomatricorum* n. sp.) aus dem Ober-Karbon (Westfalium C) des Saarlandes (Palaeodictyoptera: Homoiopteridae). *Geologica et Palaeontologica*. *26*:135–141.

Brauckmann C, Chesnut DR, Jennings JR. 1993. New spilapterid insect from the Breathitt Formation (Middle Pennsylvanian, Westphalian B) of eastern Kentucky, USA. *Neues Jahrbuch für Geologie und Paläontologie, Monatshefte. 1993*(11):641–647. https://doi.org/10.1127/njgpm/1993/1993/641

Brauckmann C, Gröning E. 1996. Ein neuer ober-karbonischer Insekten-Flügel aus Süd-Limburg (Palaeodictyoptera: Breyeriidae, Westfalium, Niederlande). *Neues Jahrbuch für Geologie und Paläontologie, Monatshefte. 1996*(2):72–82. https://doi.org/10.1127/njgpm/1996/1996/72

Brauckmann C, Gröning E. 1998. A new species of *Homaloneura* from the Namurian (Upper Carboniferous) of Hagen-Vorhalle (Germany). *Entomologia Generalis. 23*(1/2):77–84. https://doi.org/10.1127/entom.gen/23/1998/77

Brauckmann C, Hahn G. 1978. Ein neuer Palaeodictyopteren-Fund aus dem westdeutschen Namurium. *Paläontologische Zeitschrift. 52*(1/2):13–18. https://doi.org/10.1007/BF03006726

Brauckmann C, Hahn G. 1983. Ein Palaeodictyopteren-flügel aus dem Ober-Karbon der Karnischen Alpen. *Carinthia II. 173*(93):245–251.

Brauckmann C, Herd KJ. 2000. Eine weitere neue Breyeriiden-Art (Insecta: Palaeodictyoptera) aus dem Ober-Karbon von Osnabrück (Deutschland). *Neues Jahrbuch für Geologie und Paläontologie, Monatshefte. 2000*(6):333–344. https://doi.org/10.1127/njgpm/2000/2000/333

Brauckmann C, Herd KJ. 2002. Insekten-Funde aus dem Westfalium D (Ober-Karbon) des Piesberges bei Osnabrück (Deutschland). Teil 1: Palaeoptera. *Osnabrücker Naturwissenschaftliche Mitteilungen. 28*:27–69.

Brauckmann C, Herd KJ, Leipner A. 2009. Insekten-Funde aus dem Westfalium D (Ober-Karbon) des Piesberges bei Osnabrück (Deutschland). Nachtrag 1: Palaeodictyopteroidea. *Osnabrücker Naturwissenschaftliche Mitteilungen. 35*:5–29.

Brauckmann C, Herd KJ, Leipner A. 2023. First Eugereonidae (Insecta: Palaeodictyoptera) from the Pennsylvanian (Late Carboniferous) of the Piesberg site near Osnabrück, Germany. *PalZ. 97*(4):861–866. https://doi.org/10.1007/s12542-021-00578-3

Brauckmann C, Koch L. 1982. Neue Insekten aus den Vorhalle-Schichten (oberes Namurium B) von Hagen-Vorhalle. *Dortmunder Beiträge zur Landeskunde, Naturwissenschaftliche Mitteilungen. 16*:15–26.

Brauckmann C, Koch L. 1983. Eine weitere neue Insektenart aus den Vorhalle-Schichten (Ober-Karbon, oberes Namurium B) von Hagen-Vorhalle. *Dortmunder Beiträge zur Landeskunde, Naturwissenschaftliche Mitteilungen. 17*:3–8.

Brauckmann C, Koch L, Kemper M. 1985. Spinnentiere (Arachnida) und Insekten aus den Vorhalle-Schichten (Namurium B; Ober-Karbon) von Hagen-Vorhalle (West-Deutschland). *Geologie und Paläontologie in Westfalen. 3*:1–131.

Brauckmann C, Schneider J. 1996. Ein unter-karbonisches Insekt aus dem Raum Bitterfeld/Delitzsch (Pterygota, Arnsbergium, Deutschland). *Neues Jahrbuch für Geologie und Paläontologie, Monatshefte. 1996*(1):17–30. https://doi.org/10.1127/njgpm/1996/1996/17

Brauckmann C, Schöllmann L, Sippel W. 2003. Die fossilen Insekten, Spinnentiere und Eurypteriden von Hagen-Vorhalle. *Geologie und Paläontologie in Westfalen. 59*:5–89.

Bravard A. 1858. Monografía de los terrenos marinos terciarios de las cercanías del Paraná. *Anales del Museo Nacional de Buenos Aires. 3*:45–94.

Brongniart C. 1882. Sur un nouvel insecte fossile de l'ordre des Orthoptères, provenant des terrains houillers de Commentry (Allier). *Comptes rendus hebdomadaires des séances de l'Académie des sciences. 95*:1228–1230.

Brongniart C. 1885. Les insectes fossiles des terrains primaires: Coup d'œil rapide sur la faune entomologique des terrains paléozoïques. *Bulletin de la Société des amis des sciences naturelles de Rouen*:50–68. https://doi.org/10.5962/bhl.title.36372

Brongniart C. 1890. Note sur quelques insectes fossiles du terrain houiller qui présentent au prothorax des appendices aliformes. *Bulletin de la Société philomathique de Paris, 8e série. 2*(3):154–158.

Brongniart C. 1894. *Recherches pour servir à l'histoire des insectes fossiles des temps primaires, précédées d'une étude sur la nervation des ailes des insectes.* Saint-Étienne: Théolier. https://doi.org/10.5962/bhl.title.65984

Budnikov IV, Kutygin RV, Shi GR, Sivtchikov VE, Krivenko OV. 2020. Permian stratigraphy and paleogeography of Central Siberia (Angaraland) – A review. *Journal of Asian Earth Sciences. 196*:104365. https://doi.org/10.1016/j.jseaes.2020.104365

Buono MR, Cozzuol MA. 2013. A new beaked whale (Cetacea, Odontoceti) from the Late Miocene of Patagonia, Argentina. *Journal of Vertebrate Paleontology. 33*(4):986–997. https://doi.org/10.1080/02724634.2013.752377

Buono MR, Fernández MS, Cozzuol MA, Cuitiño JI, Fitzgerald EMG. 2017. The early Miocene balaenid *Morenocetus parvus* from Patagonia (Argentina) and the evolution of right whales. *PeerJ. 5*:e4148. https://doi.org/10.7717/peerj.4148

Burmeister H. 1871. On *Saurocetes argentinus*, a new type of Zeuglodontidae. *Annals and Magazine of Natural History (series 4). 7*:51–55.

Cabrera A. 1926. Cetáceos fósiles del Museo de La Plata. *Revista del Museo de La Plata. 29*:363–411.

Capellini G. 1876. Sulle balene fossili toscane. *Atti della Reale Accademia dei Lincei, serie 2. 3*:9–14.

Capellini G. 1878. Della pietra leccese e di alcuni suoi fossili. *Memorie della Reale Accademia delle Scienze dell'Istituto di Bologna. 3*(4):1–34.

Capellini G. 1883. Di un'orca fossile scoperta a Cetona in Toscana. *Memorie della Reale Accademia delle Scienze dell'Istituto di Bologna, serie 4. 4*:1–25.

Capellini G. 1901. Balenottera miocenica del Monte Titano, Repubblica di S. Marino. *Memorie della Reale Accademia delle Scienze dell'Istituto di Bologna. 9*:1–26.

Capellini G. 1903. Avanzi di squalodonte nella arenaria di Grumi dei Frati. *Memorie della Reale Accademia delle Scienze dell'Istituto di Bologna. 10*:437–445.

Capellini G. 1904. *Balaena montalionis. Memorie della Reale Accademia delle Scienze dell'Istituto di Bologna, serie 6. 1*:102–170.

Carnevale G, Godfrey SJ. 2018. Miocene bony fishes from the Calvert, Choptank, St. Marys and Eastover formations, Chesapeake group, Maryland and Virginia. In: Godfrey SJ, editor. *The Geology and Vertebrate Paleontology of Calvert Cliffs, Maryland, USA*. Washington, D.C.: Smithsonian Institution Scholarly Press. p. 161–198. https://doi.org/10.5479/si.1943-6688.100

Carpenter FM. 1930. The Lower Permian insects of Kansas. Part 3. the Protohymenoptera. *Psyche: A Journal of Entomology. 37*(4):343–374. https://doi.org/10.1155/1930/79267

Carpenter FM. 1933. The Lower Permian insects of Kansas. Part 6. Delopteridae, Protelytroptera, Plectoptera and a new collection of Protodonata, Odonata, Megasecoptera, Homoptera, and Psocoptera. *Proceedings of the American Academy of Arts and Sciences. 68*(11):411–503. https://doi.org/10.2307/20022959

Carpenter FM. 1934. A new megasecopteron from the Carboniferous of Kansas. *The University of Kansas Science Bulletin. 21*:365–367.

Carpenter FM. 1939. The Lower Permian insects of Kansas. Part 8: Additional Megasecoptera, Protodonata, Odonata, Homoptera, Psocoptera, Protelytroptera, Plectoptera, and Protoperlaria. *Proceedings of the American Academy of Arts and Sciences. 73*(3):29–70. https://doi.org/10.2307/25130151

Carpenter FM. 1940. Carboniferous insects from the Stanton Formation, Kansas. *American Journal of Science. 238*(9):636–642. https://doi.org/10.2475/ajs.238.9.636

Carpenter FM. 1943. Carboniferous insects from the vicinity of Mazon Creek, Illinois. *Illinois State Museum Scientific Papers. 3*(1):9–24.

Carpenter FM. 1947. Lower Permian insects from Oklahoma. Part 1. introduction and the orders Megasecoptera, Protodonata, and Odonata. *Proceedings of the American Academy of Arts and Sciences. 76*(2):25–54. https://doi.org/10.2307/20023497

Carpenter FM. 1962. A Permian Megasecopteron from Texas. *Psyche: A Journal of Entomology. 69*(1):37–41. https://doi.org/10.1155/1962/90353

Carpenter FM. 1963. A Megasecopteron from Upper Carboniferous strata in Spain. *Psyche: A Journal of Entomology. 70*(1):44–49. https://doi.org/10.1155/1963/10721

Carpenter FM. 1964. Studies on North American Carboniferous insects. 3. a spilapterid from the vicinity of Mazon Creek, Illinois (Palaeodictyoptera). *Psyche: A Journal of Entomology. 71*(3):117–124. https://doi.org/10.1155/1964/21384

Carpenter FM. 1967. Studies on North American Carboniferous insects. 5. Palaeodictyoptera and Megasecoptera from Illinois and Tennessee, with a discussion of the order Sypharopteroidea. *Psyche: A Journal of Entomology. 74*:58–84. https://doi.org/10.1155/1967/58796

Carpenter FM. 1970. Fossil insects from New Mexico. *Psyche: A Journal of Entomology. 77*:400–412. https://doi.org/10.1155/1970/84847

Carpenter FM. 1976. The Lower Permian insects of Kansas. Part 12. Protorthoptera (continued), Neuroptera, additional Palaeodictyoptera, and families of uncertain position. *Psyche: A Journal of Entomology. 83*(3-4):336–376. https://doi.org/10.1155/1976/932123

Carpenter FM. 1979. Lower Permian insects from Oklahoma. Part 2. orders Ephemeroptera and Palaeodictyoptera. *Psyche: A Journal of Entomology. 86*(2-3):261–290. https://doi.org/10.1155/1979/96359

Carpenter FM. 1992. Superclass Hexapoda. In: *Treatise on Invertebrate Paleontology, Part R, Arthropoda 4, Volumes 3 and 4*. Boulder, Colorado and Lawrence, Kansas: Geological Society of America and University of Kansas. p. xxii + 655.

Carpenter FM, Richardson ES. 1968. Megasecopterous nymphs in Pennsylvanian concretions from Illinois. *Psyche: A Journal of Entomology. 75*:295–309. https://doi.org/10.1155/1968/903124

Carpenter FM, Richardson ES. 1971. Additional insects in Pennsylvanian concretions from Illinois. *Psyche: A Journal of Entomology. 78*:267–295. https://doi.org/10.1155/1971/80989

Case EC. 1904. Systematic Paleontology of the Miocene Deposits of Maryland: Mammalia. In: Clark WB, Shattuck GB, Dall WH, editors. *Miocene*. Baltimore: The Johns Hopkins Press. p. 3–58.

Case EC. 1934. A specimen of a long-nosed dolphin from the Bone Valley gravels of Polk county, Florida. *Contributions from the Museum of Paleontology, University of Michigan. 4*(6):105–113.

Cedillo-Ávila C, González-Barba G, Solis-Añorve A. 2025. First record of an Eomysticetidae from the late Oligocene at the Pilón locality, San Gregorio Formation, Baja California Sur, Mexico. *Palaeontologia Electronica. 28*(1):a1. https://doi.org/10.26879/1390

Césari SN, Chiesa JO. 2017. Palynology of the Bajo de Véliz Formation, central-western Argentina: Implications for Carboniferous–Permian transition biostratigraphy. *Journal of South American Earth Sciences. 78*:238–249. https://doi.org/10.1016/j.jsames.2017.07.004

Churchill M, Martínez-Cáceres M, de Muizon C, Mnieckowski J, Geisler JH. 2016. The origin of high-frequency hearing in whales. *Current Biology. 26*:2144–2149. https://doi.org/10.1016/j.cub.2016.06.004

Clark J. 1937. The stratigraphy and paleontology of the Chadron Formation in the Big Badlands of South Dakota. *Annals of the Carnegie Museum.* *25*:261–351.

Clyde WC, Stamatakos J, Gingerich PD. 1994. Chronology of the Wasatchian land-mammal age (early Eocene): Magnetostratigraphic results from the McCullough peaks section, northern Bighorn Basin, Wyoming. *The Journal of Geology.* *102*(4):367–377. https://doi.org/10.1086/629680

Colbert EH. 1944. A new fossil whale from the Miocene of Peru. *Bulletin of the American Museum of Natural History.* *83*:195–216.

Collareta A, Cigala Fulgosi F, Bianucci G. 2019. A new kogiid sperm whale from northern Italy supports psychrospheric conditions in the early Pliocene Mediterranean sea. *Acta Palaeontologica Polonica.* *64*(3):609–626. https://doi.org/10.4202/app.00578.2018

Collareta A, Lambert O, Landini W, Di Celma C, Malinverno E, Varas-Malca RM, Urbina M, Bianucci G. 2017. *Koristocetus pescei* gen. et sp. nov., a diminutive sperm whale (Cetacea: Odontoceti: Kogiidae) from the late Miocene of Peru. *Fossil Record.* *20*(2):259–278. https://doi.org/10.5194/fr-20-259-2017

Collareta A, Lambert O, Landini W, Malinverno E, Varas-Malca RM, Urbina M, Bianucci G. 2020. A new physeteroid from the late Miocene of Peru expands the diversity of extinct dwarf and pygmy sperm whales (Cetacea: Odontoceti: Kogiidae). *Comptes Rendus Palevol.* *19*(5):79–100. https://doi.org/10.5852/cr-palevol2020v19a5

Colpaert W, Bosselaers M, Lambert O. 2015. Out of the Pacific: A second fossil porpoise from the Pliocene of the North Sea basin. *Acta Palaeontologica Polonica.* *60*(1):1–10. https://doi.org/10.4202/app.00115.2014

Committee on Taxonomy. 2026. List of Marine Mammal Species and Subspecies. Society for Marine Mammalogy. https://marinemammalscience.org/science-and-publications/list-marine-mammal-species-subspecies/ last accessed 07/27/2026.

Cooper LN, Thewissen JGM, Hussain ST. 2009. New middle Eocene archaeocetes (Cetacea: Mammalia) from the Kuldana Formation of northern Pakistan. *Journal of Vertebrate Paleontology.* *29*(4):1289–1299. https://doi.org/10.1671/039.029.0423

Cope ED. 1867. An addition to the vertebrate fauna of the Miocene period, with a synopsis of the extinct Cetacea of the United States. *Proceedings of the Academy of Natural Sciences of Philadelphia.* *19*:138–156.

Cope ED. 1868. Second contribution to the history of the Vertebrata of the Miocene period of the United States. *Proceedings of the Academy of Natural Sciences of Philadelphia.* *20*:184–194.

Cope ED. 1869a. Remarks on *Eschrichtius polyporus*, *Hypsibema crassicauda*, *Hadrosaurus tripos*, and *Polydectes biturgidus*. *Proceedings of the Academy of Natural Sciences of Philadelphia.* *21*:191–192.

Cope ED. 1869b. Third contribution to the fauna of the Miocene period of the United States. *Proceedings of the Academy of Natural Sciences of Philadelphia*. *21*:6–13.

Cope ED. 1870. Synopsis of the extinct Batrachia, Reptilia and Aves of North America. *Transactions of the American Philosophical Society, New Series*. *14*(1):1–252.

Cope ED. 1872. On an extinct whale from California. *Proceedings of the Academy of Natural Sciences of Philadelphia*. *24*:29–30.

Cope ED. 1895. Fourth contribution to the marine fauna of the Miocene period of the United States. *Proceedings of the American Philosophical Society*. *34*:135–155.

Cope ED. 1896. Sixth contribution to the knowledge of the marine Miocene fauna of North America. *Proceedings of the American Philosophical Society*. *35*(151):139–146.

Copeland MJ. 1957. *The Arthropod Fauna of the Upper Carboniferous Rocks of the Maritime Provinces*. Memoir 286. Geological Survey of Canada. https://doi.org/10.4095/101505

Correia P, Nel A, Sá AA, Domingos R, Carneiro A, Flores D. 2014. A new Palaeodictyoptera from the Late Carboniferous of Portugal. *Annales de la Société entomologique de France (N.S.)* *49*(4):398–401. https://doi.org/10.1080/00379271.2014.893645

Correia P, Šimůnek Z, Cleal CJ, Vallois B, Domingos R, Sá AA. 2019. On a new species of the calamitalean fossil-genus *Annularia* from the Douro Basin (lower Gzhelian; NW Portugal). *Historical Biology*. *33*(2):258–267. https://doi.org/10.1080/08912963.2019.1613391

Corrie JE, Fordyce RE. 2022. A redescription and re-evaluation of *Kekenodon onamata* (Mammalia: Cetacea), a late-surviving archaeocete from the late Oligocene of New Zealand. *Zoological Journal of the Linnean Society*. *196*(4):1637–1670. https://doi.org/10.1093/zoolinnean/zlac019

Corrie JE, Fordyce RE. 2024. A new genus and species of kekenodontid from the late Oligocene of New Zealand with comments on the evolution of tooth displacement in Cetacea. *Journal of the Royal Society of New Zealand*. *54*(5):722–737. https://doi.org/10.1080/03036758.2023.2297696

Coste A, Fordyce RE, Loch C. 2023. A new dolphin with tusk-like teeth from the late Oligocene of New Zealand indicates evolution of novel feeding strategies. *Proceedings of the Royal Society B: Biological Sciences*. *290*(2000):20230873. https://doi.org/10.1098/rspb.2023.0873

Coste A, Fordyce RE, Loch C. 2024. A new fossil dolphin with tusk-like teeth from New Zealand and an analysis of procumbent teeth in fossil cetaceans. *Journal of the Royal Society of New Zealand*. *54*(5):738–757. https://doi.org/10.1080/03036758.2023.2267456

Cozzuol MA. 1988. Una nueva especie de *Saurodelphis* Burmeister, 1891 (Cetacea: Iniidae) del 'Mesopotamiense' (Mioceno Tardío–Plioceno Temprano) de la Provincia de Entre Ríos, Argentina. *Ameghiniana*. *25*(1):39–45.

Cuadrado-Ríos S, Vargas-Ramírez M, Kehlmaier C, Fritz U. 2025. Complete mitochondrial genome and phylogenetic analysis of the Atrato slider, *Trachemys medemi* (Testudines, Emydidae). *ZooKeys. 1224*:253–260. https://doi.org/10.3897/zookeys.1224.136863

Cuvier G. 1823. *Recherches sur les Ossemens Fossiles.* Paris: G. Dufour et E. d'Ocagne.

Czyżewska T, Ryziewicz Z. 1976. *Pinocetus polonicus* gen. n., sp. n. (Cetacea) from the Miocene limestones of Pińczów, Poland. *Acta Palaeontologica Polonica. 21*(3):259–291.

Dal Piaz G. 1901. Alcuni resti di *Squalodon* dell'arenaria miocenica di Belluno. *Palaeontographia Italica. 6*:303–314.

Dal Piaz G. 1904. *Neosqualodon*, nuovo genere della famiglia degli Squalodontidi. *Mémoires de la Société Paléontologique Suisse. 31*:1–19.

Dal Piaz G. 1908. Sui vertebrati delle arenarie mioceniche di Belluno. *Atti della Accademia Scientifica Veneto-Trentino-Istriana. 5*:106–120.

Dal Piaz G. 1916a. Gli Odontoceti del Miocene bellunese: *Cyrtodelphis. Memorie dell'Istituto Geologico della R. Università di Padova. 5*(5):1–18.

Dal Piaz G. 1916b. Gli Odontoceti del Miocene bellunese: *Eoplatanista italica. Memorie dell'Istituto Geologico della R. Università di Padova. 5*(2):1–23.

Dal Piaz G. 1916c. Gli Odontoceti del Miocene bellunese: *Scaldicetus bolzanensis. Memorie dell'Istituto Geologico della R. Università di Padova. 5*(9):86–92.

Dal Piaz G. 1916d. Gli Odontoceti del Miocene bellunese. Parte settima: *Protodelphinus capellinii. Memorie dell'Istituto Geologico della R. Università di Padova. 5*:42–50.

Dal Piaz G. 1917. Gli Odontoceti del Miocene bellunese, Parte Terza: *Squalodelphis fabianii. Memorie dell'Istituto Geologico della R. Università di Padova. 5*(1):1–34.

Dal Piaz G. 1971. *Guida dell'Istituto e del Museo di Geologia e Paleontologia.* Padua: Università degli Studi di Padova.

Dalebout ML, Baker CS, Mead JG, Cockcroft VG, Yamada TK. 2004. A comprehensive and validated molecular taxonomy of beaked whales, family Ziphiidae. *Journal of Heredity. 95*(6):459–473. https://doi.org/10.1093/jhered/esh054

Dalebout ML, Steel D, Baker CS. 2008. Phylogeny of the beaked whale genus *Mesoplodon* (Ziphiidae: Cetacea) revealed by nuclear introns: Implications for the evolution of male tusks. *Systematic Biology. 57*(6):857–875. https://doi.org/10.1080/10635150802559257

Dames WB. 1894. Über Zeuglodonten aus Ägypten und die beziehungen der Archäoceten zu den übrigen Cetaceen. *Geologische und Paläontologische Abhandlungen. 5*:189–222.

Davies SJ, Gibling MR, Rygel MC, Calder JH, Skilliter DM. 2005. The Pennsylvanian Joggins Formation of Nova Scotia: Sedimentological log and stratigraphic framework of the historic fossil cliffs. *Atlantic Geology*. *41*:115–142. https://doi.org/10.4138/182

Davydov VI, Biakov AS, Schmitz MD, Silantiev VV. 2018. Radioisotopic calibration of the Guadalupian (middle Permian) series: Review and updates. *Earth-Science Reviews*. *176*:222–240. https://doi.org/10.1016/j.earscirev.2017.10.011

Dawson SD. 1996. A new kentriodontid dolphin (Cetacea; Delphinoidea) from the middle Miocene Choptank formation, Maryland. *Journal of Vertebrate Paleontology*. *16*(1):135–140. https://doi.org/10.1080/02724634.1996.10011291

de Muizon C. 1983a. *Pliopontos littoralis*, un nouveau Platanistidae (Cetacea) du Pliocène de la côte péruvienne. *Comptes Rendus de l'Académie des Sciences de Paris, Série II*. *296*:625–628.

de Muizon C. 1983b. Un nouveau Phocoenidae (Cetacea) du Pliocène inférieur du Pérou. *Comptes Rendus de l'Académie des Sciences de Paris, Série II*. *296*:1203–1206.

de Muizon C. 1983c. Un Ziphiidae nouveau du Pliocène inférieur du Pérou. *Comptes Rendus de l'Académie des Sciences de Paris, Série II*. *297*:85–88.

de Muizon C. 1988. Les vertébrés fossiles de la formation Pisco (Pérou). troisième partie: Les Odontocètes (Cetacea, Mammalia) du Miocène. *Travaux de l'Institut Français d'Études Andines*. *42*:1–244.

de Muizon C. 1993. Walrus-like feeding adaptation in a new cetacean from the Pliocene of Peru. *Nature*. *365*:745–748. https://doi.org/10.1038/365745a0

de Muizon C, DeVries TJ. 1985. Geology and paleontology of late Cenozoic marine deposits in the Sacaco area (Peru). *Geologische Rundschau*. *74*(3):547–563. https://doi.org/10.1007/BF01821211

de Muizon C, Domning DP. 2002. The anatomy of *Odobenocetops* (Delphinoidea, Mammalia), the walrus-like dolphin from the Pliocene of Peru. *Zoological Journal of the Linnean Society*. *134*(4):423–452. https://doi.org/10.1046/j.1096-3642.2002.00015.x

de Muizon C, Domning DP, Parrish M. 1999. Dimorphic tusks and adaptive strategies in a new species of walrus-like dolphin (Odobenocetopsidae) from the Pliocene of Peru. *Comptes Rendus de l'Académie des Sciences, Series IIA, Earth and Planetary Science*. *329*:449–455.

De Schepper S, Head MJ. 2009. Pliocene and Pleistocene dinoflagellate cyst and acritarch zonation of DSDP Hole 610A, eastern North Atlantic. *Palynology*. *33*(1):179–218. https://doi.org/10.2113/gspalynol.33.1.179

De Stefano G. 1900. Gli avanzi fossili dei misticeti in Calabria. *Bollettino della Società Geologica Italiana*. *19*(2):281–294.

Deckers J, Louwye S, Goolaerts S. 2020. The internal division of the Pliocene Lillo Formation. *Geologica Belgica*. *23*(3-4):333–343. https://doi.org/10.20341/gb.2020.027

Dehm R, Oettingen-Spielberg Tz. 1958. Paläontologische und geologische untersuchungen im tertiär von Pakistan. 2. die mitteleocänen säugetiere von Ganda Kas bei Basal in Nordwest-Pakistan. *Abhandlungen der Bayerischen Akademie der Wissenschaften, Math. Naturwiss. Klasse, Neue Folge*. *91*:1–54.

Delfortrie E. 1873. Un Zeuglodon dans les faluns du Sud-Ouest de la France. *Actes de la Société Linnéenne de Bordeaux*. *29*:25–30.

Delfortrie E. 1875. Un dauphin d'espèce nouvelle dans les faluns du sud-ouest. *Actes de la Société Linnéenne de Bordeaux*. *30*(2):3–7.

Di Celma C, Malinverno E, Bosio G, Collareta A, Gariboldi K, Gioncada A, Molli G, Basso D, Varas-Malca RM, Pierantoni PP, et al. 2016. Stratigraphic framework of the late Miocene to Pliocene vertebrate-bearing Pisco formation at Cerro Los Quesos (Ica desert, Peru). *Journal of Maps*. *12*(5):1020–1028. https://doi.org/10.1080/17445647.2015.1115783

Di Celma C, Malinverno E, Collareta A, Bosio G, Gariboldi K, Gioncada A, Molli G, Basso D, Varas-Malca RM, Pierantoni PP, et al. 2017. Sequence stratigraphy and paleontology of the upper Miocene Pisco formation along the western side of the lower Ica valley (Ica desert, Peru). *Rivista Italiana di Paleontologia e Stratigrafia*. *123*(2):255–273. https://doi.org/10.13130/2039-4942/8373

Dickson MR. 1964. The skull and other remains of *Prosqualodon marplesi*, a new species of fossil whale. *New Zealand Journal of Geology and Geophysics*. *7*:626–635. https://doi.org/10.1080/00288306.1964.10422110

Dohrn A. 1866. *Eugereon Boeckingi*, eine neue Insektenform aus dem Todtliegenden. *Palaeontographica*. *13*:333–340.

Dominici S, Danise S, Benvenuti M. 2018. Pliocene stratigraphic paleobiology in Tuscany and the fossil record of marine megafauna. *Earth-Science Reviews*. *176*:277–310. https://doi.org/10.1016/j.earscirev.2017.09.018

Dooley A. C. J. 2005. A new species of *Squalodon* (Mammalia, Cetacea) from the middle Miocene of Virginia. *Memoir of the Virginia Museum of Natural History*. *8*:1–43.

Dooley A. C. J, Fraser NC, Luo ZX. 2004. The earliest known member of the rorqual–gray whale clade (Mammalia, Cetacea). *Journal of Vertebrate Paleontology*. *24*(2):453–463. https://doi.org/10.1671/2401

Dostál O, Prokop J. 2009. New fossil insects (Diaphanopterodea: Martynoviidae) from the Lower Permian of the Boskovice Basin, southern Moravia. *Geobios*. *42*(4):495–502. https://doi.org/10.1016/j.geobios.2009.01.004

Dowsett HJ, Robinson MM, Foley KM, Herbert TD. 2021. The Yorktown formation: Improved stratigraphy, chronology, and paleoclimate interpretations from the U.S. mid-Atlantic coastal plain. *Geosciences*. *11*(12):486. https://doi.org/10.3390/geosciences11120486

Du Bus B. 1872. Mammifères nouveaux du crag d'anvers. *Bulletins de l'Académie Royale des Sciences, des Lettres et des Beaux-Arts de Belgique*. *34*:491–509.

Du Bus BAL. 1867. Sur quelques mammifères du Crag d'Anvers. *Bulletins de l'Académie Royale des Sciences, des Lettres et des Beaux-Arts de Belgique, 2e série*. *24*:562–577.

Du Bus BAL. 1868. Sur différents ziphiides nouveaux du Crag d'Anvers. *Bulletins de l'Académie Royale des Sciences, des Lettres et des Beaux-Arts de Belgique, 2e série*. *25*:621–630.

Duboys de Lavigerie G, Bosselaers M, Goolaerts S, Park T, Lambert O, Marx FG. 2020. New Pliocene right whale from Belgium informs balaenid phylogeny and function. *Journal of Systematic Palaeontology*. *18*(14):1141–1166. https://doi.org/10.1080/14772019.2020.1746422

Dubrovo IA. 1971. Novyi rod kitoobraznykh (*Sachalinocetus cholmicus* gen. et sp. nov.) iz miotsena o-va Sakhalin. *Trudy Paleontologicheskogo Instituta, Akademiya Nauk SSSR*. *130*:87–103.

Dubrovo IA, Sanders AE. 2000. A new species of *Patriocetus* (Mammalia, Cetacea) from the late Oligocene of Kazakhstan. *Journal of Vertebrate Paleontology*. *20*(3):577–590. https://doi.org/10.1671/0272-4634(2000)020[0577:ANSOPM]2.0.CO;2

Dunbar CO, Tillyard RJ. 1924. Kansas Permian insects. Part 1, the geologic occurrence and the environment of the insects. *American Journal of Science*. *207*(39):171–209. https://doi.org/10.2475/ajs.s5-7.39.171

Duncan RJ, Rule JP, Park T, Evans AR, Adams JW, Fitzgerald EMG. 2025. An immature toothed mysticete from the Oligocene of Australia and insights into mammalodontid (Cetacea: Mysticeti) morphology, systematics, and ontogeny. *Zoological Journal of the Linnean Society*. *204*(4):zlaf090. https://doi.org/10.1093/zoolinnean/zlaf090

Eddy MP, Bowring SA, Umhoefer PJ, Miller RB, McLean NM, Donaghy EE. 2016. High-resolution temporal and stratigraphic record of Siletzia's accretion and triple junction migration from nonmarine sedimentary basins in central and western Washington. *Geological Society of America Bulletin*. *128*(3-4):425–441. https://doi.org/10.1130/B31335.1

Ehret DJ, Bourque JR. 2011. An extinct map turtle *Graptemys* (Testudines, Emydidae) from the late Pleistocene of Florida. *Journal of Vertebrate Paleontology*. *31*(3):575–587. https://doi.org/10.1080/02724634.2011.562267

Ehret DJ, Bourque JR, Hulbert R. C. J. 2013. Case 3628. *Terrapene putnami* Hay, 1906 (Reptilia, Testudines, Emydidae): Replacement of the holotype by designation of a neotype. *Bulletin of Zoological Nomenclature*. *70*(3):193–198. https://doi.org/10.21805/bzn.v70i3.a13

El Adli JJ, Deméré TA, Boessenecker RW. 2014. *Herpetocetus morrowi* (Cetacea: Mysticeti), a new species of diminutive baleen whale from the Upper Pliocene (Piacenzian) of California, USA, with observations on the evolution and relationships of the Cetotheriidae. *Zoological Journal of the Linnean Society. 170*(2):400–466. https://doi.org/10.1111/zoj.12108

Emlong DR. 1966. A new archaic cetacean from the Oligocene of Northwest Oregon. *Bulletin of the Museum of Natural History, University of Oregon. 3*:1–51.

Fajardo-Mellor L, Berta A, Brownell RL, Boy CC, Goodall RNP. 2006. The phylogenetic relationships and biogeography of true porpoises (Mammalia: Phocoenidae) based on morphological data. *Marine Mammal Science. 22*(4):910–932. https://doi.org/10.1111/j.1748-7692.2006.00080.x

Figueiredo R, Bosselaers M, Póvoas L, Castanhinha R. 2024. Redescription of three fossil baleen whale skulls from the Miocene of Portugal reveals new cetotheriid phylogenetic insights. *PLoS One. 19*(3):e0298658. https://doi.org/10.1371/journal.pone.0298658

Fischer JB. 1829. *Synopsis Mammalium.* Stuttgart: J. G. Cotta.

Fitzgerald EMG. 2006. A bizarre new toothed mysticete (Cetacea) from Australia and the early evolution of baleen whales. *Proceedings of the Royal Society B: Biological Sciences. 273*(1604):2955–2963. https://doi.org/10.1098/rspb.2006.3664

Flot L. 1896. Note sur les cétacés fossiles de l'Aquitaine. *Bulletin de la Société Géologique de France, 3e série. 24*:270–282.

Fordyce RE. 1994. *Waipatia maerewhenua*, new genus and new species (Waipatiidae, new family), an archaic late Oligocene dolphin (Cetacea: Odontoceti: Platanistoidea) from New Zealand. *Proceedings of the San Diego Society of Natural History. 29*:147–176. https://doi.org/10.5962/bhl.part.10662

Fordyce RE. 2002. *Simocetus rayi* (Odontoceti: Simocetidae, new family): A bizarre new archaic Oligocene dolphin from the eastern North Pacific. *Smithsonian Contributions to Paleobiology. 93*:185–222. https://doi.org/10.5479/si.00810266.93.185

Fordyce RE, Marx FG. 2016. Mysticetes baring their teeth: A new fossil whale, *Mammalodon hakataramea*, from the southwest pacific. *Memoirs of Museum Victoria. 74*:107–116. https://doi.org/10.24199/j.mmv.2016.74.11

Fordyce RE, Marx FG. 2018. Gigantism precedes filter feeding in baleen whale evolution. *Current Biology. 28*(10):1670–1676. https://doi.org/10.1016/j.cub.2018.04.027

Fordyce RE, Quilty PG, Daniels J. 2002. *Australodelphis mirus*, a bizarre new toothless ziphiid-like fossil dolphin (Cetacea: Delphinidae) from the Pliocene of Vestfold Hills, East Antarctica. *Antarctic Science. 14*(1):37–54. https://doi.org/10.1017/S0954102002000561

Fraas E. 1904. Neue Zeuglodonten aus dem unteren Mitteleocän vom Mokattam bei Cairo. *Geologische und Palaeontologische Abhandlungen. 6*(3):199–220.

Fritsch A. 1880. Fauna der Gaskohle und der Kalksteine der Permformation Böhmens. Band I, Heft 2. In: Prag: F. Řivnáč. p. 93–126.

Fritz U, Herrmann HW, Rosen PC, Auer M, Vargas-Ramírez M, Kehlmaier C. 2024. *Trachemys* in Mexico and beyond: Beautiful turtles, taxonomic nightmare, and a mitochondrial poltergeist (Testudines: Emydidae). *Vertebrate Zoology. 74*:435–452. https://doi.org/10.3897/vz.74.e125958

Fu Y, Béthoux O, Yang Q, Ren D. 2015. The earliest and most oriental Calvertiellidae unearthed (Palaeodictyoptera; Late Carboniferous; China). *Insect Systematics & Evolution. 46*(5):485–492. https://doi.org/10.1163/1876312X-46052128

Fujiwara Y, Irizuki T, Hayashi H, Tanaka Y. 2005. Stratigraphy and age of the Miocene Awa Group, eastern Iga City, Mie prefecture, central Japan. *The Journal of the Geological Society of Japan. 111*(12):779–791. https://doi.org/10.5575/geosoc.111.779

Gaetán CM, Buono MR, Gaetano LC. 2018. *Prosqualodon australis* (Cetacea: Odontoceti) from the Early Miocene of Patagonia, Argentina: Redescription and phylogenetic analysis. *Ameghiniana. 56*(1):1–27. https://doi.org/10.5710/AMGH.21.11.2018.3208

Gaetán CM, Paolucci F, Buono MR. 2023. A new squaloziphiid-like odontocete from the Early Miocene of Patagonia expands the cetacean diversity in the southwestern Atlantic Ocean. *Journal of Vertebrate Paleontology. 42*(6):e2232425. https://doi.org/10.1080/02724634.2023.2232425

Gastaldo RA, Kamo SL, Neveling J, Geissman JW, Looy CV, Martini AM. 2020. The base of the *Lystrosaurus* Assemblage Zone, Karoo Basin, predates the end-Permian marine extinction. *Nature Communications. 11*:1428. https://doi.org/10.1038/s41467-020-15243-7

Geisler JH, Boessenecker RW, Brown M, Beatty BL. 2017. The origin of filter feeding in whales. *Current Biology. 27*(13):2036–2042. https://doi.org/10.1016/j.cub.2017.06.003

Geisler JH, Colbert MW, Carew JL. 2014. A new fossil species supports an early origin for toothed whale echolocation. *Nature. 508*(7496):383–386. https://doi.org/10.1038/nature13086

Geisler JH, Godfrey SJ, Lambert O. 2012. A new genus and species of late Miocene inioid (Cetacea, Odontoceti) from the Meherrin River, North Carolina, U.S.A. *Journal of Vertebrate Paleontology. 32*(1):198–211. https://doi.org/10.1080/02724634.2012.629016

Geisler JH, McGowen MR, Yang G, Gatesy J. 2011. A supermatrix analysis of genomic, morphological, and paleontological data from crown Cetacea. *BMC Evolutionary Biology. 11*:112. https://doi.org/10.1186/1471-2148-11-112

Geisler JH, Sanders AE, Luo ZX. 2005. A new protocetid whale (Cetacea: Archaeoceti) from the late middle Eocene of South Carolina. *American Museum Novitates. 3480*:1–65. https://doi.org/10.1206/0003-0082(2005)480[0001:ANPWCA]2.0.CO;2

Gervais P. 1849. *Zoologie et Paléontologie Françaises (Animaux Vertébrés)*. Paris: Arthus Bertrand.

Gervais P. 1859. *Zoologie et Paléontologie Françaises (Animaux Vertébrés), ou Nouvelles Recherches sur les Animaux Vivants et Fossiles de la France*. 2nd ed. Paris: Arthus Bertrand.

Gervais P. 1861. Sur différentes espèces de vertébrés fossiles observées pour la plupart dans le Midi de la France. *Mémoires de la Section des Sciences, Académie des Sciences et Lettres de Montpellier*. *5*:117–132.

Getty PR, Sproule R, Stimson MR, Lyons PC. 2017. Invertebrate trace fossils from the Pennsylvanian Rhode Island Formation of Massachusetts, USA. *Atlantic Geology*. *53*:185–206. https://doi.org/10.4138/atlgeol.2017.007

Gibbes RW. 1845. Description of the teeth of a new fossil animal found in the Green Sand of South Carolina. *Proceedings of the Academy of Natural Sciences of Philadelphia*. *2*:254–256.

Gibson ML, Geisler JH. 2009. A new Pliocene dolphin (Cetacea: Pontoporiidae), from the Lee Creek mine, North Carolina. *Journal of Vertebrate Paleontology*. *29*(3):966–971. https://doi.org/10.1671/039.029.0307

Gibson ML, Mnieckowski J, Geisler JH. 2019. *Tupelocetus palmeri*, a new species of protocetid whale (Mammalia, Cetacea) from the middle Eocene of South Carolina. *Journal of Vertebrate Paleontology*. *38*(6):e1555165. https://doi.org/10.1080/02724634.2018.1555165

Gilmore CW. 1927. On fossil turtles from the Pleistocene of Florida. *Proceedings of the United States National Museum*. *71*(6):1–10. https://doi.org/10.5479/si.00963801.71-2687.1

Gilmore CW. 1933. A new species of extinct turtle from the upper Pliocene of Idaho. *Proceedings of the United States National Museum*. *82*(9):1–7. https://doi.org/10.5479/si.00963801.82-2950.1

Gingerich PD. 1992. Marine mammals (Cetacea and Sirenia) from the Eocene of Gebel Mokattam and Fayum, Egypt: Stratigraphy, age, and paleoenvironments. *University of Michigan Papers on Paleontology*. *30*:1–84.

Gingerich PD. 2007. *Stromerius nidensis*, new archaeocete (Mammalia, Cetacea) from the upper Eocene Qasr el-Sagha Formation, Fayum, Egypt. *Contributions from the Museum of Paleontology, University of Michigan*. *31*(13):363–378.

Gingerich PD. 2015. New partial skeleton and relative brain size in the late Eocene archaeocete *Zygorhiza kochii* (Mammalia, Cetacea) from the Pachuta Marl of Alabama, with a note on contemporaneous *Pontogeneus brachyspondylus*. *Contributions from the Museum of Paleontology, University of Michigan*. *32*(10):161–188.

Gingerich PD, Antar MSM, Zalmout IS. 2019. *Aegicetus gehennae*, a new late Eocene protocetid (Cetacea, Archaeoceti) from Wadi Al Hitan, Egypt, and the transition to tail-powered swimming in whales. *PLoS One*. *14*(12):e0225391. https://doi.org/10.1371/journal.pone.0225391

Gingerich PD, Arif M, Bhatti MA, Anwar M, Sanders WJ. 1997. *Basilosaurus drazindai* and *Basiloterus hussaini*, new Archaeoceti (Mammalia, Cetacea) from the middle Eocene Drazinda Formation, Sulaiman Range, Punjab (Pakistan). *Contributions from the Museum of Paleontology, University of Michigan*. *30*(2):55–81.

Gingerich PD, Arif M, Clyde WC. 1995. New archaeocetes (Mammalia, Cetacea) from the middle Eocene Domanda Formation of the Sulaiman Range, Punjab (Pakistan). *Contributions from the Museum of Paleontology, University of Michigan*. *29*(11):291–330.

Gingerich PD, Cappetta H. 2014. A new archaeocete and other marine mammals (Cetacea and Sirenia) from lower middle Eocene phosphate deposits of Togo. *Journal of Paleontology*. *88*(1):109–129. https://doi.org/10.1666/13-040

Gingerich PD, ul-Haq M, Khan IH, Zalmout IS. 2001a. Eocene stratigraphy and archaeocete whales (Mammalia, Cetacea) of Drug Lahar in the eastern Sulaiman Range, Balochistan (Pakistan). *Contributions from the Museum of Paleontology, University of Michigan*. *30*(11):269–319.

Gingerich PD, ul-Haq M, von Koenigswald W, Sanders WJ, Smith BH, Zalmout IS. 2009. New protocetid whale from the middle Eocene of Pakistan: Birth on land, precocial development, and sexual dimorphism. *PLoS One*. *4*(2):e4366. https://doi.org/10.1371/journal.pone.0004366

Gingerich PD, ul-Haq M, Zalmout IS, Khan IH, Malkani MS. 2001b. Origin of whales from early artiodactyls: Hands and feet of Eocene Protocetidae from Pakistan. *Science*. *293*(5538):2239–2242. https://doi.org/10.1126/science.1063902

Gingerich PD, Raza SM, Arif M, Anwar M, Zhou X. 1994. New whale from the Eocene of Pakistan and the origin of cetacean swimming. *Nature*. *368*:844–847. https://doi.org/10.1038/368844a0

Gingerich PD, Russell DE. 1981. *Pakicetus inachus*, a new archaeocete (Mammalia, Cetacea) from the early-middle Eocene Kuldana Formation of Kohat (Pakistan). *Contributions from the Museum of Paleontology, University of Michigan*. *25*(11):235–246.

Gingerich PD, Uhen MD. 1996. *Ancalecetus simonsi*, a new dorudontine archaeocete (Mammalia, Cetacea) from the early late Eocene of Wadi Hitan, Egypt. *Contributions from the Museum of Paleontology, University of Michigan*. *29*(13):359–401.

Gingerich PD, Zalmout IS, ul-Haq M, Bhatti MA. 2005. *Makaracetus bidens*, a new protocetid archaeocete (Mammalia, Cetacea) from the early middle Eocene of Balochistan (Pakistan). *Contributions from the Museum of Paleontology, University of Michigan*. *31*(9):197–210.

Gingerich PD, Zouhri S. 2015. New fauna of archaeocete whales (Mammalia, Cetacea) from the Bartonian middle Eocene of southern Morocco. *Journal of African Earth Sciences*. *111*:273–286. https://doi.org/10.1016/j.jafrearsci.2015.08.006

Ginsburg L, Janvier P. 1971. Les mammifères marins des faluns miocènes de la Touraine et de l'Anjou. *Bulletin du Muséum National d'Histoire Naturelle, Sciences de la Terre.* *6*(22):161–195.

Godfrey SJ, Barnes LG. 2008. A new genus and species of late Miocene pontoporiid dolphin (Cetacea: Odontoceti) from the St. Marys formation in Maryland. *Journal of Vertebrate Paleontology.* *28*(2):520–528. https://doi.org/10.1671/0272-4634(2008)28[520:ANGASO]2.0.CO;2

Godfrey SJ, Gutstein CS, Morgan DJ. 2021. A new odontocete (Inioidea, Odontoceti) from the late Neogene of North Carolina, USA. *Fossil Record.* *24*:275–285. https://doi.org/10.5194/fr-24-275-2021

Godfrey SJ, Lambert O. 2023. Miocene Toothed Whales (Odontoceti) from Calvert Cliffs, Atlantic Coastal Plain, USA. In: *The Geology and Vertebrate Paleontology of Calvert Cliffs, Maryland, USA, Volume 2: Turtles and Toothed Whales.* Washington, D.C.: Smithsonian Institution Scholarly Press. p. 49–186.

Godfrey SJ, Uhen MD, Osborne JE, Edwards LE. 2016. A new specimen of *Agorophius pygmaeus* (Agorophiidae, Odontoceti, Cetacea) from the early Oligocene Ashley formation of South Carolina, USA. *Journal of Paleontology.* *90*(1):154–169. https://doi.org/10.1017/jpa.2016.4

Gohar AS, Antar MS, Boessenecker RW, Sabry DA, El-Sayed S, Seiffert ER, Zalmout IS, Sallam HM. 2021. A new protocetid whale offers clues to biogeography and feeding ecology in early cetacean evolution. *Proceedings of the Royal Society B: Biological Sciences.* *288*(1957):20211368. https://doi.org/10.1098/rspb.2021.1368

Gol'din P. 2018. New Paratethyan dwarf baleen whales mark the origin of cetotheres. *PeerJ.* *6*:e5800. https://doi.org/10.7717/peerj.5800

Gol'din P, Startsev D. 2014. *Brandtocetus*, a new genus of baleen whales (Cetacea, Cetotheriidae) from the late Miocene of Crimea, Ukraine. *Journal of Vertebrate Paleontology.* *34*(2):419–433. https://doi.org/10.1080/02724634.2013.799482

Gol'din P, Startsev D. 2017. A systematic review of cetothere baleen whales (Cetacea, Cetotheriidae) from the late Miocene of Crimea and Caucasus, with a new genus. *Papers in Palaeontology.* *3*(1):49–68. https://doi.org/10.1002/spp2.1066

Gol'din P, Startsev D, Krakhmalnaya T. 2014. The anatomy of the late Miocene baleen whale *Cetotherium riabinini* from Ukraine. *Acta Palaeontologica Polonica.* *59*(4):795–814. https://doi.org/10.4202/app.2012.0107

Gol'din P, Steeman ME. 2015. From problem taxa to problem solver: A new Miocene family, Tranatocetidae, brings perspective on baleen whale evolution. *PLoS One.* *10*(9):e0135500. https://doi.org/10.1371/journal.pone.0135500

Goldenberg F. 1854. *Die fossilen Insecten der Kohlenformation von Saarbrücken.* Cassel: Th. Fischer.

Goldenberg F. 1877. *Die fossilen Thiere aus der Steinkohlenformation von Saarbrücken. Fauna Saraepontana Fossilis, Heft 2*. Saarbrücken: Mollinger, 54.

Gondar D. 1975. La presencia de cetáceos Physeteridae en el Terciario superior ("Rionegrense") de la provincia de Río Negro. In: *Actas del Primer Congreso Argentino de Paleontología y Bioestratigrafía*. Vol. 2. Tucumán: Asociación Paleontológica Argentina. p. 349–356.

Goolaerts S, De Ceuster J, Mollen FH, Gijsen B, Bosselaers M, Lambert O, Uchman A, Van Herck M, Adriaens R, Houthuys R, et al. 2020. The upper Miocene Deurne Member of the Diest Formation revisited: Unexpected results from the study of a large temporary outcrop near Antwerp International Airport, Belgium. *Geologica Belgica*. *23*(3-4):219–252. https://doi.org/10.20341/gb.2020.011

Gorjanović-Kramberger D. 1892. O fosilnih cetaceih Hrvatske i Kranjske. *Rad Jugoslavenske Akademije Znanosti i Umjetnosti*. *111*:1–21.

Gradstein FM, Ogg JG, Schmitz MD, Ogg GM, eds. 2020. *Geologic Time Scale 2020*. Elsevier. https://doi.org/10.1016/C2020-1-02369-3

Graham IJ, Morgans HEG, Waghorn DB, Trotter JA, Whitford DJ. 2000. Strontium isotope stratigraphy of the Oligocene–Miocene Otekaike Limestone (Trig Z section) in southern New Zealand: Age of the Duntroonian/Waitakian stage boundary. *New Zealand Journal of Geology and Geophysics*. *43*(3):335–347. https://doi.org/10.1080/00288306.2000.9514891

Gregory WK, Kellogg R. 1927. A fossil porpoise from California. *American Museum Novitates*. *269*:1–7.

Guo Z, Kohno N. 2021. A new kentriodontid (Cetacea: Odontoceti) from the early to middle Miocene of the western North Pacific and a revision of kentriodontid phylogeny. *PeerJ*. *9*:e10945. https://doi.org/10.7717/peerj.10945

Guo Z, Kohno N. 2023. An early Miocene kentriodontoid (Cetacea: Odontoceti) from the western North Pacific, and its implications for their phylogeny and paleobiogeography. *PLoS One*. *18*(2):e0280218. https://doi.org/10.1371/journal.pone.0280218

Guthörl P. 1930. Neue Insektenfunde aus dem Saarcarbon. *Neues Jahrbuch für Mineralogie, Geologie und Paläontologie, Abteilung B*. *64*:147–164.

Guthörl P. 1931. *Cleffia sarana* n. gen. n. sp., eine neue insektenform aus dem saarcarbon. *Centralblatt für Mineralogie, Geologie und Paläontologie, Abteilung B*. *1931*:91–94.

Guthörl P. 1934. Die Arthropoden aus dem Carbon und Perm des Saar-Nahe-Pfalz-Gebietes. *Abhandlungen der Preußischen Geologischen Landesanstalt, Neue Folge*. *164*:1–219.

Guthörl P. 1939. Zur Arthropoden-Fauna des Karbons und Perms. 9. Palaeodictyoptera, Mixotermitoidea, Miomoptera und Blattariae. *Senckenbergiana*. *21*:314–329.

Guthörl P. 1940. Zur Arthropoden-Fauna des Karbons und Perms. 11. Neue Insektenfunde (Palaeodictyoptera) aus Schacht 5 der Grube Frankenholz-Saar. *Senckenbergiana. 22*:35–73.

Handlirsch A. 1904. Les insectes houillers de la Belgique. *Mémoires du Musée royal d'Histoire naturelle de Belgique. 3*:1–20.

Handlirsch A. 1906a. *Die Fossilen Insekten und die Phylogenie der Rezenten Formen: Ein Handbuch für Paläontologen und Zoologen.* Leipzig: Wilhelm Engelmann, 1430. https://doi.org/10.5962/bhl.title.34145

Handlirsch A. 1906b. Revision of American Paleozoic insects. *Proceedings of the United States National Museum. 29*(1441):661–820. https://doi.org/10.5479/si.00963801.29-1441.661

Handlirsch A. 1908. *Die Fossilen Insekten und die Phylogenie der Rezenten Formen: Ein Handbuch für Paläontologen und Zoologen.* Leipzig: Wilhelm Engelmann, 1121–1430.

Handlirsch A. 1910. Das erste fossile Insekt aus dem Oberkarbon Westfalens. *Verhandlungen der Kaiserlich-Königlichen Zoologisch-Botanischen Gesellschaft in Wien. 60*:249–251.

Handlirsch A. 1911. New Paleozoic insects from the vicinity of Mazon Creek, Illinois. *American Journal of Science, 4th series. 31*:297–326, 353–377. https://doi.org/10.2475/ajs.s4-31.185.353

Handlirsch A. 1919. Revision der paläozoischen Insekten. *Denkschriften der Akademie der Wissenschaften in Wien, Mathematisch-Naturwissenschaftliche Klasse. 96*:511–592.

Harlan R. 1842. Description of a new extinct species of dolphin from Maryland. *Bulletin of the Proceedings of the National Institute for the Promotion of Science. 2*:195–196.

Harzhauser M, Böhme M, Mandic O, Hofmann CC. 2002. The Karpatian (late Burdigalian) of the Korneuburg basin — a palaeoecological and biostratigraphical synthesis. *Beiträge zur Paläontologie. 27*:441–456.

Hatai K, Hayasaka S, Masuda K. 1963. Some fossil tympanics from the Mizuho period of northern Japan. *Saito Ho-on Kai Museum Research Bulletin. 32*:5–17.

Haug C, Haug JT. 2017. The presumed oldest flying insect: More likely a myriapod? *PeerJ. 5*:e3402. https://doi.org/10.7717/peerj.3402

Hay OP. 1899. Descriptions of two new species of tortoises from the Tertiary of the United States. *Proceedings of the United States National Museum. 22*:21–24. https://doi.org/10.5479/si.00963801.22-1181.21

Hay OP. 1906. Descriptions of two new genera (*Echmatemys* and *Xenochelys*) and two new species (*Xenochelys formosa* and *Terrapene putnami*) of fossil turtles. *Bulletin of the American Museum of Natural History. 22*:27–31.

Hay OP. 1908. *The Fossil Turtles of North America.* Publication No. 75. Washington, D.C.: Carnegie Institution of Washington.

Hector J. 1881. Notes on New Zealand Cetacea, recent and fossil. *Transactions of the New Zealand Institute. 13*:434–436.

Hernández-Cisneros AE. 2018. A new group of late Oligocene mysticetes from México. *Palaeontologia Electronica. 21*(1):1–30. https://doi.org/10.26879/746

Hernández-Cisneros AE. 2022. A new aetiocetid (Cetacea, Mysticeti, Aetiocetidae) from the Late Oligocene of Mexico. *Journal of Systematic Palaeontology. 20*(1):1–15. https://doi.org/10.1080/14772019.2022.2100725

Hernández-Cisneros AE, Schwennicke T, Rochín-Bañaga H, Tsai CH. 2023. *Echericetus novellus* n. gen. n. sp. (Cetacea, Mysticeti, Eomysticetidae), an Oligocene baleen whale from Baja California Sur, Mexico. *Journal of Paleontology. 97*(6):1309–1328. https://doi.org/10.1017/jpa.2023.80

Hirota K, Barnes LG. 1995. A new species of middle Miocene sperm whale of the genus *Scaldicetus* (Cetacea; Physeteridae) from Shiga-mura, Japan. *The Island Arc. 3*(4):453–472. https://doi.org/10.1111/j.1440-1738.1994.tb00125.x

Hoch E. 2004. Cuvierian analysis, and determination of a delphinoid cetacean from the late Miocene North Sea. *Revue de Paléobiologie. 9*:1–15.

Hofstein ID. 1948. Pakhyostoz u vykopnykh kytiv. *Zbirnyk Prats z Paleontologii i Stratygrafii, Instytut Geologichnykh Nauk URSR. 1*(2):65–75.

Hohenegger J, Ćorić S, Wagreich M. 2014. Timing of the Middle Miocene Badenian stage of the Central Paratethys. *Geologica Carpathica. 65*(1):55–66. https://doi.org/10.2478/geoca-2014-0004

Holl F. 1829. *Handbuch der Petrefactenkunde.* Vol. 1. Dresden: P. G. Hilscher'sche Buchhandlung.

Holman JA, Fritz U. 2001. A new emydine species from the middle Miocene (Barstovian) of Nebraska, USA with a new generic arrangement for the species of *Clemmys* sensu McDowell (1964) (Reptilia: Testudines: Emydidae). *Zoologische Abhandlungen (Dresden). 51*(2):331–353.

Holman JA, Fritz U. 2005. The box turtle genus *Terrapene* (Testudines: Emydidae) in the Miocene of the USA. *The Herpetological Journal. 15*(2):81–90.

Hong YC. 1985a. New fossil genera and species of Shanxi Formation in Xishan of Taiyuan. *Entomotaxonomia. 7*:83–91.

Hong YC. 1985b. *Palaeontological Atlas of North China. Volume 1: Palaeozoic.* Beijing: Geological Publishing House, 492–510.

Horikawa H. 1977. Hokkaido Kabato-sanchi, chushintosan no Sujiiruka-zoku no shinshu ni tsuite. *Chikyu Kagaku. 31*(3):97–114.

Hulbert RC, Petkewich RM, Bishop GA, Bukry D, Aleshire DP. 1998. A new middle Eocene protocetid whale (Mammalia: Cetacea: Archaeoceti) and associated biota from Georgia. *Journal of Paleontology. 72*(5):907–927. https://doi.org/10.1017/S0022336000027232

Hunt AP, Lucas SG, Pyenson ND. 2005. The Significance of the Union Chapel Mine Site: A Lower Pennsylvanian (Westphalian A) Ichnological Konzentrat-Lagerstätte, Alabama, USA. In: Buta RJ, Rindsberg AK, Kopaska-Merkel DC, editors. *Pennsylvanian Footprints in the Black Warrior Basin of Alabama*. p. 3–14.

Hutchison JH. 2013. New Turtles from the Paleogene of North America. In: Brinkman DB, Holroyd PA, Gardner JD, editors. *Morphology and Evolution of Turtles*. Dordrecht: Springer. p. 477–497. https://doi.org/10.1007/978-94-007-4309-0_26

Huxley TH. 1864. On the cetacean fossils termed "*Ziphius*" by Cuvier, with a notice of a new species (*Belemnoziphius compressus*) from the Red Crag. *Quarterly Journal of the Geological Society*. *20*(1-2):388–396. https://doi.org/10.1144/gsl.jgs.1864.020.01-02.48

Ichishima H. 1995. A new fossil kentriodontid dolphin (Cetacea; Kentriodontidae) from the middle Miocene Takinoue formation, Hokkaido, Japan. *Island Arc*. *3*(4):473–485. https://doi.org/10.1111/j.1440-1738.1994.tb00126.x

Ichishima H, Furusawa H, Tachibana M, Kimura M. 2019. First monodontid cetacean (Odontoceti, Delphinoidea) from the early Pliocene of the north-western Pacific Ocean. *Papers in Palaeontology*. *5*(2):323–342. https://doi.org/10.1002/spp2.1244

Ichishima H, Kimura M. 2000. A new fossil porpoise (Cetacea; Delphinoidea; Phocoenidae) from the early Pliocene Horokaoshirarika formation, Hokkaido, Japan. *Journal of Vertebrate Paleontology*. *20*(3):561–576. https://doi.org/10.1671/0272-4634(2000)020[0561:ANFPCD]2.0.CO;2

Ichishima H, Kimura M. 2005. *Haborophocaena toyoshimai*, a new early Pliocene porpoise (Cetacea; Phocoenidae) from Hokkaido, Japan. *Journal of Vertebrate Paleontology*. *25*(3):655–664. https://doi.org/10.1671/0272-4634(2005)025[0655:HTANEP]2.0.CO;2

Ichishima H, Kimura M. 2009. A new species of *Haborophocaena*, an early Pliocene phocoenid cetacean from Hokkaido, Japan. *Marine Mammal Science*. *25*(4):855–874. https://doi.org/10.1111/j.1748-7692.2009.00293.x

Ilger JM, Brauckmann C. 2011. The smallest Neoptera (Baryshnyalidae fam. n.) from Hagen-Vorhalle (early Late Carboniferous: Namurian B; Germany). *ZooKeys*. *130*:91–102. https://doi.org/10.3897/zookeys.130.1422

Ivanov VV. 2000. A new species of Dictyoneurida from the Kungurian deposits of the Middle Urals. *Paleontological Journal*. *34*(Suppl. 3):S288–S289.

Ives LRW, Isbell JL, Licht A. 2022. A 'local first' approach to glacigenic sediment provenance demonstrated using U–Pb detrital zircon geochronology of the Permo-Carboniferous Wynyard Formation, Tasmanian Basin. *The Sedimentary Record*. *20*(1). https://doi.org/10.2110/001c.38180

Jackson DR. 1978. The late Miocene fossil record of the chicken turtle *Deirochelys*, with a re-evaluation of the genus. *Tulane Studies in Zoology and Botany. 20*(1):35–55.

Jasinski SE. 2018. A new slider turtle (Testudines: Emydidae: Deirochelyinae: *Trachemys*) from the late Hemphillian (late Miocene/early Pliocene) of eastern Tennessee and the evolution of the deirochelyines. *PeerJ. 6*:e4338. https://doi.org/10.7717/peerj.4338

Jasinski SE. 2022. A new species of *Chrysemys* (Emydidae: Deirochelyinae) from the latest Miocene–early Pliocene of Tennessee, USA and its implications for the evolution of painted turtles. *Zoological Journal of the Linnean Society. 198*(1):149–183. https://doi.org/10.1093/zoolinnean/zlac084

Joyce WG, Parham JF, Lyson TR, Warnock RCM, Donoghue PCJ. 2013. A divergence dating analysis of turtles using fossil calibrations: An example of best practices. *Journal of Paleontology. 87*(4):612–634. https://doi.org/10.1666/12-149

Joyce WG, Petričević A, Lyson TR, Czaplewski NJ. 2012. A new box turtle from the Miocene/Pliocene boundary (latest Hemphillian) of Oklahoma and a refined chronology of box turtle diversification. *Journal of Paleontology. 86*(1):177–190. https://doi.org/10.1666/11-073.1

Kawatani A, Kohno N. 2021. The oldest fossil record of the extant genus *Berardius* (Odontoceti, Ziphiidae) from the middle to late Miocene boundary of the western North Pacific. *Royal Society Open Science. 8*(3):201152. https://doi.org/10.1098/rsos.201152

Kazár E. 2005. A new kentriodontid (Cetacea: Delphinoidea) from the middle Miocene of Hungary. *Fossil Record. 8*(1):53–73. https://doi.org/10.1002/mmng.200410004

Kazár E. 2006. *Sophianaecetus*, a replacement name for *Mediocris* (Cetacea: Delphinoidea: Kentriodontidae). *Fossil Record. 9*(2):260. https://doi.org/10.1002/mmng.200600013

Kazár E. 2010. Revision of the genus *Pachyacanthus* brandt, 1871 (mammalia: Cetacea: Odontoceti). *Annalen des Naturhistorischen Museums in Wien, Serie A. 112*:537–567.

Kazár E, Grigorescu D. 2005. Revision of *Sarmatodelphis moldavicus* Kirpichnikov, 1954 (Cetacea: Delphinoidea), from the Miocene of Kishinev, republic of Moldavia. *Journal of Vertebrate Paleontology. 25*(4):929–935. https://doi.org/10.1671/0272-4634(2005)025[0929:ROSMKC]2.0.CO;2

Kazár E, Hampe O. 2014. A new species of *Kentriodon* (Mammalia, Odontoceti, Delphinoidea) from the middle/late Miocene of Groß Pampau (Schleswig-Holstein, North Germany). *Journal of Vertebrate Paleontology. 34*(5):1216–1230. https://doi.org/10.1080/02724634.2014.857347

Kellogg R. 1922. Description of the skull of *Megaptera miocaena*, a fossil humpback whale from the Miocene diatomaceous earth of Lompoc, California. *Proceedings of the United States National Museum. 61*(14):1–18. https://doi.org/10.5479/si.00963801.61-2435.1

Kellogg R. 1923a. Description of an apparently new toothed cetacean from South Carolina. *Smithsonian Miscellaneous Collections. 76*(7):1–7.

Kellogg R. 1923b. Description of two squalodonts recently discovered in the Calvert Cliffs, Maryland; and notes on the shark-toothed cetaceans. *Proceedings of the United States National Museum. 62*(16):1–69. https://doi.org/10.5479/si.00963801.62-2462.1

Kellogg R. 1924. Description of a new genus and species of whalebone whale from the Calvert Cliffs, Maryland. *Proceedings of the United States National Museum. 63*:1–14. https://doi.org/10.5479/si.00963801.63-2483.1

Kellogg R. 1925a. Additions to the Tertiary History of the Pelagic Mammals on the Pacific Coast of North America. *Carnegie Institution of Washington Publication. 348*:1–120. https://doi.org/10.5962/t.165691

Kellogg R. 1925b. On the occurrence of remains of fossil porpoises of the genus *Eurhinodelphis* in North America. *Proceedings of the United States National Museum. 66*:1–40. https://doi.org/10.5479/si.00963801.66-2563.1

Kellogg R. 1927a. *Kentriodon pernix*, a Miocene porpoise from Maryland. *Proceedings of the United States National Museum. 69*(19):1–55. https://doi.org/10.5479/si.00963801.69-2645.1

Kellogg R. 1927b. Study of the skull of a fossil sperm-whale from the Temblor Miocene of southern California. *Carnegie Institution of Washington Publication. 346*:1–24.

Kellogg R. 1929a. A new cetothere from southern California. *University of California Publications in Geological Sciences. 18*(15):449–457.

Kellogg R. 1929b. A new fossil toothed whale from Florida. *American Museum Novitates. 389*:1–10.

Kellogg R. 1931. Pelagic mammals from the Temblor Formation of the Kern River region, California. *Proceedings of the California Academy of Sciences. 19*:217–397.

Kellogg R. 1932. A Miocene long-beaked porpoise from California. *Smithsonian Miscellaneous Collections. 87*(2):1–11.

Kellogg R. 1934. A new cetothere from the Modelo Formation at Los Angeles, California. *Carnegie Institution of Washington Publication. 447*:83–104.

Kellogg R. 1941. On the cetotheres figured by Vandelli. *Boletim do Museu de Mineralogia e Geologia da Universidade de Lisboa. 7*(8):13–22.

Kellogg R. 1955. Three Miocene porpoises from the Calvert Cliffs, Maryland. *Proceedings of the United States National Museum. 105*(3354):101–154. https://doi.org/10.5479/si.00963801.105-3354.101

Kellogg R. 1957. Two additional Miocene porpoises from the Calvert Cliffs, Maryland. *Proceedings of the United States National Museum. 107*:279–337. https://doi.org/10.5479/si.00963801.108-3387.279

Kellogg R. 1965. A new whalebone whale from the Miocene Calvert formation. *United States National Museum Bulletin. 247*:3–43.

Kellogg R. 1966. Fossil marine mammals from the Miocene Calvert formation of Maryland and Virginia. 4. A new odontocete from the Calvert Miocene of Maryland. *United States National Museum Bulletin. 247*:99–101.

Kellogg R. 1968. Fossil marine mammals from the Miocene Calvert Formation of Maryland and Virginia, Parts 5–8. *Bulletin of the United States National Museum. 247*:103–201.

Kellogg R. 1969. Cetothere skeletons from the Miocene Choptank formation of Maryland and Virginia. *United States National Museum Bulletin. 294*:1–40.

Khramov AV, Foraponova T, Węgierek P. 2023. The earliest pollen-loaded insects from the Lower Permian of Russia. *Biology Letters. 19*(3):20220523. https://doi.org/10.1098/rsbl.2022.0523

Kidwell SM, Powars DS, Edwards LE, Vogt PR. 2015. Miocene stratigraphy and paleoenvironments of the Calvert Cliffs, Maryland. In: *Tripping from the Fall Line: Field Excursions for the GSA Annual Meeting, Baltimore, 2015*. Vol. 40. Geological Society of America. p. 231–279. https://doi.org/10.1130/2015.0040(08)

Kimura T, Barnes LG. 2016. New Miocene fossil Allodelphinidae (Cetacea, Odontoceti, Platanistoidea) from the North Pacific Ocean. *Bulletin of the Gunma Museum of Natural History. 20*:1–58.

Kimura T, Hasegawa Y. 2010. A new baleen whale (Mysticeti: Cetotheriidae) from the earliest late Miocene of Japan and a reconsideration of the phylogeny of cetotheres. *Journal of Vertebrate Paleontology. 30*(2):577–591. https://doi.org/10.1080/02724631003621912

Kimura T, Hasegawa Y. 2019. A new species of *Kentriodon* (Cetacea, Odontoceti, Kentriodontidae) from the Miocene of Japan. *Journal of Vertebrate Paleontology. 39*(1):e1566739. https://doi.org/10.1080/02724634.2019.1566739

Kimura T, Hasegawa Y. 2020. *Norisdelphis annakaensis*, a new Miocene delphinid from Japan. *Journal of Vertebrate Paleontology. 40*(1):e1762628. https://doi.org/10.1080/02724634.2020.1762628

Kimura T, Hasegawa Y. 2022. A new physeteroid from the lower Miocene of Japan. *Paleontological Research. 26*(1):87–101. https://doi.org/10.2517/PR200021

Kimura T, Hasegawa Y. 2024a. A new species of Late Miocene balaenopterid, *Incakujira fordycei*, from Sacaco, Peru. *Bulletin of the Gunma Museum of Natural History*. (28):1–14.

Kimura T, Hasegawa Y. 2024b. New fossil lipotid (Cetacea, Delphinida) from the upper Miocene of Japan. *Paleontological Research. 28*(4):361–383. https://doi.org/10.2517/PR220027

Kimura T, Hasegawa Y, Kohno N. 2018. A new species of the genus *Eschrichtius* (Cetacea: Mysticeti) from the early Pleistocene of Japan. *Paleontological Research. 22*(1):1–19. https://doi.org/10.2517/2017PR007

Kimura T, Hasegawa Y, Suzuki T. 2022. A new species of baleen whale (*Isanacetus*-group) from the early Miocene, Japan. *Paleontological Research. 27*(1):85–101. https://doi.org/10.2517/PR210009

Kimura T, Narita K, Fujita T, Hasegawa Y. 2007. A new species of *Eubalaena* (Cetacea: Mysticeti: Balaenidae) from the Gonda Formation (latest Miocene–early Pliocene) of Japan. *Bulletin of the Gunma Museum of Natural History. 11*:15–27.

Kimura T, Ozawa T. 2002. A new cetothere (Cetacea: Mysticeti) from the early Miocene of Japan. *Journal of Vertebrate Paleontology. 22*(3):684–702. https://doi.org/10.1671/0272-4634(2002)022[0684:ANCCMF]2.0.CO;2

Kirpichnikov AA. 1954. Dva novykh roda del'finov iz sarmata SSSR. *Trudy Paleontologicheskogo Instituta, Akademiya Nauk SSSR. 47*:181–190.

Kliver M. 1883. Ueber einige neue Blattinarien-, zwei Dictyoneura- und zwei Arthropleura-Arten aus der Saarbrücker Steinkohlenformation. *Palaeontographica. 29*:249–280.

Kliver M. 1886. Ueber einige neue Arthropodenreste aus dem Saarbrücker und der Wettin-Löbejüner Steinkohlenformation. *Palaeontographica. 32*:99–115.

Korth WW. 2008. Cranial morphology, systematics and succession of beavers from the middle Miocene Valentine Formation of Nebraska, USA. *Acta Palaeontologica Polonica. 53*(2):169–182. https://doi.org/10.4202/app.2008.0201

Krainer K, Lucas SG, Vachard D. 2011. Lectostratotype section of the Pennsylvanian Sandia Formation, Bernalillo County, New Mexico. *Bulletin of the New Mexico Museum of Natural History and Science. 53*:71–85.

Kudrin LN, Tatarinov KA. 1966. O miotsenovykh del'finakh Zapadnoy Ukrainy. *Paleontologicheskii Zhurnal.* (4):68–74.

Kukalová J. 1955. *Permoedischia* n. g. (Protorthoptera) a *Moraviptera* n. g. (Palaeodictyoptera) z moravského permu. *Sborník Ústředního ústavu geologického. 21*:541–575.

Kukalová J. 1958. On Czechoslovakian Spilapteridae Handlirsch (Insecta–Palaeodictyoptera). *Acta Universitatis Carolinae, Geologica.* (3):231–240.

Kukalová J. 1959. *Breyeria barborae* n. sp. (Insecta, Palaeodictyoptera) of Upper Silesian coal basin (Westphalian). *Věstník Ústředního ústavu geologického. 34*:310–313.

Kukalová J. 1960. New Palaeodictyoptera of the Carboniferous and Permian of Czechoslovakia. *Sborník Ústředního ústavu geologického. 25*:239–251.

Kukalová J. 1964. Review of the Calvertiellidae, with description of a new genus from Permian strata of Moravia (Palaeodictyoptera). *Psyche: A Journal of Entomology. 71*(4):153–168. https://doi.org/10.1155/1964/34192

Kukalová J. 1969. Revisional study of the order Palaeodictyoptera in the Upper Carboniferous shales of Commentry, France. Part I. *Psyche: A Journal of Entomology. 76*(2):163–215. https://doi.org/10.1155/1969/74019

Kukalová J. 1970. Revisional study of the order Palaeodictyoptera in the Upper Carboniferous shales of Commentry, France. Part III. *Psyche: A Journal of Entomology.* *77*(1):1–44. https://doi.org/10.1155/1970/71691

Kukalová-Peck J. 1972. Unusual structures in the Paleozoic insect orders Megasecoptera and Palaeodictyoptera, with a description of a new family. *Psyche: A Journal of Entomology.* *79*(3):243–268. https://doi.org/10.1155/1972/98019

Kukalová-Peck J. 1974. Wing-folding in the Paleozoic insect order Diaphanopterodea (Paleoptera), with a description of new representatives of the family Elmoidae. *Psyche: A Journal of Entomology.* *81*(2):315–333. https://doi.org/10.1155/1974/49489

Kukalová-Peck J. 1975. Megasecoptera from the Lower Permian of Moravia. *Psyche: A Journal of Entomology.* *82*(1):1–19. https://doi.org/10.1155/1975/78215

Kukalová-Peck J, Brauckmann C. 1990. Wing folding in pterygote insects, and the oldest Diaphanopterodea from the early Late Carboniferous of West Germany. *Canadian Journal of Zoology.* *68*:1104–1111. https://doi.org/10.1139/z90-163

Kukalová-Peck J, Peck SB. 1976. Adult and immature Calvertiellidae (Insecta: Palaeodictyoptera) from the Upper Paleozoic of New Mexico and Czechoslovakia. *Psyche: A Journal of Entomology.* *83*:79–93. https://doi.org/10.1155/1976/36962

Kukalová-Peck J, Richardson ES. 1983. New Homoiopteridae (Insecta: Paleodictyoptera) with wing articulation from Upper Carboniferous strata of Mazon Creek, Illinois. *Canadian Journal of Zoology.* *61*(7):1670–1687. https://doi.org/10.1139/z83-218

Kukalová-Peck J, Sinichenkova ND. 1992. The wing venation and systematics of Lower Permian Diaphanopterodea from the Ural Mountains, Russia (Insecta: Paleoptera). *Canadian Journal of Zoology.* *70*(2):229–235. https://doi.org/10.1139/z92-036

Lambert O. 2004. Systematic revision of the Miocene long-snouted dolphin *Eurhinodelphis longirostris* Du Bus, 1872 (Cetacea, Odontoceti, Eurhinodelphinidae). *Bulletin de l'Institut royal des Sciences naturelles de Belgique, Sciences de la Terre.* *74*:147–174.

Lambert O. 2005a. Phylogenetic affinities of the long-snouted dolphin *Eurhinodelphis* (Cetacea, Odontoceti) from the Miocene of Antwerp, Belgium. *Palaeontology.* *48*(3):653–679. https://doi.org/10.1111/j.1475-4983.2005.00472.x

Lambert O. 2005b. Review of the Miocene long-snouted dolphin *Priscodelphinus cristatus* Du Bus, 1872 (Cetacea, Odontoceti) and phylogeny among eurhinodelphinids. *Bulletin de l'Institut royal des Sciences naturelles de Belgique, Sciences de la Terre.* *75*:211–235.

Lambert O. 2005c. Systematics and phylogeny of the fossil beaked whales *Ziphirostrum* du Bus, 1868 and *Choneziphius* Duvernoy, 1851 (Mammalia, Cetacea, Odontoceti), from the Neogene of Antwerp (north of Belgium). *Geodiversitas*. *27*(3):443–497.

Lambert O. 2008a. A new porpoise (Cetacea, Odontoceti, Phocoenidae) from the Pliocene of the North Sea. *Journal of Vertebrate Paleontology*. *28*(3):863–872. https://doi.org/10.1671/0272-4634(2008)28[863:ANPCOP]2.0.CO;2

Lambert O. 2008b. Sperm whales from the Miocene of the North Sea: A re-appraisal. *Bulletin de l'Institut royal des Sciences naturelles de Belgique, Sciences de la Terre*. *78*:277–316.

Lambert O, Auclair C, Cauxeiro C, Lopez M, Adnet S. 2018a. A close relative of the Amazon river dolphin in marine deposits: A new Iniidae from the late Miocene of Angola. *PeerJ*. *6*:e5556. https://doi.org/10.7717/peerj.5556

Lambert O, Bianucci G, de Muizon C. 2008. A new stem-sperm whale (Cetacea, Odontoceti, Physeteroidea) from the latest Miocene of Peru. *Comptes Rendus Palevol*. *7*(6):361–369. https://doi.org/10.1016/j.crpv.2008.06.002

Lambert O, Bianucci G, de Muizon C. 2017a. Macroraptorial sperm whales (Cetacea, Odontoceti, Physeteroidea) from the Miocene of Peru. *Zoological Journal of the Linnean Society*. *179*(2):404–474. https://doi.org/10.1111/zoj.12456

Lambert O, Bianucci G, Post K. 2009. A new beaked whale (Odontoceti, Ziphiidae) from the middle Miocene of Peru. *Journal of Vertebrate Paleontology*. *29*(3):910–922. https://doi.org/10.1671/039.029.0304

Lambert O, Bianucci G, Post K, de Muizon C, Salas-Gismondi R, Urbina M, Reumer J. 2010. The giant bite of a new raptorial sperm whale from the Miocene epoch of Peru. *Nature*. *466*(7302):105–108. https://doi.org/10.1038/nature09067

Lambert O, Bianucci G, Salas-Gismondi R, Di Celma C, Steurbaut E, Urbina M, de Muizon C. 2019a. An amphibious whale from the middle Eocene of Peru reveals early South Pacific dispersal of quadrupedal cetaceans. *Current Biology*. *29*(8):1352–1359. https://doi.org/10.1016/j.cub.2019.02.050

Lambert O, Bianucci G, Urbina M. 2014. *Huaridelphis raimondii*, a new early Miocene Squalodelphinidae (Cetacea, Odontoceti) from the Chilcatay formation, Peru. *Journal of Vertebrate Paleontology*. *34*(5):987–1004. https://doi.org/10.1080/02724634.2014.858050

Lambert O, Bianucci G, Urbina M, Geisler JH. 2017b. A new inioid (Cetacea, Odontoceti, Delphinida) from the Miocene of Peru and the origin of modern dolphin and porpoise families. *Zoological Journal of the Linnean Society*. *179*(4):919–946. https://doi.org/10.1111/zoj.12479

Lambert O, Collareta A, Benites-Palomino AM, de Muizon C, Urbina M, Bianucci G. 2021. A new small, mesorostrine inioid (Cetacea, Odontoceti, Delphinida) from four upper Miocene localities in the Pisco basin, Peru. *Papers in Palaeontology*. *7*(2):1043–1064. https://doi.org/10.1002/spp2.1332

Lambert O, Collareta A, Benites-Palomino AM, Merella M, de Muizon C, Bennion R, Urbina M, Bianucci G. 2023. A new platyrostrine sperm whale from the early Miocene of the southeastern Pacific (East Pisco basin, Peru) supports affinities with the southwestern Atlantic cetacean fauna. *Geodiversitas*. *45*(22):659–679. https://doi.org/10.5252/geodiversitas2023v45a22

Lambert O, Collareta A, Landini W, Post K, Ramassamy B, Di Celma C, Urbina M, Bianucci G. 2015a. No deep diving: Evidence of predation on epipelagic fish for a stem beaked whale from the late Miocene of Peru. *Proceedings of the Royal Society B: Biological Sciences*. *282*(1815):20151530. https://doi.org/10.1098/rspb.2015.1530

Lambert O, de Muizon C. 2013. A new long-snouted species of the Miocene pontoporiid dolphin *Brachydelphis* and a review of the Mio-Pliocene marine mammal levels in the Sacaco Basin, Peru. *Journal of Vertebrate Paleontology*. *33*(3):709–721. https://doi.org/10.1080/02724634.2013.743405

Lambert O, de Muizon C, Bennion RF, Urbina M, Bianucci G. 2025. New data on archaic homodont odontocetes from the early Miocene of Peru reveal a second species of *Chilcacetus* Lambert, Muizon & Bianucci, 2015 and a southern hemisphere record for a northeastern Pacific species. *Geodiversitas*. *47*(9):369–408. https://doi.org/10.5252/geodiversitas2025v47a9

Lambert O, de Muizon C, Bianucci G. 2013. The most basal beaked whale *Ninoziphius platyrostris* Muizon, 1983: Clues on the evolutionary history of the family Ziphiidae. *Zoological Journal of the Linnean Society*. *167*(4):569–598. https://doi.org/10.1111/zoj.12018

Lambert O, de Muizon C, Bianucci G. 2015b. A new archaic homodont toothed cetacean (Mammalia, Cetacea, Odontoceti) from the early Miocene of Peru. *Geodiversitas*. *37*(1):79–108. https://doi.org/10.5252/g2015n1a4

Lambert O, de Muizon C, Malinverno E, Di Celma C, Urbina M, Bianucci G. 2018b. A new odontocete from the early Miocene of Peru expands the morphological disparity of extinct heterodont dolphins. *Journal of Systematic Palaeontology*. *16*(12):981–1016. https://doi.org/10.1080/14772019.2017.1359689

Lambert O, de Muizon C, Urbina M, Bianucci G. 2020. A new longirostrine sperm whale (Cetacea, Physeteroidea) from the lower Miocene of the Pisco basin (southern coast of Peru). *Journal of Systematic Palaeontology*. *18*(20):1707–1742. https://doi.org/10.1080/14772019.2020.1805520

Lambert O, Estevens M, Smith R. 2005. A new kentriodontine dolphin from the middle Miocene of Portugal. *Acta Palaeontologica Polonica*. *50*(2):239–248.

Lambert O, Godfrey SJ, Fitzgerald EMG. 2019b. *Yaquinacetus meadi*, a new latest Oligocene–early Miocene dolphin (Cetacea, Odontoceti, Squaloziphiidae, fam. nov.) from the Nye mudstone (Oregon, U.S.A.) *Journal of Vertebrate Paleontology*. *38*(6):e1559174. https://doi.org/10.1080/02724634.2018.1559174

Lambert O, Louwye S. 2006. *Archaeoziphius microglenoideus*, a new primitive beaked whale (Mammalia, Cetacea, Odontoceti) from the Middle Miocene of Belgium. *Journal of Vertebrate Paleontology*. *26*(1):182–191. https://doi.org/10.1671/0272-4634(2006)26[182:AMANPB]2.0.CO;2

Lambert O, Louwye S. 2016. A new early Pliocene species of *Mesoplodon*: A calibration mark for the radiation of this species-rich beaked whale genus. *Journal of Vertebrate Paleontology*. *36*(2):e1055754. https://doi.org/10.1080/02724634.2015.1055754

Lambert O, Martínez-Cáceres M, Bianucci G, Di Celma C, Salas-Gismondi R, Steurbaut E, Urbina M, de Muizon C. 2017c. Earliest mysticete from the Late Eocene of Peru sheds new light on the origin of baleen whales. *Current Biology*. *27*(10):1535–1541. https://doi.org/10.1016/j.cub.2017.04.026

Lambert O, Oberlin M, de Muizon C, Collareta A, Urbina M, Bianucci G. 2026. A new kentriodontid dolphin (Cetacea, Odontoceti, Delphinida) from the middle Miocene of the southeastern Pacific. *Geodiversitas*. *48*(3):25–50. https://doi.org/10.5252/geodiversitas2026v48a3

Lambert O, Post K. 2005. First European pontoporiid dolphins (Mammalia: Cetacea, Odontoceti) from the Miocene of Belgium and the Netherlands. *Deinsea*. *11*:7–20.

Lameere A. 1917. Révision sommaire des insectes fossiles du Stéphanien de Commentry. *Bulletin du Muséum national d'histoire naturelle*. *23*:141–200.

Langiaux J, Parriat H. 1975. Entomofaune du bassin de Blanzy-Montceaux. nouvelles acquisitions. *La Physiophile*. *83*:35–45.

Laurentiaux D. 1949a. Description de trois nouveaux Paléodictyoptères du terrain houiller du Nord de la France. *Annales de la Société géologique du Nord*. *69*:206–223. https://doi.org/10.3406/asgn.1949.5681

Laurentiaux D. 1949b. Un insecte nouveau (Breyeriidæ) du terrain houiller du Pas-de-Calais. *Annales de la Société géologique du Nord*. *69*:47–54. https://doi.org/10.3406/asgn.1949.5671

Laurentiaux D. 1950. Les insectes carbonifères du Limbourg hollandais. *Mededelingen van de Geologische Stichting (Nieuwe Serie)*. *4*:13–22.

Laurentiaux D. 1958. *Patteiskya bouckaerti* nov. gen. et sp., insekt aus dem namur des ruhrkarbons. *Neues Jahrbuch für Geologie und Paläontologie, Monatshefte*. *1958*(7):302–306.

Laurentiaux D, Teixeira C. 1958. Um novo género de insecto paleodictióptero do Carbónico continental do Baixo-Douro (Portugal). *Boletim da Sociedade Geológica de Portugal*. *12*:41–49.

Laurentiaux-Vieira F, Laurentiaux D. 1964. Diagramme ptéronologique du genre westphalien *Breyeria* De Borre (Insectes Paléodictyoptères). *Comptes rendus hebdomadaires des séances de l'Académie des sciences. 258*:1282–1284.

Laurentiaux-Vieira F, Laurentiaux D. 1986. Paléodictyoptère nouveau du Namurien belge. *Annales de la Société géologique du Nord. 105*(3):187–193. https://doi.org/10.3406/asgn.1985.1549

Lawley R. 1876. *Nuovi Studi Sopra ai Pesci ed Altri Vertebrati Fossili delle Colline Toscane*. Firenze: Tipografia dell'Arte della Stampa.

Leidy J. 1869. The extinct mammalian fauna of Dakota and Nebraska, including an account of some allied forms from other localities, together with a synopsis of the mammalian remains of North America. *Journal of the Academy of Natural Sciences of Philadelphia, 2nd Series. 7*:23–472.

Leidy J. 1877. Description of vertebrate remains, chiefly from the phosphate beds of South Carolina. *Journal of the Academy of Natural Sciences of Philadelphia. 8*:209–261.

Leriche M. 1911. Un insecte nouveau du houiller belge (*Stenodictyoneura belgica*). *Annales de la Société géologique de Belgique. 38*:193–195.

Leslie MS, Peredo CM, Pyenson ND. 2019. *Norrisanima miocaena*, a new generic name and redescription of a stem balaenopteroid mysticete (Mammalia, Cetacea) from the Miocene of California. *PeerJ. 7*:e7629. https://doi.org/10.7717/peerj.7629

Li Y, Ren D, Pecharová M, Prokop J. 2013. A new palaeodictyopterid (Insecta: Palaeodictyoptera: Spilapteridae) from the Upper Carboniferous of China supports a close relationship between insect faunas of Quilianshian (northern China) and Laurussia. *Alcheringa: An Australasian Journal of Palaeontology. 37*(4):487–495. https://doi.org/10.1080/03115518.2013.793024

Liu X, Béthoux O, Yin X, Ren D. 2015. The smallest Palaeodictyoptera (Insecta) discovered at Xiaheyan (Late Carboniferous, China). *Comptes Rendus Palevol. 14*(5):346–352. https://doi.org/10.1016/j.crpv.2015.05.013

Longhi P. 1897. Della pietra da coti o da mola bellunese e di alcuni suoi fossili (*Schizodelphis squalodontoides capellinii* n. sp. ecc.) *Atti dell'Accademia Scientifica Veneto-Trentino-Istriana. 3*(2):41–86.

Longhi P. 1898. Sopra i resti di un cranio di *Champsodelphis* fossile. *Atti della Società Veneto-Trentina di Scienze Naturali. 3*:323–381.

Lourenço JM, Claude J, Galtier N, Chiari Y. 2012. Dating cryptodiran nodes: Origin and diversification of the turtle superfamily Testudinoidea. *Molecular Phylogenetics and Evolution. 62*(1):496–507. https://doi.org/10.1016/j.ympev.2011.10.022

Louwye S. 2002. Dinoflagellate cyst biostratigraphy of the Upper Miocene Deurne Sands (Diest Formation) of northern Belgium, southern North Sea Basin. *Geological Journal. 37*(1):55–67. https://doi.org/10.1002/gj.900

Louwye S, De Coninck J, Verniers J. 2000. Shallow marine Lower and Middle Miocene deposits at the southern margin of the North Sea Basin (northern Belgium): Dinoflagellate cyst biostratigraphy and depositional history. *Geological Magazine*. *137*(4):381–394. https://doi.org/10.1017/S0016756800004258

Louwye S, Deckers J, Verhaegen J, Adriaens R, Vandenberghe N. 2020. A review of the lower and middle Miocene of northern Belgium. *Geologica Belgica*. *23*(3-4):137–156. https://doi.org/10.20341/gb.2020.010

Louwye S, Head MJ, De Schepper S. 2004. Dinoflagellate cyst stratigraphy and palaeoecology of the Pliocene in northern Belgium, southern North Sea Basin. *Geological Magazine*. *141*(3):353–378. https://doi.org/10.1017/S0016756804009136

Lucas SG, Shen SZ. 2018. The Permian timescale: An introduction. *Geological Society, London, Special Publications*. *450*(1):1–19. https://doi.org/10.1144/sp450.15

Lull RS. 1914. Fossil dolphin from California. *American Journal of Science, Series 4*. *37*:209–220.

Lydekker R. 1892. On zeuglodont and other cetacean remains from the Tertiary of the Caucasus. *Proceedings of the Geological Society of London*. *1892*:558–564.

Lydekker R. 1894. Contributions to a knowledge of the fossil vertebrates of Argentina. II. Cetacean skulls from Patagonia. *Anales del Museo de La Plata, Paleontología Argentina*. *2*:1–13.

Makiyama J. 1936. *Sinanodelphis izumidaensis*, a new Miocene dolphin of Japan. *Memoirs of the College of Science, Kyoto Imperial University, Series B*. *11*(2):115–134.

Mantell GA. 1844. *The Medals of Creation; or, First Lessons in Geology, and the Study of Organic Remains*. London: Henry G. Bohn.

Marples BJ. 1956. Cetotheres (Cetacea) from the Oligocene of New Zealand. *Proceedings of the Zoological Society of London*. *126*(4):565–580.

Martin JE, Mallory VS. 2011. Vertebrate paleontology of the late Miocene (Hemphillian) Wilbur Locality of central Washington. *Paludicola*. *8*(3):155–185.

Martínez-Cáceres M, de Muizon C. 2011. A new basilosaurid (Cetacea, Pelagiceti) from the Late Eocene to Early Oligocene Otuma Formation of Peru. *Comptes Rendus Palevol*. *10*(7):517–526. https://doi.org/10.1016/j.crpv.2011.03.006

Martínez-Cáceres M, Lambert O, Muizon Cd. 2017. The anatomy and phylogenetic affinities of *Cynthiacetus peruvianus*, a large *Dorudon*-like basilosaurid (Cetacea, Mammalia) from the late Eocene of Peru. *Geodiversitas*. *39*(1):7–163. https://doi.org/10.5252/g2017n1a1

Martynov AV. 1930a. O novykh permskikh nasekomykh Tikhikh Gor. I. Palaeoptera. *Trudy Geologicheskogo Muzeya Akademii Nauk SSSR*. *6*:69–86.

Martynov AV. 1930b. Paleozoiskie nasekomye Kuznetskogo basseina. *Izvestiya Glavnogo Geologo-Razvedochnogo Upravleniya*. *49*(10):73–111.

Martynov AV. 1932. New Permian Palaeoptera with the discussion of some problems of their evolution. *Trudy Paleozoologicheskogo Instituta*. *1*:1–44.

Martynov AV. 1937. Permskie iskopaemye nasekomye Kargaly i ikh otnosheniya. *Trudy Paleontologicheskogo Instituta*. *7*(2):1–92.

Martynov AV. 1940. Permskie iskopaemye nasekomye Chekardy. *Trudy Paleontologicheskogo Instituta Akademii Nauk SSSR*. *11*:1–63.

Marx FG. 2011. Lost in translation — a history of systematic confusion and comments on the type species of *Squalodon* and *Patriocetus* (Cetacea, Odontoceti). *Palaeontology*. *54*(2):303–307. https://doi.org/10.1111/j.1475-4983.2010.01018.x

Marx FG, Bosselaers M, Louwye S. 2016. A new species of *Metopocetus* (Cetacea, Mysticeti, Cetotheriidae) from the Late Miocene of the Netherlands. *PeerJ*. *4*:e1572. https://doi.org/10.7717/peerj.1572

Marx FG, Coste A, Richards MD, Palin JM, Fordyce RE. 2024. Strontium isotopes reveal a globally unique assemblage of early Miocene baleen whales. *Journal of the Royal Society of New Zealand*. *54*(5):711–721. https://doi.org/10.1080/03036758.2023.2278732

Marx FG, Fordyce RE. 2015. Baleen boom and bust: A synthesis of mysticete phylogeny, diversity and disparity. *Royal Society Open Science*. *2*(4):140434. https://doi.org/10.1098/rsos.140434

Marx FG, Kohno N. 2016. A new Miocene baleen whale from the Peruvian desert. *Royal Society Open Science*. *3*(10):160542. https://doi.org/10.1098/rsos.160542

Marx FG, Lambert O, de Muizon C. 2017. A new Miocene baleen whale from Peru deciphers the dawn of cetotheriids. *Royal Society Open Science*. *4*(9):170560. https://doi.org/10.1098/rsos.170560

Marx FG, Post K, Bosselaers M, Munsterman DK. 2019. A large Late Miocene cetotheriid (Cetacea, Mysticeti) from the Netherlands clarifies the status of Tranatocetidae. *PeerJ*. *7*:e6426. https://doi.org/10.7717/peerj.6426

Marx FG, Tsai CH, Fordyce RE. 2015. A new early Oligocene toothed 'baleen' whale (Mysticeti: Aetiocetidae) from western North America: One of the oldest and the smallest. *Royal Society Open Science*. *2*(12):150476. https://doi.org/10.1098/rsos.150476

Matsumoto H. 1926. On some fossil cetaceans of Japan. *Science Reports of the Tohoku Imperial University, Series 2 (Geology)*. *10*:23–25.

Matsumoto H. 1937. A new species of orc from the basal Calabrian at Naganuma, Minato town, province of Kazusa, Japan. *Zoological Magazine (Tokyo)*. *49*(5):191–193.

Matthew GF. 1910. Remarkable forms of the Little River Group. *Proceedings and Transactions of the Royal Society of Canada, Third Series*. *3*(4):115–133.

McGowen MR, Tsagkogeorga G, Álvarez-Carretero S, dos Reis M, Struebig M, Deaville R, Jepson PD, Jarman S, Polanowski A, Morin PA, et al. 2020. Phylogenomic resolution of the cetacean tree of life using target sequence capture. *Systematic Biology*. *69*(3):479–501. https://doi.org/10.1093/sysbio/syz068

Mchedlidze GA. 1959. Iskopaemyi del'fin *Imerodelphis thabagarii* iz sarmata okrestnostei Zestafoni. *Soobshcheniya Akademii Nauk Gruzinskoi SSR*. *23*(6):687–694.

Mchedlidze GA. 1964. *Iskopaemye Kitoobraznye Kavkaza*. Tbilisi: Metsniereba.

Mchedlidze GA. 1970. *Nekotorye Obshchie Cherty Istorii Kitoobraznykh, Chast' I*. Tbilisi: Metsniereba.

Mchedlidze GA. 1976. *Osnovnye Cherty Paleobiologicheskoi Istorii Kitoobraznykh*. Tbilisi: Metsniereba.

Mchedlidze GA. 1984. *Iskopaemyi Kit iz Miotsenovykh Otlozhenii Okrestnostei Stanitsy Otradnaya (Severnyi Kavkaz)*. Tbilisi: Metsniereba.

McLeod SA, Barnes LG. 2008. A new genus and species of Eocene protocetid archaeocete whale (Mammalia, Cetacea) from the Atlantic coastal plain. *Science Series, Natural History Museum of Los Angeles County*. *41*:73–98.

Meekin S, Fordyce RE, Coste A. 2024. *Aureia rerehua*, a new platanistoid dolphin from the Oligocene of New Zealand with a unique feeding method. *Journal of the Royal Society of New Zealand*. *54*(5):758–777. https://doi.org/10.1080/03036758.2024.2314505

Meredith RW, Janečka JE, Gatesy J, Ryder OA, Fisher CA, Teeling EC, Goodbla A, Eizirik E, Simão TLL, Stadler T, et al. 2011. Impacts of the Cretaceous terrestrial revolution and KPg extinction on mammal diversification. *Science*. *334*(6055):521–524. https://doi.org/10.1126/science.1211028

Meunier F. 1908a. Nouveaux Mégasécoptérides et nouveau Paléodictyoptère de Commentry. *Bulletin du Muséum national d'histoire naturelle*. *14*(3):172–175.

Meunier F. 1908b. Nouveaux Paléodictyoptères du Stéphanien de Commentry. *Bulletin du Muséum national d'histoire naturelle*. *14*(1):34–37.

Meunier F. 1908c. Quatrième note sur de nouveaux insectes du Stéphanien de Commentry. *Bulletin du Muséum national d'histoire naturelle*. *14*(5):244–249.

Meunier F. 1908d. Sur quelques Sténodictyoptères du terrain houiller de Commentry. *Annales de la Société scientifique de Bruxelles*. *32*:154–155.

Meunier F. 1908e. Un Platyptéride et un Sténodictyoptéride du Stéphanien de Commentry. *Annales de la Société scientifique de Bruxelles*. *32*:153–154.

Meunier F. 1909. Nouveaux Paléodictyoptères et Protorthoptères du Stéphanien de Commentry. *Annales de la Société scientifique de Bruxelles*. *33*:139–140.

Meunier F. 1910. Nouveaux Paléodictyoptères du houiller de Commentry. *Bulletin du Muséum national d'histoire naturelle*. *16*:233–237. https://doi.org/10.5962/bhl.part.20426

Meunier F. 1911. Nouveaux insectes du houiller de Commentry. *Bulletin du Muséum national d'histoire naturelle. 17*:117–128.

Meunier F. 1914. Un nouveau Paléodictyoptère (Sténodictyoptère) du houiller de Commentry (Allier). *Bulletin de la Société entomologique de France. 19*(3):119–122. https://doi.org/10.3406/bsef.1914.25505

Michel LA, Tabor NJ, Montañez IP, Schmitz MD, Davydov VI. 2015. Chronostratigraphy and paleoclimatology of the Lodève Basin, France: Evidence for a pan-tropical aridification event across the Carboniferous–Permian boundary. *Palaeogeography, Palaeoclimatology, Palaeoecology. 430*:118–131. https://doi.org/10.1016/j.palaeo.2015.03.020

Miján I, Louwye S, Lambert O. 2017. A new *Beneziphius* beaked whale from the ocean floor off Galicia, Spain and biostratigraphic reassessment of the type species. *Acta Palaeontologica Polonica. 62*(1):211–220. https://doi.org/10.4202/app.00309.2016

Misof B, Liu S, Meusemann K, Peters RS, Donath A, Mayer C, Frandsen PB, Ware J, Flouri T, Beutel RG, et al. 2014. Phylogenomics resolves the timing and pattern of insect evolution. *Science. 346*(6210):763–767. https://doi.org/10.1126/science.1257570

Missimer TM. 1999. Sequence stratigraphy of the late Miocene–early Pliocene Peace River formation, southwestern Florida. *Transactions of the Gulf Coast Association of Geological Societies. 49*:358–368.

Mitchell ED. 1989. A new cetacean from the late Eocene La Meseta Formation, Seymour Island, Antarctic Peninsula. *Canadian Journal of Fisheries and Aquatic Sciences. 46*(12):2219–2235. https://doi.org/10.1139/f89-273

Młynarski M. 1956. On a new species of emydid-tortoise from the Pliocene of Poland. *Acta Palaeontologica Polonica. 1*:153–164.

Moreno FP. 1892. Lijeros apuntes sobre dos géneros de cetáceos fósiles de la república Argentina. *Revista del Museo de La Plata. 3*:393–400.

Morgan GS. 1994. Miocene and Pliocene marine mammal faunas from the Bone Valley formation of central Florida. In: Berta A, Deméré TA, editors. *Contributions in Marine Mammal Paleontology Honoring Frank C. Whitmore, Jr.* Vol. 29. p. 239–268.

Muizon Cd. 1991. A new Ziphiidae (Cetacea) from the early Miocene of Washington State (USA) and phylogenetic analysis of the major groups of odontocetes. *Bulletin du Muséum national d'Histoire naturelle, 4e série, Section C. 12*(3–4):279–326.

Müller J. 1849. *Über die fossilen Reste der Zeuglodonten von Nordamerica, mit Rücksicht auf die europäischen Reste aus dieser Familie*. Berlin: G. Reimer.

Murakami M. 2016. A new extinct inioid (Cetacea, Odontoceti) from the upper Miocene Senhata Formation, Chiba, central Japan: The first record of Inioidea from the North Pacific Ocean. *Paleontological Research*. *20*(3):207–225. https://doi.org/10.2517/2015PR031

Murakami M, Shimada C, Hikida Y, Hirano H. 2012a. A new basal porpoise, *Pterophocaena nishinoi* (Cetacea, Odontoceti, Delphinoidea), from the upper Miocene of Japan and its phylogenetic relationships. *Journal of Vertebrate Paleontology*. *32*(5):1157–1171. https://doi.org/10.1080/02724634.2012.677299

Murakami M, Shimada C, Hikida Y, Hirano H. 2012b. Two new extinct basal phocoenids (Cetacea, Odontoceti, Delphinoidea), from the upper Miocene Koetoi formation of Japan and their phylogenetic significance. *Journal of Vertebrate Paleontology*. *32*(5):1172–1185. https://doi.org/10.1080/02724634.2012.694337

Murakami M, Shimada C, Hikida Y, Soeda Y, Hirano H. 2014. *Eodelphis kabatensis*, a new name for the oldest true dolphin *Stenella kabatensis* Horikawa, 1977 (Cetacea, Odontoceti, Delphinidae), from the upper Miocene of Japan, and the phylogeny and paleobiogeography of Delphinoidea. *Journal of Vertebrate Paleontology*. *34*(3):491–511. https://doi.org/10.1080/02724634.2013.816720

Naugolnykh SV. 2015. Fossil flora from the Kazanian (Middle Permian) Iva-Gora locality, Soyana River, Arkhangelsk Region, Russia. *Paleontological Journal*. *49*(11):1193–1205. https://doi.org/10.1134/S0031030115110076

Near TJ, Meylan PA, Shaffer HB. 2005. Assessing concordance of fossil calibration points in molecular clock studies: An example using turtles. *The American Naturalist*. *165*(2):137–146. https://doi.org/10.1086/427734

Nel A, Lapeyrie J, Garrouste R. 2019. The first European representative of the South American family Xenopteraidae (Insecta: Megasecoptera) discovered in the Guadalupian of Lodève (France). *Alcheringa: An Australasian Journal of Palaeontology*. *43*(3):436–440. https://doi.org/10.1080/03115518.2019.1597162

Nelson CR, Tidwell WD. 1987. *Brodioptera stricklani* n. sp. (Megasecoptera: Brodiopteridae), a new fossil insect from the upper Manning Canyon Shale Formation, Utah (lowermost Namurian B). *Psyche: A Journal of Entomology*. *94*:309–316. https://doi.org/10.1155/1987/52032

Nelson MD, Lambert O, Uhen MD. 2025a. Reassessment of the iconic Oligo-Miocene heterodont dolphin *Squalodon*: A redescription of the type species *S. grateloupii*. *Papers in Palaeontology*. *11*(1):e70003. https://doi.org/10.1002/spp2.70003

Nelson MD, Lambert O, Uhen MD. 2025b. Redescription of the late Oligocene heterodont odontocete *Eosqualodon langewieschei* from the historic Doberg of Bünde, Germany. *Journal of Systematic Palaeontology*. *23*(1):2489940. https://doi.org/10.1080/14772019.2025.2489940

Nelson MD, Lambert O, Uhen MD. 2025c. Taxonomic revision of the family Squalodontidae (Cetacea, Odontoceti): Emptying the wastebasket of fragmentary holotypes. *Papers in Palaeontology. 11*(2):e70002. https://doi.org/10.1002/spp2.70002

Nelson MD, Uhen MD. 2020. A new platanistoid, *Perditicetus yaconensis* gen. et sp. nov. (Cetacea, Odontoceti), from the Chattian–Aquitanian Nye formation of Oregon. *Journal of Systematic Palaeontology. 18*(18):1497–1517. https://doi.org/10.1080/14772019.2020.1783379

Novák OP. 1880. Ueber *Gryllacris Bohemica*, einen neuen Locustidenrest aus der Steinkohlenformation von Stradonitz in Böhmen. *Jahrbuch der Kaiserlich-Königlichen Geologischen Reichsanstalt. 30*:69–74.

Novokshonov VG. 1995. New fossil insects from the Kungurian of the Middle Urals. *Paleontological Journal. 29*(3):76–85.

Novokshonov VG. 1998. New insects (Insecta: Hypoperlida, Mischopterida, Jurinida) from the Lower Permian of the Middle Urals. *Paleontological Journal. 32*:46–53.

Ochoa D, Baby P, DeVries TJ, Barbosa-Espitia A, Foster DA, Altamirano A, de Muizon C, Salas-Gismondi R, Urbina M, Carré M. 2022. Age and provenance of the mio-Pleistocene sediments from the Sacaco area, Peruvian continental margin. *Journal of South American Earth Sciences. 116*:103799. https://doi.org/10.1016/j.jsames.2022.103799

Ochoa D, Salas-Gismondi R, DeVries TJ, Baby P, de Muizon C, Altamirano A, Barbosa-Espitia A, Foster DA, Quispe K, Cardich J, et al. 2021. Late Neogene evolution of the Peruvian margin and its ecosystems: A synthesis from the Sacaco record. *International Journal of Earth Sciences. 110*:995–1025. https://doi.org/10.1007/s00531-021-02003-1

Okazaki Y. 1982. A lower Miocene squalodontid from the Ashiya Group, Kyushu, Japan. *Bulletin of the Kitakyushu Museum of Natural History. 4*:107–112.

Okazaki Y. 2012. A new mysticete from the upper Oligocene Ashiya Group, Kyushu, Japan and its significance to mysticete evolution. *Bulletin of the Kitakyushu Museum of Natural History and Human History, Series A (Natural History). 10*:129–152. https://doi.org/10.34522/kmnh.10.0_129

Opluštil S, Jirásek J, Schmitz MD, Matýsek D. 2017. Biotic changes around the radioisotopically constrained Carboniferous–Permian boundary in the Boskovice Basin (Czech Republic). *Bulletin of Geosciences. 92*(1):95–122. https://doi.org/10.3140/bull.geosci.1638

Opluštil S, Laurin J, Hýlová L, Jirásek J, Schmitz MD, Sivek M. 2022. Coal-bearing fluvial cycles of the late Paleozoic tropics; astronomical control on sediment supply constrained by high-precision radioisotopic ages, Upper Silesian Basin. *Earth-Science Reviews. 228*:103998. https://doi.org/10.1016/j.earscirev.2022.103998

Opluštil S, Schmitz M, Cleal CJ, Martínek K. 2016. A review of the Middle–Late Pennsylvanian west European regional substages and floral biozones, and their correlation to the Geological Time Scale based on new U–Pb ages. *Earth-Science Reviews*. *154*:301–335. https://doi.org/10.1016/j.earscirev.2016.01.004

Otsuka H, Ota Y. 2008. Cetotheres from the early middle Miocene Bihoku Group in Shobara district, Hiroshima prefecture, west Japan. *Miscellaneous Reports of the Hiwa Museum for Natural History*. *49*:1–66.

Owen R. 1839. Observations on the *Basilosaurus* of Dr. Harlan (*Zeuglodon cetoides*, Owen). *Transactions of the Geological Society of London, 2nd series*. *6*(1):69–79. https://doi.org/10.1144/transgslb.6.1.69

Packard EL, Kellogg R. 1934. A New Cetothere from the Miocene Astoria Formation of Newport, Oregon. *Carnegie Institution of Washington Publication*. *447*:1–62.

Palcu DV, Golovina LA, Vernyhorova YV, Popov SV, Krijgsman W. 2017. Middle Miocene paleoenvironmental crises in Central Eurasia caused by changes in marine gateway configuration. *Global and Planetary Change*. *158*:57–71. https://doi.org/10.1016/j.gloplacha.2017.09.013

Palcu DV, Patina IS, Şandric I, Lazarev S, Vasiliev I, Stoica M, Krijgsman W. 2021. Late Miocene megalake regressions in Eurasia. *Scientific Reports*. *11*:11471. https://doi.org/10.1038/s41598-021-91001-z

Paolucci F, Buono MR, Fernández MS. 2025. Awakening Patagonia's sleeping sperm whale: A new description of the Early Miocene *Idiorophus patagonicus* (Odontoceti, Physeteroidea). *Papers in Palaeontology*. *11*(2):e70007. https://doi.org/10.1002/spp2.70007

Paolucci F, Buono MR, Fernández MS, Marx FG, Cuitiño JI. 2020. *Diaphorocetus poucheti* (Cetacea, Odontoceti, Physeteroidea) from Patagonia, Argentina: One of the earliest sperm whales. *Journal of Systematic Palaeontology*. *18*(4):335–355. https://doi.org/10.1080/14772019.2019.1605544

Paolucci F, Fernández MS, Buono MR, Cuitiño JI. 2021. '*Aulophyseter*' *rionegrensis* (Cetacea: Odontoceti: Physeteroidea) from the Miocene of Patagonia (Argentina): A reappraisal. *Zoological Journal of the Linnean Society*. *192*(4):1293–1322. https://doi.org/10.1093/zoolinnean/zlaa137

Papp K. 1905. *Heterodelphis leiodontus* nova forma aus den miocänen schichten des comitates sopron in ungarn. *Mitteilungen aus dem Jahrbuche der Königlich Ungarischen Geologischen Anstalt*. *14*(2):25–60.

Parham JF, Barron JA, Velez-Juarbe J. 2022. Middle and Late Miocene Marine Mammal Assemblages from the Monterey Formation of Orange County, California. In: *Understanding the Monterey Formation and Similar Biosiliceous Units across Space and Time*. Vol. 556. p. 229–241. https://doi.org/10.1130/2021.2556(10)

Parham JF, Papenfuss TJ, Buskirk JR, Parra-Olea G, Chen JY, Simison WB. 2015. *Trachemys ornata* or not *ornata*: Reassessment of a taxonomic revision for Mexican *Trachemys*. *Proceedings of the California Academy of Sciences, 4th series*. *62*(12):359–367.

Pecharová M, Prokop J. 2018. The morphology of mouthparts, wings and genitalia of Paleozoic insect families Protohymenidae and Scytohymenidae reveals new details and supposed function. *Arthropod Structure & Development*. *47*(1):117–129. https://doi.org/10.1016/j.asd.2017.11.006

Pecharová M, Prokop J, Ren D. 2015a. Early Pennsylvanian aykhalids from Xiaheyan, northern China and their palaeogeographical significance (Insecta: Megasecoptera). *Comptes Rendus Palevol*. *14*(8):613–624. https://doi.org/10.1016/j.crpv.2015.06.006

Pecharová M, Ren D, Prokop J. 2015b. A new palaeodictyopteroid (Megasecoptera: Brodiopteridae) from the Early Pennsylvanian of northern China reveals unique morphological traits and intra-specific variability. *Alcheringa: An Australasian Journal of Palaeontology*. *39*(2):236–249. https://doi.org/10.1080/03115518.2015.993299

Pecharová M, Sinitshenkova ND, Prokop J. 2020. On the morphology of the Late Paleozoic insect families Bardohymenidae and Aspidothoracidae (Palaeodictyopterida: Megasecoptera). *Arthropod Structure & Development*. *55*:100916. https://doi.org/10.1016/j.asd.2020.100916

Peppers RA. 1996. *Palynological Correlation of Major Pennsylvanian (Middle and Upper Carboniferous) Chronostratigraphic Boundaries in the Illinois and Other Coal Basins*. Geological Society of America Memoir 188. Boulder, Colorado: Geological Society of America. https://doi.org/10.1130/MEM188

Peredo CM, Pyenson ND. 2018. *Salishicetus meadi*, a new aetiocetid from the late Oligocene of Washington State and implications for feeding transitions in early mysticete evolution. *Royal Society Open Science*. *5*(4):172336. https://doi.org/10.1098/rsos.172336

Peredo CM, Pyenson ND, Marshall CD, Uhen MD. 2018a. Tooth loss precedes the origin of baleen in whales. *Current Biology*. *28*(24):3992–4000. https://doi.org/10.1016/j.cub.2018.10.047

Peredo CM, Uhen MD. 2016. A new basal chaeomysticete (Mammalia: Cetacea) from the late Oligocene Pysht formation of Washington, USA. *Papers in Palaeontology*. *2*(4):533–554. https://doi.org/10.1002/spp2.1051

Peredo CM, Uhen MD, Nelson MD. 2018b. A new kentriodontid (Cetacea: Odontoceti) from the early Miocene Astoria formation and a revision of the stem delphinidan family Kentriodontidae. *Journal of Vertebrate Paleontology*. *38*(2):e1411357. https://doi.org/10.1080/02724634.2017.1411357

Pereira AG, Sterli J, Moreira FRR, Schrago CG. 2017. Multilocus phylogeny and statistical biogeography clarify the evolutionary history of major lineages of turtles. *Molecular Phylogenetics and Evolution*. *113*:59–66. https://doi.org/10.1016/j.ympev.2017.05.008

Pérez Loinaze VS. 2009. New palynological data from the Malanzán Formation (Carboniferous), La Rioja Province, Argentina. *Ameghiniana*. *46*(3):385–408.

Peri E, Collareta A, Aringhieri G, Caramella D, Foresi LM, Bianucci G. 2022. A new physeteroid cetacean from the lower Miocene of southern Italy: CT imaging, retrodeformation, systematics and palaeobiology of a sperm whale from the Pietra leccese. *Bollettino della Società Paleontologica Italiana*. *61*(2):187–206. https://doi.org/10.4435/BSPI.2022.15

Peters SE, Antar MSM, Zalmout IS, Gingerich PD. 2009. Sequence stratigraphic control on preservation of late Eocene whales and other vertebrates at Wadi Al-Hitan, Egypt. *Palaios*. *24*(5):290–302. https://doi.org/10.2110/palo.2008.p08-080r

Petrulevičius JF, Gutiérrez PR. 2025. Oldest winged insects: First Megasecoptera from the early Carboniferous (Serpukhovian) of Argentina. *Palaeontology*. *68*(5):e70024. https://doi.org/10.1111/pala.70024

Pilleri G. 1985. The Miocene Cetacea of the Belluno sandstones (Eastern Southern Alps). *Memorie degli Istituti di Geologia e Mineralogia dell'Università di Padova*. *37*:1–250.

Pilleri G. 1987. *The Cetacea of the Italian Pliocene: With a Descriptive Catalogue of the Species in the Florence Museum of Paleontology*. Berne: Brain Anatomy Institute.

Pilleri G. 1988. *Eoplatanista taurinensis*, a new platanistid (cetacea) from the miocene of piedmont. In: *Contributions to the Paleontology of Some Tethyan Cetacea and Sirenia*. Ostermundigen: Brain Anatomy Institute. p. 7–18.

Pilleri G. 1989. Bemerkungen zur Arbeit 'Christian de Muizon: Le polyphýlétisme des Acrodelphidae'. In: *Contributions to the Paleontology of Some Tethyan Cetacea and Sirenia*. Ostermundigen: Brain Anatomy Institute. p. 115–123.

Pilleri G, Siber HJ. 1989. *Beiträge zur Paläontologie der Cetaceen Perus*. Ostermundigen: Hirnanatomisches Institut der Universität Bern.

Pinto ID. 1986. Carboniferous insects from Argentina. III. família Xenopteridae Pinto, nov. Ordo Megasecoptera. *Pesquisas*. *18*:23–29.

Pinto ID. 1992. New Upper Carboniferous palaeodictyopteran insect from Piedra Shotle Formation, Argentina. *Pesquisas em Geociências*. *19*(1):55–58.

Pinto ID. 1994. *Sphecorydaloides lucchesei*, a new Carboniferous megasecopteran insect from Argentina. *Pesquisas*. *21*:87–95.

Pinto ID. 1999. Taxonomy: South American Paleozoic Faunulae and Two New Insects. Chronological, Paleogeographical and Systematic Interpretation. In: *Proceedings of the First International Palaeoentomological Conference, Moscow, 1998*. p. 177–184.

Pinto ID, Adami-Rodrigues K. 1997. *Velisoptera taschi* nov. gen., nov. sp., a Carboniferous insect from Bajo de Velis, Argentina. *Pesquisas*. *24*(1-2):47–52.

Pinto ID, de Ornellas LP. 1978. Upper Carboniferous insects from Argentina. I – família Diaphanopteridae (Megasecopteroidea). *Pesquisas*. *10*:87–95.

Popov SV, Sychevskaya EK, Akhmet'ev MA, Zaporozhets NI. 2008. Stratigraphy of the Maikop group and Pteropoda beds in northern Azerbaijan. *Stratigraphy and Geological Correlation*. *16*(6):664–677. https://doi.org/10.1134/S0869593808060063

Post K, Bosselaers M. 2005. Late Pliocene occurrence of *Hemisyntrachelus* (Odontoceti, Delphinidae) in the southern North Sea. *Deinsea*. *11*:29–46.

Post K, Louwye S, Lambert O. 2017. *Scaldiporia vandokkumi*, a new pontoporiid (Mammalia, Cetacea, Odontoceti) from the Late Miocene to earliest Pliocene of the Westerschelde estuary (The Netherlands). *PeerJ*. *5*:e3991. https://doi.org/10.7717/peerj.3991

Poust AW, Boessenecker RW. 2018. Expanding the geographic and geochronologic range of early pinnipeds: New specimens of *Enaliarctos* from northern California and Oregon. *Acta Palaeontologica Polonica*. *63*(1):25–40. https://doi.org/10.4202/app.00399.2017

Preudhomme de Borre A. 1875. Note sur des empreintes d'insectes fossiles, découvertes dans les schistes houillers des environs de Mons. *Comptes-rendus des séances de la Société entomologique de Belgique*. *18*:XXXIX–XLIV.

Pritchard BG. 1939. On the discovery of a fossil whale in the older tertiaries of Torquay, Victoria. *The Victorian Naturalist*. *55*(9):151–159.

Prokop J, Kukalová-Peck J. 2017. New insects from the earliest Permian of Carrizo Arroyo (New Mexico, USA) bridging the gap between the Carboniferous and Permian entomofaunas. *Insect Systematics & Evolution*. *48*:493–511. https://doi.org/10.1163/1876312x-48022160

Prokop J, Nel A. 2004. A new genus and species of Homoiopteridae from the Upper Carboniferous of the Intra-Sudetic Basin, Czech Republic (Insecta: Palaeodictyoptera). *European Journal of Entomology*. *101*(4):583–589. https://doi.org/10.14411/eje.2004.082

Prokop J, Nel A. 2011. New Middle Permian palaeopteran insects from Lodève Basin in southern France (Ephemeroptera, Diaphanopterodea, Megasecoptera). *ZooKeys*. *130*:41–55. https://doi.org/10.3897/zookeys.130.1311

Prokop J, Nel A, Engel MS, Pecharová M, Hörnschemeyer T. 2016. New Carboniferous fossils of Spilapteridae enlighten postembryonic wing development in Palaeodictyoptera. *Systematic Entomology*. *41*(1):178–190. https://doi.org/10.1111/syen.12148

Prokop J, Pecharová M, Jarzembowski EA, Ross AJ. 2018. New palaeodictyopterans from the Late Carboniferous of the UK (Insecta: Palaeodictyopterida). *Earth and Environmental Science Transactions of the Royal Society of Edinburgh. 107*(2-3):99–107. https://doi.org/10.1017/s1755691017000408

Prokop J, Pecharová M, Nel A, Grey M, Hörnschemeyer T. 2017. A remarkable insect from the Pennsylvanian of the Joggins Formation in Nova Scotia, Canada: Insights into unusual venation of Brodiidae and nymphs of Megasecoptera. *Journal of Systematic Palaeontology. 15*:1051–1065. https://doi.org/10.1080/14772019.2017.1283364

Prokop J, Ren D. 2007. New significant fossil insects from the Upper Carboniferous of Ningxia in northern China (Palaeodictyoptera, Archaeorthoptera). *European Journal of Entomology. 104*(2):267–275. https://doi.org/10.14411/eje.2007.041

Prokop J, Roques P, Nel A. 2014. New non-holometabolous insects from the Pennsylvanian of the Avion locality in Pas-de-Calais, France (Insecta: ‘Exopterygota’). *Alcheringa. 38*(2):155–169. https://doi.org/10.1080/03115518.2014.848620

Prokop J, Rosová K, Leipner A, Sroka P. 2023. Thoracic and abdominal outgrowths in early pterygotes: A clue to the common ancestor of winged insects? *Communications Biology. 6*:1262. https://doi.org/10.1038/s42003-023-05568-6

Prokop J, Smith R, Jarzembowski EA, Nel A. 2006. New homoiopterids from the Late Carboniferous of England (Insecta: Palaeodictyoptera). *Comptes Rendus Palevol. 5*(7):867–873. https://doi.org/10.1016/j.crpv.2006.03.008

Prokop J, Tippeltová Z, Roques P, Nel A. 2013. A new genus and species of Breyeriidae and wings of immature stages from the Upper Carboniferous, Nord-Pas-de-Calais, France (Insecta: Palaeodictyoptera). *Insect Systematics & Evolution. 44*(2):117–128. https://doi.org/10.1163/1876312x-44032098

Prothero DR, Hoffman JM, Goedert JL. 2008. Paleomagnetism of the Oligocene and Miocene Lincoln Creek and Astoria formations, Knappton, Washington. In: Wang X, Barnes LG, editors. *Geology and Vertebrate Paleontology of Western and Southern North America: Contributions in Honor of David P. Whistler.* Vol. 41. Natural History Museum of Los Angeles County. p. 63–72.

Prothero DR, Streig A, Burns C. 2001. Magnetic Stratigraphy and Tectonic Rotation of the Upper Oligocene Pysht Formation, Clallam County, Washington. In: Prothero DR, editor. *Magnetic Stratigraphy of the Pacific Coast Cenozoic.* Vol. 91. p. 224–233.

Pruvost P. 1919. *Introduction à l’Étude du Terrain Houiller du Nord et du Pas-de-Calais. La Faune Continentale du Terrain Houiller du Nord de la France.* Mémoires pour servir à l’explication de la Carte géologique détaillée de la France. Paris: Imprimerie Nationale, 584.

Pruvost P. 1930. La faune continentale du terrain houiller de la Belgique. *Mémoires du Musée royal d'Histoire naturelle de Belgique. 44*:103–280.

Pyenson ND, Irmis RB, Lipps JH, Barnes LG, Mitchell ED, McLeod SA. 2009. Origin of a widespread marine bonebed deposited during the middle Miocene Climatic Optimum. *Geology. 37*(6):519–522. https://doi.org/10.1130/G25509A.1

Pyenson ND, Vélez-Juarbe J, Gutstein CS, Little H, Vigil D, O'Dea A. 2015. *Isthminia panamensis*, a new fossil inioid (Mammalia, Cetacea) from the Chagres formation of Panama and the evolution of "river dolphins" in the Americas. *PeerJ. 3*:e1227. https://doi.org/10.7717/peerj.1227

Racicot RA, Deméré TA, Beatty BL, Boessenecker RW. 2014. Unique feeding morphology in a new prognathous extinct porpoise from the Pliocene of California. *Current Biology. 24*(7):774–779. https://doi.org/10.1016/j.cub.2014.02.031

Racicot RA, Gearty W, Kohno N, Flynn JJ. 2016. Comparative anatomy of the bony labyrinth of extant and extinct porpoises (Cetacea: Phocoenidae). *Biological Journal of the Linnean Society. 119*(4):831–846. https://doi.org/10.1111/bij.12857

Ramassamy B. 2016. Description of a new long-snouted beaked whale from the Late Miocene of Denmark: Evolution of suction feeding and sexual dimorphism in the Ziphiidae (Cetacea: Odontoceti). *Zoological Journal of the Linnean Society. 178*(2):381–409. https://doi.org/10.1111/zoj.12418

Rasmussen ES, Dybkjær K, Piasecki S. 2010. Lithostratigraphy of the Upper Oligocene–Miocene succession of Denmark. *Geological Survey of Denmark and Greenland Bulletin. 22*:1–92. https://doi.org/10.34194/geusb.v22.4733

Rasnitsyn AP, Aristov DS, Gorochov AV, Rowland JM, Sinitshenkova ND. 2004. Important new insect fossils from Carrizo Arroyo and the Permo-Carboniferous faunal boundary. *Bulletin of the New Mexico Museum of Natural History and Science. 25*:215–246.

Reichenbach HGL. 1847. Systematisches. In: Carus CG, editor. *Resultate Geologischer, Anatomischer und Zoologischer Untersuchungen über das unter dem Namen Hydrarchos von Dr. A. C. Koch zuerst nach Europa gebrachte und in Dresden aufgestellte grosse fossile Skelett.* Dresden and Leipzig: Arnoldische Buchhandlung. p. 13–15.

Rensberger JM. 1969. A new iniid cetacean from the Miocene of California. *University of California Publications in Geological Sciences. 82*:1–43.

Richardson ES. 1956. *Pennsylvanian Invertebrates of the Mazon Creek Area, Illinois. Insects.* Vol. 12. Fieldiana: Geology 2. Chicago: Chicago Natural History Museum.

Riek EF. 1976a. Neosecoptera, a new insect suborder based on a specimen discovered in the Late Carboniferous of Tasmania. *Alcheringa: An Australasian Journal of Palaeontology. 1*(2):227–234. https://doi.org/10.1080/03115517608619072

Riek EF. 1976b. New Upper Permian insects from Natal, South Africa. *Annals of the Natal Museum*. *22*(3):755–789.

Robinson JH, Lee DE, Richards MD, White SEM, Fordyce RE. 2024. The fossil vertebrate primary type specimens in the collection of the University of Otago Department of Geology. *Journal of the Royal Society of New Zealand*. *54*(5):566–583. https://doi.org/10.1080/03036758.2024.2363436

Rochín-Bañaga H, Davis DW, Schwennicke T, Hernández-Cisneros AE. 2024. Age constraints on marine-fluvial deposits at Rancho La Palma, Baja California Sur, Mexico. *Boletín de la Sociedad Geológica Mexicana*. *76*(3):A040424. https://doi.org/10.18268/BSGM2024v76n3a040424

Rohdendorf BB. 1940a. Another genus of the family Scytohymenidae found in the Permian of the Urals. *Comptes Rendus (Doklady) de l'Académie des Sciences de l'URSS*. *26*(1):106–107.

Rohdendorf BB. 1940b. Body structure of Archodonata and position of this order. *Comptes Rendus (Doklady) de l'Académie des Sciences de l'URSS*. *26*(1):108–110.

Rohdendorf BB. 1947. Ob iskopaemykh nasekomykh iz Vorkutskogo ugol'nogo basseyna. *Doklady Akademii Nauk SSSR*. *57*:391–393.

Rohdendorf BB, Becker-Migdisova EE, Martynova OM, Sharov AG. 1961. *Paleozoiskie Nasekomye Kuznetskogo Basseina*. Vol. 85. Trudy Paleontologicheskogo Instituta, 1–705.

Rose FL, Weaver WG. 1966. Two new species of *Chrysemys* (= *Pseudemys*) from the Florida Pliocene. *Tulane Studies in Geology and Paleontology*. *5*(1):41–48.

Rosová K, Heising M, Leipner A, Prokop J. 2024. A new palaeodictyopteran *Haseneura jarmilae* gen. et sp. nov. from the Pennsylvanian of Piesberg reveals the structure of alinota. *Historical Biology*. *37*(9):2032–2039. https://doi.org/10.1080/08912963.2024.2407499

Rosová K, Pecharová M, Leipner A, Prokop J. 2022. New palaeodictyopterid insects from the Pennsylvanian of Piesberg reveal spectacular disparity in the patterns of the wing venation (Megasecoptera, Palaeodictyoptera). *Historical Biology*. *35*(1):152–162. https://doi.org/10.1080/08912963.2021.2025229

Roth F. 1978. *Mesocetus argillarius* sp.n. (Cetacea, Mysticeti) from Upper Miocene of Denmark, with remarks on the lower jaw and the echolocation system in whale phylogeny. *Zoologica Scripta*. *7*:63–79.

Rothausen K. 1968. Die systematische Stellung der europäischen Squalodontidae (Odontoceti, Mamm.) *Paläontologische Zeitschrift*. *42*(1-2):83–104. https://doi.org/10.1007/BF02987130

Rothausen K. 1971. *Cetotheriopsis tobieni* n. sp., der erste paläogene Bartenwal (Cetotheriidae, Mysticeti, Mamm.) nördlich des Tethysraumes. *Abhandlungen des Hessischen Landesamtes für Bodenforschung*. *60*:131–148.

Russell LS. 1968. A new cetacean from the Oligocene Sooke Formation of Vancouver Island, British Columbia. *Canadian Journal of Earth Sciences. 5*(4):929–933. https://doi.org/10.1139/e68-089

Rust BR, Gibling MR, Best MA, Dilles SJ, Masson AG. 1987. A sedimentological overview of the coal-bearing Morien Group (Pennsylvanian), Sydney Basin, Nova Scotia, Canada. *Canadian Journal of Earth Sciences. 24*(9):1869–1885. https://doi.org/10.1139/e87-177

Rýznar V. 2021. *Engisoptera ivanae* sp. n. (Megasecoptera: Engisopteridae) a new fossil insect from the Lower Permian of Obora, Czech Republic. *Humanitarian Space. 10*(1):77–82. https://doi.org/10.24412/2226-0773-2021-10-1-77-82

Sahni A, Mishra VP. 1972. A new species of *Protocetus* (Cetacea) from the middle Eocene of Kutch, western India. *Palaeontology. 15*(3):490–495.

Sahni A, Mishra VP. 1975. Lower Tertiary vertebrates from western India. *Monograph of the Palaeontological Society of India. 3*:1–48.

Salinas-Márquez FM, Barnes LG, Flores-Trujillo JG, Aranda-Manteca FJ. 2014. Una especie de delfín fósil (Cetacea; Delphinoidea; Kentriodontidae) del Mioceno medio de Baja California. *Boletín de la Sociedad Geológica Mexicana. 66*(1):145–164. https://doi.org/10.18268/bsgm2014v66n1a11

Sanchez-Posada C, Racicot RA, Ruf I, Krings M, Rössner GE. 2024. *Romaleodelphis pollerspoecki*, gen. et sp. nov., an archaic dolphin from the Central Paratethys (early Miocene, Austria). *Journal of Vertebrate Paleontology. 44*(2):e2401503. https://doi.org/10.1080/02724634.2024.2401503

Sanders AE, Barnes LG. 2002a. Paleontology of the Late Oligocene Ashley and Chandler Bridge Formations of South Carolina, 2: *Micromysticetus rothauseni*, a primitive cetotheriid mysticete (Mammalia, Cetacea). *Smithsonian Contributions to Paleobiology. 93*:271–293.

Sanders AE, Barnes LG. 2002b. Paleontology of the Late Oligocene Ashley and Chandler Bridge Formations of South Carolina, 3: *Eomysticetidae*, a New Family of Primitive Mysticetes (Mammalia: Cetacea). In: *Cenozoic Mammals of Land and Sea: Tributes to the Career of Clayton E. Ray.* Vol. 93. Washington, D.C.: Smithsonian Institution Press. p. 313–356.

Sanders AE, Geisler JH. 2015. A new basal odontocete from the upper Rupelian of South Carolina, U.S.A., with contributions to the systematics of *Xenorophus* and *Mirocetus* (Mammalia, Cetacea). *Journal of Vertebrate Paleontology. 35*(1):e890107. https://doi.org/10.1080/02724634.2014.890107

Santos AA, Hernández-Orúe A, Peñalver E, McLoughlin S, Diez JB, Nel A. 2025. First Iberian aspidothoracid megasecopteran insect and associated plants evidencing herbivory in a tropical Carboniferous forest from León, Spain. *Acta Palaeontologica Polonica. 70*(1):115–124. https://doi.org/10.4202/app.01203.2024

Santos AA, Wappler T, McLoughlin S. 2024. Earliest evidence of granivory from China (Shanxi Formation) points to seeds as a food source and nursing habitat for insects in the earliest Permian humid tropical forests of Cathaysia. *PLoS One*. *19*(10):e0311737. https://doi.org/10.1371/journal.pone.0311737

Sasaki T, Nikaido M, Wada S, Yamada TK, Cao Y, Hasegawa M, Okada N. 2006. *Balaenoptera omurai* is a newly discovered baleen whale that represents an ancient evolutionary lineage. *Molecular Phylogenetics and Evolution*. *41*(1):40–52. https://doi.org/10.1016/j.ympev.2006.03.032

Schachat SR, Goldstein PZ, DeSalle R, Bobo DM, Boyce CK, Payne JL, Labandeira CC. 2023. Illusion of flight? absence, evidence and the age of winged insects. *Biological Journal of the Linnean Society*. *138*(2):143–168. https://doi.org/10.1093/biolinnean/blac137

Scheirer AH, Magoon LB. 2008. Age, Distribution, and Stratigraphic Relationship of Rock Units in the San Joaquin Basin Province, California. In: Scheirer AH, editor. *Petroleum Systems and Geologic Assessment of Oil and Gas in the San Joaquin Basin Province, California*. Vol. 1713. U.S. Geological Survey. https://doi.org/10.3133/pp17135

Schneider JW, Lucas SG, Scholze F, Voigt S, Marchetti L, Klein H, Opluštil S, Werneburg R, Golubev VK, Barrick JE, et al. 2020. Late Palaeozoic–early Mesozoic continental biostratigraphy — Links to the Standard Global Chronostratigraphic Scale. *Palaeoworld*. *29*(2):186–238. https://doi.org/10.1016/j.palwor.2019.09.001

Schubnel T, Roques P, Oudard J, Garrouste R, Nel A. 2020. A new paleopteran genus and species from the late Carboniferous of Avion, France (Insecta: Palaeodictyopterida: ?Diaphanopterodea). *Palaeoentomology*. *3*(1):32–35. https://doi.org/10.11646/palaeoentomology.3.1.3

Scudder SH. 1867. Notice of fossil insects from the Devonian rocks of New Brunswick and of *Haplophlebium barnesii*. *Proceedings of the Boston Society of Natural History*. *11*:150–151.

Scudder SH. 1880. *The Devonian Insects of New Brunswick*. Anniversary Memoirs of the Boston Society of Natural History. Boston: Boston Society of Natural History, 41.

Scudder SH. 1881. Two new British Carboniferous insects, with remarks on those already known. *Geological Magazine*. *8*(7):293–300. https://doi.org/10.1017/s0016756800117285

Scudder SH. 1885. Dictyoneura and the allied insects of the Carboniferous epoch. *Proceedings of the American Academy of Arts and Sciences*. *20*:167–173. https://doi.org/10.2307/25138771

Scudder SH. 1893. Insect fauna of the Rhode Island coal field. *Bulletin of the United States Geological Survey*. *101*:1–27. https://doi.org/10.3133/b101

Seidel ME, Ernst CH. 2017. A systematic review of the turtle family Emydidae. *Vertebrate Zoology*. *67*(1):1–122. https://doi.org/10.3897/vz.67.e31535

Sharov AG, Sinitshenkova ND. 1977. New Palaeodictyoptera from the USSR. *Paleontological Journal*. *11*(1):44–59.

Shcherbakov DE, Makarkin VN, Aristov DS, Vasilenko DV. 2009. Permian insects from the Russky Island, South Primorye. *Russian Entomological Journal*. *18*(1):7–16.

Shen B, Shen S, Wu Q, Zhang S, Zhang B, Wang X, Hou Z, Yuan D, Zhang Y, Liu F, et al. 2022. Carboniferous and Permian integrative stratigraphy and timescale of North China Block. *Science China Earth Sciences*. *65*(6):983–1011. https://doi.org/10.1007/s11430-021-9909-9

Shipps BK, Peredo CM, Pyenson ND. 2019. *Borealodon osedax*, a new stem mysticete (Mammalia, Cetacea) from the Oligocene of Washington state and its implications for fossil whale-fall communities. *Royal Society Open Science*. *6*(7):182168. https://doi.org/10.1098/rsos.182168

Silantiev VV, Gutak YM, Tichomirowa M, Käßner A, Kulikova AV, Arbuzov SI, Nourgalieva NG, Karasev EV, Felker AS, Naumcheva MA, et al. 2025. Revisiting the Permian stratigraphy of the Kuznetsk Coal Basin (Siberia, Russia) using radioisotopic data: Sedimentology, biotic events, and palaeoclimate. *Minerals*. *15*(6):643. https://doi.org/10.3390/min15060643

Sinitshenkova ND. 1979. A new family of the Palaeodictyoptera from the Carboniferous of Siberia. *Paleontological Journal*. *13*(2):192–205.

Sinitshenkova ND. 1980. A revision of the order Permothemistida (Insecta). *Paleontological Journal*. *14*(4):97–112.

Sinitshenkova ND. 1981. A new species of the Tchirkovaeidae (Insecta, Dictyoneurida) from the Upper Carboniferous of the Tunguska basin. *Paleontological Journal*. *15*(1):121–123.

Sinitshenkova ND. 1992. Two New Insect Species (Insecta: Dictyoneurida = Palaeodictyoptera, Perlida = Plecoptera) from the Late Permian of South Mongolia. In: *Novye Taksony Iskopaemykh Bespozvonochnykh Mongolii*. Vol. 41. p. 98–100.

Sinitshenkova ND. 1993. A new insect family Aykhalidae from the Upper Palaeozoic of Yakutia-Sakha (Insecta: Mischopteridae = Megasecoptera). *Paleontological Journal*. *27*(1A):131–134.

Sinitshenkova ND. 2019. New species of Mischopterida = Megasecoptera (Insecta) from the Permian deposits of the Vorkuta Basin. *Paleontological Journal*. *53*(1):52–55. https://doi.org/10.1134/S003103011901009X

Sinitshenkova ND. 2020. New dictyoneuridans (Insecta: Dictyoneurida Handlirsch = Palaeodictyoptera Goldenberg) from the Permian deposits of the Vorkuta Basin. *Paleontological Journal*. *54*(2):126–131. https://doi.org/10.1134/S0031030120020136

Sinitshenkova ND. 2025. A new species *Dunbaria elkunensis* sp. nov. (Insecta: Palaeodictyoptera: Spilapteridae) from the Middle Permian Golyusherma locality, Udmurtia, Russia: The youngest known Spilapteridae. *Paleontological Journal*. *59*(2):168–174. https://doi.org/10.1134/S0031030125600076

Sinitshenkova ND, Aristov DS. 2013. The latest member of the order Mischopterida = Megasecoptera (Insecta) from the Upper Permian Isady locality. *Paleontological Journal*. *47*(7):704–705.

Sinitshenkova ND, Ponomareva GY, Aristov DS. 2020. A new megasecopteran species of the genus *Aspidothorax* Brongniart, 1893 (Insecta: Mischopterida = Megasecoptera: Aspidothoracidae) from the Middle Permian of Perm Region, Russia. *Paleontological Journal. 54*(3):273–278. https://doi.org/10.1134/S0031030120030144

Snoodijk F, Wallaard JJW, Schulp AS, Reumer JWF. 2024. A Miocene sperm whale (Cetacea, Physeteroidea) tooth from Liessel (Noord-Brabant, the Netherlands). *Netherlands Journal of Geosciences. 103*:e5. https://doi.org/10.1017/njg.2024.5

Solís-Añorve A, González-Barba G, Hernández-Rivera R. 2019. Description of a new toothed mysticete from the Late Oligocene of San Juan de La Costa, B.C.S., México. *Journal of South American Earth Sciences. 89*:337–346. https://doi.org/10.1016/j.jsames.2018.11.015

Solís-Añorve A, González-Barba G, Hernández-Rivera R, Schwennicke T. 2021. Late Miocene balaenopterid (Cetacea: Mysticeti) from Baja California Sur, Mexico. *Journal of South American Earth Sciences. 111*:103498. https://doi.org/10.1016/j.jsames.2021.103498

Spasskii PI. 1951. Ostatki tsetoteriev iz severnogo Kavkaza. *Izvestiya Akademii Nauk Azerbaidzhanskoi SSR. 2*:57–65.

Spinks PQ, Thomson RC, Lovely GA, Shaffer HB. 2009. Assessing what is needed to resolve a molecular phylogeny: Simulations and empirical data from emydid turtles. *BMC Evolutionary Biology. 9*:56. https://doi.org/10.1186/1471-2148-9-56

Spinks PQ, Thomson RC, McCartney-Melstad E, Shaffer HB. 2016. Phylogeny and temporal diversification of the New World pond turtles (Emydidae). *Molecular Phylogenetics and Evolution. 103*:85–97. https://doi.org/10.1016/j.ympev.2016.07.007

Steeman ME. 2009. A new baleen whale from the Late Miocene of Denmark and early mysticete hearing. *Palaeontology. 52*(5):1169–1190. https://doi.org/10.1111/j.1475-4983.2009.00893.x

Steeman ME. 2010. The extinct baleen whale fauna from the Miocene–Pliocene of Belgium and the diagnostic cetacean ear bones. *Journal of Systematic Palaeontology. 8*(1):63–80. https://doi.org/10.1080/14772011003594961

Strobel P. 1881. *Iconografia Comparata delle Ossa Fossili del Gabinetto di Storia Naturale dell'Università di Parma*. Parma: Libreria Editrice Luigi Battei.

Sugarman PJ, Miller KG, Owens JP, Feigenson MD. 1993. Strontium-isotope and sequence stratigraphy of the Miocene Kirkwood formation, southern New Jersey. *Geological Society of America Bulletin. 105*(4):423–436. https://doi.org/10.1130/0016-7606(1993)105<0423:SIASSO>2.3.CO;2

Tanaka Y, Abella J, Aguirre-Fernández G, Gregori M, Fordyce RE. 2017. A new tropical Oligocene dolphin from Montañita/Olón, Santa Elena, Ecuador. *PLoS One. 12*(12):e0188380. https://doi.org/10.1371/journal.pone.0188380

Tanaka Y, Ando T, Sawamura H. 2018. A new species of middle Miocene baleen whale from the Nupinai group, Hikatagawa formation of Hokkaido, Japan. *PeerJ*. *6*:e4934. https://doi.org/10.7717/peerj.4934

Tanaka Y, Fordyce RE. 2014. Fossil dolphin *Otekaikea marplesi* (latest Oligocene, New Zealand) expands the morphological and taxonomic diversity of Oligocene cetaceans. *PLoS One*. *9*(9):e107972. https://doi.org/10.1371/journal.pone.0107972

Tanaka Y, Fordyce RE. 2015a. A new Oligo-Miocene dolphin from New Zealand: *Otekaikea huata* expands diversity of the early Platanistoidea. *Palaeontologia Electronica*. *18*(1):18.1.5A. https://doi.org/10.26879/518

Tanaka Y, Fordyce RE. 2015b. Historically significant late oligocene dolphin *Microcetus hectori* Benham 1935: A new species of *Waipatia* (Platanistoidea). *Journal of the Royal Society of New Zealand*. *45*(3):135–150. https://doi.org/10.1080/03036758.2015.1016046

Tanaka Y, Fordyce RE. 2016. *Awamokoa tokarahi*, a new basal dolphin in the Platanistoidea (late Oligocene, New Zealand). *Journal of Systematic Palaeontology*. *15*(5):365–386. https://doi.org/10.1080/14772019.2016.1202339

Tanaka Y, Furusawa H, Kimura M. 2020. A new member of fossil balaenid (Mysticeti, Cetacea) from the early Pliocene of Hokkaido, Japan. *Royal Society Open Science*. *7*(4):192182. https://doi.org/10.1098/rsos.192182

Tanaka Y, Kimura T, Shinmura T, Ohira H, Furusawa H. 2025. A new member of a large and archaic balaenid from the Late Miocene of Sapporo, Hokkaido, Japan partly fills a gap of right whale evolution. *Palaeontologia Electronica*. *28*(2):a37. https://doi.org/10.26879/1549

Tanaka Y, Nakagawa R. 2024. A new platanistoid (Odontoceti: Squalodelphinidae) from the early Miocene of Japan. *Journal of Systematic Palaeontology*. *22*(1):e2378783. https://doi.org/10.1080/14772019.2024.2378783

Tanaka Y, Ortega M, Fordyce RE. 2026. A new squalodelphinid species with a domed supraorbital process from the Lower Miocene of New Zealand. *Journal of Vertebrate Paleontology*. *45*(5):e2641081. https://doi.org/10.1080/02724634.2026.2641081

Tanaka Y, Watanabe M. 2019. An early and new member of Balaenopteridae from the upper Miocene of Hokkaido, Japan. *Journal of Systematic Palaeontology*. *17*(16):1417–1431. https://doi.org/10.1080/14772019.2018.1532968

Tarasenko KK. 2014. New genera of baleen whales (Cetacea, Mammalia) from the Miocene of the northern Caucasus and Ciscaucasia: 3. *Zygiocetus* gen. nov. (middle Sarmatian, Adygea). *Paleontological Journal*. *48*(5):551–562. https://doi.org/10.1134/S0031030114050116

Tarasenko KK. 2024. Finds of skulls of juvenile *Kurdalagonus maicopicus* (Cetacea, Cetotheriidae) in the upper Miocene of the Republic of Adygea. *Doklady Biological Sciences*. *517*(1):73–76. https://doi.org/10.1134/S0012496624701011

Tarasenko KK, Lopatin AV. 2012a. New baleen whale genera (Cetacea, Mammalia) from the Miocene of the northern Caucasus and Ciscaucasia: 1. *Kurdalagonus* gen. nov. from the middle–late Sarmatian of Adygea. *Paleontological Journal*. *46*(5):531–542. https://doi.org/10.1134/S0031030112050115

Tarasenko KK, Lopatin AV. 2012b. New baleen whale genera (Cetacea, Mammalia) from the Miocene of the northern Caucasus and Ciscaucasia: 2. *Vampalus* gen. nov. from the middle–late Miocene of Chechnya and Krasnodar region. *Paleontological Journal*. *46*(6):620–629. https://doi.org/10.1134/S003103011206010X

Tasch P, Zimmerman JR. 1962. The *Asthenohymen–Delopterum* bed, a new Leonardian insect horizon in the Wellington Formation of Kansas and Oklahoma. *Journal of Paleontology*. *36*(6):1319–1333.

Teixeira C. 1941. Nouveaux insectes du Stéphanien portugais. *Boletim da Sociedade Geológica de Portugal*. *1*(1):13–31.

Teixeira C. 1946. Sur les premiers insectes fossiles rencontrés dans la mine de charbon de S. Pedro de Cova, au Portugal. *Boletim do Museu e Laboratório Mineralógico e Geológico da Universidade de Lisboa*. *13*:79–83.

Thewissen JGM, Hussain ST. 1998. Systematic review of the Pakicetidae, early and middle Eocene Cetacea (Mammalia) from Pakistan and India. *Bulletin of Carnegie Museum of Natural History*. *34*:220–238.

Thewissen JGM, Hussain ST. 2000. *Attockicetus praecursor*, a new remingtonocetid cetacean from marine Eocene sediments of Pakistan. *Journal of Mammalian Evolution*. *7*(3):133–146. https://doi.org/10.1023/A:1009458618729

Thewissen JGM, Hussain ST, Arif M. 1994. Fossil evidence for the origin of aquatic locomotion in archaeocete whales. *Science*. *263*(5144):210–212. https://doi.org/10.1126/science.263.5144.210

Thomson RC, Spinks PQ, Shaffer HB. 2021. A global phylogeny of turtles reveals a burst of climate-associated diversification on continental margins. *Proceedings of the National Academy of Sciences of the United States of America*. *118*(7):e2012215118. https://doi.org/10.1073/pnas.2012215118

Tillyard RJ. 1924. Kansas Permian insects. Part 3, the new Protohymenoptera. *American Journal of Science*. *208*(44):111–122. https://doi.org/10.2475/ajs.s5-8.44.111

Tillyard RJ. 1925. Kansas Permian insects. Part 4, the order Paleodictyoptera. *American Journal of Science*. *209*(52):328–335. https://doi.org/10.2475/ajs.s5-9.52.328

Tillyard RJ. 1926. Kansas Permian insects. Part 6, additions to the orders Protohymenoptera and Odonata. *American Journal of Science*. *211*(61):58–73. https://doi.org/10.2475/ajs.s5-11.61.58

Tillyard RJ. 1932. Kansas Permian insects. Part 14, the order Neuroptera. *American Journal of Science. 223*(133):1–30. https://doi.org/10.2475/ajs.s5-23.133.1

Tillyard RJ. 1937. Kansas Permian insects. Part 17, the order Megasecoptera and additions to the Palaeodictyoptera, Odonata, Protoperlaria, Copeognatha, and Neuroptera. *American Journal of Science. 233*(194):81–110. https://doi.org/10.2475/ajs.s5-33.194.81

Tosetto V, Damarco P, Daniello R, Pavia M, Carnevale G, Bisconti M. 2023. Cranial material of long-snouted dolphins (Cetacea, Odontoceti, Eurhinodelphinidae) from the Early Miocene of Rosignano Monferrato, Piedmont (NW Italy): Anatomy, paleoneurology, phylogenetic relationships and paleobiogeography. *Diversity. 15*(2):227. https://doi.org/10.3390/d15020227

Trevisan L. 1942. Una nuova specie di *Balaenula* pliocenica. *Palaeontographia Italica. 40*:1–13.

Trivedi AN, Satsangi PP. 1984. A new archaeocete (whale) from the Eocene of India. In: *27th International Geological Congress, Moscow*. Vol. 1. p. 322–323.

True FW. 1909. A new genus of fossil cetaceans from Santa Cruz Territory, Patagonia; and description of a mandible and vertebrae of *Prosqualodon*. *Smithsonian Miscellaneous Collections. 52*:441–455.

True FW. 1912. Description of a new fossil porpoise of the genus *Delphinodon* from the Miocene formation of Maryland. *Journal of the Academy of Natural Sciences of Philadelphia, 2nd Series. 15*:165–194.

Trümper S, Schneider JW, Nemyrovska T, Korn D, Linnemann U, Ren D, Béthoux O. 2020. Age and depositional environment of the Xiaheyan insect fauna, embedded in marine black shales (Early Pennsylvanian, China). *Palaeogeography, Palaeoclimatology, Palaeoecology. 538*:109444. https://doi.org/10.1016/j.palaeo.2019.109444

Tsai CH. 2017. A Miocene breeding ground of an extinct baleen whale (Cetacea: Mysticeti). *PeerJ. 5*:e3711. https://doi.org/10.7717/peerj.3711

Tsai CH. 2023. In search of the origin of crown mysticeti. *Journal of the Royal Society of New Zealand. 54*(5):682–695. https://doi.org/10.1080/03036758.2023.2249410

Tsai CH, Fordyce RE. 2015. The earliest gulp-feeding mysticete (Cetacea: Mysticeti) from the Oligocene of New Zealand. *Journal of Mammalian Evolution. 22*(4):535–560. https://doi.org/10.1007/s10914-015-9290-0

Tsai CH, Fordyce RE. 2016. Archaic baleen whale from the Kokoamu Greensand: Earbones distinguish a new late Oligocene mysticete (Cetacea: Mysticeti) from New Zealand. *Journal of the Royal Society of New Zealand. 46*(2):117–138. https://doi.org/10.1080/03036758.2016.1156552

Tsai CH, Fordyce RE. 2018. A new archaic baleen whale *Toipahautea waitaki* (early late Oligocene, New Zealand) and the origins of crown Mysticeti. *Royal Society Open Science. 5*(4):172453. https://doi.org/10.1098/rsos.172453

Tsai CH, Goedert JL, Boessenecker RW. 2024a. The oldest mysticete in the Northern Hemisphere. *Current Biology*. *34*(8):1794–1800. https://doi.org/10.1016/j.cub.2024.03.011

Tsai CH, Kimura T, Hasegawa Y. 2024b. Coexistence of Oligocene toothed and baleen-assisted mysticetes in the northwestern Pacific. *Fossil Record*. *27*(1):95–100. https://doi.org/10.3897/fr.27.e111567

Turtle Taxonomy Working Group. 2025. Turtles of the world: Annotated checklist and atlas of taxonomy, synonymy, distribution, and conservation status (10th ed.) *Chelonian Research Monographs*. *10*:1–575. https://doi.org/10.3854/crm.10.checklist.atlas.v10.2025

Uhen MD. 1998. New protocetid (Mammalia, Cetacea) from the late middle Eocene Cook Mountain formation of Louisiana. *Journal of Vertebrate Paleontology*. *18*(3):664–668. https://doi.org/10.1080/02724634.1998.10011093

Uhen MD. 1999. New species of protocetid archaeocete whale, *Eocetus wardii* (Mammalia: Cetacea) from the middle Eocene of North Carolina. *Journal of Paleontology*. *73*(3):512–528. https://doi.org/10.1017/S002233600002802X

Uhen MD. 2004. Form, function, and anatomy of *Dorudon atrox* (Mammalia, Cetacea): An archaeocete from the middle to late Eocene of Egypt. *University of Michigan Papers on Paleontology*. *34*:1–222.

Uhen MD. 2005. A new genus and species of archaeocete whale from Mississippi. *Southeastern Geology*. *43*(3):157–172.

Uhen MD. 2008. A new *Xenorophus*-like odontocete cetacean from the Oligocene of North Carolina and a discussion of the basal odontocete radiation. *Journal of Systematic Palaeontology*. *6*(4):433–452. https://doi.org/10.1017/S1477201908002472

Uhen MD. 2013. A review of North American Basilosauridae. *Bulletin of the Alabama Museum of Natural History*. *31*(2):1–45.

Uhen MD, Gingerich PD. 2001. New genus of dorudontine archaeocete (Cetacea) from the middle-to-late Eocene of South Carolina. *Marine Mammal Science*. *17*(1):1–34. https://doi.org/10.1111/j.1748-7692.2001.tb00979.x

Uhen MD, Pyenson ND, DeVries TJ, Urbina M, Renne PR. 2011. New middle Eocene whales from the Pisco Basin of Peru. *Journal of Paleontology*. *85*(5):955–969. https://doi.org/10.1666/10-162.1

Van Beneden PJ. 1859. Sur la découverte d'ossements fossiles faite à Saint-Nicolas. *Bulletin de l'Académie Royale des Sciences, des Lettres et des Beaux-Arts de Belgique, Series 2*. *8*:123–146.

Van Beneden PJ. 1865. Recherches sur les ossements provenant du Crag d'Anvers. les Squalodons. *Mémoires de l'Académie Royale des Sciences, des Lettres et des Beaux-Arts de Belgique*. *35*:1–85.

Van Beneden PJ. 1869. Sur un nouveau genre de Ziphioïde fossile (*Placoziphius*), trouvé à Edeghem, près d'Anvers. *Mémoires de l'Académie Royale des Sciences, des Lettres et des Beaux-Arts de Belgique*. *37*:1–12.

Van Beneden PJ. 1872. Les baleines fossiles d'Anvers. *Bulletin de l'Académie Royale des Sciences, des Lettres et des Beaux-Arts de Belgique, 2e série*. *34*(7):6–20.

Van Beneden PJ. 1877. Note sur un cachalot nain du Crag d'Anvers (*Physeterula dubusii*). *Bulletins de l'Académie Royale des Sciences, des Lettres et des Beaux-Arts de Belgique, Series 2*. *44*(12):851–856.

Van Beneden PJ. 1880. Les mysticètes à courts fanons des sables des environs d'Anvers. *Bulletins de l'Académie Royale des Sciences, des Lettres et des Beaux-Arts de Belgique*. *49*:11–27.

Van Beneden PJ. 1883. Sur quelques ossements de cétacés fossiles, recueillis dans des couches phosphatées entre l'Elbe et le Weser. *Bulletins de l'Académie Royale des Sciences, des Lettres et des Beaux-Arts de Belgique, Series 3*. *6*:27–33.

Van Beneden PJ. 1884. Une baleine fossile de Croatie, appartenant au genre *Mesocetus*. *Mémoires de l'Académie Royale des Sciences, des Lettres et des Beaux-Arts de Belgique*. *45*:1–29.

Van Beneden PJ, Gervais P. 1880. *Ostéographie des Cétacés Vivants et Fossiles, Comprenant la Description et l'Iconographie du Squelette et du Système Dentaire de Ces Animaux, Ainsi que des Documents Relatifs à Leur Histoire Naturelle*. Paris: Arthus Bertrand. https://doi.org/10.5962/bhl.title.50302

Vanden Broeck E. 1874. Esquisse géologique et paléontologique des dépôts pliocènes des environs d'Anvers. *Mémoires de la Société Malacologique de Belgique*. *9*:87–374.

van Dijk DE, Geertsema H. 1999. Permian insects from the Beaufort Group of Natal, South Africa. *Annals of the Natal Museum*. *40*:137–171.

van Vliet HJ, Bosselaers M, Vahldiek BW, Paymans T, Verheijen I. 2020. Eocene cetaceans from the Helmstedt region, Germany, with some remarks on *Platyosphys*, *Basilotritus* and *Pachycetus*. *Cainozoic Research*. *20*(1):121–148.

Vélez-Juarbe J. 2017. A new stem odontocete from the late Oligocene Pysht formation in Washington state, U.S.A. *Journal of Vertebrate Paleontology*. *37*(5):e1366916. https://doi.org/10.1080/02724634.2017.1366916

Vélez-Juarbe J. 2023. New heterodont odontocetes from the Oligocene Pysht Formation in Washington State, U.S.A., and a reevaluation of Simocetidae (Cetacea, Odontoceti). *PeerJ*. *11*:e15576. https://doi.org/10.7717/peerj.15576

Vélez-Juarbe J, Pyenson ND. 2012. *Bohaskaia monodontoides*, a new monodontid (Cetacea, Odontoceti, Delphinoidea) from the Pliocene of the western North Atlantic ocean. *Journal of Vertebrate Paleontology*. *32*(2):476–484. https://doi.org/10.1080/02724634.2012.641705

Vélez-Juarbe J, Wood AR, De Gracia C, Hendy AJW. 2015. Evolutionary patterns among living and fossil kogiid sperm whales: Evidence from the Neogene of Central America. *PLoS One*. *10*(4):e0123909. https://doi.org/10.1371/journal.pone.0123909

Viglino M, Buono MR, Fordyce RE, Cuitiño JI, Fitzgerald EMG. 2018a. Anatomy and phylogeny of the large shark-toothed dolphin *Phoberodon arctirostris* Cabrera, 1926 (Cetacea: Odontoceti) from the early Miocene of Patagonia (Argentina). *Zoological Journal of the Linnean Society*. *185*(2):511–542. https://doi.org/10.1093/zoolinnean/zly053

Viglino M, Buono MR, Gutstein CS, Cozzuol MA, Cuitiño JI. 2018b. A new dolphin from the early Miocene of Patagonia, Argentina: Insights into the evolution of Platanistoidea in the Southern Hemisphere. *Acta Palaeontologica Polonica*. *63*(2):261–277. https://doi.org/10.4202/app.00441.2017

Viglino M, Gaetán CM, Cuitiño JI, Buono MR. 2021. First toothless platanistoid from the Early Miocene of Patagonia: The golden age of diversification of the Odontoceti. *Journal of Mammalian Evolution*. *28*(2):337–358. https://doi.org/10.1007/s10914-020-09505-w

Vlachos E. 2018. A review of the fossil record of North American turtles of the clade Pan-Testudinoidea. *Bulletin of the Peabody Museum of Natural History*. *59*(1):3–94. https://doi.org/10.3374/014.059.0101

Vogt RC, Bulté G, Iverson JB. 2018. *Graptemys geographica* (LeSueur 1817) – Northern map turtle, common map turtle. *Chelonian Research Monographs*. *5*:104.1–104.18. https://doi.org/10.3854/crm.5.104.geographica.v1.2018

von Meyer H. 1841. *Arionius servatus*, ein den Delphinen verwandtes Meeres-Säugethier aus der Molasse von Baltringen in Württemberg. *Neues Jahrbuch für Mineralogie, Geognosie, Geologie und Petrefaktenkunde*. *1841*:315–331.

von Meyer H. 1843. Mittheilungen an Professor Bronn gerichtet. *Neues Jahrbuch für Mineralogie, Geognosie, Geologie und Petrefaktenkunde*. *1843*:698–704.

von Meyer H. 1849. Briefliche mittheilung. *Neues Jahrbuch für Mineralogie, Geognosie, Geologie und Petrefaktenkunde*:547–550.

Wade BS, Aljahdali MH, Mufrreh YA, Memesh AM, AlSoubhi SA, Zalmout IS. 2021. Upper Eocene planktonic foraminifera from northern Saudi Arabia: Implications for stratigraphic ranges. *Journal of Micropalaeontology*. *40*(2):145–161. https://doi.org/10.5194/jm-40-145-2021

Wagner RH, Paz Castro M. 2011. Compositional changes in a mid-Stephanian (Kasimovian) flora in relation to alluvial-plain deposits derived from westward-receding mountains and bordered by the Paleotethys: La Magdalena Coalfield, northwestern Spain. *Palaios. 26*(1):33–54. https://doi.org/10.2110/palo.2010.p10-004r

Wang YH, Engel MS, Rafael JA, Wu HY, Rédei D, Xie Q, Wang G, Liu XG, Bu WJ. 2016. Fossil record of stem groups employed in evaluating the chronogram of insects (Arthropoda: Hexapoda). *Scientific Reports. 6*:38939. https://doi.org/10.1038/srep38939

Waqas M, Weppe R, Lazzari V, Blondel C, Marivaux L, Charruault AL, Tarry M, Euriat A, Lebrun R, Rana RS, et al. 2026. A basal representative of Cetacea from the Eocene of India. *Nature Ecology & Evolution*:1195–1203. https://doi.org/10.1038/s41559-026-03055-9

Waterlot G. 1934. *Bassin houiller de la Sarre et de la Lorraine. II. Faune fossile. Étude de la faune continentale du terrain houiller sarro-lorrain.* Études des Gîtes Minéraux de la France. Lille: Imprimerie L. Danel, 317.

Weaver WG, Robertson JS. 1967. A re-evaluation of fossil turtles of the *Chrysemys scripta* group. *Tulane Studies in Geology and Paleontology. 5*(2):570–588.

Weems RE, Bybell LM, Edwards LE, Lewis WC, Self-Trail JM. 2016. Stratigraphic revision of the Cooper Group and the Chandler Bridge and Edisto formations in the coastal plain of South Carolina. *South Carolina Geology. 49*:1–24.

Weems RE, Edwards LE. 2007. The age and provenance of "*Eschrichtius*" *cephalus* Cope (Mammalia: Cetacea). *Journal of Vertebrate Paleontology. 27*(3):752–756. https://doi.org/10.1671/0272-4634(2007)27[752:TAAPOE]2.0.CO;2

Weems RE, Edwards LE, Landacre B. 2017. Geology and Biostratigraphy of the Potomac River Cliffs at Stratford Hall, Westmoreland County, Virginia. In: *From the Blue Ridge to the Beach: Geological Field Excursions across Virginia.* Vol. 47. Geological Society of America. p. 125–152. https://doi.org/10.1130/2017.0047(05)

Weems RE, Edwards LE, Osborne J, Alford AA. 2011. An occurrence of the protocetid whale "*Eocetus*" *wardii* in the middle Eocene Piney Point formation of Virginia. *Journal of Paleontology. 85*(2):271–278. https://doi.org/10.1666/10-083.1

West RM. 1980. Middle Eocene large mammal assemblage with Tethyan affinities, Ganda Kas region, Pakistan. *Journal of Paleontology. 54*(3):508–533.

Westgate JW, Whitmore F. C. J. 2002. *Balaena ricei*, a new species of bowhead whale from the Yorktown formation (Pliocene) of Hampton, Virginia. *Smithsonian Contributions to Paleobiology. 93*:295–312. https://doi.org/10.5479/si.00810266.93.295

Whitmore FC, Barnes LG. 2008. The Herpetocetinae, a new subfamily of extinct baleen whales (Mammalia, Cetacea, Cetotheriidae). *Virginia Museum of Natural History Special Publication. 14*:141–180.

Whitmore F. C. J, Kaltenbach JA. 2008. Neogene Cetacea of the Lee Creek Phosphate Mine, North Carolina. *Virginia Museum of Natural History Special Publication. 14*:181–269.

Wilson LE. 1935. Miocene marine mammals from the Bakersfield region, California. *Bulletin of the Peabody Museum of Natural History. 4*:1–143.

Wilson LE. 1973. A delphinid (Mammalia, Cetacea) from the Miocene of Palos Verdes Hills, California. *University of California Publications in Geological Sciences. 103*:1–34.

Woodward H. 1876. On a remarkable fossil orthopterous insect from the Coal-measures of Scotland. *Quarterly Journal of the Geological Society of London. 32*(1-4):60–65. https://doi.org/10.1144/gsl.jgs.1876.032.01-04.09

Woodward H. 1887. On the discovery of the larval stage of a cockroach, *Etoblattina peachii*, H. Woodw., from the Coal-Measures of Kilmaurs, Ayrshire. *Geological Magazine. 4*(10):433–435. https://doi.org/10.1017/s0016756800190284

Woodward H. 1913. *Rochdalia parkeri*, a new branchiopod crustacean from the Middle Coal-Measures of Sparth, Rochdale. *Geological Magazine. 10*(8):352–356. https://doi.org/10.1017/s0016756800127025

Wootton RJ. 1972. Nymphs of Palaeodictyoptera (Insecta) from the Westphalian of England. *Palaeontology. 15*(4):662–675.

Wu Q, Ramezani J, Zhang H, Wang J, Zeng F, Zhang Y, Liu F, Chen J, Cai Y, Hou Z, et al. 2021. High-precision U–Pb age constraints on the Permian floral turnovers, paleoclimate change, and tectonics of the North China block. *Geology. 49*(6):677–681. https://doi.org/10.1130/g48051.1

Xu ZQ, Yang N, Oyama N, Xu Y, Dong L, Ren D, Béthoux O. 2025. A new, large Permian Parelmoidae (Insecta, Megasecopteromorpha, Diaphanopterodea) found in Shanxi Province (China). *Geodiversitas. 47*(18):705–712. https://doi.org/10.5252/geodiversitas2025v47a18

Yang N, Cui Y, Xu Z, Xu Y, Ren D, Béthoux O. 2024. The first Permian Diaphanopterodea (Insecta, Megasecopteromorpha) from China. *Fossil Record. 27*(2):247–258. https://doi.org/10.3897/fr.27.128892

Yang N, Ren D, Béthoux O. 2020. The 'Megasecoptera–Diaphanopterodea' twilight zone epitomized by a new insect from Xiaheyan (Early Pennsylvanian; China). *Alcheringa: An Australasian Journal of Palaeontology. 44*(2):273–278. https://doi.org/10.1080/03115518.2020.1737228

Yang N, Ren D, Béthoux O. 2023. Small-sized Megasecopteromorpha (Insecta: Palaeoptera) occurred also in the Pennsylvanian. *Historical Biology. 37*(1):1–6. https://doi.org/10.1080/08912963.2023.2282675

Yoshida K, Kimura T, Hasegawa Y. 2003. New cetothere (Cetacea: Mysticeti) from the Miocene Chichibumachi Group, Japan. *Bulletin of the Saitama Museum of Natural History. 20*:1–10.

Zalessky G. 1937a. Études des insectes permiens du bassin de la Sylva et problèmes de l'évolution dans la classe des insectes. *Problemy Paleontologii. 2–3*:601–607.

Zalessky G. 1937b. Nouveaux représentants des protohyménoptères et des archodonates provenant du terrain permien du bassin de la Rivière Kama et leurs liens de parenté. *Annales de la Société entomologique de France. 106*(2):101–114. https://doi.org/10.1080/21686351.1937.12278973

Zalessky GM. 1939. Permskie nasekomye basseyna reki Sylvy i voprosy evolyutsii v klasse nasekomykh. III. *Problemy Paleontologii. 5*:33–91.

Zalessky GM. 1946. Sur le nouveau représentant de l'ordre des Palaeodictyoptères du Paléozoïque supérieur du bassin de Kousnetzk. *Bulletin de la Société des Naturalistes de Moscou, Série Biologique. 51*:58–62.

Zalessky GM. 1956. Novye predstaviteli otryada Protohymenoptera iz permskikh otlozhenii Urala. *Doklady Akademii Nauk SSSR. 110*:1089–1092.

Zalessky MD. 1931. Observations sur les insectes trouvés dans les dépôts à charbon du bassin de Kousnetzk. *Bulletin de la Société géologique de France, 5e série. 1*:209–218.

Zeigler CV, Chan GL, Barnes LG. 1997. A new late Miocene balaenopterid whale (Cetacea: Mysticeti), *Parabalaenoptera baulinensis*, (new genus and species) from the Santa Cruz Mudstone, Point Reyes Peninsula, California. *Proceedings of the California Academy of Sciences. 50*(4):115–138.

Zessin W. 2006. Zwei neue Insektenreste (Megasecoptera, Odonatoptera) aus dem Westfalium D (Oberkarbon) des Piesberges bei Osnabrück, Deutschland. *Virgo, Mitteilungsblatt des Entomologischen Vereins Mecklenburg. 9*(1):37–45.

Zimmt JB, Kidwell SM, Lockwood R, Thirlwall MF. 2022. Strontium isotope stratigraphy reveals 100 ky-scale condensation, beveling, and internal shingling of transgressive shell beds in the Maryland Miocene. *Palaios. 37*(10):553–573. https://doi.org/10.2110/palo.2022.004

Zurano JP, Magalhães FM, Asato AE, Silva G, Bidau CJ, Mesquita DO, Costa GC. 2019. Cetartiodactyla: Updating a time-calibrated molecular phylogeny. *Molecular Phylogenetics and Evolution. 133*:256–262. https://doi.org/10.1016/j.ympev.2018.12.015
